# Ki-67 promotes sequential stages of tumourigenesis by enabling cellular plasticity

**DOI:** 10.1101/712380

**Authors:** K. Mrouj, P. Singh, M. Sobecki, G. Dubra, E. Al Ghoul, A. Aznar, S. Prieto, N. Pirot, F. Bernex, B. Bordignon, C. Hassen-Khodja, M. Pouzolles, V. Zimmerman, V. Dardalhon, L. Krasinska, D. Fisher

**Affiliations:** Institut de Génétique Moléculaire de Montpellier (IGMM), University of Montpellier, CNRS, Montpellier, France; Equipe Labellisée LIGUE 2018, Ligue Nationale Contre le Cancer, Paris, France; Institute for Stem Cell Biology and Regenerative Medicine, Stanford University School of Medicine, Stanford, CA 94305, USA; Columbia University Medical Center, New York, USA; University of Zurich, Institute of Anatomy, Zurich, Switzerland; Institut de Recherche en Cancérologie de Montpellier (IRCM), Inserm, Montpellier, France; Réseau d’Histologie Expérimentale de Montpellier (RHEM), CNRS, Montpellier, France; Montpellier Ressources en Imagerie (MRI), CNRS, Montpellier, France

## Abstract

Recent studies have shown that the cell proliferation antigen Ki-67 is not required for cell proliferation. Here, we demonstrate that Ki-67 enables implementation of transcriptional programmes conferring cellular plasticity, and is required for each step of tumour initiation, growth and metastasis. Ki-67 knockout causes global transcriptome remodelling, which, in mammary carcinoma cells, inhibits the epithelial-mesenchymal transition in a polycomb-repressive complex 2-dependent manner. This results in suppression of stem cell characteristics and sensitisation to various drug classes. Cancer cells lacking Ki-67 proliferate normally *in vivo*, but tumour growth is inhibited due to disrupted angiogenesis, and metastasis is abrogated. Finally, mice lacking Ki-67 are resistant to chemical or genetic induction of intestinal tumourigenesis. Thus, Ki-67, which is expressed in all proliferating cancer cells, confers the plasticity required for different steps of carcinogenesis.

## INTRODUCTION

Ki-67 is a nuclear protein expressed only in proliferating vertebrate cells, a property underlying its widespread use in oncology as a biomarker (1). It is routinely used in histopathology to grade tumours; there are also indications for its application as prognostic marker (2). However, its cellular functions are not well understood, and whether it is involved in tumourigenesis remains unclear.

For a long time, Ki-67 was thought to be required for cell proliferation (3–8) and early work suggested that it promotes ribosomal RNA transcription (4, 9). However, recent genetic studies have shown that despite promoting formation of the perichromosomal layer of mitotic chromosomes (9–12), it is not required at all for cell proliferation (10–13). It is also dispensable for ribosomal RNA synthesis and processing, and mice lacking Ki-67 develop and age normally (11). In addition, Ki-67 is not overexpressed in cancers; rather, Ki-67 expression is controlled by cell cycle regulators, including CDKs, the activating subunit of the ubiquitin ligase APC/C-CDH1, which is required for destruction of mitotic cyclins, and the cell cycle transcription factor B-Myb (11, 14, 15). These recent studies raise the question of whether Ki-67 is required for tumourigenesis, which has not been addressed genetically.

Despite not being essential for cell proliferation, Ki-67 might be important in carcinogenesis for other reasons. We previously identified over 50 Ki-67-interacting proteins that are involved in transcription and chromatin regulation. We also found that Ki-67 organises heterochromatin: in NIH3T3 cells with TALEN-mediated biallelic disruption of the Ki-67 gene, or human cancer cells with stable knockdown of Ki-67, localisation of repressive histone marks H3K9me3 and H4K20me3 is dispersed, the intensity of heterochromatin is reduced, and association between centromeres and nucleoli is disrupted (11). Conversely, overexpression of Ki-67 caused ectopic heterochromatin formation (11). Involvement of Ki-67 in heterochromatin organisation was corroborated by a study showing that Ki-67 is required to maintain heterochromatin marks at inactive X-chromosomes in non-transformed cells (16). Heterochromatin is a phenotypic marker of multiple cancers (17), suggesting that it might facilitate carcinogenesis. We thus tested whether and how Ki-67 is required for different steps of carcinogenesis.

## RESULTS

### Loss of Ki-67 causes global transcriptome changes and deregulates pathways involved in cancer

We first investigated the impact of the reorganisation of chromatin observed upon loss of Ki-67 on gene expression. RNA-seq analysis of NIH/3T3 cells revealed wide-ranging transcriptome changes upon Ki-67 knockout, with 2558 genes significantly deregulated in independent clones of *Mki67*^*-/-*^ NIH/3T3 cells (q < 0.05) (Fig. 1A; Table S1). This level of transcriptome alteration suggests a global effect rather than a direct involvement of Ki-67 in controlling specific transcription factors, which is consistent with our previous finding that Ki-67 interacts with many general chromatin regulators (11). If so, then Ki-67 knockout should also globally affect the transcriptome of cancer cells, with possible consequences for tumourigenicity. To investigate this, we used the 4T1 mouse mammary carcinoma model, which is derived from BALB/c mice. This cell line mimics human triple-negative breast cancer and spontaneously metastasises to distant organs (18, 19). Consistent with our recent results showing that Ki-67 is not required for cell proliferation (11), 4T1 cell proliferation rates were unaffected by CRISPR-Cas9-mediated *Mki67* gene knockout (Fig. S1A-D). In 4T1 cells, as in NIH/3T3 cells, *Mki67* knockout caused genome-scale gene expression alterations: 4979 genes were deregulated, of which 1239 and 585 genes were >2-fold down-regulated and up-regulated, respectively (Fig. 1B,C; Table S2). There was little overlap in the deregulated genes between *Mki67*^*-/-*^ 4T1 (epithelial) and NIH/3T3 (mesenchymal) cells (Fig. 1C, Tables S1, S2) in accordance with our hypothesis that, by maintaining chromatin organisation, Ki-67 ensures global gene expression programmes in different cell types rather than directly regulating specific genes.

**Figure 1.**
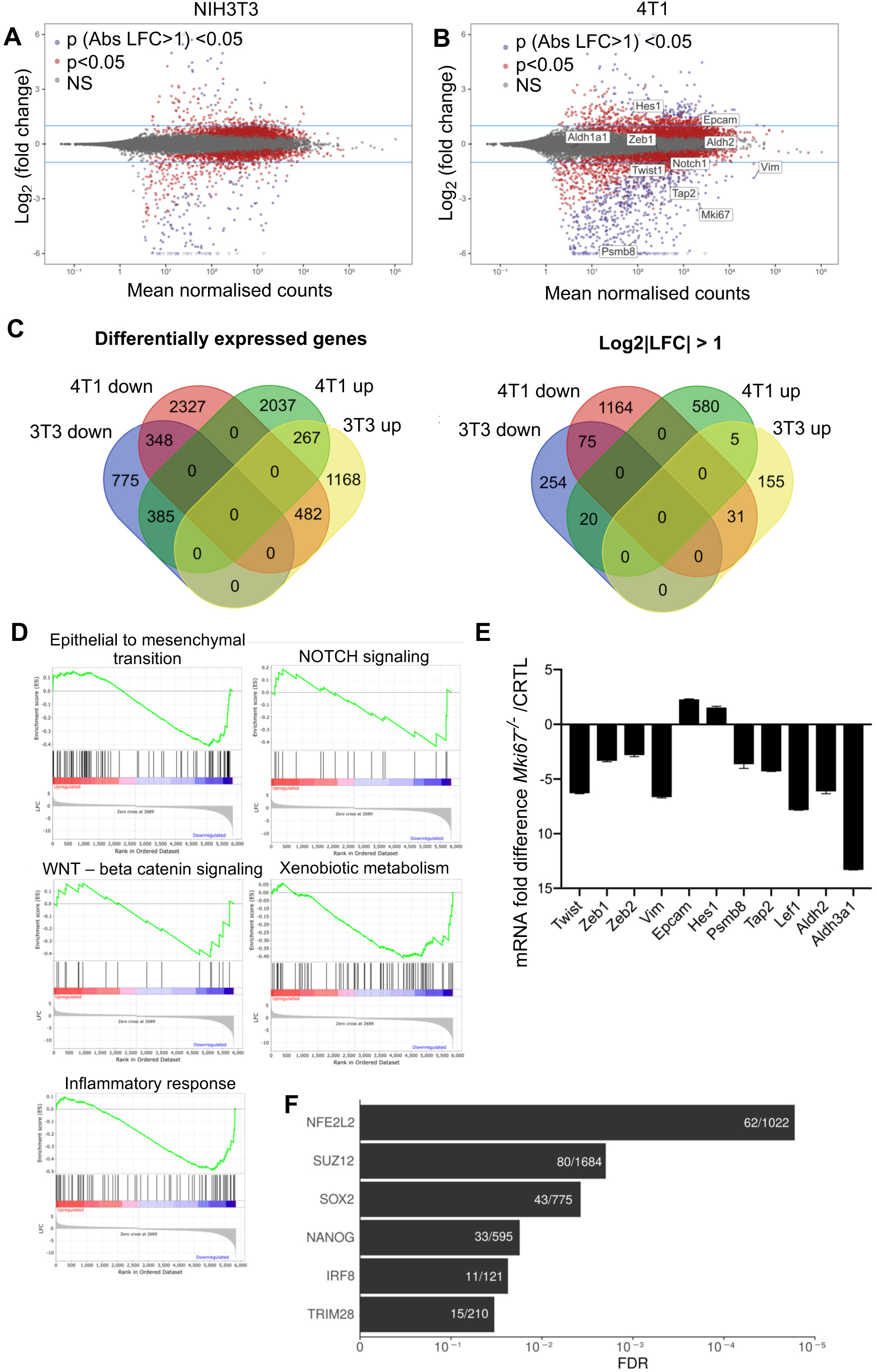
Ki-67 ablation deregulates global gene expression programs. (A, B) Dot plot analysis of differentially expressed genes (DEGs) in *Mki67*^-/-^ NIH/3T3 (A) and 4T1 (B) cells. Red dots: DEGs with p-value < 0.05; purple dots: log_2_ fold change, LFC >1 or <-1, p-value < 0.05; grey dots: not significant, NS. (C) Venn diagrams of DEGs in NIH/3T3 and 4T1 *Mki67*^-/-^ cells under condition of p-value < 0.05 (left), and p-value (LFC >1 or <-1) < 0.05 (right). (D) Gene Set Enrichment Analysis (GSEA) of highly down-regulated genes in 4T1 *Mki67*^-/-^ cells. (E) Quantitative RT-PCR analysis of differentially expressed genes in 4T1 *Mki67*^-/-^ cells; fold change in expression ± SD is shown. (F) Gene set enrichment analysis of ENCODE and ChEA Consensus transcription factors from ChIP-X gene sets in *Mki67*^*-/-*^ 4T1 cells. FDR, false discovery rate adjusted p-values.

We then investigated whether the extensive transcriptome changes seen in cancer cells upon Ki-67 knockout deregulate transcription programmes involved in tumourigenesis. Pathway analysis of highly deregulated genes revealed upregulation of Notch targets and downregulation of genes involved in the inflammatory response, Wnt pathway, the epithelial-mesenchymal transition (EMT) and xenobiotic metabolism (Fig. 1D), which we validated by qRT-PCR (Fig. 1E). Downregulated genes were enriched in targets of nuclear factor erythroid 2-related factor 2 (NFE2L2), one of the major orchestrators of responses to oxidative stress; polycomb-repression complex 2 (PRC2), which mediates H3K27me3 and is a well-characterised regulator of the EMT (20, 21); the pluripotency factors Nanog and Sox2; and interferon regulatory factor 8 (IRF8) (Fig. 1F). All of these pathways have previously been implicated in tumourigenesis.

### Ki-67 enables the epithelial-mesenchymal transition and cell plasticity

We next investigated the biological relevance of these transcriptome alterations. Importantly, although the Notch pathway is oncogenic in T cell acute lymphoid leukaemia, it can act as a tumour suppressor in specific cellular contexts (22), can block Wnt signalling (23, 24) (a driver of tumourigenesis, cell stemness and the EMT) and induce drug resistance (25). We confirmed Notch pathway upregulation at the protein level (Fig. 2A). We also confirmed that Ki-67 knockout 4T1 cells had lost expression of the mesenchymal marker vimentin but upregulated E-cadherin, and had a more epithelial morphology (Fig. 2B,C), indicative of a loss of EMT. Since the EMT is closely associated with a stem-like state (26–28), we next analysed aldehyde dehydrogenase activity, a marker of stem and progenitor cells (29, 30). This was strongly reduced in 4T1 Ki-67 knockout cells (Fig. 2D), suggesting a loss of stem-like character. In agreement, the ability to form spheroids in the absence of adhesion to a surface (26, 31) was also strongly reduced (Fig. 2E).

**Figure 2.**
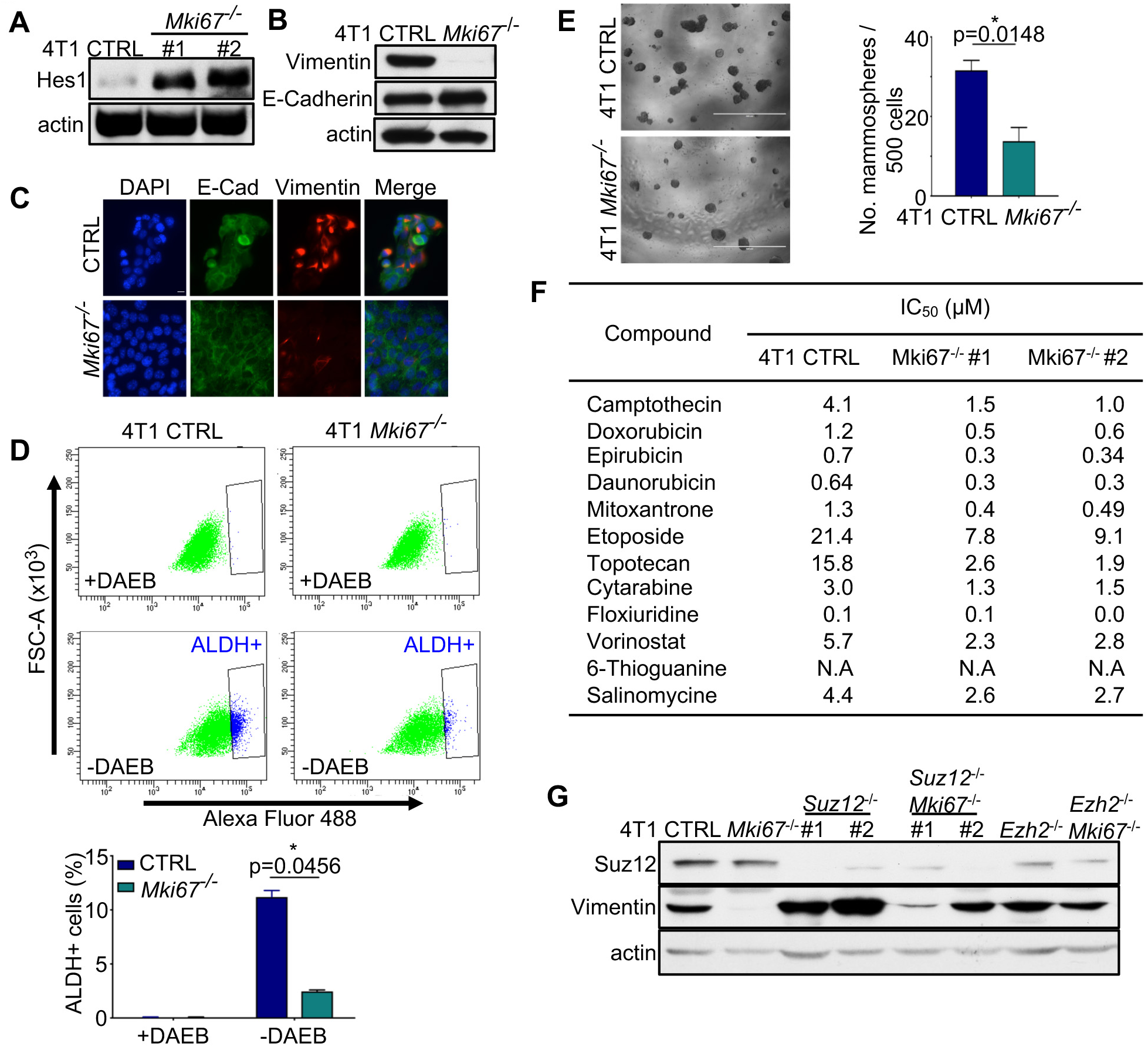
Ki-67 enables implementation of an EMT via PRC2. (A) Immunoblot analysis of Hes1 expression in 4T1 CTRL and *Mki67*^-/-^ cells. (B, C) Immunoblot (B) and immunofluorescence (C) (scale bar, 10µm) analysis of vimentin and E-cadherin expression in 4T1 CTRL and *Mki67*^-/-^ cells. (D) Aldehyde dehydrogenase 1 activity measured using a flow-cytometry assay in 4T1 CTRL and *Mki67*^-/-^ cells. DAEB, inhibitor of ALDH was used as a negative control. Top, flow cytometry profiles. Bottom, quantification of ALDH+ cells. Error bars, SEM (n=2 independent analyses). (E) Mammosphere formation assay of 4T1 CTRL or *Mki67*^-/-^ cells. Representative images (left; scale bar, 400 µm) and quantification (right; error bars, SEM, n=10) after 7 days. (F) IC_50_ of 4T1 CTRL or *Mki67*^-/-^ cells to indicated compounds, derived from dose-response experiments. (G) Western-blot analysis of Suz12 and Vimentin levels in 4T1 cells, CRTL, and single or double knockouts for *Mki67, Suz12* and *Ezh2*. Actin serves as loading control.

In addition to stem-like characteristics, the EMT has also been associated with resistance to chemotherapeutic drugs (32). We noticed that 26 genes involved in drug metabolism were downregulated in Ki-67 knockout 4T1 cells, while only one was upregulated, suggesting that Ki-67 expression might determine sensitivity to chemotherapeutic drugs (Fig. 1D, Fig. S2A). To test this, we performed an automated gene-drug screen using the Prestwick chemical library, composed of 1283 FDA-approved small molecules. We also included salinomycin, a positive control found to target cancer stem cells (33), and 6-thioguanine, which was originally used to isolate 4T1 cells (18). Control 4T1 cells were sensitive to 102 drugs at 10µM concentration, while the two *Mki67*^*-/-*^ clones were sensitive to 99 and 98 respectively, with 82 hits common to the three cell lines (Fig. S2B; Supporting Data S1). This suggests that Ki-67 loss does not qualitatively alter the drug-sensitivity profiles. We next determined the IC_50_ of 10 common hits used in cancer therapy. Importantly, *Mki67*^*-/-*^ cells were markedly more sensitive to all the molecules tested (Fig. 2F). As such, by supporting expression of xenobiotic metabolism genes, Ki-67 provides cancer cells with a degree of protection against drugs used in therapies.

We previously found that in human osteosarcoma U2OS cells, Ki-67 binds to the essential PRC2 component SUZ12 (11). To test whether the repression of the EMT in Ki-67 knockout depends on PRC2, we disrupted *Suz12* or *Ezh2* in WT and *Mki67*^*-/-*^ 4T1 cells using CRISPR-Cas9 (Fig. 2G). This partially rescued expression of vimentin (Fig. 2G), suggesting that loss of Ki-67 releases a pool of PRC2 which can then repress target genes, including EMT genes.

### Ki-67 is essential for normal mammary tumour growth and metastasis

Hybrid EMT states are important for tumourigenesis (34–36), and most evidence supports an essential role for the EMT in metastasis (37, 38). To determine the physiological consequences of Ki-67 loss for tumourigenesis, we engrafted WT and Ki-67 knockout 4T1 cells orthotopically into mouse mammary fat pads. Since Ki-67 knockout caused alteration of inflammatory response genes (Fig. 1D), we used athymic-nude and NOD/SCID mice, allowing us to assess roles of Ki-67 in tumour growth and metastasis without confounding effects of a potentially altered immune response. RNA-seq of early-stage tumours from WT and Ki-67 mutant 4T1 cells grafted into nude mice showed that Ki-67-dependent transcriptome changes were generally preserved *in vivo* (Fig. 3A, Table S3). As assessed by PCNA staining, cell proliferation *in vivo* was unaffected by loss of Ki-67, but tumours from *Mki67*^*-/-*^ 4T1 cells grew significantly slower than from Ki-67-expressing 4T1 cells in both types of immunodeficient mice (Fig. 3B-D). Control 4T1 cells metastasised within 3-4 weeks in nude mice, but metastases were largely absent at this point in mice bearing *Mki67*^*-/-*^ 4T1 cells (Fig. 3E). We quantified micrometastases by dissociating lung tissue after 3 weeks and growing cells in the presence of 6-thioguanine, to which 4T1 cells are resistant. The number of metastatic cells that formed colonies was reduced nearly 100-fold in Ki-67 knockouts (Fig. 3F). This points to an essential requirement for Ki-67 in metastasis, in accordance with its role in enabling the EMT.

**Figure 3.**
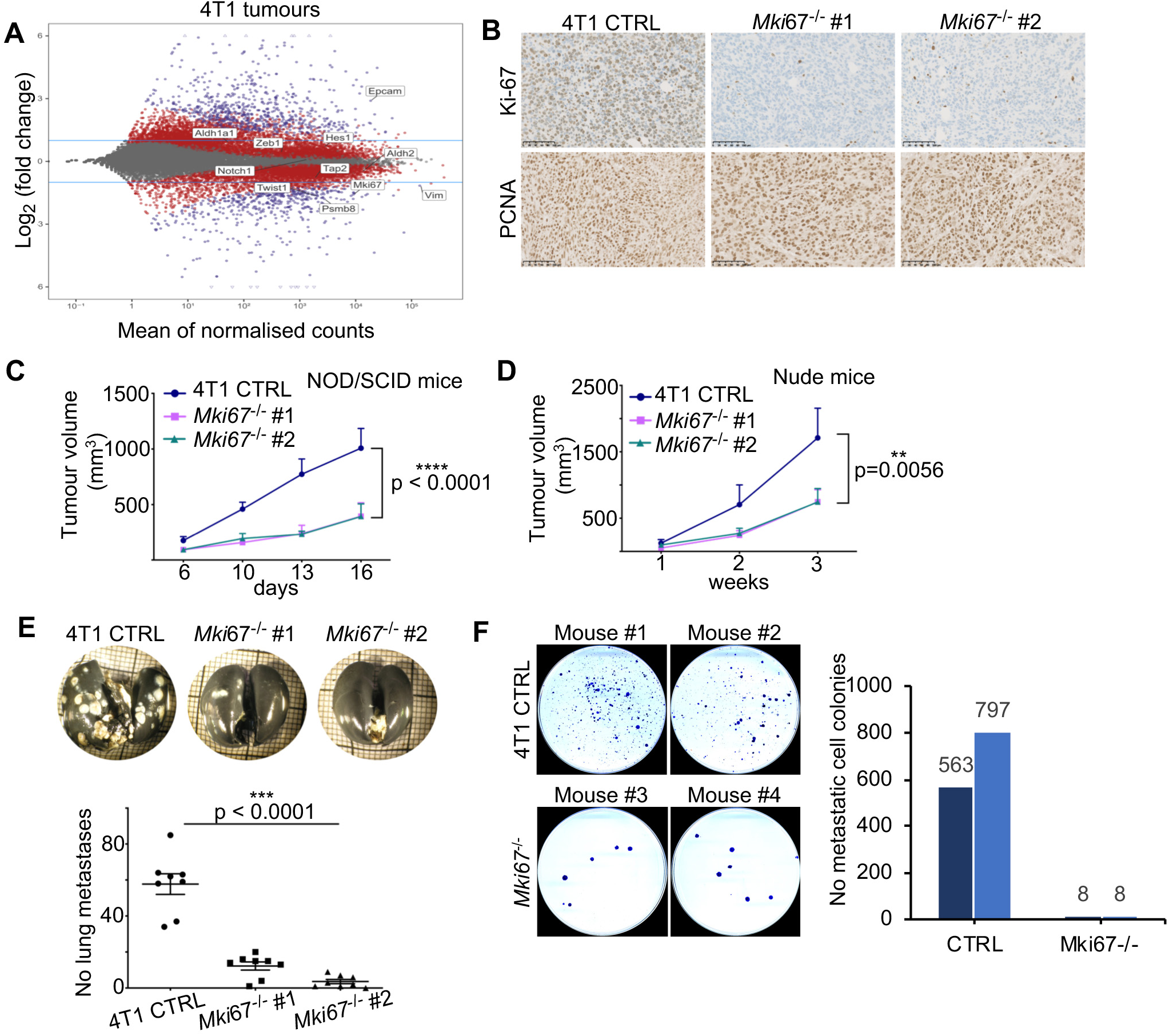
Ki-67 promotes tumour growth and metastasis. (A) Dot plot analysis of differentially expressed genes (DEGs) in tumours derived from grafting 4T1 *Mki67*^-/-^ cells or WT cells into nude mice. (B) IHC staining for Ki-67 and PCNA in 4T1 CTRL or *Mki67*^-/-^ tumours. Scale bar, 100µm. (C) 4T1 CTRL or *Mki67*^-/-^ orthotopic xenografts in NOD/SCID mice. Tumour growth was monitored for 3 weeks. Error bars, SEM (n=6 mice). (D) 4T1 CTRL or *Mki67*^-/-^ orthotopic xenografts in athymic nude mice, tumour burden measured over 3 weeks; error bars, SEM (n=8 mice). (E) Top: representative images of lungs stained to visualise metastases (white nodules); scale bar, 10mm. Below: quantification of lung metastases in each group. Error bars, SEM. (F) Lung tissue from mice injected via tail vein with 4T1 CTRL or *Mki67*^-/-^ cells (2 per condition) was dissociated and resulting cells were maintained in the presence of 6-thioguanine, to select for 4T1 cells. Left, crystal violet staining of resulting colonies. Right, quantification.

To assess the generality of requirement for Ki-67 in tumour growth in human cancer cells, we disrupted the *MKI67* gene by CRISPR-Cas9 in human MDA-MB-231 triple negative breast cancer cells (Fig. S3A-D). As expected, *MKI67*^*-/-*^ cells proliferated normally *in vitro* (Fig. S3C,D). In contrast, tumours from xenografts in nude mice grew much slower than controls (Fig. 4A). Cell proliferation was normal, as expected (Fig. 4B), but, while a regular thin fibrovascular capillary network was present in WT tumours, this was disorganised and irregular in *MKI67*^*-/-*^ tumours, suggesting defective angiogenesis (Fig. 4C). In accordance, mutant tumours showed increased oedema of lymphatic vessels, indicative of slow blood flow, and a larger necrotic area. Thus, Ki-67 is required to promote tumour growth not through sustaining cell proliferation, but by inducing angiogenesis.

**Figure 4.**
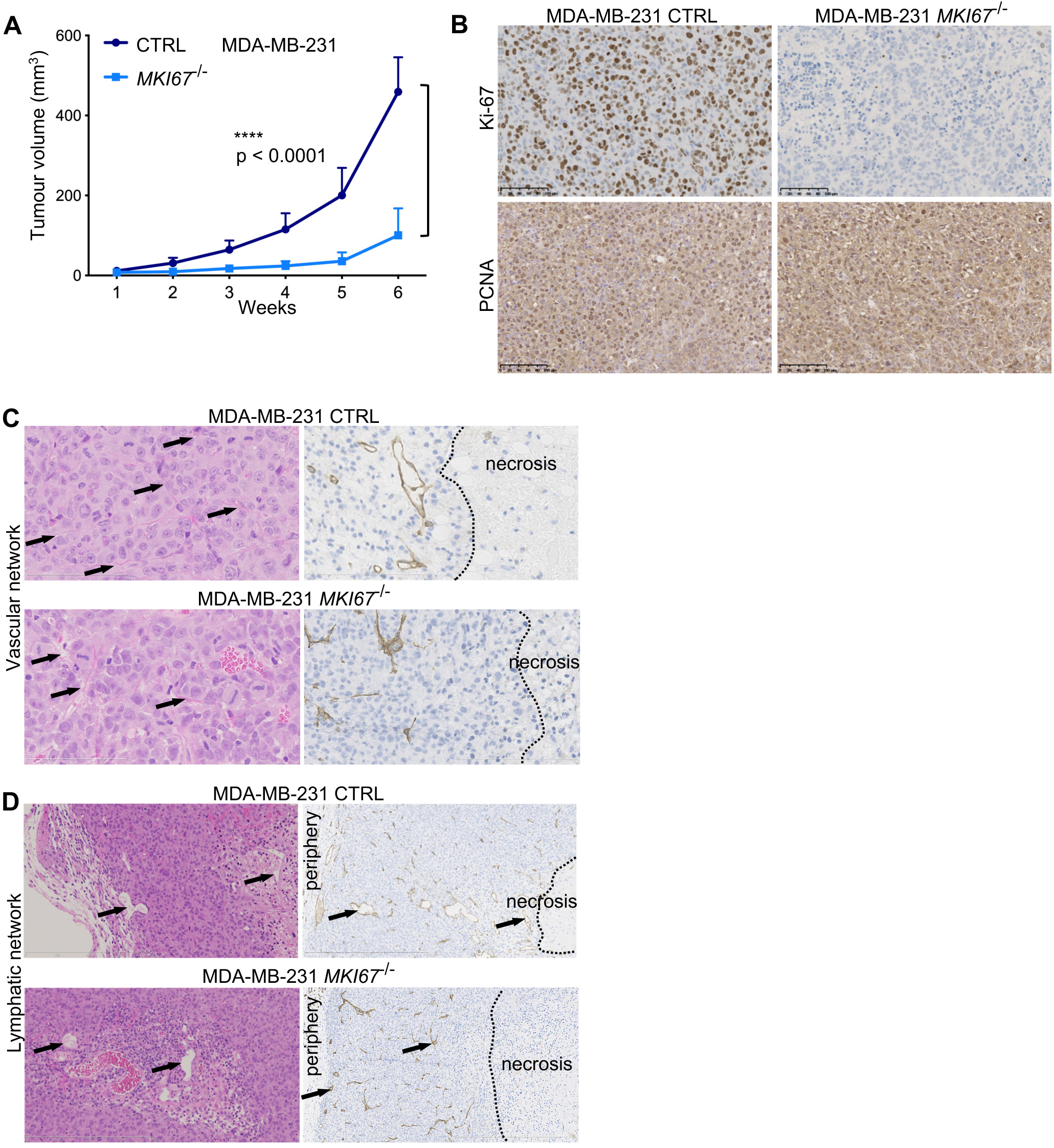
Ki-67 sustains tumour growth by enabling angiogenesis. (A-D) Xenografts of MDA-MB-231 CTRL or Ki-67 KO cells in mammary fat pads of nude mice. (A) Tumour burden. Error bars, SEM (n=8 mice). (B) IHC images of tumours stained for Ki-67 and PCNA. Scale bar, 100µm. (C, D) IHC analysis of vascular (C) and lymphatic (D) networks (arrows). Left, H&E; scale bar, 400µm. Right, CD31 staining; scale bar, 800µm. Areas of necrosis are indicated.

### Mice lacking Ki-67 are resistant to intestinal tumourigenesis

The above experiments show that Ki-67 is required for normal tumour growth and metastasis. To see whether Ki-67 knockout protects against tumourigenesis *in vivo*, we used a germline TALEN-disruption of Ki-67 (*Mki67*^*2ntΔ/2ntΔ*^) that we generated (11). We first employed a mouse model of colon carcinogenesis induced chemically by azoxymethane / dextran sodium sulphate (AOM-DSS) treatment (39). We confirmed that DSS alone triggered similar inflammatory responses in mice of all genotypes (Fig. S4A,B). Furthermore, *Mki67*^*2ntΔ/2ntΔ*^ mice have no apparent defects in immune cell proliferation and differentiation in response to haemolytic anaemia induced by phenylhydrazine treatment (Fig. S5). As expected, AOM-DSS efficiently induced colon tumours within 16 weeks in wild-type and *Mki67*^*2ntΔ/+*^ mice; however, no macroscopic lesions were observed in *Mki67*^*2ntΔ/2ntΔ*^ mice (Fig. 5A, B). This suggests that Ki-67 is specifically required for initiation of tumourigenesis. To confirm this, we used a genetic model of intestinal tumourigenesis. We crossed *Mki67*^*2ntΔ/2ntΔ*^ mice with *Apc*^*Δ14/+*^ mice, which rapidly develop tumours in the intestine due to loss of the second allele of the *Apc* tumour suppressor gene (40). While, as expected, *Apc*^*Δ14/+*^ *Mki67*^*2ntΔ/+*^ mice formed multiple intestinal tumours, tumour burden was strongly reduced in *Apc*^*Δ14/+*^ *Mki67*^*2ntΔ/2ntΔ*^ mice (Fig. 5C, D). Thus, Ki-67 is required for efficient initiation of tumourigenesis induced by chemical mutagenesis or loss of a tumour suppressor *in vivo*. We hypothesised that the altered transcriptomes that accompany loss of Ki-67 in non-transformed cells might hinder their oncogenic transformation. While testing this *in vivo* is not feasible, we found that overexpression of oncogenic *RAS* efficiently transformed wild-type NIH/3T3 cells, but not *Mki67*^*-/-*^ NIH/3T3 cells (Fig. S6).

**Figure 5.**
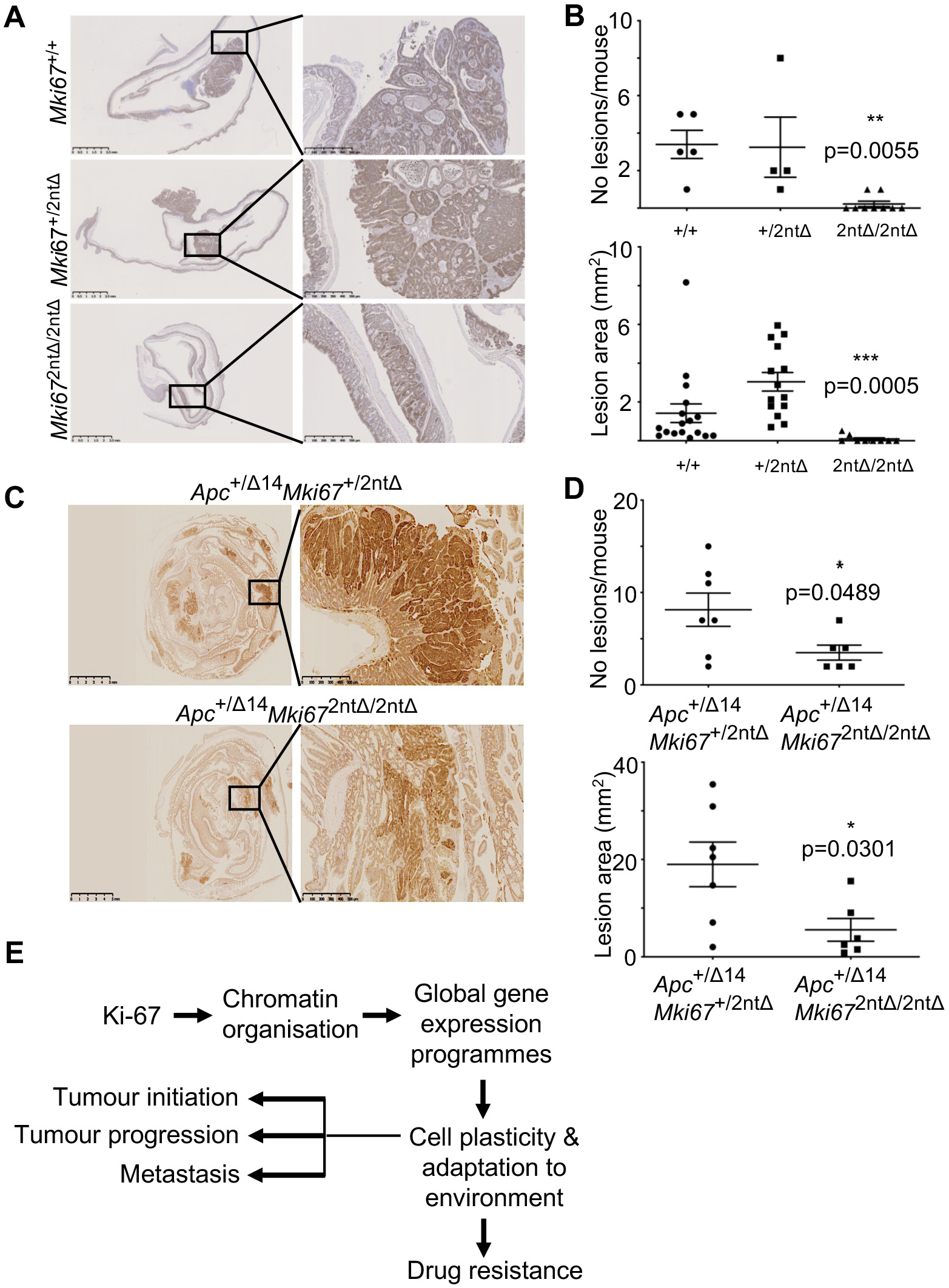
Germline disruption of Ki-67 protects mice against intestinal tumourigenesis. (A, C) IHC staining of β-catenin in whole intestines from 6-7-month old *Apc*^+/Δ14^*Mki67*^+/2ntΔ^ (A) and Apc^+/Δ14^*Mki67*^2ntΔ/2ntΔ^ (C) mice. Scale bar, 5mm. Insets show accumulation of β-catenin in nuclei. Scale bar, 500µm. (B, D) Quantification of the number (top) and total area (bottom) of neoplastic lesions. Error bars, SEM (n= 7 *Apc*^+/Δ14^*Mki67*^+/2ntΔ^ mice; n= 6 Apc^+/Δ14^*Mki67*^2ntΔ/2ntΔ^ mice). (E) Model of Ki-67 mechanism in tumour progression and cancer cell dissemination. See text for details.

## DISCUSSION

The above results show that Ki-67 is not required for cancer cell proliferation *in vivo*, but its expression critically influences all steps of tumourigenesis tested, including initiation, progression, metastasis and drug sensitivity. We present evidence that this is because Ki-67 is required to implement transcriptional programmes needed for tumour cell plasticity (Fig. 5E). This is indicated by the failure to generate intestinal tumours in Ki-67-mutant mice, the inability of cancer cells mutated for Ki-67 to induce angiogenesis, the extensive presence of EMT characteristics in epithelial cancer cells but their absence in cells lacking Ki-67, and the increased sensitivity of the latter to drugs.

A recent study in the human colorectal cancer cell line DLD-1 showed that Ki-67 knockout reduces the number of cells expressing the antigen CD133, commonly assumed to be a marker of cancer stem cells (CSC) (13). However, in intestinal crypts, CD133 expression is not specific to stem cells, and CD133-negative cancer cells were equally capable of sustaining tumourigenesis in a long-term serial transplantation model (41). Furthermore, the concept that CSCs are rare pre-existing populations of cells with hard-wired CSC properties, a model which emerged from xenograft experiments using sorted populations of hematopoietic stem cells, no longer appears valid, at least for solid tumours (42). Lineage tracing experiments in the intestine showed that tumours arise from stem cells which divide rapidly, make up around 10% of the cell in intestinal crypts, and can be regenerated from non-stem cells (42). A more recent concept is that epithelial cells are phenotypically plastic, in a manner dependent on EMT-inducing transcription factors (43), with no such fixed entity as CSC or non-CSC. Emerging evidence suggests that cells can reside in various phenotypic stages along the EMT-spectrum with concurrent expression of epithelial and mesenchymal traits, and that such hybrid states are important for carcinogenesis (34–36). Our data showing that ALDH activity, which is a bona fide marker for intestinal stem cells and breast and colon CSC (29, 30), is expressed by a significant fraction (≈10%) of WT epithelial cancer cells but far fewer (<3%) Ki-67 knockout cells, are consistent with this model.

Clues to the mechanism of action of Ki-67 are suggested by the fact that it is an intrinsically disordered protein (IDP) and behaves as a surfactant for mitotic chromatin (10). We propose that these properties may also allow Ki-67 to regulate interphase chromatin, since, as we showed previously, heterochromatin is more dispersed in cells lacking Ki-67 (11). Importantly, heterochromatin is a liquid-like compartment whose surface-tension properties depend partly on HP1 proteins (44, 45), which bind directly to Ki-67 (46). Such liquid-like compartments rely on proteins that can induce phase-separation, which often include IDPs (47). Due to their conformational flexibility, IDPs can also act as hubs to bind multiple protein partners (48), and we previously identified many general chromatin regulators interacting with Ki-67, including components of PRC1 and PRC2, REST, NuRD, NIF1, NuA4, MLL, SET1, NoRC and NCOA6 complexes (11). Disrupting one such interactor, the PRC2 complex, partially, but not fully, rescued EMT traits in Ki-67 mutant cancer cells, consistent with the idea that Ki-67 acts through multiple chromatin regulatory complexes. We speculate on several ways in which Ki-67 might generally confer transcriptome plasticity to cells in response to environmental conditions. First, it might directly sequester and release general chromatin regulators, providing a pool of these factors for rapid cellular adaptation. In this model, environmental cues could alter Ki-67 interactions through changes in post-translational modifications, such as phosphorylation, releasing sequestered factors. Alternatively, Ki-67 could influence structural properties of membrane-less nuclear compartments, such as perinucleolar heterochromatin, which is a marker of malignancy in multiple solid tumours that correlates with aggressiveness (17). Modification of physico-chemical properties of Ki-67 would result in spatial reorganisation of chromatin, that could alter binding of general transcriptional co-factors with knockon effects on the transcriptome. Further studies will be required to test these models and define the exact biochemical mechanisms by which Ki-67 confers transcriptional plasticity to cells.

In conclusion, Ki-67, which is universally expressed in proliferating cells, enables multiple steps in carcinogenesis in different cancer types by promoting global transcriptional programmes. Therapeutic targeting of Ki-67 itself will likely be challenging, since it is an IDP with no intrinsic enzyme activity. However, it could be of therapeutic benefit to inhibit its effectors in control of cellular plasticity.

## MATERIALS AND METHODS

### Cell lines and mice

4T1 cells were provided by Robert Hipskind (IGMM, Montpellier); MDA-MB-231 cells were obtained from SIRIC, Montpellier. NIH 3T3, 4T1, MDA-MB-231 cells were grown in Dulbecco modified Eagle medium (DMEM - high glucose, pyruvate, GlutaMAX – Gibco^®^ Life LifeTechnologies) supplemented with 10% foetal bovine serum (SIGMA or DUTSCHER). Cells were grown under standard conditions at 37°C in humidified incubator containing 5% CO_2_.

6-8 weeks-old female athymic nude (Hsd:Athymic Foxn1^nu^/Foxn1^+^), and NOD.SCID (NOD.CB17-Prkdc^scid^/NCrHsd) mice were purchased from Envigo. C57BL/6 Apc^Δ14^ mice were provided by Philippe Jay (IGF, Montpellier).

### Animal studies

All animal experiments were performed in accordance with international ethics standards and were subjected to approval by the Animal Experimentation Ethics Committee of Languedoc Roussillon (n°APAFIS#8637-2017012311188975v3).

### Antibodies and plasmids

Antibodies: Ki-67 (clone SP6; Abcam), cyclin A2 (6E6; Novocastra), PCNA (ab18197; Abcam), β-catenin (BD610154; BD-Bioscience), Ras (G12V Mutant Specific D2H12, #14412; CST), actin (A2066; Sigma), vimentin (D21H3, #5741; CST), E-Cadherin (24E10, #3195; CST), Suz12 (D39F6, #3737; CST), H3K27me3 (#39155, Active Motif).

Lentiviral Vectors used: LentiCRISPRv2 (Addgene #52961), pMD2.G (Addgene #12259), psPAX2 (Addgene #12260). pHIV-Luc-ZsGreen (Addgene #39196) was used to generate bicistronic lentivirus that expresses both Luciferase and Zsgreen.

Retroviral vectors used: pBabe-puro (Addgene #1764), pBabe-puro H-Ras^G12V^ (Addgene #1768). gag/pol (retroviral packaging) and Maloney (envelope) were gift from Leatitia Linares (IRCM, Montpellier).

pUC57-U6-sgRNA (#51132, Addgene); Cas9-VP12 (#72247, Addgene), modified by adding T2A-GFP at the C-terminal end (gift from Edouard Bertrand, IGMM).

### CRISPR/Cas9-mediated genome editing

The sgRNAs targeting mouse *Mki67* exon 3 or human *MKI67* exon 6 and non-targeting control sequences were previously designed (49), and cloning of the target sequence into the LentiCRISPRV2 lentiviral vector was conducted as described (54). Lentiviruses encoding the sgRNA targeting sequences were produced. Transduced cells (4T1 and MDA-MB-231) expressing CRISPR/Cas9 were selected using puromycin. Resistant cells were isolated and seeded as single cell clones in 96 well-plates.

For Suz12 and Ezh2 knockout, sgRNA targeting sequences (49) were synthesised with BpiI sticky ends and cloned into the pUC57-U6-sgRNA vector. Cas9-VP12-T2A-GFP and sgRNA vectors were transfected into 4T1 WT and Ki-67^-/-^ #2 clone using Lipofectamine 2000 (ThermoFisher). 24h post-transfection, GFP-positive cells were sorted i96-well plates by flow cytometry (FACS Aria, BD). After 10-12 days of culture, colonies were picked and screened (Cellomics, Thermo), using Suz12 or H3K9me3 antibodies. Positive knockout clones were confirmed by PCR, IF and western blotting.

### AOM-DSS-induced colon carcinogenesis

Mice (*Mki67*^*+/+*^; *Mki67*^*+/2ntΔ*^ & *Mki67*^*2ntΔ/2ntΔ*^) were given a single intraperitoneal injection of AOM (10 mg/kg in 0.9% saline; A5486, Sigma-Aldrich), and one week later, 2% Dextran Sodium Sulfate (DSS; MP Biomedicals) was administered in the drinking water for 7 consecutive days. Mice were sacrificed at week 16-post AOM-DSS treatment and colon tissues were removed.

Colons were flushed and fixed overnight in neutral buffered formalin (10%) before paraffin embedding. Briefly, 4µm thick sections were dewaxed in xylene and rehydrated in graded alcohol baths. Slides were incubated in 3% H_2_O_2_ for 20 min and washed in PBS to quench endogenous peroxidase activity. Antigen retrieval was performed by boiling slides for 20 min in 10 mM sodium citrate buffer, pH 6.0. Nonspecific binding sites were blocked in blocking buffer (TBS, pH 7.4, 5% dried milk, 0.5% Triton X-100) for 60 min at RT. Sections were incubated with anti-β-catenin antibody diluted in blocking buffer overnight at 4°C. Envision+ (Dako) was used as a secondary reagent. Signals were developed with DAB (Sigma-Aldrich). After dehydration, sections were mounted in Pertex (Histolab) and imaged using the Nanozoomer-XR Digital slide Scanner C12000-01 (Hamamatsu).

### DNA replication assay, EdU labelling

Cells were treated with 10 µM 5-ethynyl-2’-deoxyuridine (EdU; LifeTechnologies) for the indicated time, harvested, washed once with cold PBS, resuspended in 300µL cold PBS and fixed with 700µL ice-cold 100% ethanol. Click reaction was performed according to the manufacturer instructions (Click-iT™ Plus EdU Alexa Fluor™ 647 Flow Cytometry Assay Kit; Invitrogen) and cells were analysed by flow-cytometry (BD FACSCanto II). FlowJo^®^ software was used for analysis.

### Mammosphere assay

The mammosphere formation assay was performed as previously described (50). Briefly, 500 cells were plated per well in a low-adherence 96-well plate coated with poly-2-hydroxyethyl-methacrylate (poly-Hema). After 7 days in culture (37°C, 5% CO_2_), images of formed mammospheres were acquired and counted using automated high-content microscopy analysis (Cellomics, Thermo).

### Aldehyde Dehydrogenase 1 (ALDH1) activity

ALDH1 enzymatic activity was determined using the ALDEFLUOR kit (Stem Cell Technologies) according to the manufacturer instructions. For each sample, half the cell/substrate mixture was treated with diethylaminobenzaldehyde (DEAB; ALDH inhibitor). ALDEFLUOR/DEAB treated cells were used to define negative gates. FACS was performed with ≥10^5^ cells.

### Xenografts

Animals were housed in the animal facility of IGMM and were maintained in a pathogen-free environment and fed *ad libitum*.

To generate primary tumours, 10^6^ cells (4T1) or 3×10^6^ cells (MDA-MB-231) of log-phase viable ‘mouse pathogen-free’ (Test: IMPACT1, Iddex) were implanted into the fourth inguinal mammary gland (in 50 µl PBS (4T1) or 200 µl PBS (MDA-MB-231)).

Primary tumour volume was measured every week by electronic calliper using the formula “π/6*S^2^ (Smaller radius)*L (Larger radius)”.

At the end of the experiment, following sacrifice, primary tumours were excised and fixed over-night in neutral buffered formalin (10%) before paraffin embedding (see above). IHC analysis of Ki-67 expression of the different tumour tissue sections was conducted as described above.

### Visualisation of lung metastases

Dissected lungs were stained with 15% India Ink diluted in distilled water, and subsequently washed with 5 ml Fekete’s solution (40 ml glacial acid acetic, 80 ml formalin, 580 ml ethanol 100%, 200 ml water). A binocular microscope connected to a digital camera was used to visualise and count the metastatic nodules.

### Automated drug library screen

We performed an initial screen of all 1,280 FDA-approved drugs in the Prestwick Chemical Library (Prestwick Chemical, Illkirch-Graffenstaden, France) at 10 μM in white clear-bottom 96-well plates. Briefly, we prepared seeding solutions at a density of 2 × 10^5^ cells/mL in DMEM complete medium (10%FBS), and dispensed 100 μL into 96-well plates using a Multidrop Combi Dispenser (Thermo Scientific™). Next, we prepared 1x treatment solutions of the Prestwick compounds in DMEM complete medium at a concentration of 10 μM using a Tecan EVO200 robotic liquid handling system (Tecan Trading AG). We removed medium from the pre-seeded 96-well plates and we dispensed 100 μL of 1x treatment solutions using the Tecan robot. We incubated plates at 37°C, 5% CO_2_ for 48 h in an automated incubator (Cytomat, Thermo Scientific™) associated with the Tecan robot. After treatment, cell survival was determined using the CellTiter-Glo^®^ Luminescent Cell Viability Assay (Promega Corporation) in accordance with the manufacturer’s instructions. Luminescence signal was automatically measured with a Tecan Infinity F500 microplate reader (Tecan Trading AG). “Hits” were identified applying the Z-score statistical method (see statistical analysis section). The same method was used for subsequent dose-response experiments on selected “hits”.

### Induction of anemia with PHZ

Anemia was induced in adult *Mki67*^2ntΔ/2ntΔ^ mice and littermate controls (*Mki67*^+/2ntΔ^) by injection of 40 mg/kg phenylhydrazine (PHZ) on days 0, 1, and 3. Animals were sacrificed on day 5 and blood, spleen and bone marrow were collected. Hematocrit was measured as previously described (51). Cell suspensions were generated and immunophenotyping performed by flow cytometry. Cells were stained with Sytox blue or Live/dead fixable viability dye (Life Technologies and eBioscience, respectively) together with the appropriate conjugated anti-CD3, CD62L, CD4, CD8, CD44, CD25, B220, NK1.1, CD11b, Ter119, and CD71 antibodies (eBioscience/ThermoFisher or Becton Dickinson, San Diego, CA). Cells were analysed on a FACS Fortessa (BD Biosciences) flow cytometer. Data analyses were performed using FlowJo Mac v.10 (Tree Star) or DIVA (Becton Dickinson) software.

### Antibodies used for immunophenotying

Antibodies used for immunophenotyping
|  | clone | fluorochrome | reference |
| --- | --- | --- | --- |
| Murine anti-CD3 | 145-2C11 | V421 | 562600 |
| Murine anti-CD4 | RM4-5 | BV711 | 563726 |
| Murine anti-CD8 | 53-6.7 | FITC | 553031 |
| Murine anti-CD8 | 53-6.7 | AF700 | 564983 |
| Murine anti-CD62L | MEL-14 | APC | 553152 |
| Murine anti-CD44 | IM7 | PE-CF594 | 562464 |
| Murine anti-CD25 | PC61 | AAF | 47-0251-82 |
| Murine anti-NK11 | PK136 | APC | 550627 |
| Murine anti-CD11b | M1/70 | BV605 | 563015 |
| Murine anti-Ter119 | Ter-119 (hamster) | FITC | 557915 |
| Murine anti-CD71 | C2 | BV605 | 563013 |
| Murine anti-B220 | RA3-6B2 | eF780 | 47-0452-82 |

### Cell extracts and Western-blotting

Frozen cell pellets (harvested by trypsinization, washed with cold PBS) were lysed directly in Laemmli buffer at 95°C. Protein concentrations were determined by BCA protein assay (Pierce Biotechnology). Proteins were separated by SDS-polyacrylamide gel electrophoresis (SDS-PAGE) (7.5% and 12.5% gels) and transferred to Immobilon membranes (Milipore) at 1.15 mA/cm^2^ for 120 min with a semi-dry blotting apparatus. Membranes were blocked in TBS-T pH 7.6 (20 mM Tris, 140mM NaCl, 0.1% Tween-20) containing non-fat dry milk (5%), incubated with the primary antibody for 2 hours at RT or over-night at 4°C, washed several times with TBS-T for a total of 45 minutes, incubated with secondary antibody at 1/5000 dilution for 1 hour at RT and washed several times in TBS-T. Signals were detected using Western Lightning Plus-ECL (PerkinElmer) and Amersham Hyperfilm^™^ (GE Healthcare).

### qRT-PCR

For reverse transcription (cDNA synthesis), 500 ng of purified RNA in total volume of 10µl, extracted by RNeasy Mini Kit (Qiagen), were mixed with 1µl of 10mM dNTPs mix (LifeTechnologies) and 1µl of 50µM random hexaprimers (NEB). Samples were incubated at 65°C for 5 minutes, transferred to ice, 5µl 5x First Strand Buffer, 2µLl100mM DTT and 1µl RNasin RNase Inhibitor (Promega) were added and samples were incubated at 25°C for 10 minutes, 42°C for 2 minutes. 1µl of SuperScript^®^ III Reverse Transcriptase (LifeTechnologies) was added to each sample and incubated at 50°C for 60 minutes, 70°C for 15 minutes.

qPCR was performed using LightCycler^®^ 480 SYBR Green I Master (Roche) and LightCycler^®^ 480 qPCR machine. The reaction contained 5ng of cDNA, 2µl of 1μM qPCR primer pair, 5µl 2x Master Mix, and final volume made up to 10µl with DNase free water. Primers used for both mouse (Mki67) and human (MKI67) Ki-67 in addition to the housekeeping genes are similar to those used by Sobecki et al., 2016. qPCR was conducted at 95°C for 10 min, and then 40 cycles of 95°C 20s, 58°C 20s and 72°C 20s. The specificity of the reaction was verified by melting curve analysis.

### PCR primers used in this study

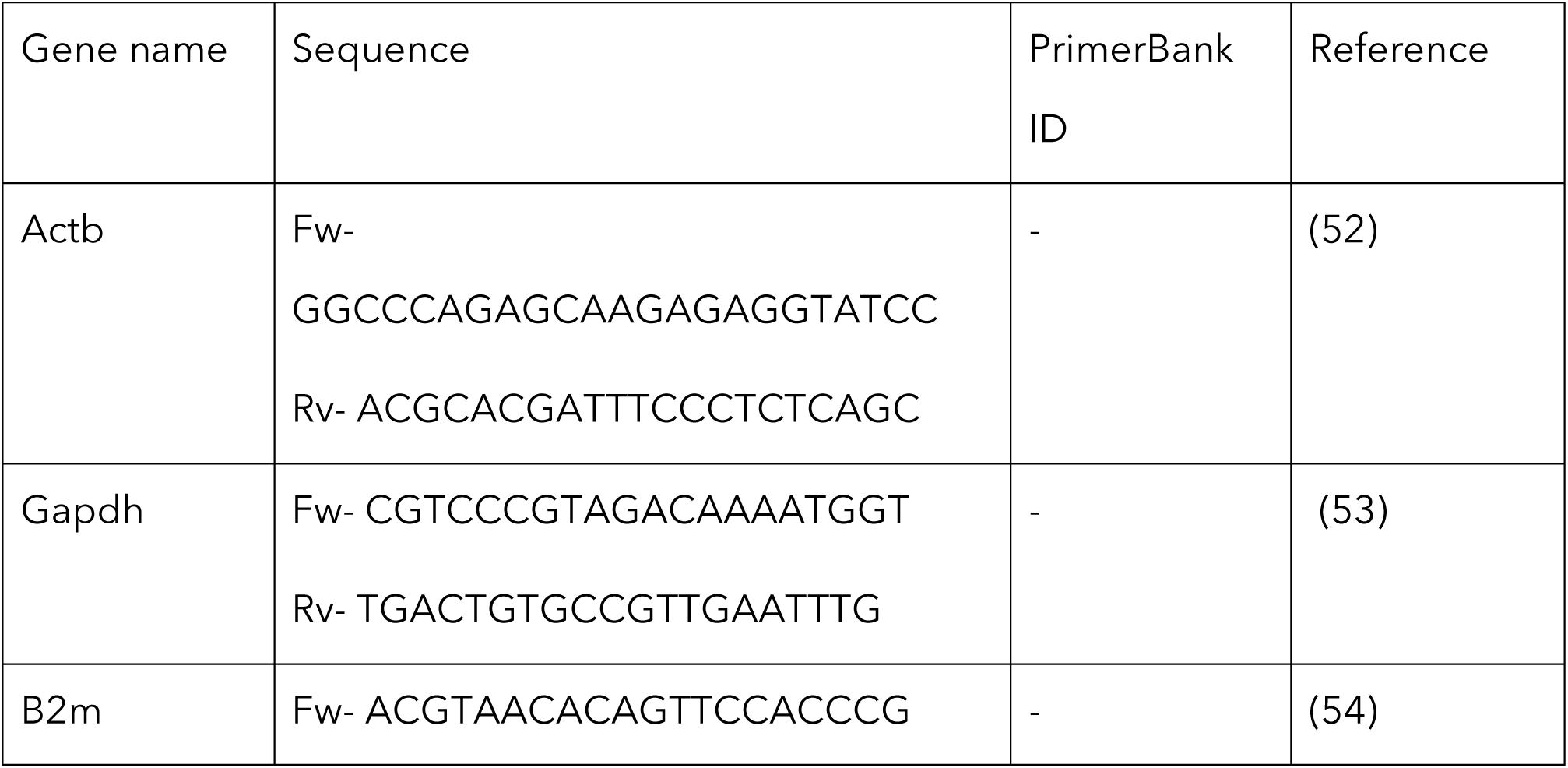
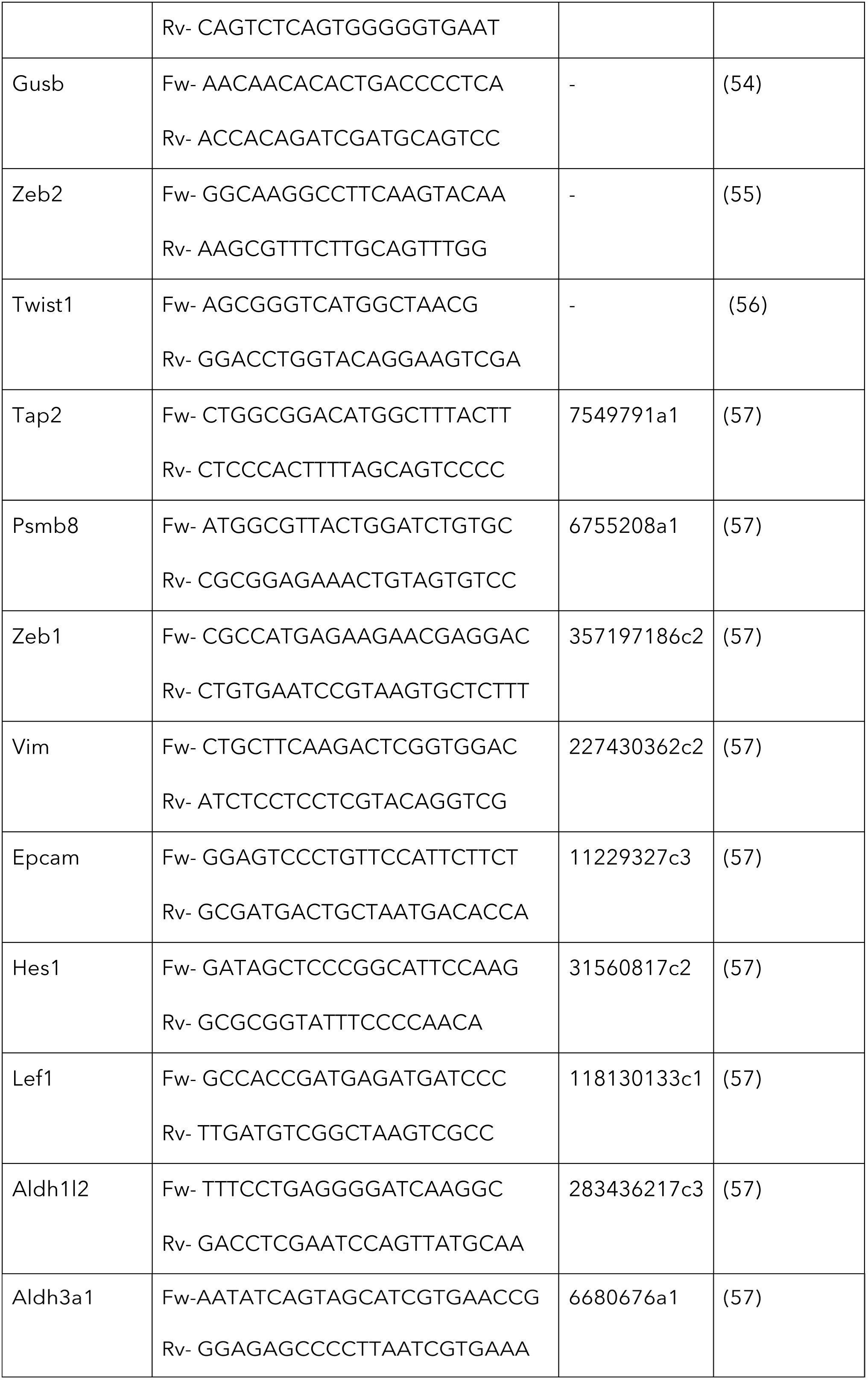

### Colony formation assay

3T3 wild-type (WT) and two Ki-67 TALEN-mutant clones were transduced with either empty control or H-Ras^G12V^ expressing retroviruses. After verification of Ras^G12V^ expression, cells were seeded at 10^5^ cells/well (6 well-plate) in triplicate and allowed to grow for 2-weeks (media were changed every 2 days). Cells were then fixed (4% formaldehyde) and stained with 0.5% (w/v) crystal violet to visualise the colonies formed.

### RNA sequencing library preparation

Total RNA was extracted using Trizol (Life Technologies) following manufacturer’s instructions from wild type and two clones of 4T1 Ki-67 knockout cell lines. RNA integrity was analysed on Agilent 2100 bioanalyzer. For library preparation, cDNA synthesis was performed on rRNA-depleted samples using the TruSeq Stranded Total RNA Library Preparation (RS-122-2301). All sequencing libraries were prepared in two or three biological replicates. Indexed cDNA libraries were sequenced by MGX (Montpellier) on Illumina HiSeq2000 with a single 50 bp read and a 10 bp index read.

### Sequencing of cDNA libraries and data processing

FastQC was used to perform quality control of the sequencing. Using the tool STAR 2.6.0a (2), all the reads that passed the quality control were aligned to the mouse reference genome (GRCm38.p6) and the counts per gene were quantified. The release 93 of the Ensembl mouse genome annotations were used for establishing the coordinates of each gene and their transcripts. Differential expression analysis was performed in R using the DESeq2 (2) library embedded in an in-house script. After normalization of the read counts per gene, a negative binomial generalised linear model was fitted considering single factor design for assessing the differential expression between Mki67 knockout and wild-type samples. Gene set enrichment analysis of the 4T1 *Mki67*^-/-^ cell line vs WT cell line was performed using the javaGSEA (3) desktop application with a log-fold-change pre-ranked gene list.

### Gene set enrichment for transcription factors

The gene set enrichment was performed over the ENCODE and ChEA Consensus TFs from ChIP-X gene set, which contains consensus target genes for transcriptions factors present in ENCODE and ChEA databases. The p value was computed with the Fisher Exact test and then adjusted for multiple hypotesis testing in order to obtain the FDR (false discovery rate) adjusted p-value.

### Statistical analysis

Significant differences between the different experimental groups were tested using an unpaired two-tailed Student’s *t*-test or ANOVA in Prism 5 (GraphPad). For all analyses, p values < 0,05 (*), p values < 0,01 (**), p values < 0,001 (***) and p values < 0,0001 (****) were considered to indicate a statistically significant result.

## Data availability

All RNA-seq raw data will be deposited in GEO and will be publicly available prior to publication of the manuscript.

## SUPPORTING INFORMATION

Supporting information includes six figures, three tables and one dataset.

## ACKNOWLEDGEMENTS

We thank D. Grimanelli for help with transcriptome analysis, and P. Jay, D. Santamaria, M.E. Hochberg and M. Serrano for comments on the manuscript. K.M and M.S. were funded by the Ligue Nationale Contre le Cancer (LNCC); P.S. and K.M. received funding from Worldwide Cancer Research (WWCR). D.F. and L.K. are Inserm employees. This work was initiated and finalised with support from the LNCC and continued with support from Worldwide Cancer Research (WWCR) and the French National Cancer Institute (INCa). The chemical library screen relied on support provided by the Programme Opérationnel FEDER-FSE 2014-2020 Languedoc Roussillon.

## AUTHOR CONTRIBUTIONS

D.F. conceived and supervised the project. K.M., L.K., V.Z., and D.F. designed experiments and interpreted the data. K.M., M.S., P.S., E.A., A.A., S.P., N.P., F.B., B.B., M.P., C.H-K. and V.D. performed experiments. G.D. performed data analysis. K.M., L.K. and D.F. wrote the manuscript.

## Supporting information figure legends

**Fig. S1.**
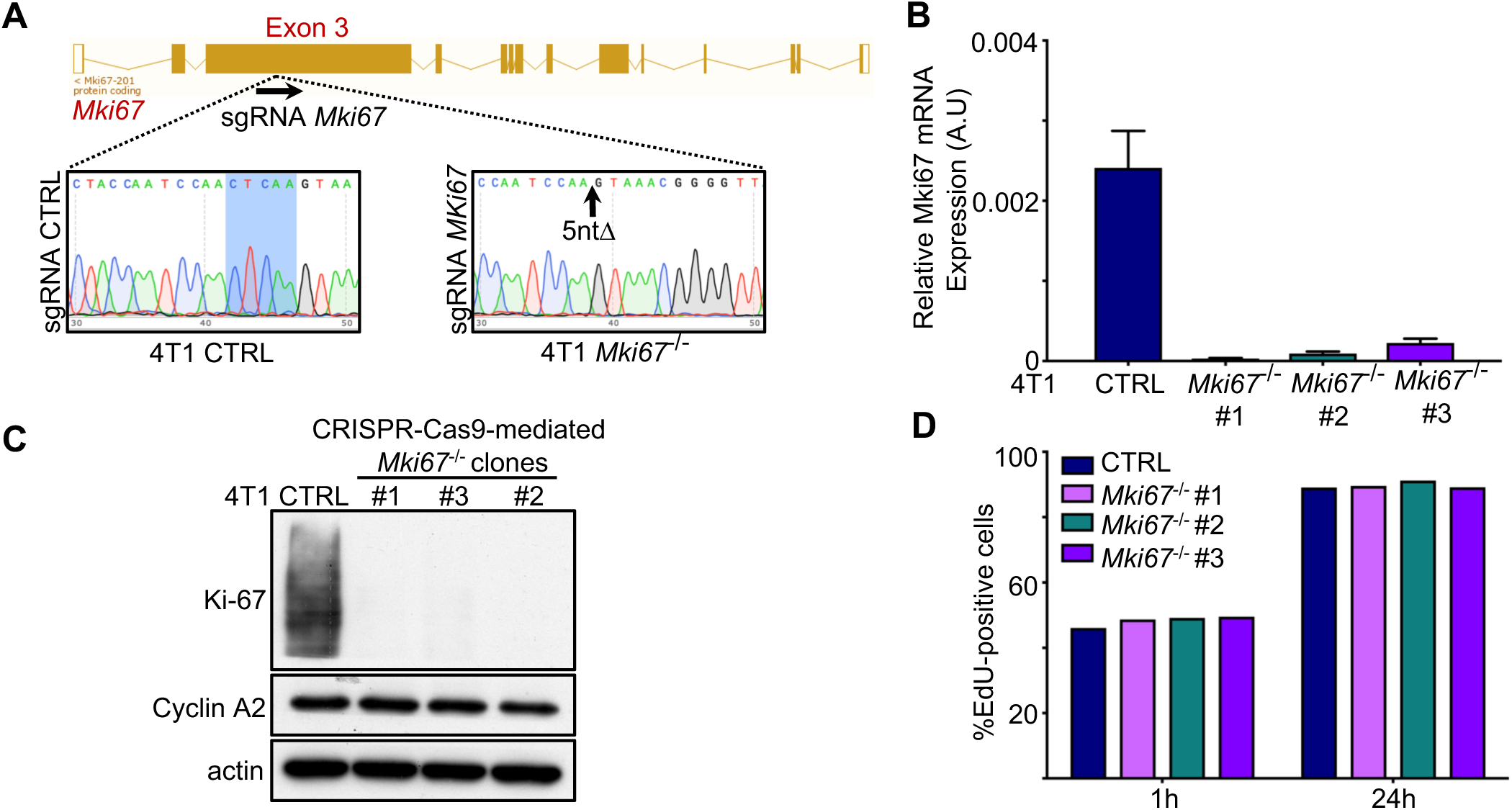
Knockout of *Mki67* does not affect cell proliferation in 4T1 cells. (A) Scheme of CRISPR-Cas9-mediated disruption of *Mki67* gene in 4T1 cells (targeting exon 3 and resulting in a 5nt-deletion). (B) Relative Ki-67 mRNA expression in parental 4T1 cells and three *Mki67*^*-/-*^ clones. (C) Immunoblotting for the indicated proteins of parental 4T1 cells and *Mki67*^*-/-*^ clones. (D) Quantification of the number of CTRL and *Mki67*^*-/-*^ EdU-positive cells, pulsed with EdU for either 1h or 24h.

**Fig. S2.**
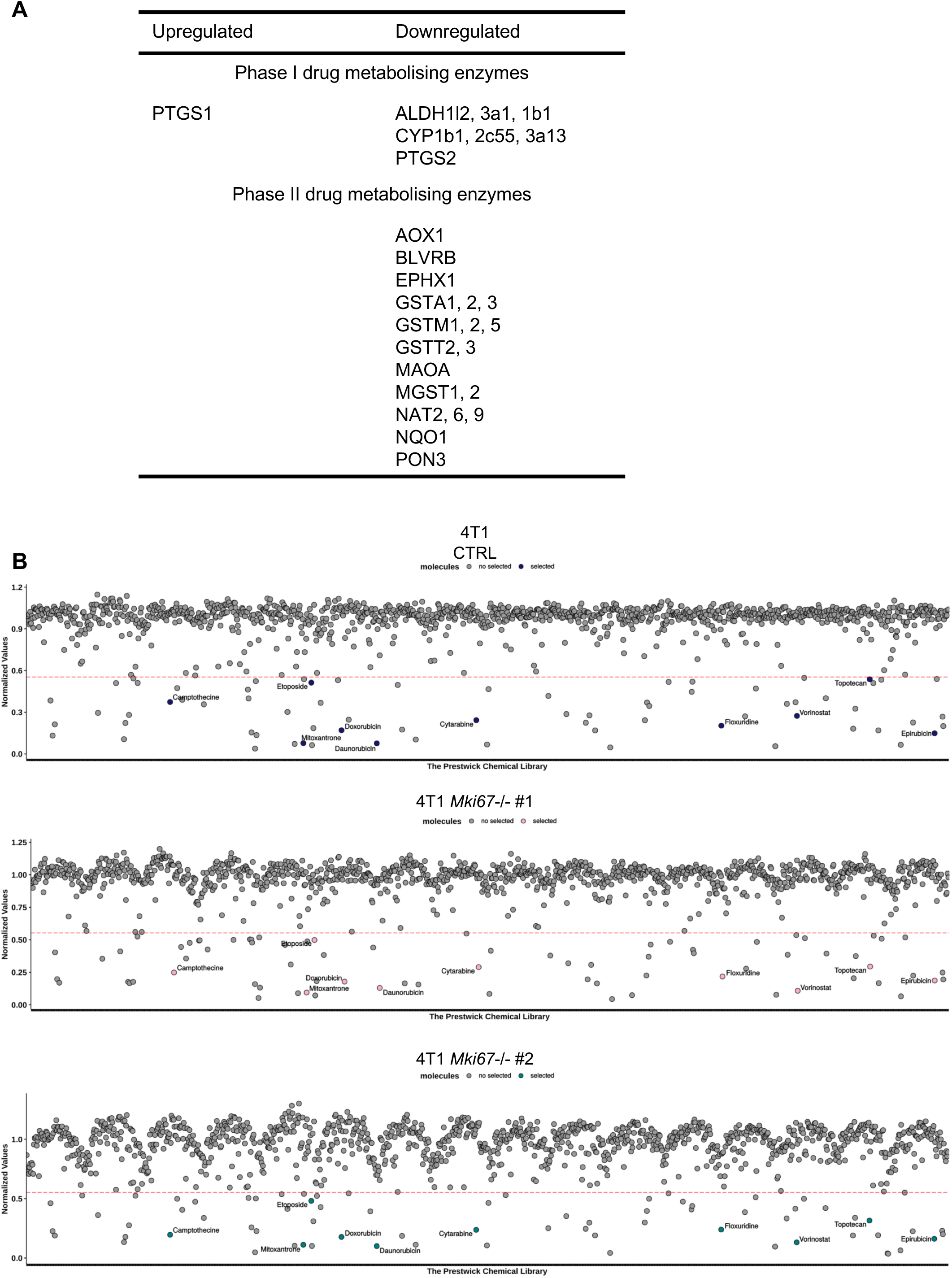
Ki-67 promotes cancer cell drug resistance. (A) List of drug-metabolising enzymes with altered expression in *Mki67*^-/-^ cells. (B) Results of an automated gene-drug screen using the Prestwick chemical library. ‘Hits’ were identified applying the Z-score statistical method and are located below the dashed red line. Drugs chosen for subsequent dose-response experiments are highlighted in colour for CTRL and *Mki67*^-/-^ clones.

**Fig. S3.**
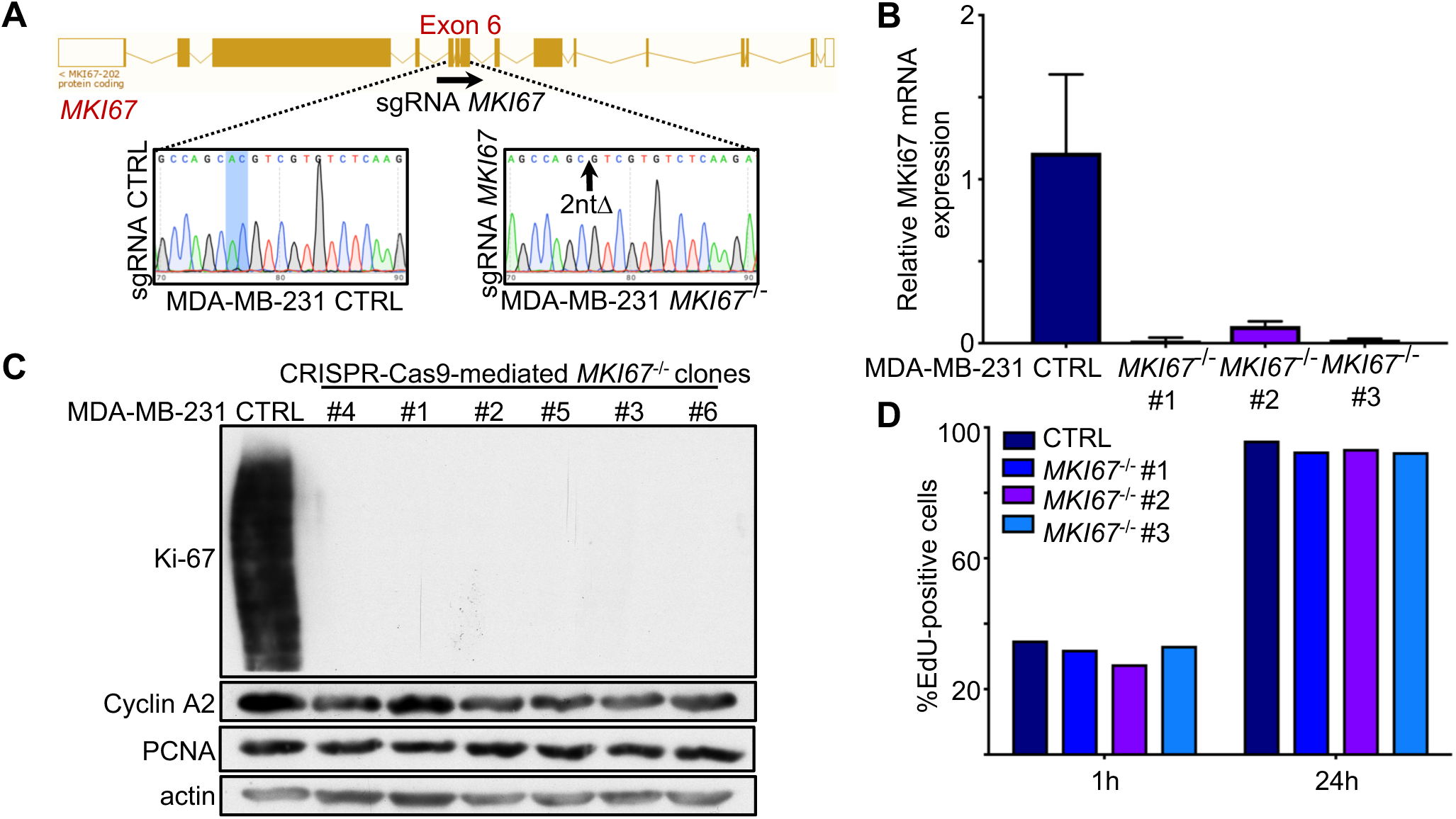
Knockout of *Mki67* does not affect cell proliferation in MDA-MB-231 cells. (A-D) Generation and characterisation of MDA-MB-231 CTRL and *MKI67*^*-/-*^ cells. (A) Scheme of CRISPR-Cas9-mediated disruption of *MKI67* gene (targeting exon 6 and resulting in a 2nt-deletion). (B) Relative Ki-67 mRNA expression in parental MDA-MB-231 cells and three *MKI67*^-/-^ clones. (C) Immunoblotting for the indicated proteins of parental MDA-MB-231 cells and six *MKI67*^-/-^ clones. (D) Quantification of the number of CTRL and *MKI67*^-/-^ MDA-MB-231 EdU-positive cells, pulsed with EdU for either 1h or 24h.

**Fig. S4.**
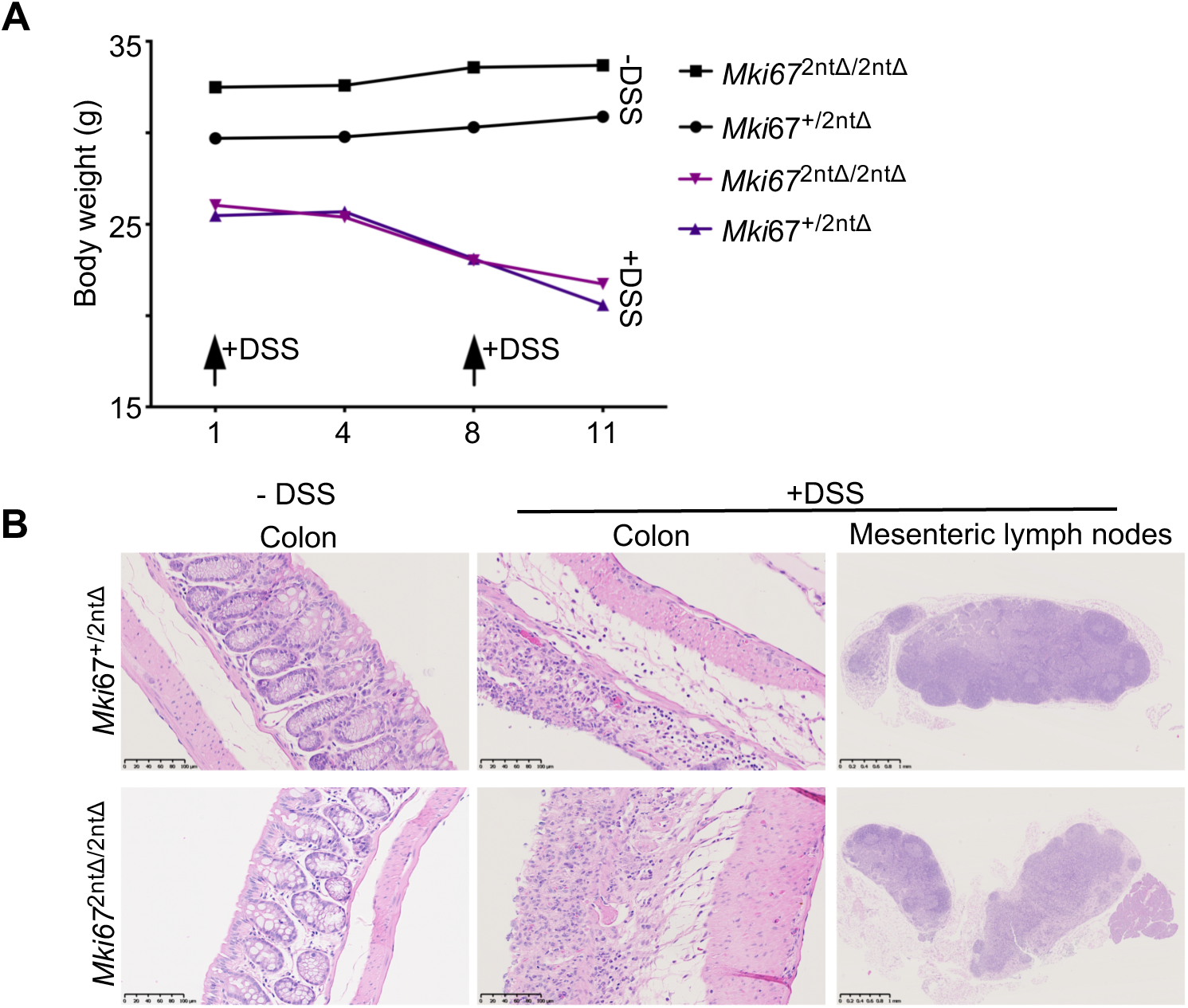
*Mki67*^mt/mt^ mice have normal inflammatory responses and haematopoiesis. (A) Body weight (g) of *Mki67*^+/2ntΔ^ and *Mki67*^2ntΔ/2ntΔ^ mice during the AOM-DSS experiment. (B) Assessment of inflammatory response in colon and mesenteric lymph nodes in *Mki67*^+/2ntΔ^ and *Mki67*^2ntΔ/2ntΔ^ mice, treated or not with DSS (H&E staining; scale bar, 1mm).

**Fig. S5.**
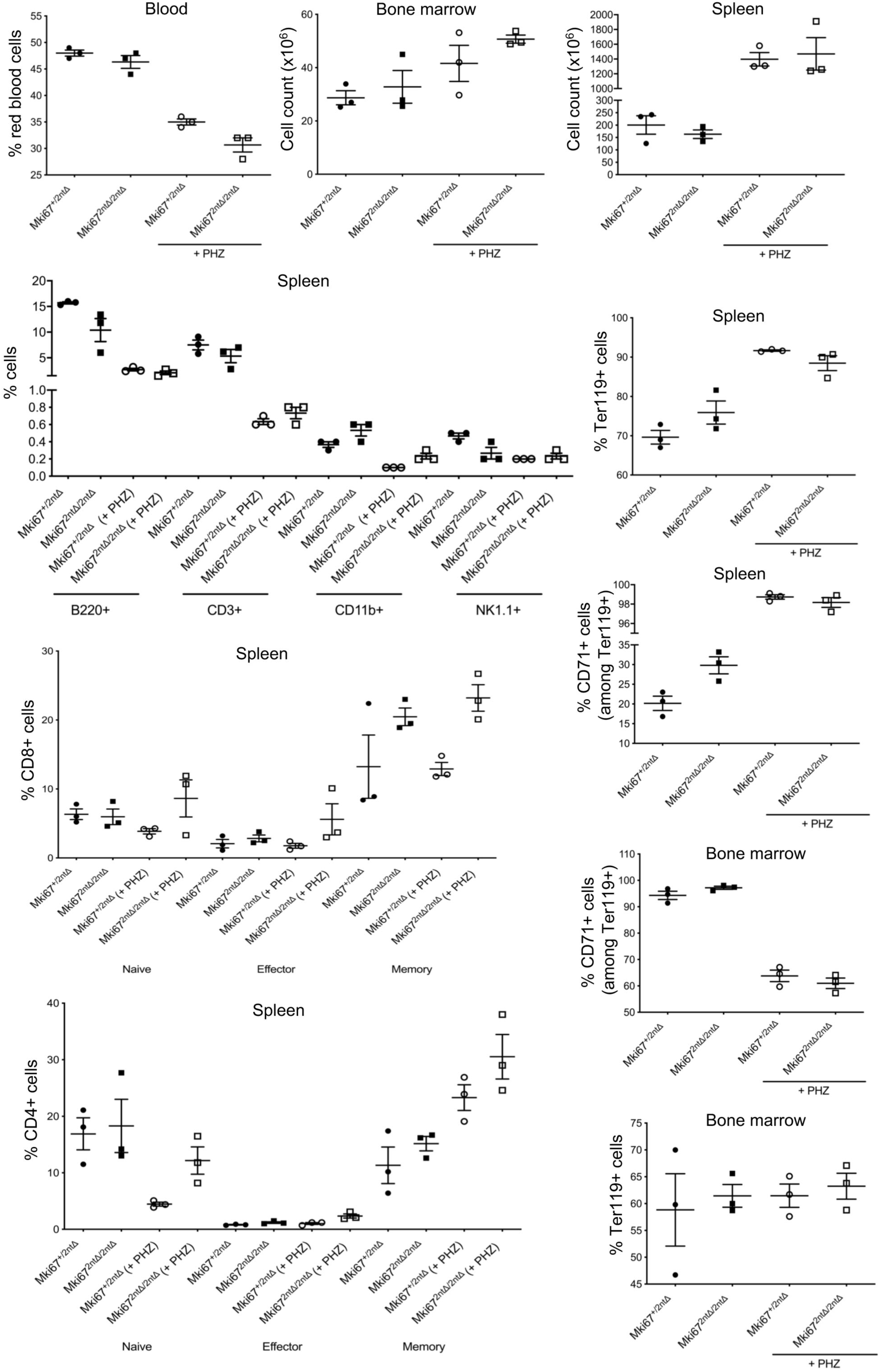
*Mki67*^mt/mt^ mice have normal haematopoiesis. *Mki67*^mt/mt^ mice have normal response to haemolytic anaemia triggered by phenylhydrazine (PHZ) treatment. Analysis of number and percentage of indicated populations of blood, bone marrow and spleen cells in *Mki67*^+/2ntΔ^ and *Mki67*^2ntΔ/2ntΔ^ mouse, treated or not with PHZ.

**Fig. S6.**
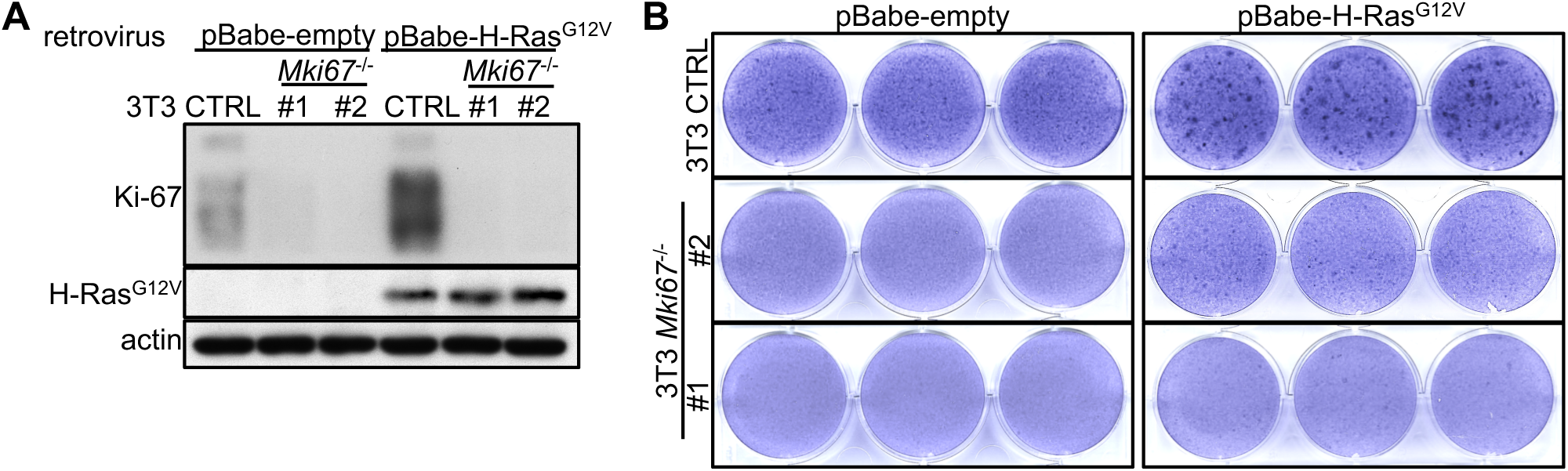
Ki-67 is required for oncogene-induced transformation of 3T3 fibroblasts. (A, B) CTRL or TALEN-mutated *Mki67*^-/-^ 3T3 fibroblasts were transduced with either empty or H-Ras^G12V^-expressing retroviruses. (A) Immunoblot analysis of the expression of the indicated proteins. Actin serves as loading control. (B) Crystal violet staining of the colonies formed.

**Table S1. (separate file)**

Table showing genes statistically significantly deregulated between WT and Ki-67 knockout 3T3 cells (adjusted p-value, padj<0.05), as determined by RNA-sequencing analysis.

**Table S2. (separate file)**

Table showing genes statistically significantly deregulated between WT and Ki-67 knockout 4T1 cells (adjusted p-value, padj<0.05), as determined by RNA-sequencing analysis.

**Table S3. (separate file)**

Table showing genes statistically significantly deregulated between tumours derived from WT and Ki-67 knockout 4T1 cells (adjusted p-value, padj<0.05), as determined by RNA-sequencing analysis.

**Supporting Data S1. (separate file)**

Report of Prestwick chemical library screen for WT and two clones (clone 101 and 119) of Ki-67 knockout 4T1 cells, showing raw luminescence data from 96-well plates of CellTiter-Glo (Promega) 72 hours after treatment with 10µM of each compound.

