## Supplementary Data for "Ki-67 promotes sequential stages of tumourigenesis by enabling cellular plasticity"

### Screen

*Cédric HASSEN-KHODJA*

*15 février 2019*

#### Quality Control

##### Z-Factor

##### Description

The Z-factor is a measure that quantifies the separation between the distribution of positive and negative controls.

$$Z - factor = 1 - \frac{3(\sigma p + \sigma n)}{\mid \mu p - \mu n \mid}$$

If robust, the Z-factor is calculated using robust estimates of location (median) and spread (mad).

##### Theoretical interpretation

| Z-Factor | Interpretation |
|---|---|
| 1.0 | Ideal assay |
| [0.5 - 1.0] | Excellent assay |
| [0 - 0.5] | Marginal assay |
| < 0 | Bad assay |

##### Visualization

for the screen we have as positive control salinomycine (SAL) and as negative control DMSO.
Let's look at the distribution of the controls for each clones: WT, cl101 and cl119.

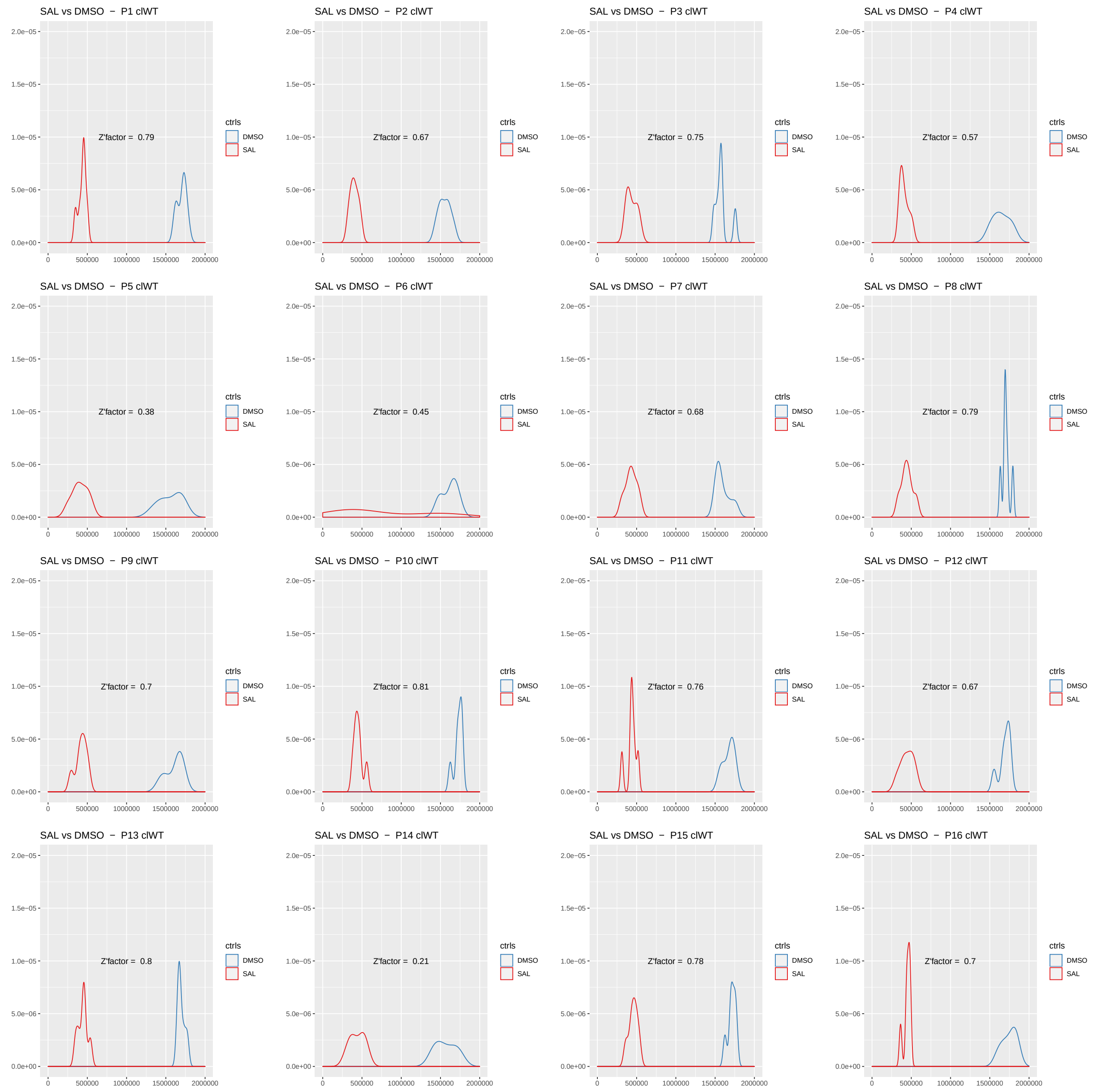
SAL vs DMSO − P1 clWT
Z'factor = 0.79
SAL vs DMSO − P2 clWT
Z'factor = 0.67
SAL vs DMSO − P3 clWT
Z'factor = 0.75
SAL vs DMSO − P4 clWT
Z'factor = 0.57
SAL vs DMSO − P5 clWT
Z'factor = 0.38
SAL vs DMSO − P6 clWT
Z'factor = 0.45
SAL vs DMSO − P7 clWT
Z'factor = 0.68
SAL vs DMSO − P8 clWT
Z'factor = 0.79
SAL vs DMSO − P9 clWT
Z'factor = 0.7
SAL vs DMSO − P10 clWT
Z'factor = 0.81
SAL vs DMSO − P11 clWT
Z'factor = 0.76
SAL vs DMSO − P12 clWT
Z'factor = 0.67
SAL vs DMSO − P13 clWT
Z'factor = 0.8
SAL vs DMSO − P14 clWT
Z'factor = 0.21
SAL vs DMSO − P15 clWT
Z'factor = 0.78
SAL vs DMSO − P16 clWT
Z'factor = 0.7
ctrls
DMSO
SAL
0.0e+00
5.0e−06
1.0e−05
1.5e−05
2.0e−05
0
500000
1000000
1500000
2000000

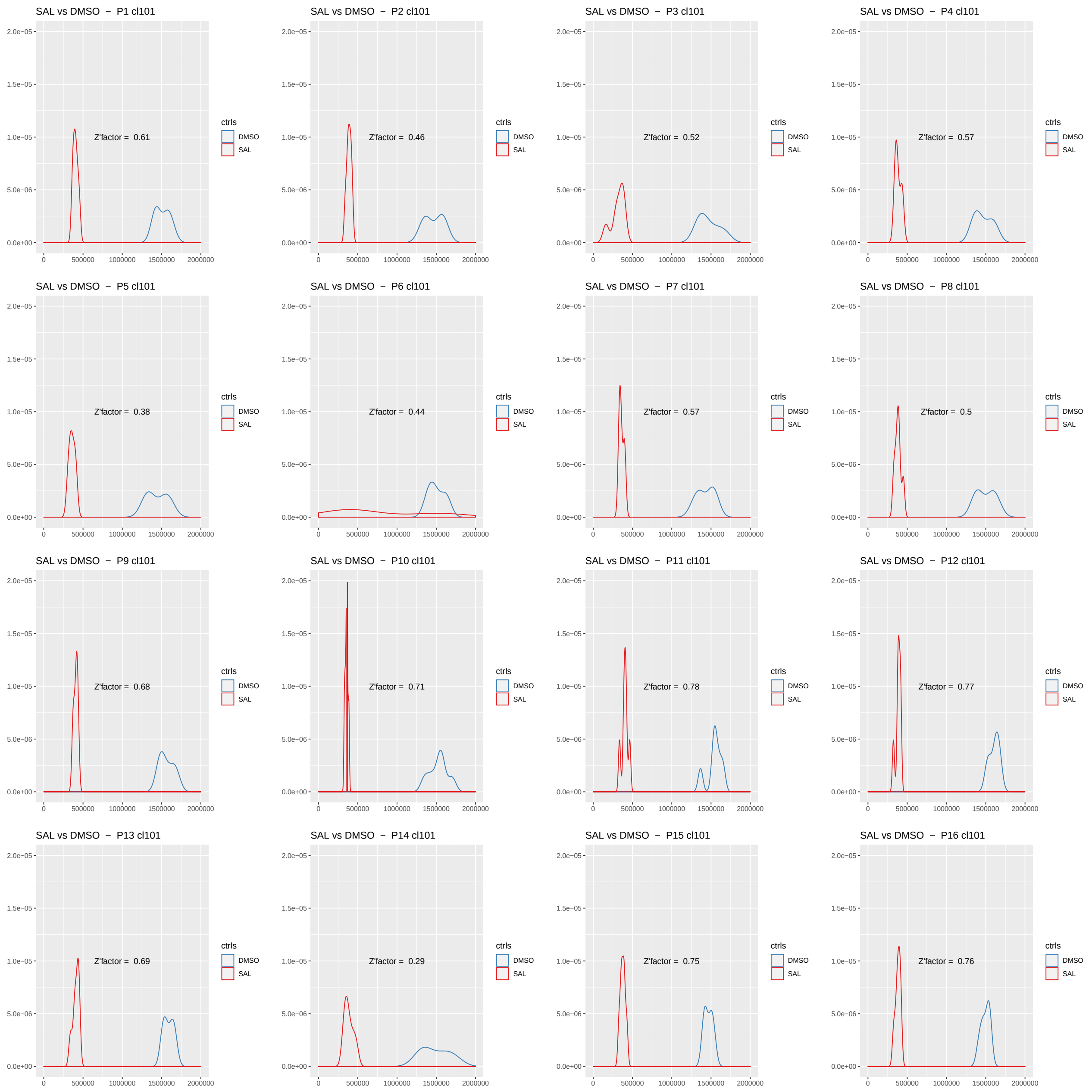
SAL vs DMSO − P1 cl101
Z'factor = 0.61
SAL vs DMSO − P2 cl101
Z'factor = 0.46
SAL vs DMSO − P3 cl101
Z'factor = 0.52
SAL vs DMSO − P4 cl101
Z'factor = 0.57
SAL vs DMSO − P5 cl101
Z'factor = 0.38
SAL vs DMSO − P6 cl101
Z'factor = 0.44
SAL vs DMSO − P7 cl101
Z'factor = 0.57
SAL vs DMSO − P8 cl101
Z'factor = 0.5
SAL vs DMSO − P9 cl101
Z'factor = 0.68
SAL vs DMSO − P10 cl101
Z'factor = 0.71
SAL vs DMSO − P11 cl101
Z'factor = 0.78
SAL vs DMSO − P12 cl101
Z'factor = 0.77
SAL vs DMSO − P13 cl101
Z'factor = 0.69
SAL vs DMSO − P14 cl101
Z'factor = 0.29
SAL vs DMSO − P15 cl101
Z'factor = 0.75
SAL vs DMSO − P16 cl101
Z'factor = 0.76
ctrls
DMSO
SAL
2.0e−05
1.5e−05
1.0e−05
5.0e−06
0.0e+00
0
500000
1000000
1500000
2000000

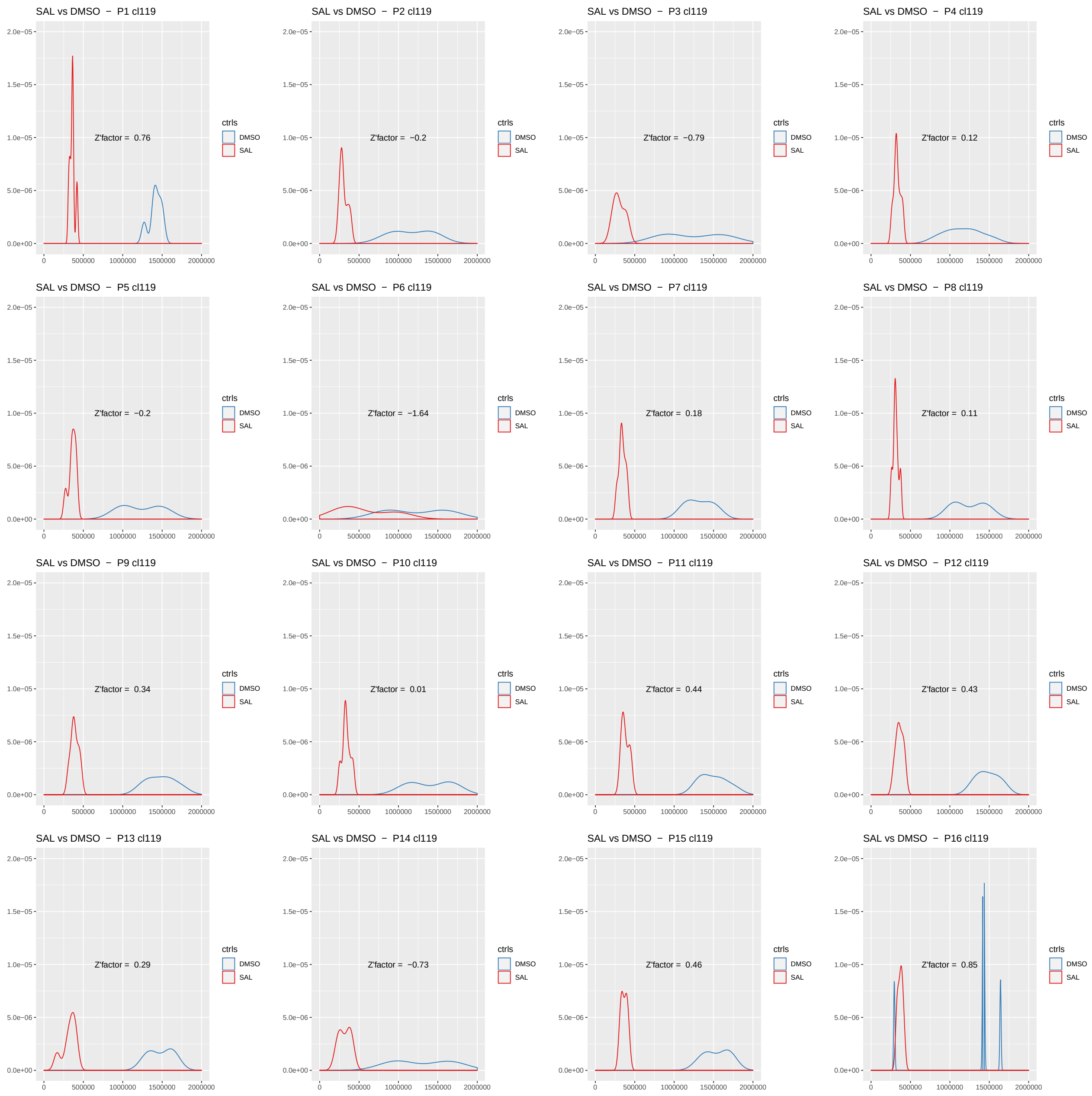


#### Z-Factor interpretation

Overall, the values are good for clones WT and cl101; however, aberrant dots of negative control or a large dispersion of values for DMSO are observed for the plates 5, 6 and 14.
For the cl119 clone, the bad values of z-factor are due essentially to a greater dispersion of the values of the DMSO, thus less reproducible. However we have a good positive control (SAL) that we can take into account to define a threshold.

#### Data normalization

##### Description

Normalization is performed separately for each plate. Different normalization methods exist. We have chosen two of these methods:

- **median based normalization**: plates effects are corrected by the median value across wells that are annotated as sample.
- **Normalized percent inhibition (NPI)**: each measurement is subtracted from the average of the intensities on the plate positive controls, and this result is divided by the difference between the means of the measurements on the positive and the negative controls.

$$x' = \frac{\mu pi - Xki}{\mu pi - \mu ni} * 100$$

where X is the raw value, n the negative control, p the positive control, k-th well and i-th plate

#### Scores normalized values

##### Description

This method scores the normalized values, this is calculated by subtracting the overall median from each measurement and dividing the result by the mad.

$$score = \frac{x'ki - median(x'i)}{mad(x'i)}$$

where x' is the normalized values, k-th well and i-th plate

#### List of results for the median based normalization

For each clone, we have a result table that contains the raw, the normalized and scored values per plate and wells.

#### clone WT

| Plate | Well | Value | Treatment | Norm | Score |
|---|---|---|---|---|---|
| P1 clWT | A01 | 348840 | SAL | 0.2119513 | -14.8023902 |
| P1 clWT | A02 | 1588600 | sample | 0.9652155 | -0.6533773 |
| P1 clWT | A03 | 1627700 | sample | 0.9889723 | -0.2071406 |
| P1 clWT | A04 | 1632900 | sample | 0.9921317 | -0.1477945 |
| P1 clWT | A05 | 1651100 | sample | 1.0031898 | 0.0599167 |
| P1 clWT | A06 | 1532300 | sample | 0.9310083 | -1.2959124 |
| P1 clWT | A07 | 1682600 | sample | 1.0223289 | 0.4194168 |
| P1 clWT | A08 | 1547100 | sample | 0.9400006 | -1.1270044 |
| P1 clWT | A09 | 1636500 | sample | 0.9943190 | -0.1067088 |
| P1 clWT | A10 | 1606100 | sample | 0.9758483 | -0.4536550 |
| P1 clWT | A11 | 1492700 | sample | 0.9069478 | -1.7478555 |
| P1 clWT | A12 | 1485300 | empty | 0.9024516 | -1.8323095 |
| P1 clWT | B01 | 404510 | SAL | 0.2457757 | -14.1670450 |
| P1 clWT | B02 | 1735000 | sample | 1.0541665 | 1.0174425 |
| P1 clWT | B03 | 1614500 | sample | 0.9809521 | -0.3577882 |
| P1 clWT | B04 | 1711700 | sample | 1.0400097 | 0.7515265 |
| P1 clWT | B05 | 1709600 | sample | 1.0387338 | 0.7275598 |
| P1 clWT | B06 | 1712800 | sample | 1.0406781 | 0.7640805 |
| P1 clWT | B07 | 1634900 | sample | 0.9933469 | -0.1249691 |
| P1 clWT | B08 | 1691900 | sample | 1.0279795 | 0.5255550 |
| P1 clWT | B09 | 1502400 | sample | 0.9128414 | -1.6371523 |
| P1 clWT | B10 | 1748000 | sample | 1.0620652 | 1.1658076 |
| P1 clWT | B11 | 1642000 | sample | 0.9976608 | -0.0439389 |
| P1 clWT | B12 | 1695500 | empty | 1.0301668 | 0.5666407 |
| P1 clWT | C01 | 458970 | SAL | 0.2788650 | -13.5455092 |
| P1 clWT | C02 | 1713300 | sample | 1.0409819 | 0.7697868 |
| P1 clWT | C03 | 1712000 | sample | 1.0401920 | 0.7549503 |
| P1 clWT | C04 | 1658800 | sample | 1.0078683 | 0.1477945 |
| P1 clWT | C05 | 1584100 | sample | 0.9624814 | -0.7047344 |
| P1 clWT | C06 | 1322900 | sample | 0.8037792 | -3.6857325 |
| P1 clWT | C07 | 1705300 | sample | 1.0361212 | 0.6784852 |
| P1 clWT | C08 | 1681500 | sample | 1.0216605 | 0.4068629 |
| P1 clWT | C09 | 1673700 | sample | 1.0169213 | 0.3178438 |
| P1 clWT | C10 | 1720200 | sample | 1.0451742 | 0.8485345 |
| P1 clWT | C11 | 1704600 | sample | 1.0356958 | 0.6704963 |
| P1 clWT | C12 | 1725800 | DMSO | 1.0485767 | 0.9124456 |
| P1 clWT | D01 | 1777400 | DMSO | 1.0799283 | 1.5013411 |
| P1 clWT | D02 | 1613600 | sample | 0.9804053 | -0.3680597 |
| P1 clWT | D03 | 1663700 | sample | 1.0108455 | 0.2037168 |
| P1 clWT | D04 | 1603400 | sample | 0.9742079 | -0.4844693 |
| P1 clWT | D05 | 632520 | sample | 0.3843121 | -11.5648345 |
| P1 clWT | D06 | 1655900 | sample | 1.0061063 | 0.1146977 |
| P1 clWT | D07 | 1690900 | sample | 1.0273719 | 0.5141423 |
| P1 clWT | D08 | 216120 | sample | 0.1313121 | -16.3170842 |
| P1 clWT | D09 | 1693200 | sample | 1.0287693 | 0.5403915 |
| P1 clWT | D10 | 1745700 | sample | 1.0606677 | 1.1395584 |
| P1 clWT | D11 | 350470 | sample | 0.2129416 | -14.7837875 |
| P1 clWT | D12 | 1620900 | DMSO | 0.9848407 | -0.2847469 |
| P1 clWT | E01 | 1639200 | DMSO | 0.9959595 | -0.0758945 |
| P1 clWT | E02 | 1715700 | sample | 1.0424401 | 0.7971773 |
| P1 clWT | E03 | 1705800 | sample | 1.0364249 | 0.6841916 |
| P1 clWT | E04 | 1651000 | sample | 1.0031291 | 0.0587754 |
| P1 clWT | E05 | 1683800 | sample | 1.0230580 | 0.4331121 |
| P1 clWT | E06 | 1644300 | sample | 0.9990582 | -0.0176897 |

*(continued)*

| Plate | Well | Value | Treatment | Norm | Score |
|---|---|---|---|---|---|
| P1 clWT | E07 | 1647400 | sample | 1.0009418 | 0.0176897 |
| P1 clWT | E08 | 1611600 | sample | 0.9791901 | -0.3908851 |
| P1 clWT | E09 | 1587300 | sample | 0.9644257 | -0.6682138 |
| P1 clWT | E10 | 1703800 | sample | 1.0352098 | 0.6613662 |
| P1 clWT | E11 | 1728900 | sample | 1.0504602 | 0.9478250 |
| P1 clWT | E12 | 1732200 | DMSO | 1.0524653 | 0.9854869 |
| P1 clWT | F01 | 1719900 | DMSO | 1.0449919 | 0.8451107 |
| P1 clWT | F02 | 1772200 | sample | 1.0767688 | 1.4419951 |
| P1 clWT | F03 | 1688100 | sample | 1.0256706 | 0.4821867 |
| P1 clWT | F04 | 1231200 | sample | 0.7480633 | -4.7322774 |
| P1 clWT | F05 | 1683800 | sample | 1.0230580 | 0.4331121 |
| P1 clWT | F06 | 1664200 | sample | 1.0111493 | 0.2094231 |
| P1 clWT | F07 | 1685100 | sample | 1.0238479 | 0.4479486 |
| P1 clWT | F08 | 1548000 | sample | 0.9405474 | -1.1167330 |
| P1 clWT | F09 | 1514500 | sample | 0.9201932 | -1.4990586 |
| P1 clWT | F10 | 1745700 | sample | 1.0606677 | 1.1395584 |
| P1 clWT | F11 | 1673700 | sample | 1.0169213 | 0.3178438 |
| P1 clWT | F12 | 454800 | SAL | 0.2763314 | -13.5931002 |
| P1 clWT | G01 | 1618800 | empty | 0.9835647 | -0.3087136 |
| P1 clWT | G02 | 1716900 | sample | 1.0431692 | 0.8108726 |
| P1 clWT | G03 | 1636700 | sample | 0.9944406 | -0.1044262 |
| P1 clWT | G04 | 1603800 | sample | 0.9744509 | -0.4799042 |
| P1 clWT | G05 | 1366500 | sample | 0.8302701 | -3.1881386 |
| P1 clWT | G06 | 1587900 | sample | 0.9647902 | -0.6613662 |
| P1 clWT | G07 | 1633200 | sample | 0.9923140 | -0.1443707 |
| P1 clWT | G08 | 1551800 | sample | 0.9428563 | -1.0733647 |
| P1 clWT | G09 | 1627900 | sample | 0.9890938 | -0.2048580 |
| P1 clWT | G10 | 1723200 | sample | 1.0469970 | 0.8827726 |
| P1 clWT | G11 | 1735200 | sample | 1.0542881 | 1.0197250 |
| P1 clWT | G12 | 497500 | SAL | 0.3022754 | -13.1057777 |
| P1 clWT | H01 | 1476200 | empty | 0.8969226 | -1.9361651 |
| P1 clWT | H02 | 1290400 | sample | 0.7840326 | -4.0566454 |
| P1 clWT | H03 | 1416900 | sample | 0.8608925 | -2.6129384 |
| P1 clWT | H04 | 1536500 | sample | 0.9335602 | -1.2479791 |
| P1 clWT | H05 | 1412100 | sample | 0.8579761 | -2.6677194 |
| P1 clWT | H06 | 1533700 | sample | 0.9318589 | -1.2799347 |
| P1 clWT | H07 | 1069700 | sample | 0.6499377 | -6.5754290 |
| P1 clWT | H08 | 1461000 | sample | 0.8876872 | -2.1096382 |
| P1 clWT | H09 | 1095000 | sample | 0.6653097 | -6.2866876 |
| P1 clWT | H10 | 1672300 | sample | 1.0160707 | 0.3018660 |
| P1 clWT | H11 | 1679400 | sample | 1.0203846 | 0.3828962 |
| P1 clWT | H12 | 446080 | SAL | 0.2710332 | -13.6926189 |
| P2 clWT | A01 | 327460 | SAL | 0.2074107 | -8.9172453 |
| P2 clWT | A02 | 1547200 | sample | 0.9799848 | -0.2251866 |
| P2 clWT | A03 | 1366100 | sample | 0.8652774 | -1.5157336 |
| P2 clWT | A04 | 1414900 | sample | 0.8961870 | -1.1679771 |
| P2 clWT | A05 | 1560600 | sample | 0.9884723 | -0.1296961 |
| P2 clWT | A06 | 1480500 | sample | 0.9377375 | -0.7005012 |
| P2 clWT | A07 | 1497100 | sample | 0.9482518 | -0.5822070 |
| P2 clWT | A08 | 1452400 | sample | 0.9199392 | -0.9007462 |
| P2 clWT | A09 | 1423000 | sample | 0.9013175 | -1.1102553 |
| P2 clWT | A10 | 1567000 | sample | 0.9925260 | -0.0840887 |
| P2 clWT | A11 | 1468600 | sample | 0.9302002 | -0.7853025 |
| P2 clWT | A12 | 1452100 | empty | 0.9197492 | -0.9028841 |
| P2 clWT | B01 | 362750 | SAL | 0.2297631 | -8.6657632 |
| P2 clWT | B02 | 1693900 | sample | 1.0729035 | 0.8202207 |
| P2 clWT | B03 | 1650500 | sample | 1.0454142 | 0.5109455 |
| P2 clWT | B04 | 1609700 | sample | 1.0195718 | 0.2201983 |
| P2 clWT | B05 | 1645400 | sample | 1.0421839 | 0.4746021 |
| P2 clWT | B06 | 1697000 | sample | 1.0748670 | 0.8423118 |
| P2 clWT | B07 | 1546300 | sample | 0.9794147 | -0.2316001 |

*(continued)*

| Plate | Well | Value | Treatment | Norm | Score |
|---|---|---|---|---|---|
| P2 clWT | B08 | 1207800 | sample | 0.7650114 | -2.6438042 |
| P2 clWT | B09 | 1696500 | sample | 1.0745503 | 0.8387487 |
| P2 clWT | B10 | 1809200 | sample | 1.1459336 | 1.6418666 |
| P2 clWT | B11 | 465380 | sample | 0.2947682 | -7.9344057 |
| P2 clWT | B12 | 1630600 | empty | 1.0328097 | 0.3691349 |
| P2 clWT | C01 | 387470 | SAL | 0.2454206 | -8.4896046 |
| P2 clWT | C02 | 1524800 | sample | 0.9657968 | -0.3848125 |
| P2 clWT | C03 | 1509200 | sample | 0.9559159 | -0.4959805 |
| P2 clWT | C04 | 1483900 | sample | 0.9398911 | -0.6762723 |
| P2 clWT | C05 | 1679700 | sample | 1.0639093 | 0.7190292 |
| P2 clWT | C06 | 1671800 | sample | 1.0589055 | 0.6627326 |
| P2 clWT | C07 | 1486200 | sample | 0.9413479 | -0.6598821 |
| P2 clWT | C08 | 1584900 | sample | 1.0038637 | 0.0434696 |
| P2 clWT | C09 | 1635800 | sample | 1.0361034 | 0.4061909 |
| P2 clWT | C10 | 1759900 | sample | 1.1147074 | 1.2905470 |
| P2 clWT | C11 | 1687800 | sample | 1.0690398 | 0.7767511 |
| P2 clWT | C12 | 1589600 | DMSO | 1.0068406 | 0.0769625 |
| P2 clWT | D01 | 1444400 | DMSO | 0.9148721 | -0.9577555 |
| P2 clWT | D02 | 1668500 | sample | 1.0568153 | 0.6392163 |
| P2 clWT | D03 | 1659500 | sample | 1.0511148 | 0.5750809 |
| P2 clWT | D04 | 1624500 | sample | 1.0289460 | 0.3256654 |
| P2 clWT | D05 | 1604800 | sample | 1.0164682 | 0.1852801 |
| P2 clWT | D06 | 1462200 | sample | 0.9261464 | -0.8309099 |
| P2 clWT | D07 | 1594300 | sample | 1.0098176 | 0.1104554 |
| P2 clWT | D08 | 1717700 | sample | 1.0879782 | 0.9898232 |
| P2 clWT | D09 | 1763900 | sample | 1.1172409 | 1.3190517 |
| P2 clWT | D10 | 1715800 | sample | 1.0867748 | 0.9762835 |
| P2 clWT | D11 | 1765500 | sample | 1.1182544 | 1.3304535 |
| P2 clWT | D12 | 1595200 | DMSO | 1.0103876 | 0.1168690 |
| P2 clWT | E01 | 1517700 | DMSO | 0.9612997 | -0.4354082 |
| P2 clWT | E02 | 1730600 | sample | 1.0961490 | 1.0817506 |
| P2 clWT | E03 | 1668100 | sample | 1.0565619 | 0.6363658 |
| P2 clWT | E04 | 1676600 | sample | 1.0619458 | 0.6969382 |
| P2 clWT | E05 | 1532400 | sample | 0.9706106 | -0.3306537 |
| P2 clWT | E06 | 805110 | sample | 0.5099506 | -5.5134364 |
| P2 clWT | E07 | 1588500 | sample | 1.0061439 | 0.0691237 |
| P2 clWT | E08 | 1495100 | sample | 0.9469851 | -0.5964593 |
| P2 clWT | E09 | 1678400 | sample | 1.0630859 | 0.7097652 |
| P2 clWT | E10 | 1661200 | sample | 1.0521915 | 0.5871953 |
| P2 clWT | E11 | 1747500 | sample | 1.1068533 | 1.2021827 |
| P2 clWT | E12 | 1665600 | DMSO | 1.0549785 | 0.6185504 |
| P2 clWT | F01 | 1503600 | DMSO | 0.9523689 | -0.5358870 |
| P2 clWT | F02 | 1731800 | sample | 1.0969090 | 1.0903020 |
| P2 clWT | F03 | 1585600 | sample | 1.0043071 | 0.0484579 |
| P2 clWT | F04 | 1339900 | sample | 0.8486825 | -1.7024389 |
| P2 clWT | F05 | 1664200 | sample | 1.0540917 | 0.6085738 |
| P2 clWT | F06 | 1565200 | sample | 0.9913859 | -0.0969157 |
| P2 clWT | F07 | 167120 | sample | 0.1058525 | -10.0598533 |
| P2 clWT | F08 | 1643400 | sample | 1.0409172 | 0.4603497 |
| P2 clWT | F09 | 351210 | sample | 0.2224538 | -8.7479991 |
| P2 clWT | F10 | 1673200 | sample | 1.0597922 | 0.6727092 |
| P2 clWT | F11 | 1794800 | sample | 1.1368128 | 1.5392499 |
| P2 clWT | F12 | 444250 | SAL | 0.2813846 | -8.0849814 |
| P2 clWT | G01 | 1573800 | empty | 0.9968330 | -0.0356308 |
| P2 clWT | G02 | 1511200 | sample | 0.9571827 | -0.4817282 |
| P2 clWT | G03 | 1579900 | sample | 1.0006967 | 0.0078388 |
| P2 clWT | G04 | 443550 | sample | 0.2809412 | -8.0899697 |
| P2 clWT | G05 | 1361000 | sample | 0.8620471 | -1.5520770 |
| P2 clWT | G06 | 1587100 | sample | 1.0052572 | 0.0591471 |
| P2 clWT | G07 | 898550 | sample | 0.5691348 | -4.8475683 |
| P2 clWT | G08 | 1577700 | sample | 0.9993033 | -0.0078388 |

*(continued)*

| Plate | Well | Value | Treatment | Norm | Score |
|---|---|---|---|---|---|
| P2 clWT | G09 | 1530900 | sample | 0.9696605 | -0.3413429 |
| P2 clWT | G10 | 857890 | sample | 0.5433810 | -5.1373178 |
| P2 clWT | G11 | 1649000 | sample | 1.0444641 | 0.5002562 |
| P2 clWT | G12 | 479370 | SAL | 0.3036293 | -7.8347108 |
| P2 clWT | H01 | 1291300 | empty | 0.8178997 | -2.0487701 |
| P2 clWT | H02 | 1559200 | sample | 0.9875855 | -0.1396727 |
| P2 clWT | H03 | 990660 | sample | 0.6274766 | -4.1911780 |
| P2 clWT | H04 | 1483000 | sample | 0.9393210 | -0.6826858 |
| P2 clWT | H05 | 1520200 | sample | 0.9628832 | -0.4175928 |
| P2 clWT | H06 | 806670 | sample | 0.5109387 | -5.5023196 |
| P2 clWT | H07 | 1311700 | sample | 0.8308209 | -1.9033965 |
| P2 clWT | H08 | 1443300 | sample | 0.9141753 | -0.9655943 |
| P2 clWT | H09 | 1486400 | sample | 0.9414745 | -0.6584569 |
| P2 clWT | H10 | 1605500 | sample | 1.0169116 | 0.1902684 |
| P2 clWT | H11 | 1656900 | sample | 1.0494680 | 0.5565529 |
| P2 clWT | H12 | 409130 | SAL | 0.2591399 | -8.3352521 |
| P3 clWT | A01 | 377220 | SAL | 0.2356594 | -14.9227117 |
| P3 clWT | A02 | 1540100 | sample | 0.9621416 | -0.7391345 |
| P3 clWT | A03 | 1482100 | sample | 0.9259074 | -1.4465570 |
| P3 clWT | A04 | 1392400 | sample | 0.8698694 | -2.5406225 |
| P3 clWT | A05 | 1482400 | sample | 0.9260948 | -1.4428980 |
| P3 clWT | A06 | 1515300 | sample | 0.9466483 | -1.0416186 |
| P3 clWT | A07 | 1443300 | sample | 0.9016680 | -1.9197983 |
| P3 clWT | A08 | 1484500 | sample | 0.9274068 | -1.4172844 |
| P3 clWT | A09 | 1457800 | sample | 0.9107266 | -1.7429427 |
| P3 clWT | A10 | 1588700 | sample | 0.9925033 | -0.1463633 |
| P3 clWT | A11 | 1483900 | sample | 0.9270319 | -1.4246025 |
| P3 clWT | A12 | 1580100 | empty | 0.9871306 | -0.2512570 |
| P3 clWT | B01 | 384500 | SAL | 0.2402074 | -14.8339179 |
| P3 clWT | B02 | 1681000 | sample | 1.0501656 | 0.9794142 |
| P3 clWT | B03 | 1665500 | sample | 1.0404823 | 0.7903617 |
| P3 clWT | B04 | 1573000 | sample | 0.9826951 | -0.3378552 |
| P3 clWT | B05 | 1248200 | sample | 0.7797838 | -4.2994212 |
| P3 clWT | B06 | 1708200 | sample | 1.0671581 | 1.3111710 |
| P3 clWT | B07 | 1627900 | sample | 1.0169926 | 0.3317568 |
| P3 clWT | B08 | 1568500 | sample | 0.9798838 | -0.3927415 |
| P3 clWT | B09 | 1659600 | sample | 1.0367964 | 0.7183997 |
| P3 clWT | B10 | 1718700 | sample | 1.0737177 | 1.4392389 |
| P3 clWT | B11 | 1697500 | sample | 1.0604735 | 1.1806638 |
| P3 clWT | B12 | 1704300 | empty | 1.0647217 | 1.2636029 |
| P3 clWT | C01 | 385990 | SAL | 0.2411383 | -14.8157445 |
| P3 clWT | C02 | 1559000 | sample | 0.9739489 | -0.5086124 |
| P3 clWT | C03 | 1656200 | sample | 1.0346723 | 0.6769301 |
| P3 clWT | C04 | 1685900 | sample | 1.0532267 | 1.0391793 |
| P3 clWT | C05 | 1581900 | sample | 0.9882551 | -0.2293025 |
| P3 clWT | C06 | 1671300 | sample | 1.0441057 | 0.8611039 |
| P3 clWT | C07 | 1599000 | sample | 0.9989380 | -0.0207348 |
| P3 clWT | C08 | 1638200 | sample | 1.0234273 | 0.4573852 |
| P3 clWT | C09 | 1613900 | sample | 1.0082464 | 0.1609996 |
| P3 clWT | C10 | 1752300 | sample | 1.0947086 | 1.8490560 |
| P3 clWT | C11 | 1682500 | sample | 1.0511026 | 0.9977097 |
| P3 clWT | C12 | 1582900 | DMSO | 0.9888799 | -0.2171055 |
| P3 clWT | D01 | 1483400 | DMSO | 0.9267196 | -1.4307010 |
| P3 clWT | D02 | 1575100 | sample | 0.9840070 | -0.3122417 |
| P3 clWT | D03 | 1614500 | sample | 1.0086212 | 0.1683178 |
| P3 clWT | D04 | 1590500 | sample | 0.9936278 | -0.1244088 |
| P3 clWT | D05 | 1553200 | sample | 0.9703255 | -0.5793546 |
| P3 clWT | D06 | 1537300 | sample | 0.9603923 | -0.7732860 |
| P3 clWT | D07 | 1625100 | sample | 1.0152433 | 0.2976053 |
| P3 clWT | D08 | 1602400 | sample | 1.0010620 | 0.0207348 |
| P3 clWT | D09 | 1631600 | sample | 1.0193041 | 0.3768854 |
| P3 clWT | D10 | 1615900 | sample | 1.0094958 | 0.1853935 |

*(continued)*

| Plate | Well | Value | Treatment | Norm | Score |
|---|---|---|---|---|---|
| P3 clWT | D11 | 598280 | sample | 0.3737615 | -12.2264562 |
| P3 clWT | D12 | 1577600 | DMSO | 0.9855688 | -0.2817493 |
| P3 clWT | E01 | 1528400 | DMSO | 0.9548323 | -0.8818387 |
| P3 clWT | E02 | 1618200 | sample | 1.0109327 | 0.2134464 |
| P3 clWT | E03 | 1634000 | sample | 1.0208034 | 0.4061581 |
| P3 clWT | E04 | 1661100 | sample | 1.0377335 | 0.7366952 |
| P3 clWT | E05 | 1655800 | sample | 1.0344224 | 0.6720514 |
| P3 clWT | E06 | 1639700 | sample | 1.0243643 | 0.4756806 |
| P3 clWT | E07 | 1595900 | sample | 0.9970013 | -0.0585453 |
| P3 clWT | E08 | 1513200 | sample | 0.9453364 | -1.0672322 |
| P3 clWT | E09 | 1686900 | sample | 1.0538514 | 1.0513762 |
| P3 clWT | E10 | 1731900 | sample | 1.0819641 | 1.6002385 |
| P3 clWT | E11 | 758110 | sample | 0.4736115 | -10.2770193 |
| P3 clWT | E12 | 1756500 | DMSO | 1.0973324 | 1.9002832 |
| P3 clWT | F01 | 1566500 | DMSO | 0.9786343 | -0.4171353 |
| P3 clWT | F02 | 1650500 | sample | 1.0311114 | 0.6074076 |
| P3 clWT | F03 | 1616000 | sample | 1.0095583 | 0.1866132 |
| P3 clWT | F04 | 1528100 | sample | 0.9546448 | -0.8854978 |
| P3 clWT | F05 | 1627600 | sample | 1.0168051 | 0.3280977 |
| P3 clWT | F06 | 1680600 | sample | 1.0499157 | 0.9745355 |
| P3 clWT | F07 | 1649900 | sample | 1.0307366 | 0.6000894 |
| P3 clWT | F08 | 622910 | sample | 0.3891485 | -11.9260456 |
| P3 clWT | F09 | 1614200 | sample | 1.0084338 | 0.1646587 |
| P3 clWT | F10 | 1628100 | sample | 1.0171175 | 0.3341961 |
| P3 clWT | F11 | 1653700 | sample | 1.0331105 | 0.6464378 |
| P3 clWT | F12 | 527780 | SAL | 0.3297182 | -13.0863404 |
| P3 clWT | G01 | 1518500 | empty | 0.9486475 | -1.0025884 |
| P3 clWT | G02 | 1687300 | sample | 1.0541013 | 1.0562550 |
| P3 clWT | G03 | 931710 | sample | 0.5820641 | -8.1596306 |
| P3 clWT | G04 | 1567800 | sample | 0.9794465 | -0.4012793 |
| P3 clWT | G05 | 1623800 | sample | 1.0144312 | 0.2817493 |
| P3 clWT | G06 | 1576900 | sample | 0.9851315 | -0.2902872 |
| P3 clWT | G07 | 1551900 | sample | 0.9695133 | -0.5952107 |
| P3 clWT | G08 | 1648200 | sample | 1.0296745 | 0.5793546 |
| P3 clWT | G09 | 1568400 | sample | 0.9798213 | -0.3939611 |
| P3 clWT | G10 | 1664000 | sample | 1.0395452 | 0.7720663 |
| P3 clWT | G11 | 1620600 | sample | 1.0124321 | 0.2427191 |
| P3 clWT | G12 | 505480 | SAL | 0.3157868 | -13.3583322 |
| P3 clWT | H01 | 1503800 | empty | 0.9394640 | -1.1818834 |
| P3 clWT | H02 | 1520400 | sample | 0.9498344 | -0.9794142 |
| P3 clWT | H03 | 1521100 | sample | 0.9502718 | -0.9708764 |
| P3 clWT | H04 | 1495900 | sample | 0.9345286 | -1.2782393 |
| P3 clWT | H05 | 1548900 | sample | 0.9676392 | -0.6318015 |
| P3 clWT | H06 | 903400 | sample | 0.5643781 | -8.5049260 |
| P3 clWT | H07 | 994300 | sample | 0.6211657 | -7.3962242 |
| P3 clWT | H08 | 1545200 | sample | 0.9653277 | -0.6769301 |
| P3 clWT | H09 | 1572100 | sample | 0.9821328 | -0.3488325 |
| P3 clWT | H10 | 1576000 | sample | 0.9845693 | -0.3012644 |
| P3 clWT | H11 | 1621700 | sample | 1.0131193 | 0.2561357 |
| P3 clWT | H12 | 450340 | SAL | 0.2813394 | -14.0308714 |
| P4 clWT | A01 | 358270 | SAL | 0.2312314 | -10.2019832 |
| P4 clWT | A02 | 1457600 | sample | 0.9407513 | -0.7862635 |
| P4 clWT | A03 | 1527200 | sample | 0.9856719 | -0.1901422 |
| P4 clWT | A04 | 1414800 | sample | 0.9131277 | -1.1528439 |
| P4 clWT | A05 | 1540400 | sample | 0.9941913 | -0.0770847 |
| P4 clWT | A06 | 1508200 | sample | 0.9734091 | -0.3528764 |
| P4 clWT | A07 | 1567200 | sample | 1.0114883 | 0.1524563 |
| P4 clWT | A08 | 552470 | sample | 0.3565703 | -8.5386676 |
| P4 clWT | A09 | 1476100 | sample | 0.9526914 | -0.6278117 |
| P4 clWT | A10 | 1542300 | sample | 0.9954176 | -0.0608112 |
| P4 clWT | A11 | 1537500 | sample | 0.9923196 | -0.1019230 |
| P4 clWT | A12 | 1692800 | empty | 1.0925520 | 1.2282156 |

*(continued)*

| Plate | Well | Value | Treatment | Norm | Score |
|---|---|---|---|---|---|
| P4 clWT | B01 | 405960 | SAL | 0.2620111 | -9.7935202 |
| P4 clWT | B02 | 1696100 | sample | 1.0946818 | 1.2564799 |
| P4 clWT | B03 | 1540400 | sample | 0.9941913 | -0.0770847 |
| P4 clWT | B04 | 1653900 | sample | 1.0674455 | 0.8950385 |
| P4 clWT | B05 | 1402700 | sample | 0.9053182 | -1.2564799 |
| P4 clWT | B06 | 1628500 | sample | 1.0510520 | 0.6774885 |
| P4 clWT | B07 | 1438000 | sample | 0.9281012 | -0.9541368 |
| P4 clWT | B08 | 1471000 | sample | 0.9493998 | -0.6714930 |
| P4 clWT | B09 | 1677400 | sample | 1.0826126 | 1.0963151 |
| P4 clWT | B10 | 1573700 | sample | 1.0156835 | 0.2081286 |
| P4 clWT | B11 | 1584700 | sample | 1.0227830 | 0.3023432 |
| P4 clWT | B12 | 1751800 | empty | 1.1306312 | 1.7335483 |
| P4 clWT | C01 | 370160 | SAL | 0.2389054 | -10.1001458 |
| P4 clWT | C02 | 1598900 | sample | 1.0319479 | 0.4239656 |
| P4 clWT | C03 | 1652900 | sample | 1.0668001 | 0.8864736 |
| P4 clWT | C04 | 1519300 | sample | 0.9805731 | -0.2578054 |
| P4 clWT | C05 | 1531800 | sample | 0.9886408 | -0.1507433 |
| P4 clWT | C06 | 1623000 | sample | 1.0475023 | 0.6303812 |
| P4 clWT | C07 | 1663900 | sample | 1.0738996 | 0.9806882 |
| P4 clWT | C08 | 973710 | sample | 0.6284433 | -4.9307630 |
| P4 clWT | C09 | 1627300 | sample | 1.0502775 | 0.6672105 |
| P4 clWT | C10 | 1528900 | sample | 0.9867691 | -0.1755817 |
| P4 clWT | C11 | 1429700 | sample | 0.9227443 | -1.0252260 |
| P4 clWT | C12 | 1792700 | DMSO | 1.1570285 | 2.0838553 |
| P4 clWT | D01 | 1628200 | DMSO | 1.0508584 | 0.6749190 |
| P4 clWT | D02 | 1653600 | sample | 1.0672518 | 0.8924690 |
| P4 clWT | D03 | 1561100 | sample | 1.0075513 | 0.1002101 |
| P4 clWT | D04 | 1637800 | sample | 1.0570543 | 0.7571426 |
| P4 clWT | D05 | 1604700 | sample | 1.0356912 | 0.4736424 |
| P4 clWT | D06 | 1567700 | sample | 1.0118110 | 0.1567388 |
| P4 clWT | D07 | 1666200 | sample | 1.0753840 | 1.0003876 |
| P4 clWT | D08 | 1572200 | sample | 1.0147154 | 0.1952811 |
| P4 clWT | D09 | 1664500 | sample | 1.0742868 | 0.9858271 |
| P4 clWT | D10 | 1578700 | sample | 1.0189105 | 0.2509534 |
| P4 clWT | D11 | 1724800 | sample | 1.1132051 | 1.5022943 |
| P4 clWT | D12 | 1647500 | DMSO | 1.0633148 | 0.8402228 |
| P4 clWT | E01 | 1554200 | DMSO | 1.0030980 | 0.0411118 |
| P4 clWT | E02 | 1660700 | sample | 1.0718343 | 0.9532803 |
| P4 clWT | E03 | 1688500 | sample | 1.0897767 | 1.1913862 |
| P4 clWT | E04 | 1010600 | sample | 0.6522525 | -4.6148015 |
| P4 clWT | E05 | 1652800 | sample | 1.0667355 | 0.8856171 |
| P4 clWT | E06 | 1623900 | sample | 1.0480831 | 0.6380897 |
| P4 clWT | E07 | 1406400 | sample | 0.9077062 | -1.2247896 |
| P4 clWT | E08 | 1376300 | sample | 0.8882793 | -1.4825949 |
| P4 clWT | E09 | 1581800 | sample | 1.0209113 | 0.2775048 |
| P4 clWT | E10 | 1686900 | sample | 1.0887440 | 1.1776823 |
| P4 clWT | E11 | 1683500 | sample | 1.0865496 | 1.1485614 |
| P4 clWT | E12 | 1750300 | DMSO | 1.1296631 | 1.7207009 |
| P4 clWT | F01 | 1501600 | DMSO | 0.9691493 | -0.4094052 |
| P4 clWT | F02 | 1630400 | sample | 1.0522783 | 0.6937619 |
| P4 clWT | F03 | 1546500 | sample | 0.9981283 | -0.0248384 |
| P4 clWT | F04 | 1498300 | sample | 0.9670195 | -0.4376696 |
| P4 clWT | F05 | 1427700 | sample | 0.9214535 | -1.0423559 |
| P4 clWT | F06 | 1358400 | sample | 0.8767265 | -1.6359077 |
| P4 clWT | F07 | 1584300 | sample | 1.0225248 | 0.2989172 |
| P4 clWT | F08 | 1000600 | sample | 0.6457984 | -4.7004512 |
| P4 clWT | F09 | 1576600 | sample | 1.0175552 | 0.2329670 |
| P4 clWT | F10 | 1556500 | sample | 1.0045824 | 0.0608112 |
| P4 clWT | F11 | 1627500 | sample | 1.0504066 | 0.6689235 |
| P4 clWT | F12 | 506870 | SAL | 0.3271395 | -8.9292299 |
| P4 clWT | G01 | 1535200 | empty | 0.9908352 | -0.1216225 |

*(continued)*

| Plate | Well | Value | Treatment | Norm | Score |
|---|---|---|---|---|---|
| P4 clWT | G02 | 1613500 | sample | 1.0413709 | 0.5490141 |
| P4 clWT | G03 | 1580400 | sample | 1.0200077 | 0.2655138 |
| P4 clWT | G04 | 1135100 | sample | 0.7326062 | -3.5484638 |
| P4 clWT | G05 | 1535000 | sample | 0.9907061 | -0.1233355 |
| P4 clWT | G06 | 1584500 | sample | 1.0226539 | 0.3006302 |
| P4 clWT | G07 | 918810 | sample | 0.5930102 | -5.4009794 |
| P4 clWT | G08 | 1583000 | sample | 1.0216858 | 0.2877827 |
| P4 clWT | G09 | 716790 | sample | 0.4626242 | -7.1312730 |
| P4 clWT | G10 | 618180 | sample | 0.3989803 | -7.9758639 |
| P4 clWT | G11 | 1621800 | sample | 1.0467278 | 0.6201033 |
| P4 clWT | G12 | 450020 | SAL | 0.2904479 | -9.4161480 |
| P4 clWT | H01 | 1565500 | empty | 1.0103911 | 0.1378959 |
| P4 clWT | H02 | 1226100 | sample | 0.7913386 | -2.7690522 |
| P4 clWT | H03 | 1516000 | sample | 0.9784433 | -0.2860697 |
| P4 clWT | H04 | 239710 | sample | 0.1547115 | -11.2174451 |
| P4 clWT | H05 | 1547100 | sample | 0.9985156 | -0.0196994 |
| P4 clWT | H06 | 1500500 | sample | 0.9684394 | -0.4188266 |
| P4 clWT | H07 | 1551700 | sample | 1.0014844 | 0.0196994 |
| P4 clWT | H08 | 1460700 | sample | 0.9427520 | -0.7597121 |
| P4 clWT | H09 | 58771 | sample | 0.0379315 | -12.7671808 |
| P4 clWT | H10 | 810470 | sample | 0.5230864 | -6.3289074 |
| P4 clWT | H11 | 209340 | sample | 0.1351104 | -11.4775630 |
| P4 clWT | H12 | 370560 | SAL | 0.2391635 | -10.0967198 |
| P5 clWT | A01 | 254570 | SAL | 0.1599008 | -11.6252307 |
| P5 clWT | A02 | 1529100 | sample | 0.9604598 | -0.5471546 |
| P5 clWT | A03 | 1605100 | sample | 1.0081970 | 0.1134292 |
| P5 clWT | A04 | 1302600 | sample | 0.8181904 | -2.5158679 |
| P5 clWT | A05 | 1591200 | sample | 0.9994661 | -0.0073881 |
| P5 clWT | A06 | 1682300 | sample | 1.0566879 | 0.7844432 |
| P5 clWT | A07 | 1558900 | sample | 0.9791778 | -0.2881362 |
| P5 clWT | A08 | 1505200 | sample | 0.9454477 | -0.7548908 |
| P5 clWT | A09 | 1509900 | sample | 0.9483999 | -0.7140389 |
| P5 clWT | A10 | 1653600 | sample | 1.0386608 | 0.5349859 |
| P5 clWT | A11 | 1654700 | sample | 1.0393518 | 0.5445470 |
| P5 clWT | A12 | 1627800 | empty | 1.0224553 | 0.3107351 |
| P5 clWT | B01 | 351330 | SAL | 0.2206777 | -10.7842033 |
| P5 clWT | B02 | 1641400 | sample | 1.0309978 | 0.4289448 |
| P5 clWT | B03 | 1631300 | sample | 1.0246537 | 0.3411567 |
| P5 clWT | B04 | 1669200 | sample | 1.0484595 | 0.6705794 |
| P5 clWT | B05 | 1681400 | sample | 1.0561226 | 0.7766205 |
| P5 clWT | B06 | 1780900 | sample | 1.1186206 | 1.6414637 |
| P5 clWT | B07 | 1442500 | sample | 0.9060645 | -1.2998723 |
| P5 clWT | B08 | 1608300 | sample | 1.0102070 | 0.1412432 |
| P5 clWT | B09 | 1729200 | sample | 1.0861468 | 1.1920929 |
| P5 clWT | B10 | 1770800 | sample | 1.1122766 | 1.5536756 |
| P5 clWT | B11 | 1611800 | sample | 1.0124054 | 0.1716649 |
| P5 clWT | B12 | 1688600 | empty | 1.0606451 | 0.8392021 |
| P5 clWT | C01 | 371690 | SAL | 0.2334663 | -10.6072364 |
| P5 clWT | C02 | 1553100 | sample | 0.9755347 | -0.3385492 |
| P5 clWT | C03 | 1620200 | sample | 1.0176816 | 0.2446767 |
| P5 clWT | C04 | 1427900 | sample | 0.8968939 | -1.4267739 |
| P5 clWT | C05 | 1599400 | sample | 1.0046167 | 0.0638854 |
| P5 clWT | C06 | 1700000 | sample | 1.0678057 | 0.9382897 |
| P5 clWT | C07 | 1509100 | sample | 0.9478974 | -0.7209924 |
| P5 clWT | C08 | 1611700 | sample | 1.0123426 | 0.1707957 |
| P5 clWT | C09 | 1205300 | sample | 0.7570742 | -3.3615889 |
| P5 clWT | C10 | 1682900 | sample | 1.0570648 | 0.7896583 |
| P5 clWT | C11 | 1596800 | sample | 1.0029836 | 0.0412865 |
| P5 clWT | C12 | 1684100 | DMSO | 1.0578185 | 0.8000886 |
| P5 clWT | D01 | 1342000 | DMSO | 0.8429383 | -2.1734074 |
| P5 clWT | D02 | 1599200 | sample | 1.0044911 | 0.0621470 |
| P5 clWT | D03 | 1664900 | sample | 1.0457586 | 0.6332043 |

*(continued)*

| Plate | Well | Value | Treatment | Norm | Score |
|---|---|---|---|---|---|
| P5 clWT | D04 | 1592900 | sample | 1.0005339 | 0.0073881 |
| P5 clWT | D05 | 1539100 | sample | 0.9667410 | -0.4602356 |
| P5 clWT | D06 | 904050 | sample | 0.5678528 | -5.9800212 |
| P5 clWT | D07 | 1736400 | sample | 1.0906693 | 1.2546745 |
| P5 clWT | D08 | 1551100 | sample | 0.9742784 | -0.3559329 |
| P5 clWT | D09 | 1704300 | sample | 1.0705066 | 0.9756648 |
| P5 clWT | D10 | 1562000 | sample | 0.9811250 | -0.2611913 |
| P5 clWT | D11 | 1671100 | sample | 1.0496530 | 0.6870940 |
| P5 clWT | D12 | 1674200 | DMSO | 1.0516001 | 0.7140389 |
| P5 clWT | E01 | 1505700 | DMSO | 0.9457618 | -0.7505448 |
| P5 clWT | E02 | 1670100 | sample | 1.0490248 | 0.6784021 |
| P5 clWT | E03 | 1665000 | sample | 1.0458214 | 0.6340735 |
| P5 clWT | E04 | 480350 | sample | 0.3017179 | -9.6627755 |
| P5 clWT | E05 | 1321100 | sample | 0.8298106 | -2.3550679 |
| P5 clWT | E06 | 1632500 | sample | 1.0254075 | 0.3515870 |
| P5 clWT | E07 | 1564900 | sample | 0.9829465 | -0.2359848 |
| P5 clWT | E08 | 1485800 | sample | 0.9332621 | -0.9235134 |
| P5 clWT | E09 | 1522800 | sample | 0.9565026 | -0.6019135 |
| P5 clWT | E10 | 1636100 | sample | 1.0276687 | 0.3828778 |
| P5 clWT | E11 | 1665200 | sample | 1.0459470 | 0.6358118 |
| P5 clWT | E12 | 1691400 | DMSO | 1.0624038 | 0.8635394 |
| P5 clWT | F01 | 1456500 | DMSO | 0.9148582 | -1.1781859 |
| P5 clWT | F02 | 1699500 | sample | 1.0674916 | 0.9339437 |
| P5 clWT | F03 | 1638700 | sample | 1.0293018 | 0.4054767 |
| P5 clWT | F04 | 114310 | sample | 0.0718005 | -12.8443553 |
| P5 clWT | F05 | 1595600 | sample | 1.0022298 | 0.0308562 |
| P5 clWT | F06 | 1000400 | sample | 0.6283722 | -5.1425574 |
| P5 clWT | F07 | 1167500 | sample | 0.7333312 | -3.6901424 |
| P5 clWT | F08 | 1569200 | sample | 0.9856474 | -0.1986097 |
| P5 clWT | F09 | 1603700 | sample | 1.0073176 | 0.1012605 |
| P5 clWT | F10 | 1657400 | sample | 1.0410477 | 0.5680151 |
| P5 clWT | F11 | 1698900 | sample | 1.0671147 | 0.9287286 |
| P5 clWT | F12 | 489750 | SAL | 0.3076222 | -9.5810717 |
| P5 clWT | G01 | 1298300 | empty | 0.8154895 | -2.5532430 |
| P5 clWT | G02 | 1596300 | sample | 1.0026695 | 0.0369405 |
| P5 clWT | G03 | 1453400 | sample | 0.9129110 | -1.2051307 |
| P5 clWT | G04 | 1382300 | sample | 0.8682516 | -1.8231242 |
| P5 clWT | G05 | 1544000 | sample | 0.9698188 | -0.4176454 |
| P5 clWT | G06 | 123420 | sample | 0.0775227 | -12.7651722 |
| P5 clWT | G07 | 856670 | sample | 0.5380924 | -6.3918430 |
| P5 clWT | G08 | 1166200 | sample | 0.7325147 | -3.7014419 |
| P5 clWT | G09 | 1495800 | sample | 0.9395434 | -0.8365945 |
| P5 clWT | G10 | 1318900 | sample | 0.8284288 | -2.3741901 |
| P5 clWT | G11 | 1600800 | sample | 1.0054961 | 0.0760540 |
| P5 clWT | G12 | 540580 | SAL | 0.3395496 | -9.1392629 |
| P5 clWT | H01 | 1276200 | empty | 0.8016080 | -2.7453338 |
| P5 clWT | H02 | 1535500 | sample | 0.9644798 | -0.4915264 |
| P5 clWT | H03 | 1516900 | sample | 0.9527967 | -0.6531956 |
| P5 clWT | H04 | 1479500 | sample | 0.9293050 | -0.9782724 |
| P5 clWT | H05 | 1660200 | sample | 1.0428064 | 0.5923524 |
| P5 clWT | H06 | 1580700 | sample | 0.9928708 | -0.0986530 |
| P5 clWT | H07 | 816780 | sample | 0.5130367 | -6.7385625 |
| P5 clWT | H08 | 99622 | sample | 0.0625747 | -12.9720218 |
| P5 clWT | H09 | 1512400 | sample | 0.9499702 | -0.6923091 |
| P5 clWT | H10 | 295400 | sample | 0.1855469 | -11.2703408 |
| P5 clWT | H11 | 1708600 | sample | 1.0732075 | 1.0130399 |
| P5 clWT | H12 | 431020 | SAL | 0.2707327 | -10.0915465 |
| P6 clWT | A01 | 281910 | SAL | 0.1778444 | -9.9268587 |
| P6 clWT | A02 | 1544300 | sample | 0.9742296 | -0.3111569 |
| P6 clWT | A03 | 1372700 | sample | 0.8659748 | -1.6182446 |
| P6 clWT | A04 | 1292500 | sample | 0.8153802 | -2.2291329 |
| P6 clWT | A05 | 923290 | sample | 0.5824622 | -5.0414281 |

*(continued)*

| Plate | Well | Value | Treatment | Norm | Score |
|---|---|---|---|---|---|
| P6 clWT | A06 | 1413900 | sample | 0.8919661 | -1.3044217 |
| P6 clWT | A07 | 1424600 | sample | 0.8987162 | -1.2229192 |
| P6 clWT | A08 | 1482900 | sample | 0.9354951 | -0.7788445 |
| P6 clWT | A09 | 1350600 | sample | 0.8520329 | -1.7865817 |
| P6 clWT | A10 | 582350 | sample | 0.3673785 | -7.6383889 |
| P6 clWT | A11 | 1375600 | sample | 0.8678043 | -1.5961552 |
| P6 clWT | A12 | 1645700 | empty | 1.0381983 | 0.4612130 |
| P6 clWT | B01 | 1458500 | SAL | 0.9201022 | -0.9647008 |
| P6 clWT | B02 | 1631700 | sample | 1.0293663 | 0.3545742 |
| P6 clWT | B03 | 1543300 | sample | 0.9735987 | -0.3187740 |
| P6 clWT | B04 | 1496800 | sample | 0.9442639 | -0.6729673 |
| P6 clWT | B05 | 1421500 | sample | 0.8967606 | -1.2465320 |
| P6 clWT | B06 | 1662100 | sample | 1.0485443 | 0.5861329 |
| P6 clWT | B07 | 1512200 | sample | 0.9539791 | -0.5556646 |
| P6 clWT | B08 | 1543600 | sample | 0.9737880 | -0.3164889 |
| P6 clWT | B09 | 1673000 | sample | 1.0554206 | 0.6691588 |
| P6 clWT | B10 | 1756600 | sample | 1.1081601 | 1.3059451 |
| P6 clWT | B11 | 1718200 | sample | 1.0839353 | 1.0134500 |
| P6 clWT | B12 | 1806000 | empty | 1.1393244 | 1.6822279 |
| P6 clWT | C01 | 1522800 | SAL | 0.9606662 | -0.4749238 |
| P6 clWT | C02 | 1314100 | sample | 0.8290067 | -2.0646044 |
| P6 clWT | C03 | 1364900 | sample | 0.8610542 | -1.6776577 |
| P6 clWT | C04 | 1474300 | sample | 0.9300697 | -0.8443512 |
| P6 clWT | C05 | 1387500 | sample | 0.8753115 | -1.5055121 |
| P6 clWT | C06 | 1253600 | sample | 0.7908400 | -2.5254366 |
| P6 clWT | C07 | 1623200 | sample | 1.0240040 | 0.2898292 |
| P6 clWT | C08 | 1551600 | sample | 0.9788348 | -0.2555524 |
| P6 clWT | C09 | 1662800 | sample | 1.0489859 | 0.5914648 |
| P6 clWT | C10 | 1745200 | sample | 1.1009684 | 1.2191106 |
| P6 clWT | C11 | 1653100 | sample | 1.0428666 | 0.5175793 |
| P6 clWT | C12 | 1683300 | DMSO | 1.0619184 | 0.7476145 |
| P6 clWT | D01 | 1479900 | DMSO | 0.9336025 | -0.8016957 |
| P6 clWT | D02 | 1560600 | sample | 0.9845125 | -0.1869988 |
| P6 clWT | D03 | 1662400 | sample | 1.0487336 | 0.5884180 |
| P6 clWT | D04 | 841400 | sample | 0.5308015 | -5.6651892 |
| P6 clWT | D05 | 1770200 | sample | 1.1167397 | 1.4095372 |
| P6 clWT | D06 | 1643000 | sample | 1.0364950 | 0.4406470 |
| P6 clWT | D07 | 1686700 | sample | 1.0640633 | 0.7735126 |
| P6 clWT | D08 | 1668100 | sample | 1.0523294 | 0.6318352 |
| P6 clWT | D09 | 269740 | sample | 0.1701669 | -10.0195583 |
| P6 clWT | D10 | 1592600 | sample | 1.0046999 | 0.0567471 |
| P6 clWT | D11 | 1577800 | sample | 0.9953632 | -0.0559854 |
| P6 clWT | D12 | 1651700 | DMSO | 1.0419834 | 0.5069154 |
| P6 clWT | E01 | 1641700 | DMSO | 1.0356749 | 0.4307448 |
| P6 clWT | E02 | 1641000 | sample | 1.0352333 | 0.4254129 |
| P6 clWT | E03 | 1629100 | sample | 1.0277261 | 0.3347698 |
| P6 clWT | E04 | 1322500 | sample | 0.8343059 | -2.0006211 |
| P6 clWT | E05 | 1685000 | sample | 1.0629909 | 0.7605635 |
| P6 clWT | E06 | 1624200 | sample | 1.0246349 | 0.2974462 |
| P6 clWT | E07 | 1660800 | sample | 1.0477242 | 0.5762307 |
| P6 clWT | E08 | 1397700 | sample | 0.8817462 | -1.4278181 |
| P6 clWT | E09 | 390880 | sample | 0.2465887 | -9.0968275 |
| P6 clWT | E10 | 1685100 | sample | 1.0630540 | 0.7613253 |
| P6 clWT | E11 | 1745100 | sample | 1.1009053 | 1.2183489 |
| P6 clWT | E12 | 1731700 | DMSO | 1.0924518 | 1.1162803 |
| P6 clWT | F01 | 1502100 | DMSO | 0.9476075 | -0.6325969 |
| P6 clWT | F02 | 1674500 | sample | 1.0563669 | 0.6805844 |
| P6 clWT | F03 | 1695100 | sample | 1.0693625 | 0.8374959 |
| P6 clWT | F04 | 1496400 | sample | 0.9440116 | -0.6760142 |
| P6 clWT | F05 | 1629400 | sample | 1.0279153 | 0.3370550 |
| P6 clWT | F06 | 1224300 | sample | 0.7723559 | -2.7486165 |

*(continued)*

| Plate | Well | Value | Treatment | Norm | Score |
|---|---|---|---|---|---|
| P6 clWT | F07 | 1419700 | sample | 0.8956250 | -1.2602428 |
| P6 clWT | F08 | 1652300 | sample | 1.0423619 | 0.5114857 |
| P6 clWT | F09 | 1630400 | sample | 1.0285462 | 0.3446720 |
| P6 clWT | F10 | 1718800 | sample | 1.0843138 | 1.0180202 |
| P6 clWT | F11 | 1641800 | sample | 1.0357379 | 0.4315065 |
| P6 clWT | F12 | 424500 | SAL | 0.2677980 | -8.8407420 |
| P6 clWT | G01 | 1505600 | empty | 0.9498155 | -0.6059372 |
| P6 clWT | G02 | 1562200 | sample | 0.9855219 | -0.1748116 |
| P6 clWT | G03 | 1688700 | sample | 1.0653250 | 0.7887467 |
| P6 clWT | G04 | 1609700 | sample | 1.0154875 | 0.1869988 |
| P6 clWT | G05 | 1507300 | sample | 0.9508879 | -0.5929882 |
| P6 clWT | G06 | 1543500 | sample | 0.9737249 | -0.3172506 |
| P6 clWT | G07 | 1531400 | sample | 0.9660915 | -0.4094170 |
| P6 clWT | G08 | 1560600 | sample | 0.9845125 | -0.1869988 |
| P6 clWT | G09 | 1616300 | sample | 1.0196511 | 0.2372715 |
| P6 clWT | G10 | 1718900 | sample | 1.0843769 | 1.0187819 |
| P6 clWT | G11 | 1603600 | sample | 1.0116393 | 0.1405348 |
| P6 clWT | G12 | 421960 | SAL | 0.2661956 | -8.8600893 |
| P6 clWT | H01 | 1436700 | empty | 0.9063496 | -1.1307527 |
| P6 clWT | H02 | 1592500 | sample | 1.0046368 | 0.0559854 |
| P6 clWT | H03 | 1660600 | sample | 1.0475980 | 0.5747073 |
| P6 clWT | H04 | 1609400 | sample | 1.0152982 | 0.1847137 |
| P6 clWT | H05 | 1609200 | sample | 1.0151721 | 0.1831903 |
| P6 clWT | H06 | 1452500 | sample | 0.9163171 | -1.0104032 |
| P6 clWT | H07 | 1297000 | sample | 0.8182191 | -2.1948562 |
| P6 clWT | H08 | 1257500 | sample | 0.7933003 | -2.4957301 |
| P6 clWT | H09 | 989120 | sample | 0.6239914 | -4.5399969 |
| P6 clWT | H10 | 1638000 | sample | 1.0333407 | 0.4025617 |
| P6 clWT | H11 | 1654400 | sample | 1.0436867 | 0.5274815 |
| P6 clWT | H12 | 385980 | SAL | 0.2434975 | -9.1341511 |
| P7 clWT | A01 | 314080 | SAL | 0.1945370 | -15.0967517 |
| P7 clWT | A02 | 1620600 | sample | 1.0037783 | 0.0708157 |
| P7 clWT | A03 | 1532000 | sample | 0.9489006 | -0.9577537 |
| P7 clWT | A04 | 1543600 | sample | 0.9560855 | -0.8230877 |
| P7 clWT | A05 | 1608500 | sample | 0.9962837 | -0.0696548 |
| P7 clWT | A06 | 1681300 | sample | 1.0413750 | 0.7754902 |
| P7 clWT | A07 | 1575800 | sample | 0.9760297 | -0.4492735 |
| P7 clWT | A08 | 123030 | sample | 0.0762032 | -17.3146770 |
| P7 clWT | A09 | 1450100 | sample | 0.8981728 | -1.9085418 |
| P7 clWT | A10 | 1551900 | sample | 0.9612264 | -0.7267319 |
| P7 clWT | A11 | 1544600 | sample | 0.9567049 | -0.8114786 |
| P7 clWT | A12 | 1676900 | empty | 1.0386497 | 0.7244100 |
| P7 clWT | B01 | 381230 | SAL | 0.2361288 | -14.3171983 |
| P7 clWT | B02 | 1145000 | sample | 0.7091979 | -5.4504890 |
| P7 clWT | B03 | 1641200 | sample | 1.0165376 | 0.3099639 |
| P7 clWT | B04 | 1653300 | sample | 1.0240322 | 0.4504344 |
| P7 clWT | B05 | 1653000 | sample | 1.0238464 | 0.4469517 |
| P7 clWT | B06 | 1505300 | sample | 0.9323630 | -1.2677176 |
| P7 clWT | B07 | 1613600 | sample | 0.9994426 | -0.0104482 |
| P7 clWT | B08 | 1565100 | sample | 0.9694023 | -0.5734913 |
| P7 clWT | B09 | 1547100 | sample | 0.9582533 | -0.7824557 |
| P7 clWT | B10 | 1565500 | sample | 0.9696500 | -0.5688476 |
| P7 clWT | B11 | 1710700 | sample | 1.0595850 | 1.1167988 |
| P7 clWT | B12 | 1800900 | empty | 1.1154537 | 2.1639428 |
| P7 clWT | C01 | 433660 | SAL | 0.2686033 | -13.7085313 |
| P7 clWT | C02 | 1659900 | sample | 1.0281202 | 0.5270547 |
| P7 clWT | C03 | 1484000 | sample | 0.9191700 | -1.5149922 |
| P7 clWT | C04 | 1729500 | sample | 1.0712295 | 1.3350506 |
| P7 clWT | C05 | 1691200 | sample | 1.0475070 | 0.8904207 |
| P7 clWT | C06 | 1712600 | sample | 1.0607618 | 1.1388562 |
| P7 clWT | C07 | 1663500 | sample | 1.0303500 | 0.5688476 |
| P7 clWT | C08 | 1638200 | sample | 1.0146795 | 0.2751365 |

*(continued)*

| Plate | Well | Value | Treatment | Norm | Score |
|---|---|---|---|---|---|
| P7 clWT | C09 | 1563800 | sample | 0.9685971 | -0.5885832 |
| P7 clWT | C10 | 1283000 | sample | 0.7946733 | -3.8484283 |
| P7 clWT | C11 | 1672500 | sample | 1.0359244 | 0.6733298 |
| P7 clWT | C12 | 1658000 | DMSO | 1.0269433 | 0.5049974 |
| P7 clWT | D01 | 1533300 | DMSO | 0.9497058 | -0.9426618 |
| P7 clWT | D02 | 1358700 | sample | 0.8415609 | -2.9696168 |
| P7 clWT | D03 | 1571500 | sample | 0.9733664 | -0.4991928 |
| P7 clWT | D04 | 1499600 | sample | 0.9288325 | -1.3338896 |
| P7 clWT | D05 | 1631800 | sample | 1.0107154 | 0.2008380 |
| P7 clWT | D06 | 1608400 | sample | 0.9962217 | -0.0708157 |
| P7 clWT | D07 | 801070 | sample | 0.4961722 | -9.4432189 |
| P7 clWT | D08 | 1667800 | sample | 1.0330133 | 0.6187669 |
| P7 clWT | D09 | 1719800 | sample | 1.0652214 | 1.2224419 |
| P7 clWT | D10 | 1641100 | sample | 1.0164757 | 0.3088030 |
| P7 clWT | D11 | 1690200 | sample | 1.0468876 | 0.8788115 |
| P7 clWT | D12 | 1539800 | DMSO | 0.9537318 | -0.8672024 |
| P7 clWT | E01 | 1518300 | DMSO | 0.9404150 | -1.1167988 |
| P7 clWT | E02 | 1694600 | sample | 1.0496129 | 0.9298917 |
| P7 clWT | E03 | 282600 | sample | 0.1750387 | -15.4622073 |
| P7 clWT | E04 | 1663700 | sample | 1.0304738 | 0.5711695 |
| P7 clWT | E05 | 1638000 | sample | 1.0145556 | 0.2728147 |
| P7 clWT | E06 | 1683100 | sample | 1.0424899 | 0.7963867 |
| P7 clWT | E07 | 1654700 | sample | 1.0248993 | 0.4666872 |
| P7 clWT | E08 | 1556300 | sample | 0.9639517 | -0.6756517 |
| P7 clWT | E09 | 1690300 | sample | 1.0469495 | 0.8799725 |
| P7 clWT | E10 | 1733700 | sample | 1.0738309 | 1.3838089 |
| P7 clWT | E11 | 1674400 | sample | 1.0371013 | 0.6953872 |
| P7 clWT | E12 | 1756200 | DMSO | 1.0877671 | 1.6450145 |
| P7 clWT | F01 | 1568500 | DMSO | 0.9715082 | -0.5340202 |
| P7 clWT | F02 | 1611300 | sample | 0.9980180 | -0.0371492 |
| P7 clWT | F03 | 1665700 | sample | 1.0317126 | 0.5943877 |
| P7 clWT | F04 | 1616800 | sample | 1.0014246 | 0.0267010 |
| P7 clWT | F05 | 1682900 | sample | 1.0423661 | 0.7940649 |
| P7 clWT | F06 | 1616800 | sample | 1.0014246 | 0.0267010 |
| P7 clWT | F07 | 1625900 | sample | 1.0070610 | 0.1323441 |
| P7 clWT | F08 | 1664100 | sample | 1.0307216 | 0.5758131 |
| P7 clWT | F09 | 1616400 | sample | 1.0011768 | 0.0220574 |
| P7 clWT | F10 | 1615400 | sample | 1.0005574 | 0.0104482 |
| P7 clWT | F11 | 1748300 | sample | 1.0828740 | 1.5533023 |
| P7 clWT | F12 | 534360 | SAL | 0.3309755 | -12.5394914 |
| P7 clWT | G01 | 1429400 | empty | 0.8853515 | -2.1488509 |
| P7 clWT | G02 | 166550 | sample | 0.1031589 | -16.8094474 |
| P7 clWT | G03 | 1629100 | sample | 1.0090430 | 0.1694934 |
| P7 clWT | G04 | 1584100 | sample | 0.9811706 | -0.3529177 |
| P7 clWT | G05 | 1598500 | sample | 0.9900898 | -0.1857462 |
| P7 clWT | G06 | 1607300 | sample | 0.9955404 | -0.0835858 |
| P7 clWT | G07 | 1547600 | sample | 0.9585630 | -0.7766511 |
| P7 clWT | G08 | 1672100 | sample | 1.0356767 | 0.6686862 |
| P7 clWT | G09 | 1522900 | sample | 0.9432642 | -1.0633968 |
| P7 clWT | G10 | 1669400 | sample | 1.0340043 | 0.6373415 |
| P7 clWT | G11 | 1769000 | sample | 1.0956953 | 1.7936114 |
| P7 clWT | G12 | 487770 | SAL | 0.3021183 | -13.0803610 |
| P7 clWT | H01 | 1337500 | empty | 0.8284299 | -3.2157305 |
| P7 clWT | H02 | 1536800 | sample | 0.9518736 | -0.9020298 |
| P7 clWT | H03 | 1569900 | sample | 0.9723753 | -0.5177674 |
| P7 clWT | H04 | 1404200 | sample | 0.8697430 | -2.4414011 |
| P7 clWT | H05 | 1610200 | sample | 0.9973366 | -0.0499193 |
| P7 clWT | H06 | 1491900 | sample | 0.9240632 | -1.4232800 |
| P7 clWT | H07 | 1519700 | sample | 0.9412821 | -1.1005460 |
| P7 clWT | H08 | 1600900 | sample | 0.9915763 | -0.1578842 |
| P7 clWT | H09 | 1559800 | sample | 0.9661195 | -0.6350197 |
| P7 clWT | H10 | 1674700 | sample | 1.0372871 | 0.6988699 |

*(continued)*

| Plate | Well | Value | Treatment | Norm | Score |
|---|---|---|---|---|---|
| P7 clWT | H11 | 1320600 | sample | 0.8179622 | -3.4119249 |
| P7 clWT | H12 | 425950 | SAL | 0.2638278 | -13.7980377 |
| P8 clWT | A01 | 336200 | SAL | 0.2021952 | -14.5133125 |
| P8 clWT | A02 | 1495900 | sample | 0.8996542 | -1.8254466 |
| P8 clWT | A03 | 1111400 | sample | 0.6684108 | -6.0321246 |
| P8 clWT | A04 | 1582200 | sample | 0.9515562 | -0.8812689 |
| P8 clWT | A05 | 1665600 | sample | 1.0017140 | 0.0311808 |
| P8 clWT | A06 | 1598900 | sample | 0.9615998 | -0.6985602 |
| P8 clWT | A07 | 1576500 | sample | 0.9481281 | -0.9436306 |
| P8 clWT | A08 | 1117300 | sample | 0.6719591 | -5.9675748 |
| P8 clWT | A09 | 1508400 | sample | 0.9071719 | -1.6886885 |
| P8 clWT | A10 | 1552300 | sample | 0.9335739 | -1.2083942 |
| P8 clWT | A11 | 1672900 | sample | 1.0061043 | 0.1110475 |
| P8 clWT | A12 | 1827500 | empty | 1.0990828 | 1.8024713 |
| P8 clWT | B01 | 401260 | SAL | 0.2413231 | -13.8015142 |
| P8 clWT | B02 | 1636200 | sample | 0.9840325 | -0.2904741 |
| P8 clWT | B03 | 1793300 | sample | 1.0785145 | 1.4283012 |
| P8 clWT | B04 | 1778700 | sample | 1.0697339 | 1.2685678 |
| P8 clWT | B05 | 1737500 | sample | 1.0449556 | 0.8178132 |
| P8 clWT | B06 | 1682500 | sample | 1.0118779 | 0.2160777 |
| P8 clWT | B07 | 1618500 | sample | 0.9733875 | -0.4841235 |
| P8 clWT | B08 | 1620100 | sample | 0.9743497 | -0.4666185 |
| P8 clWT | B09 | 1692600 | sample | 1.0179522 | 0.3265783 |
| P8 clWT | B10 | 1677300 | sample | 1.0087506 | 0.1591864 |
| P8 clWT | B11 | 1730200 | sample | 1.0405653 | 0.7379465 |
| P8 clWT | B12 | 1866100 | empty | 1.1222974 | 2.2247801 |
| P8 clWT | C01 | 454350 | SAL | 0.2732521 | -13.2206753 |
| P8 clWT | C02 | 1698400 | sample | 1.0214404 | 0.3900340 |
| P8 clWT | C03 | 1702400 | sample | 1.0238460 | 0.4337966 |
| P8 clWT | C04 | 1419900 | sample | 0.8539468 | -2.6569356 |
| P8 clWT | C05 | 1773900 | sample | 1.0668471 | 1.2160527 |
| P8 clWT | C06 | 1720400 | sample | 1.0346715 | 0.6307282 |
| P8 clWT | C07 | 1518100 | sample | 0.9130056 | -1.5825643 |
| P8 clWT | C08 | 1625400 | sample | 0.9775372 | -0.4086331 |
| P8 clWT | C09 | 1651900 | sample | 0.9934747 | -0.1187060 |
| P8 clWT | C10 | 1208400 | sample | 0.7267479 | -4.9708820 |
| P8 clWT | C11 | 1753500 | sample | 1.0545783 | 0.9928635 |
| P8 clWT | C12 | 1794200 | DMSO | 1.0790558 | 1.4381478 |
| P8 clWT | D01 | 1692500 | DMSO | 1.0178920 | 0.3254842 |
| P8 clWT | D02 | 1687000 | sample | 1.0145843 | 0.2653106 |
| P8 clWT | D03 | 1660100 | sample | 0.9984063 | -0.0289927 |
| P8 clWT | D04 | 1681300 | sample | 1.0111562 | 0.2029490 |
| P8 clWT | D05 | 1736000 | sample | 1.0440535 | 0.8014022 |
| P8 clWT | D06 | 1605700 | sample | 0.9656894 | -0.6241638 |
| P8 clWT | D07 | 1628500 | sample | 0.9794016 | -0.3747171 |
| P8 clWT | D08 | 1705800 | sample | 1.0258908 | 0.4709948 |
| P8 clWT | D09 | 1712500 | sample | 1.0299203 | 0.5442971 |
| P8 clWT | D10 | 1707500 | sample | 1.0269132 | 0.4895939 |
| P8 clWT | D11 | 1754400 | sample | 1.0551195 | 1.0027101 |
| P8 clWT | D12 | 1690900 | DMSO | 1.0169298 | 0.3079792 |
| P8 clWT | E01 | 1725500 | DMSO | 1.0377387 | 0.6865255 |
| P8 clWT | E02 | 1773600 | sample | 1.0666667 | 1.2127705 |
| P8 clWT | E03 | 1780200 | sample | 1.0706360 | 1.2849787 |
| P8 clWT | E04 | 1746500 | sample | 1.0503684 | 0.9162790 |
| P8 clWT | E05 | 1763000 | sample | 1.0602917 | 1.0967997 |
| P8 clWT | E06 | 1693400 | sample | 1.0184333 | 0.3353308 |
| P8 clWT | E07 | 1651700 | sample | 0.9933544 | -0.1208941 |
| P8 clWT | E08 | 1558100 | sample | 0.9370621 | -1.1449385 |
| P8 clWT | E09 | 969440 | sample | 0.5830341 | -7.5852585 |
| P8 clWT | E10 | 1686300 | sample | 1.0141633 | 0.2576522 |
| P8 clWT | E11 | 1636200 | sample | 0.9840325 | -0.2904741 |

*(continued)*

| Plate | Well | Value | Treatment | Norm | Score |
|---|---|---|---|---|---|
| P8 clWT | E12 | 1703200 | DMSO | 1.0243272 | 0.4425491 |
| P8 clWT | F01 | 1634000 | DMSO | 0.9827094 | -0.3145435 |
| P8 clWT | F02 | 1662300 | sample | 0.9997294 | -0.0049233 |
| P8 clWT | F03 | 1722200 | sample | 1.0357540 | 0.6504213 |
| P8 clWT | F04 | 1707400 | sample | 1.0268531 | 0.4884998 |
| P8 clWT | F05 | 1722000 | sample | 1.0356337 | 0.6482332 |
| P8 clWT | F06 | 1747600 | sample | 1.0510299 | 0.9283137 |
| P8 clWT | F07 | 1594000 | sample | 0.9586528 | -0.7521693 |
| P8 clWT | F08 | 1731800 | sample | 1.0415276 | 0.7554515 |
| P8 clWT | F09 | 1663200 | sample | 1.0002706 | 0.0049233 |
| P8 clWT | F10 | 1686000 | sample | 1.0139829 | 0.2543700 |
| P8 clWT | F11 | 1783900 | sample | 1.0728612 | 1.3254591 |
| P8 clWT | F12 | 562040 | SAL | 0.3380183 | -12.0424773 |
| P8 clWT | G01 | 1545100 | empty | 0.9292437 | -1.2871669 |
| P8 clWT | G02 | 1715600 | sample | 1.0317847 | 0.5782131 |
| P8 clWT | G03 | 1687300 | sample | 1.0147647 | 0.2685928 |
| P8 clWT | G04 | 1688300 | sample | 1.0153661 | 0.2795335 |
| P8 clWT | G05 | 1596800 | sample | 0.9603368 | -0.7215355 |
| P8 clWT | G06 | 403910 | sample | 0.2429169 | -13.7725215 |
| P8 clWT | G07 | 1598700 | sample | 0.9614795 | -0.7007483 |
| P8 clWT | G08 | 1704100 | sample | 1.0248684 | 0.4523957 |
| P8 clWT | G09 | 1637500 | sample | 0.9848143 | -0.2762513 |
| P8 clWT | G10 | 1626900 | sample | 0.9784393 | -0.3922221 |
| P8 clWT | G11 | 1322400 | sample | 0.7953090 | -3.7236485 |
| P8 clWT | G12 | 488040 | SAL | 0.2935138 | -12.8520850 |
| P8 clWT | H01 | 1537100 | empty | 0.9244324 | -1.3746920 |
| P8 clWT | H02 | 1671200 | sample | 1.0050819 | 0.0924484 |
| P8 clWT | H03 | 1542400 | sample | 0.9276199 | -1.3167066 |
| P8 clWT | H04 | 1617600 | sample | 0.9728462 | -0.4939701 |
| P8 clWT | H05 | 1561700 | sample | 0.9392272 | -1.1055522 |
| P8 clWT | H06 | 1513400 | sample | 0.9101789 | -1.6339853 |
| P8 clWT | H07 | 1581100 | sample | 0.9508946 | -0.8933037 |
| P8 clWT | H08 | 1544900 | sample | 0.9291234 | -1.2893550 |
| P8 clWT | H09 | 1510100 | sample | 0.9081943 | -1.6700894 |
| P8 clWT | H10 | 1653300 | sample | 0.9943166 | -0.1033891 |
| P8 clWT | H11 | 111510 | sample | 0.0670636 | -16.9715660 |
| P8 clWT | H12 | 427270 | SAL | 0.2569659 | -13.5169480 |
| P9 clWT | A01 | 294700 | SAL | 0.1856027 | -19.0849891 |
| P9 clWT | A02 | 1507100 | sample | 0.9491750 | -1.1910592 |
| P9 clWT | A03 | 740270 | sample | 0.4662237 | -12.5087780 |
| P9 clWT | A04 | 1463400 | sample | 0.9216526 | -1.8360317 |
| P9 clWT | A05 | 1672600 | sample | 1.0534072 | 1.2515715 |
| P9 clWT | A06 | 1718400 | sample | 1.0822522 | 1.9275381 |
| P9 clWT | A07 | 1471900 | sample | 0.9270059 | -1.7105794 |
| P9 clWT | A08 | 1606400 | sample | 1.0117143 | 0.2745192 |
| P9 clWT | A09 | 1565400 | sample | 0.9858924 | -0.3306038 |
| P9 clWT | A10 | 1673200 | sample | 1.0537851 | 1.2604269 |
| P9 clWT | A11 | 1543900 | sample | 0.9723517 | -0.6479244 |
| P9 clWT | A12 | 1836000 | empty | 1.1563169 | 3.6632080 |
| P9 clWT | B01 | 388580 | SAL | 0.2447286 | -17.6994050 |
| P9 clWT | B02 | 1663600 | sample | 1.0477390 | 1.1187396 |
| P9 clWT | B03 | 1649500 | sample | 1.0388588 | 0.9106363 |
| P9 clWT | B04 | 1597800 | sample | 1.0062980 | 0.1475910 |
| P9 clWT | B05 | 1609100 | sample | 1.0134148 | 0.3143688 |
| P9 clWT | B06 | 1541000 | sample | 0.9705253 | -0.6907258 |
| P9 clWT | B07 | 1597800 | sample | 1.0062980 | 0.1475910 |
| P9 clWT | B08 | 1509700 | sample | 0.9508124 | -1.1526855 |
| P9 clWT | B09 | 1666000 | sample | 1.0492505 | 1.1541614 |
| P9 clWT | B10 | 1699200 | sample | 1.0701600 | 1.6441635 |
| P9 clWT | B11 | 1417200 | sample | 0.8925557 | -2.5179020 |
| P9 clWT | B12 | 1737000 | empty | 1.0939665 | 2.2020574 |
| P9 clWT | C01 | 416470 | SAL | 0.2622937 | -17.2877738 |

*(continued)*

| Plate | Well | Value | Treatment | Norm | Score |
|---|---|---|---|---|---|
| P9 clWT | C02 | 1614300 | sample | 1.0166898 | 0.3911161 |
| P9 clWT | C03 | 1665700 | sample | 1.0490616 | 1.1497337 |
| P9 clWT | C04 | 1579900 | sample | 0.9950246 | -0.1165969 |
| P9 clWT | C05 | 1509200 | sample | 0.9504975 | -1.1600651 |
| P9 clWT | C06 | 1534100 | sample | 0.9661796 | -0.7925635 |
| P9 clWT | C07 | 1236300 | sample | 0.7786245 | -5.1878228 |
| P9 clWT | C08 | 1523600 | sample | 0.9595667 | -0.9475341 |
| P9 clWT | C09 | 1616600 | sample | 1.0181383 | 0.4250620 |
| P9 clWT | C10 | 1673700 | sample | 1.0541000 | 1.2678065 |
| P9 clWT | C11 | 1679800 | sample | 1.0579418 | 1.3578370 |
| P9 clWT | C12 | 1695900 | DMSO | 1.0680816 | 1.5954584 |
| P9 clWT | D01 | 1431200 | DMSO | 0.9013730 | -2.3112747 |
| P9 clWT | D02 | 1582800 | sample | 0.9968510 | -0.0737955 |
| P9 clWT | D03 | 1556400 | sample | 0.9802242 | -0.4634357 |
| P9 clWT | D04 | 1193500 | sample | 0.7516690 | -5.8195122 |
| P9 clWT | D05 | 1552800 | sample | 0.9779569 | -0.5165684 |
| P9 clWT | D06 | 1545500 | sample | 0.9733594 | -0.6243098 |
| P9 clWT | D07 | 1587200 | sample | 0.9996221 | -0.0088555 |
| P9 clWT | D08 | 1614800 | sample | 1.0170047 | 0.3984956 |
| P9 clWT | D09 | 1626300 | sample | 1.0242474 | 0.5682253 |
| P9 clWT | D10 | 1627500 | sample | 1.0250031 | 0.5859362 |
| P9 clWT | D11 | 1649800 | sample | 1.0390477 | 0.9150641 |
| P9 clWT | D12 | 1661400 | DMSO | 1.0463534 | 1.0862696 |
| P9 clWT | E01 | 1654300 | DMSO | 1.0418818 | 0.9814800 |
| P9 clWT | E02 | 1659900 | sample | 1.0454087 | 1.0641309 |
| P9 clWT | E03 | 1628700 | sample | 1.0257589 | 0.6036471 |
| P9 clWT | E04 | 1581900 | sample | 0.9962842 | -0.0870787 |
| P9 clWT | E05 | 1572100 | sample | 0.9901121 | -0.2317178 |
| P9 clWT | E06 | 1574000 | sample | 0.9913087 | -0.2036755 |
| P9 clWT | E07 | 1552100 | sample | 0.9775161 | -0.5268998 |
| P9 clWT | E08 | 1553500 | sample | 0.9783978 | -0.5062370 |
| P9 clWT | E09 | 1644700 | sample | 1.0358357 | 0.8397927 |
| P9 clWT | E10 | 1670600 | sample | 1.0521476 | 1.2220533 |
| P9 clWT | E11 | 1620100 | sample | 1.0203426 | 0.4767189 |
| P9 clWT | E12 | 1702100 | DMSO | 1.0719864 | 1.6869649 |
| P9 clWT | F01 | 1507900 | DMSO | 0.9496788 | -1.1792519 |
| P9 clWT | F02 | 1624400 | sample | 1.0230508 | 0.5401830 |
| P9 clWT | F03 | 1481000 | sample | 0.9327371 | -1.5762716 |
| P9 clWT | F04 | 1465000 | sample | 0.9226603 | -1.8124172 |
| P9 clWT | F05 | 1560300 | sample | 0.9826804 | -0.4058752 |
| P9 clWT | F06 | 1562100 | sample | 0.9838141 | -0.3793088 |
| P9 clWT | F07 | 1560400 | sample | 0.9827434 | -0.4043993 |
| P9 clWT | F08 | 1628400 | sample | 1.0255700 | 0.5992194 |
| P9 clWT | F09 | 1588400 | sample | 1.0003779 | 0.0088555 |
| P9 clWT | F10 | 1563500 | sample | 0.9846958 | -0.3586461 |
| P9 clWT | F11 | 1649700 | sample | 1.0389848 | 0.9135881 |
| P9 clWT | F12 | 513630 | SAL | 0.3234853 | -15.8537798 |
| P9 clWT | G01 | 1413300 | empty | 0.8900995 | -2.5754625 |
| P9 clWT | G02 | 1579400 | sample | 0.9947097 | -0.1239764 |
| P9 clWT | G03 | 1609500 | sample | 1.0136667 | 0.3202724 |
| P9 clWT | G04 | 1602000 | sample | 1.0089432 | 0.2095792 |
| P9 clWT | G05 | 1597400 | sample | 1.0060461 | 0.1416873 |
| P9 clWT | G06 | 1009600 | sample | 0.6358483 | -8.5337102 |
| P9 clWT | G07 | 1401600 | sample | 0.8827308 | -2.7481440 |
| P9 clWT | G08 | 1538000 | sample | 0.9686358 | -0.7350031 |
| P9 clWT | G09 | 941790 | sample | 0.5931415 | -9.5345246 |
| P9 clWT | G10 | 1648200 | sample | 1.0380401 | 0.8914495 |
| P9 clWT | G11 | 1652100 | sample | 1.0404963 | 0.9490100 |
| P9 clWT | G12 | 472160 | SAL | 0.2973674 | -16.4658396 |
| P9 clWT | H01 | 1431300 | empty | 0.9014359 | -2.3097988 |
| P9 clWT | H02 | 1654900 | sample | 1.0422597 | 0.9903354 |
| P9 clWT | H03 | 1645100 | sample | 1.0360877 | 0.8456963 |

*(continued)*

| Plate | Well | Value | Treatment | Norm | Score |
|---|---|---|---|---|---|
| P9 clWT | H04 | 1547500 | sample | 0.9746190 | -0.5947916 |
| P9 clWT | H05 | 1603700 | sample | 1.0100139 | 0.2346697 |
| P9 clWT | H06 | 1574400 | sample | 0.9915606 | -0.1977719 |
| P9 clWT | H07 | 1246600 | sample | 0.7851115 | -5.0358041 |
| P9 clWT | H08 | 1522300 | sample | 0.9587480 | -0.9667209 |
| P9 clWT | H09 | 1621900 | sample | 1.0214763 | 0.5032852 |
| P9 clWT | H10 | 1644000 | sample | 1.0353949 | 0.8294613 |
| P9 clWT | H11 | 1632400 | sample | 1.0280892 | 0.6582558 |
| P9 clWT | H12 | 448270 | SAL | 0.2823215 | -16.8184345 |
| P10 clWT | A01 | 381730 | SAL | 0.2346436 | -21.5063235 |
| P10 clWT | A02 | 1678800 | sample | 1.0319329 | 0.8973059 |
| P10 clWT | A03 | 1763400 | sample | 1.0839352 | 2.3585586 |
| P10 clWT | A04 | 1601600 | sample | 0.9844792 | -0.4361304 |
| P10 clWT | A05 | 1767100 | sample | 1.0862095 | 2.4224668 |
| P10 clWT | A06 | 678530 | sample | 0.4170821 | -16.3798483 |
| P10 clWT | A07 | 1658400 | sample | 1.0193933 | 0.5449471 |
| P10 clWT | A08 | 1566700 | sample | 0.9630267 | -1.0389403 |
| P10 clWT | A09 | 1629200 | sample | 1.0014445 | 0.0405904 |
| P10 clWT | A10 | 1706300 | sample | 1.0488367 | 1.3722994 |
| P10 clWT | A11 | 1691200 | sample | 1.0395550 | 1.1114848 |
| P10 clWT | A12 | 1800100 | empty | 1.1064941 | 2.9924590 |
| P10 clWT | B01 | 427330 | SAL | 0.2626733 | -20.7186980 |
| P10 clWT | B02 | 1692300 | sample | 1.0402311 | 1.1304845 |
| P10 clWT | B03 | 1695000 | sample | 1.0418908 | 1.1771202 |
| P10 clWT | B04 | 1752400 | sample | 1.0771737 | 2.1685612 |
| P10 clWT | B05 | 1674800 | sample | 1.0294741 | 0.8282159 |
| P10 clWT | B06 | 1653500 | sample | 1.0163814 | 0.4603119 |
| P10 clWT | B07 | 1632700 | sample | 1.0035959 | 0.1010441 |
| P10 clWT | B08 | 1606800 | sample | 0.9876756 | -0.3463134 |
| P10 clWT | B09 | 1716900 | sample | 1.0553524 | 1.5553878 |
| P10 clWT | B10 | 1736100 | sample | 1.0671543 | 1.8870196 |
| P10 clWT | B11 | 1679000 | sample | 1.0320558 | 0.9007604 |
| P10 clWT | B12 | 1752700 | empty | 1.0773581 | 2.1737429 |
| P10 clWT | C01 | 456830 | SAL | 0.2808065 | -20.2091595 |
| P10 clWT | C02 | 1632100 | sample | 1.0032271 | 0.0906806 |
| P10 clWT | C03 | 1612900 | sample | 0.9914251 | -0.2409512 |
| P10 clWT | C04 | 1639700 | sample | 1.0078987 | 0.2219515 |
| P10 clWT | C05 | 1591200 | sample | 0.9780865 | -0.6157643 |
| P10 clWT | C06 | 1668300 | sample | 1.0254787 | 0.7159447 |
| P10 clWT | C07 | 1598700 | sample | 0.9826966 | -0.4862206 |
| P10 clWT | C08 | 1641900 | sample | 1.0092510 | 0.2599510 |
| P10 clWT | C09 | 1300900 | sample | 0.7996435 | -5.6299683 |
| P10 clWT | C10 | 1664600 | sample | 1.0232044 | 0.6520365 |
| P10 clWT | C11 | 466030 | sample | 0.2864616 | -20.0502526 |
| P10 clWT | C12 | 1721000 | DMSO | 1.0578726 | 1.6262050 |
| P10 clWT | D01 | 1767100 | DMSO | 1.0862095 | 2.4224668 |
| P10 clWT | D02 | 1626000 | sample | 0.9994775 | -0.0146816 |
| P10 clWT | D03 | 1599600 | sample | 0.9832498 | -0.4706754 |
| P10 clWT | D04 | 1609500 | sample | 0.9893352 | -0.2996777 |
| P10 clWT | D05 | 1645300 | sample | 1.0113409 | 0.3186775 |
| P10 clWT | D06 | 1602800 | sample | 0.9852168 | -0.4154034 |
| P10 clWT | D07 | 1558700 | sample | 0.9581092 | -1.1771202 |
| P10 clWT | D08 | 1679500 | sample | 1.0323632 | 0.9093966 |
| P10 clWT | D09 | 1665200 | sample | 1.0235732 | 0.6624000 |
| P10 clWT | D10 | 1699500 | sample | 1.0446569 | 1.2548464 |
| P10 clWT | D11 | 1653800 | sample | 1.0165658 | 0.4654936 |
| P10 clWT | D12 | 1713300 | DMSO | 1.0531395 | 1.4932068 |
| P10 clWT | E01 | 1773800 | DMSO | 1.0903279 | 2.5381925 |
| P10 clWT | E02 | 1670500 | sample | 1.0268310 | 0.7539442 |
| P10 clWT | E03 | 1608000 | sample | 0.9884132 | -0.3255864 |
| P10 clWT | E04 | 1614800 | sample | 0.9925930 | -0.2081335 |

*(continued)*

| Plate | Well | Value | Treatment | Norm | Score |
|---|---|---|---|---|---|
| P10 clWT | E05 | 1612600 | sample | 0.9912407 | -0.2461330 |
| P10 clWT | E06 | 1618900 | sample | 0.9951133 | -0.1373163 |
| P10 clWT | E07 | 1575300 | sample | 0.9683130 | -0.8903969 |
| P10 clWT | E08 | 1539900 | sample | 0.9465532 | -1.5018431 |
| P10 clWT | E09 | 1655500 | sample | 1.0176107 | 0.4948569 |
| P10 clWT | E10 | 1723600 | sample | 1.0594708 | 1.6711135 |
| P10 clWT | E11 | 1628100 | sample | 1.0007684 | 0.0215906 |
| P10 clWT | E12 | 1758400 | DMSO | 1.0808618 | 2.2721961 |
| P10 clWT | F01 | 1625100 | DMSO | 0.9989243 | -0.0302269 |
| P10 clWT | F02 | 1760800 | sample | 1.0823370 | 2.3136501 |
| P10 clWT | F03 | 1467500 | sample | 0.9020500 | -2.7523714 |
| P10 clWT | F04 | 1610000 | sample | 0.9896426 | -0.2910415 |
| P10 clWT | F05 | 1606500 | sample | 0.9874912 | -0.3514952 |
| P10 clWT | F06 | 1674800 | sample | 1.0294741 | 0.8282159 |
| P10 clWT | F07 | 1533700 | sample | 0.9427421 | -1.6089325 |
| P10 clWT | F08 | 364520 | sample | 0.2240649 | -21.8035831 |
| P10 clWT | F09 | 1474100 | sample | 0.9061069 | -2.6383729 |
| P10 clWT | F10 | 1658300 | sample | 1.0193318 | 0.5432198 |
| P10 clWT | F11 | 1657300 | sample | 1.0187172 | 0.5259473 |
| P10 clWT | F12 | 559860 | SAL | 0.3441374 | -18.4295748 |
| P10 clWT | G01 | 1661500 | empty | 1.0212988 | 0.5984918 |
| P10 clWT | G02 | 1651500 | sample | 1.0151520 | 0.4257669 |
| P10 clWT | G03 | 451880 | sample | 0.2777638 | -20.2946583 |
| P10 clWT | G04 | 1610800 | sample | 0.9901343 | -0.2772235 |
| P10 clWT | G05 | 1556800 | sample | 0.9569413 | -1.2099380 |
| P10 clWT | G06 | 1551000 | sample | 0.9533762 | -1.3101184 |
| P10 clWT | G07 | 1598800 | sample | 0.9827581 | -0.4844934 |
| P10 clWT | G08 | 1425700 | sample | 0.8763561 | -3.4743615 |
| P10 clWT | G09 | 1626900 | sample | 1.0000307 | 0.0008636 |
| P10 clWT | G10 | 1448700 | sample | 0.8904939 | -3.0770942 |
| P10 clWT | G11 | 274770 | sample | 0.1688969 | -23.3537891 |
| P10 clWT | G12 | 475060 | SAL | 0.2920122 | -19.8942820 |
| P10 clWT | H01 | 1577300 | empty | 0.9695424 | -0.8558519 |
| P10 clWT | H02 | 1703400 | sample | 1.0470541 | 1.3222092 |
| P10 clWT | H03 | 1609800 | sample | 0.9895196 | -0.2944960 |
| P10 clWT | H04 | 1283300 | sample | 0.7888250 | -5.9339642 |
| P10 clWT | H05 | 1626800 | sample | 0.9999693 | -0.0008636 |
| P10 clWT | H06 | 1611700 | sample | 0.9906875 | -0.2616782 |
| P10 clWT | H07 | 1587100 | sample | 0.9755663 | -0.6865815 |
| P10 clWT | H08 | 1357300 | sample | 0.8343117 | -4.6557999 |
| P10 clWT | H09 | 1660400 | sample | 1.0206227 | 0.5794921 |
| P10 clWT | H10 | 1647700 | sample | 1.0128162 | 0.3601314 |
| P10 clWT | H11 | 1656500 | sample | 1.0182254 | 0.5121293 |
| P10 clWT | H12 | 418930 | SAL | 0.2575099 | -20.8637869 |
| P11 clWT | A01 | 312610 | SAL | 0.1932853 | -13.2635279 |
| P11 clWT | A02 | 1732700 | sample | 1.0713204 | 1.1726075 |
| P11 clWT | A03 | 1672800 | sample | 1.0342845 | 0.5636852 |
| P11 clWT | A04 | 1478200 | sample | 0.9139642 | -1.4145499 |
| P11 clWT | A05 | 1674900 | sample | 1.0355829 | 0.5850331 |
| P11 clWT | A06 | 1730600 | sample | 1.0700219 | 1.1512597 |
| P11 clWT | A07 | 1370000 | sample | 0.8470646 | -2.5144731 |
| P11 clWT | A08 | 1400700 | sample | 0.8660463 | -2.2023877 |
| P11 clWT | A09 | 1473500 | sample | 0.9110582 | -1.4623285 |
| P11 clWT | A10 | 1690700 | sample | 1.0453520 | 0.7456503 |
| P11 clWT | A11 | 1609000 | sample | 0.9948372 | -0.0848832 |
| P11 clWT | A12 | 1814700 | empty | 1.1220206 | 2.0061907 |
| P11 clWT | B01 | 425320 | SAL | 0.2629734 | -12.1177576 |
| P11 clWT | B02 | 75553 | sample | 0.0467141 | -15.6733659 |
| P11 clWT | B03 | 1518700 | sample | 0.9390052 | -1.0028412 |
| P11 clWT | B04 | 1671000 | sample | 1.0331715 | 0.5453870 |
| P11 clWT | B05 | 1685200 | sample | 1.0419513 | 0.6897392 |
| P11 clWT | B06 | 1683900 | sample | 1.0411476 | 0.6765239 |

*(continued)*

| Plate | Well | Value | Treatment | Norm | Score |
|---|---|---|---|---|---|
| P11 clWT | B07 | 1752000 | sample | 1.0832535 | 1.3688045 |
| P11 clWT | B08 | 1683500 | sample | 1.0409002 | 0.6724576 |
| P11 clWT | B09 | 1698700 | sample | 1.0502983 | 0.8269755 |
| P11 clWT | B10 | 1740300 | sample | 1.0760194 | 1.2498664 |
| P11 clWT | B11 | 1693800 | sample | 1.0472687 | 0.7771638 |
| P11 clWT | B12 | 1813800 | empty | 1.1214641 | 1.9970416 |
| P11 clWT | C01 | 444200 | SAL | 0.2746468 | -11.9258302 |
| P11 clWT | C02 | 1675400 | sample | 1.0358920 | 0.5901159 |
| P11 clWT | C03 | 1680100 | sample | 1.0387980 | 0.6378944 |
| P11 clWT | C04 | 1603200 | sample | 0.9912511 | -0.1438439 |
| P11 clWT | C05 | 1568800 | sample | 0.9699818 | -0.4935422 |
| P11 clWT | C06 | 1582600 | sample | 0.9785142 | -0.3532563 |
| P11 clWT | C07 | 1661000 | sample | 1.0269886 | 0.4437305 |
| P11 clWT | C08 | 1617400 | sample | 1.0000309 | 0.0005083 |
| P11 clWT | C09 | 1508900 | sample | 0.9329459 | -1.1024645 |
| P11 clWT | C10 | 1326300 | sample | 0.8200451 | -2.9587119 |
| P11 clWT | C11 | 1171400 | sample | 0.7242712 | -4.5333708 |
| P11 clWT | C12 | 1712100 | DMSO | 1.0585835 | 0.9631952 |
| P11 clWT | D01 | 1558200 | DMSO | 0.9634278 | -0.6012981 |
| P11 clWT | D02 | 1562700 | sample | 0.9662102 | -0.5555527 |
| P11 clWT | D03 | 1527400 | sample | 0.9443843 | -0.9144001 |
| P11 clWT | D04 | 1549700 | sample | 0.9581723 | -0.6877061 |
| P11 clWT | D05 | 1687100 | sample | 1.0431261 | 0.7090540 |
| P11 clWT | D06 | 1656200 | sample | 1.0240208 | 0.3949354 |
| P11 clWT | D07 | 1558300 | sample | 0.9634897 | -0.6002815 |
| P11 clWT | D08 | 1596600 | sample | 0.9871704 | -0.2109372 |
| P11 clWT | D09 | 1669400 | sample | 1.0321823 | 0.5291220 |
| P11 clWT | D10 | 1698800 | sample | 1.0503602 | 0.8279920 |
| P11 clWT | D11 | 1499300 | sample | 0.9270102 | -1.2000548 |
| P11 clWT | D12 | 1720100 | DMSO | 1.0635298 | 1.0445204 |
| P11 clWT | E01 | 1609100 | DMSO | 0.9948991 | -0.0838666 |
| P11 clWT | E02 | 1617300 | sample | 0.9999691 | -0.0005083 |
| P11 clWT | E03 | 1453200 | sample | 0.8985068 | -1.6686912 |
| P11 clWT | E04 | 1681800 | sample | 1.0398491 | 0.6551760 |
| P11 clWT | E05 | 1569600 | sample | 0.9704764 | -0.4854097 |
| P11 clWT | E06 | 1557500 | sample | 0.9629950 | -0.6084140 |
| P11 clWT | E07 | 1617200 | sample | 0.9999073 | -0.0015248 |
| P11 clWT | E08 | 1541400 | sample | 0.9530405 | -0.7720810 |
| P11 clWT | E09 | 1641000 | sample | 1.0146227 | 0.2404176 |
| P11 clWT | E10 | 1689600 | sample | 1.0446718 | 0.7344681 |
| P11 clWT | E11 | 1601100 | sample | 0.9899527 | -0.1651918 |
| P11 clWT | E12 | 1760200 | DMSO | 1.0883235 | 1.4521628 |
| P11 clWT | F01 | 1680200 | DMSO | 1.0388599 | 0.6389110 |
| P11 clWT | F02 | 1593700 | sample | 0.9853773 | -0.2404176 |
| P11 clWT | F03 | 1592600 | sample | 0.9846972 | -0.2515998 |
| P11 clWT | F04 | 1622100 | sample | 1.0029369 | 0.0482868 |
| P11 clWT | F05 | 1557800 | sample | 0.9631805 | -0.6053644 |
| P11 clWT | F06 | 1634300 | sample | 1.0104801 | 0.1723077 |
| P11 clWT | F07 | 1581600 | sample | 0.9778959 | -0.3634219 |
| P11 clWT | F08 | 1654800 | sample | 1.0231552 | 0.3807035 |
| P11 clWT | F09 | 1138400 | sample | 0.7038674 | -4.8688372 |
| P11 clWT | F10 | 391910 | sample | 0.2423161 | -12.4573920 |
| P11 clWT | F11 | 1646500 | sample | 1.0180233 | 0.2963286 |
| P11 clWT | F12 | 518910 | SAL | 0.3208396 | -11.1663546 |
| P11 clWT | G01 | 1622900 | empty | 1.0034315 | 0.0564193 |
| P11 clWT | G02 | 1704800 | sample | 1.0540699 | 0.8889859 |
| P11 clWT | G03 | 598150 | sample | 0.3698334 | -10.3608287 |
| P11 clWT | G04 | 1690600 | sample | 1.0452901 | 0.7446337 |
| P11 clWT | G05 | 1537700 | sample | 0.9507528 | -0.8096939 |
| P11 clWT | G06 | 558550 | sample | 0.3453489 | -10.7633883 |
| P11 clWT | G07 | 1563200 | sample | 0.9665193 | -0.5504699 |
| P11 clWT | G08 | 1578200 | sample | 0.9757937 | -0.3979851 |

*(continued)*

| Plate | Well | Value | Treatment | Norm | Score |
|---|---|---|---|---|---|
| P11 clWT | G09 | 1620400 | sample | 1.0018858 | 0.0310052 |
| P11 clWT | G10 | 1682200 | sample | 1.0400965 | 0.6592423 |
| P11 clWT | G11 | 1665900 | sample | 1.0300182 | 0.4935422 |
| P11 clWT | G12 | 471320 | SAL | 0.2914150 | -11.6501378 |
| P11 clWT | H01 | 1656400 | empty | 1.0241444 | 0.3969686 |
| P11 clWT | H02 | 1710100 | sample | 1.0573469 | 0.9428639 |
| P11 clWT | H03 | 1700300 | sample | 1.0512876 | 0.8432405 |
| P11 clWT | H04 | 1673000 | sample | 1.0344081 | 0.5657183 |
| P11 clWT | H05 | 1683200 | sample | 1.0407147 | 0.6694079 |
| P11 clWT | H06 | 1251600 | sample | 0.7738585 | -3.7180858 |
| P11 clWT | H07 | 1584400 | sample | 0.9796272 | -0.3349581 |
| P11 clWT | H08 | 1407000 | sample | 0.8699416 | -2.1383441 |
| P11 clWT | H09 | 1690000 | sample | 1.0449192 | 0.7385343 |
| P11 clWT | H10 | 1701400 | sample | 1.0519677 | 0.8544227 |
| P11 clWT | H11 | 1729200 | sample | 1.0691563 | 1.1370278 |
| P11 clWT | H12 | 435040 | SAL | 0.2689832 | -12.0189475 |
| P12 clWT | A01 | 317320 | SAL | 0.1966413 | -18.2928103 |
| P12 clWT | A02 | 1723200 | sample | 1.0678565 | 1.5451200 |
| P12 clWT | A03 | 1763300 | sample | 1.0927062 | 2.1109585 |
| P12 clWT | A04 | 1449600 | sample | 0.8983082 | -2.3155635 |
| P12 clWT | A05 | 1509900 | sample | 0.9356758 | -1.4646891 |
| P12 clWT | A06 | 1601600 | sample | 0.9925017 | -0.1707393 |
| P12 clWT | A07 | 1440200 | sample | 0.8924831 | -2.4482039 |
| P12 clWT | A08 | 1516300 | sample | 0.9396418 | -1.3743808 |
| P12 clWT | A09 | 1511300 | sample | 0.9365433 | -1.4449342 |
| P12 clWT | A10 | 1545300 | sample | 0.9576129 | -0.9651709 |
| P12 clWT | A11 | 1614000 | sample | 1.0001859 | 0.0042332 |
| P12 clWT | A12 | 1655700 | empty | 1.0260271 | 0.5926488 |
| P12 clWT | B01 | 395240 | SAL | 0.2449278 | -17.1933057 |
| P12 clWT | B02 | 1775500 | sample | 1.1002665 | 2.2831089 |
| P12 clWT | B03 | 835470 | sample | 0.5177356 | -10.9813587 |
| P12 clWT | B04 | 1661200 | sample | 1.0294355 | 0.6702576 |
| P12 clWT | B05 | 1721400 | sample | 1.0667410 | 1.5197208 |
| P12 clWT | B06 | 1738200 | sample | 1.0771519 | 1.7567803 |
| P12 clWT | B07 | 1533000 | sample | 0.9499907 | -1.1387323 |
| P12 clWT | B08 | 1613400 | sample | 0.9998141 | -0.0042332 |
| P12 clWT | B09 | 1644300 | sample | 1.0189626 | 0.4317870 |
| P12 clWT | B10 | 1615300 | sample | 1.0009915 | 0.0225771 |
| P12 clWT | B11 | 1622900 | sample | 1.0057012 | 0.1298183 |
| P12 clWT | B12 | 1687500 | empty | 1.0457334 | 1.0413686 |
| P12 clWT | C01 | 417880 | SAL | 0.2589577 | -16.8738397 |
| P12 clWT | C02 | 1618900 | sample | 1.0032224 | 0.0733756 |
| P12 clWT | C03 | 1631900 | sample | 1.0112784 | 0.2568145 |
| P12 clWT | C04 | 1627000 | sample | 1.0082419 | 0.1876721 |
| P12 clWT | C05 | 1651100 | sample | 1.0231766 | 0.5277396 |
| P12 clWT | C06 | 1582900 | sample | 0.9809134 | -0.4346091 |
| P12 clWT | C07 | 1557800 | sample | 0.9653591 | -0.7887873 |
| P12 clWT | C08 | 1614500 | sample | 1.0004958 | 0.0112885 |
| P12 clWT | C09 | 1592700 | sample | 0.9869864 | -0.2963244 |
| P12 clWT | C10 | 1715800 | sample | 1.0632707 | 1.4407010 |
| P12 clWT | C11 | 1716900 | sample | 1.0639524 | 1.4562227 |
| P12 clWT | C12 | 1691000 | DMSO | 1.0479023 | 1.0907560 |
| P12 clWT | D01 | 1555000 | DMSO | 0.9636240 | -0.8282972 |
| P12 clWT | D02 | 1590600 | sample | 0.9856851 | -0.3259568 |
| P12 clWT | D03 | 1293900 | sample | 0.8018219 | -4.5125972 |
| P12 clWT | D04 | 1646100 | sample | 1.0200781 | 0.4571862 |
| P12 clWT | D05 | 1567300 | sample | 0.9712462 | -0.6547358 |
| P12 clWT | D06 | 1633800 | sample | 1.0124558 | 0.2836248 |
| P12 clWT | D07 | 1649900 | sample | 1.0224329 | 0.5108068 |
| P12 clWT | D08 | 1664900 | sample | 1.0317283 | 0.7224671 |
| P12 clWT | D09 | 1677300 | sample | 1.0394125 | 0.8974396 |

*(continued)*

| Plate | Well | Value | Treatment | Norm | Score |
|---|---|---|---|---|---|
| P12 clWT | D10 | 1614700 | sample | 1.0006197 | 0.0141107 |
| P12 clWT | D11 | 1587800 | sample | 0.9839499 | -0.3654668 |
| P12 clWT | D12 | 1736400 | DMSO | 1.0760364 | 1.7313811 |
| P12 clWT | E01 | 1665600 | DMSO | 1.0321621 | 0.7323446 |
| P12 clWT | E02 | 1647600 | sample | 1.0210076 | 0.4783522 |
| P12 clWT | E03 | 1576000 | sample | 0.9766375 | -0.5319728 |
| P12 clWT | E04 | 1588100 | sample | 0.9841358 | -0.3612335 |
| P12 clWT | E05 | 721500 | sample | 0.4471091 | -12.5895535 |
| P12 clWT | E06 | 838330 | sample | 0.5195080 | -10.9410021 |
| P12 clWT | E07 | 1604300 | sample | 0.9941749 | -0.1326404 |
| P12 clWT | E08 | 1522300 | sample | 0.9433600 | -1.2897166 |
| P12 clWT | E09 | 1615400 | sample | 1.0010535 | 0.0239882 |
| P12 clWT | E10 | 1662500 | sample | 1.0302411 | 0.6886014 |
| P12 clWT | E11 | 1400400 | sample | 0.8678193 | -3.0098092 |
| P12 clWT | E12 | 1738300 | DMSO | 1.0772139 | 1.7581914 |
| P12 clWT | F01 | 1761500 | DMSO | 1.0915908 | 2.0855593 |
| P12 clWT | F02 | 1674600 | sample | 1.0377394 | 0.8593407 |
| P12 clWT | F03 | 1100000 | sample | 0.6816633 | -7.2486591 |
| P12 clWT | F04 | 1680300 | sample | 1.0412716 | 0.9397716 |
| P12 clWT | F05 | 1628600 | sample | 1.0092334 | 0.2102492 |
| P12 clWT | F06 | 1695800 | sample | 1.0508769 | 1.1584873 |
| P12 clWT | F07 | 1587700 | sample | 0.9838880 | -0.3668778 |
| P12 clWT | F08 | 1591600 | sample | 0.9863048 | -0.3118461 |
| P12 clWT | F09 | 1595700 | sample | 0.9888455 | -0.2539923 |
| P12 clWT | F10 | 1536700 | sample | 0.9522836 | -1.0865228 |
| P12 clWT | F11 | 1641300 | sample | 1.0171036 | 0.3894549 |
| P12 clWT | F12 | 553200 | SAL | 0.3428146 | -14.9643818 |
| P12 clWT | G01 | 1503200 | empty | 0.9315238 | -1.5592307 |
| P12 clWT | G02 | 1579400 | sample | 0.9787445 | -0.4839965 |
| P12 clWT | G03 | 1647000 | sample | 1.0206358 | 0.4698858 |
| P12 clWT | G04 | 1527900 | sample | 0.9468303 | -1.2106968 |
| P12 clWT | G05 | 1615900 | sample | 1.0013633 | 0.0310435 |
| P12 clWT | G06 | 1348700 | sample | 0.8357811 | -3.7393316 |
| P12 clWT | G07 | 1541900 | sample | 0.9555060 | -1.0131472 |
| P12 clWT | G08 | 1500900 | sample | 0.9300985 | -1.5916853 |
| P12 clWT | G09 | 1668300 | sample | 1.0338353 | 0.7704434 |
| P12 clWT | G10 | 1667100 | sample | 1.0330917 | 0.7535106 |
| P12 clWT | G11 | 1611700 | sample | 0.9987606 | -0.0282214 |
| P12 clWT | G12 | 510360 | SAL | 0.3162670 | -15.5688836 |
| P12 clWT | H01 | 1461100 | empty | 0.9054347 | -2.1532906 |
| P12 clWT | H02 | 1712600 | sample | 1.0612877 | 1.3955468 |
| P12 clWT | H03 | 1485300 | sample | 0.9204313 | -1.8118120 |
| P12 clWT | H04 | 1654000 | sample | 1.0249737 | 0.5686606 |
| P12 clWT | H05 | 1742100 | sample | 1.0795687 | 1.8118120 |
| P12 clWT | H06 | 1436200 | sample | 0.8900043 | -2.5046466 |
| P12 clWT | H07 | 1592500 | sample | 0.9868625 | -0.2991465 |
| P12 clWT | H08 | 1578700 | sample | 0.9783107 | -0.4938740 |
| P12 clWT | H09 | 1629800 | sample | 1.0099771 | 0.2271820 |
| P12 clWT | H10 | 1583500 | sample | 0.9812852 | -0.4261427 |
| P12 clWT | H11 | 1661800 | sample | 1.0298073 | 0.6787240 |
| P12 clWT | H12 | 479700 | SAL | 0.2972672 | -16.0015172 |
| P13 clWT | A01 | 351350 | SAL | 0.2177699 | -21.3076612 |
| P13 clWT | A02 | 1665600 | sample | 1.0323540 | 0.8813121 |
| P13 clWT | A03 | 1555800 | sample | 0.9642990 | -0.9724823 |
| P13 clWT | A04 | 1577300 | sample | 0.9776249 | -0.6094898 |
| P13 clWT | A05 | 1445100 | sample | 0.8956861 | -2.8414717 |
| P13 clWT | A06 | 327790 | sample | 0.2031672 | -21.7054334 |
| P13 clWT | A07 | 1701900 | sample | 1.0548531 | 1.4941785 |
| P13 clWT | A08 | 1646900 | sample | 1.0207636 | 0.5655930 |
| P13 clWT | A09 | 1648100 | sample | 1.0215074 | 0.5858531 |
| P13 clWT | A10 | 1530400 | sample | 0.9485558 | -1.4013200 |
| P13 clWT | A11 | 1311500 | sample | 0.8128796 | -5.0970904 |

*(continued)*

| Plate | Well | Value | Treatment | Norm | Score |
|---|---|---|---|---|---|
| P13 clWT | A12 | 1670300 | empty | 1.0352671 | 0.9606639 |
| P13 clWT | B01 | 451400 | SAL | 0.2797818 | -19.6184797 |
| P13 clWT | B02 | 1715300 | sample | 1.0631585 | 1.7204157 |
| P13 clWT | B03 | 1710000 | sample | 1.0598736 | 1.6309339 |
| P13 clWT | B04 | 1269900 | sample | 0.7870956 | -5.7994387 |
| P13 clWT | B05 | 1726400 | sample | 1.0700384 | 1.9078212 |
| P13 clWT | B06 | 1571800 | sample | 0.9742159 | -0.7023483 |
| P13 clWT | B07 | 1606500 | sample | 0.9957233 | -0.1164953 |
| P13 clWT | B08 | 1633600 | sample | 1.0125201 | 0.3410441 |
| P13 clWT | B09 | 1724400 | sample | 1.0687988 | 1.8740544 |
| P13 clWT | B10 | 1736300 | sample | 1.0761745 | 2.0749666 |
| P13 clWT | B11 | 1687200 | sample | 1.0457419 | 1.2459929 |
| P13 clWT | B12 | 1813600 | empty | 1.1240858 | 3.3800513 |
| P13 clWT | C01 | 456460 | SAL | 0.2829181 | -19.5330498 |
| P13 clWT | C02 | 1546200 | sample | 0.9583488 | -1.1345627 |
| P13 clWT | C03 | 1599900 | sample | 0.9916326 | -0.2279255 |
| P13 clWT | C04 | 1540300 | sample | 0.9546920 | -1.2341746 |
| P13 clWT | C05 | 1599100 | sample | 0.9911367 | -0.2414322 |
| P13 clWT | C06 | 1603300 | sample | 0.9937399 | -0.1705221 |
| P13 clWT | C07 | 1624900 | sample | 1.0071278 | 0.1941588 |
| P13 clWT | C08 | 1642900 | sample | 1.0182844 | 0.4980595 |
| P13 clWT | C09 | 1701200 | sample | 1.0544192 | 1.4823602 |
| P13 clWT | C10 | 1625900 | sample | 1.0077476 | 0.2110422 |
| P13 clWT | C11 | 1713900 | sample | 1.0622908 | 1.6967790 |
| P13 clWT | C12 | 1729200 | DMSO | 1.0717739 | 1.9550946 |
| P13 clWT | D01 | 1671800 | DMSO | 1.0361969 | 0.9859890 |
| P13 clWT | D02 | 1633500 | sample | 1.0124582 | 0.3393558 |
| P13 clWT | D03 | 1625800 | sample | 1.0076856 | 0.2093538 |
| P13 clWT | D04 | 1388400 | sample | 0.8605430 | -3.7987590 |
| P13 clWT | D05 | 1611000 | sample | 0.9985125 | -0.0405201 |
| P13 clWT | D06 | 1396100 | sample | 0.8653155 | -3.6687570 |
| P13 clWT | D07 | 1631700 | sample | 1.0113425 | 0.3089657 |
| P13 clWT | D08 | 1572900 | sample | 0.9748977 | -0.6837766 |
| P13 clWT | D09 | 436220 | sample | 0.2703731 | -19.8747693 |
| P13 clWT | D10 | 1291600 | sample | 0.8005454 | -5.4330695 |
| P13 clWT | D11 | 1629800 | sample | 1.0101649 | 0.2768873 |
| P13 clWT | D12 | 1685600 | DMSO | 1.0447502 | 1.2189795 |
| P13 clWT | E01 | 1647000 | DMSO | 1.0208256 | 0.5672813 |
| P13 clWT | E02 | 1680800 | sample | 1.0417751 | 1.1379394 |
| P13 clWT | E03 | 1644200 | sample | 1.0190901 | 0.5200079 |
| P13 clWT | E04 | 1613500 | sample | 1.0000620 | 0.0016883 |
| P13 clWT | E05 | 1524700 | sample | 0.9450229 | -1.4975552 |
| P13 clWT | E06 | 1639600 | sample | 1.0162390 | 0.4423444 |
| P13 clWT | E07 | 1613300 | sample | 0.9999380 | -0.0016883 |
| P13 clWT | E08 | 1563000 | sample | 0.9687616 | -0.8509220 |
| P13 clWT | E09 | 1616400 | sample | 1.0018594 | 0.0506501 |
| P13 clWT | E10 | 1645100 | sample | 1.0196479 | 0.5352029 |
| P13 clWT | E11 | 1683300 | sample | 1.0433247 | 1.1801478 |
| P13 clWT | E12 | 1773400 | DMSO | 1.0991695 | 2.7013397 |
| P13 clWT | F01 | 1669200 | DMSO | 1.0345853 | 0.9420922 |
| P13 clWT | F02 | 1639800 | sample | 1.0163630 | 0.4457211 |
| P13 clWT | F03 | 1626500 | sample | 1.0081195 | 0.2211722 |
| P13 clWT | F04 | 759170 | sample | 0.4705405 | -14.4222839 |
| P13 clWT | F05 | 1275300 | sample | 0.7904425 | -5.7082685 |
| P13 clWT | F06 | 1553800 | sample | 0.9630594 | -1.0062490 |
| P13 clWT | F07 | 1606500 | sample | 0.9957233 | -0.1164953 |
| P13 clWT | F08 | 1624800 | sample | 1.0070658 | 0.1924705 |
| P13 clWT | F09 | 1568300 | sample | 0.9720466 | -0.7614401 |
| P13 clWT | F10 | 1633500 | sample | 1.0124582 | 0.3393558 |
| P13 clWT | F11 | 1683500 | sample | 1.0434486 | 1.1835245 |
| P13 clWT | F12 | 537500 | SAL | 0.3331474 | -18.1648212 |
| P13 clWT | G01 | 1584000 | empty | 0.9817776 | -0.4963712 |

*(continued)*

| Plate | Well | Value | Treatment | Norm | Score |
|---|---|---|---|---|---|
| P13 clWT | G02 | 1508800 | sample | 0.9351680 | -1.7660008 |
| P13 clWT | G03 | 1661100 | sample | 1.0295649 | 0.8053369 |
| P13 clWT | G04 | 1577400 | sample | 0.9776869 | -0.6078014 |
| P13 clWT | G05 | 1589100 | sample | 0.9849386 | -0.4102660 |
| P13 clWT | G06 | 1583600 | sample | 0.9815297 | -0.5031245 |
| P13 clWT | G07 | 1618500 | sample | 1.0031610 | 0.0861052 |
| P13 clWT | G08 | 1574300 | sample | 0.9757655 | -0.6601399 |
| P13 clWT | G09 | 1597200 | sample | 0.9899591 | -0.2735106 |
| P13 clWT | G10 | 1632900 | sample | 1.0120863 | 0.3292258 |
| P13 clWT | G11 | 1630300 | sample | 1.0104748 | 0.2853290 |
| P13 clWT | G12 | 461410 | SAL | 0.2859861 | -19.4494771 |
| P13 clWT | H01 | 1681400 | empty | 1.0421470 | 1.1480694 |
| P13 clWT | H02 | 1581500 | sample | 0.9802281 | -0.5385796 |
| P13 clWT | H03 | 1730800 | sample | 1.0727656 | 1.9821080 |
| P13 clWT | H04 | 1672000 | sample | 1.0363208 | 0.9893657 |
| P13 clWT | H05 | 1612100 | sample | 0.9991942 | -0.0219484 |
| P13 clWT | H06 | 1574000 | sample | 0.9755795 | -0.6652049 |
| P13 clWT | H07 | 1532600 | sample | 0.9499194 | -1.3641766 |
| P13 clWT | H08 | 1618700 | sample | 1.0032850 | 0.0894819 |
| P13 clWT | H09 | 1574600 | sample | 0.9759514 | -0.6550749 |
| P13 clWT | H10 | 1667000 | sample | 1.0332218 | 0.9049488 |
| P13 clWT | H11 | 89563 | sample | 0.0555120 | -25.7275088 |
| P13 clWT | H12 | 392440 | SAL | 0.2432379 | -20.6139233 |
| P14 clWT | A01 | 305650 | SAL | 0.1940019 | -21.1481998 |
| P14 clWT | A02 | 1556700 | sample | 0.9880673 | -0.3130969 |
| P14 clWT | A03 | 1633400 | sample | 1.0367502 | 0.9642720 |
| P14 clWT | A04 | 1236300 | sample | 0.7847033 | -5.6490683 |
| P14 clWT | A05 | 1595000 | sample | 1.0123770 | 0.3247548 |
| P14 clWT | A06 | 1619900 | sample | 1.0281815 | 0.7394417 |
| P14 clWT | A07 | 1547200 | sample | 0.9820374 | -0.4713108 |
| P14 clWT | A08 | 1536500 | sample | 0.9752460 | -0.6495096 |
| P14 clWT | A09 | 998640 | sample | 0.6338559 | -9.6070800 |
| P14 clWT | A10 | 1637800 | sample | 1.0395430 | 1.0375500 |
| P14 clWT | A11 | 1517700 | sample | 0.9633132 | -0.9626066 |
| P14 clWT | A12 | 1565900 | empty | 0.9939067 | -0.1598793 |
| P14 clWT | B01 | 384320 | SAL | 0.2439353 | -19.8380223 |
| P14 clWT | B02 | 1663300 | sample | 1.0557283 | 1.4622294 |
| P14 clWT | B03 | 1651600 | sample | 1.0483021 | 1.2673765 |
| P14 clWT | B04 | 1549700 | sample | 0.9836242 | -0.4296756 |
| P14 clWT | B05 | 1666700 | sample | 1.0578864 | 1.5188533 |
| P14 clWT | B06 | 1489900 | sample | 0.9456680 | -1.4255903 |
| P14 clWT | B07 | 566580 | sample | 0.3596192 | -16.8026473 |
| P14 clWT | B08 | 1553000 | sample | 0.9857188 | -0.3747171 |
| P14 clWT | B09 | 1534900 | sample | 0.9742304 | -0.6761562 |
| P14 clWT | B10 | 1618900 | sample | 1.0275468 | 0.7227876 |
| P14 clWT | B11 | 1573700 | sample | 0.9988575 | -0.0299774 |
| P14 clWT | B12 | 1599400 | empty | 1.0151698 | 0.3980328 |
| P14 clWT | C01 | 376840 | SAL | 0.2391876 | -19.9625949 |
| P14 clWT | C02 | 1568800 | sample | 0.9957474 | -0.1115824 |
| P14 clWT | C03 | 1589300 | sample | 1.0087591 | 0.2298265 |
| P14 clWT | C04 | 1564200 | sample | 0.9928277 | -0.1881912 |
| P14 clWT | C05 | 1551300 | sample | 0.9846398 | -0.4030290 |
| P14 clWT | C06 | 1600100 | sample | 1.0156141 | 0.4096907 |
| P14 clWT | C07 | 1561100 | sample | 0.9908600 | -0.2398189 |
| P14 clWT | C08 | 1563900 | sample | 0.9926373 | -0.1931875 |
| P14 clWT | C09 | 585490 | sample | 0.3716217 | -16.4877184 |
| P14 clWT | C10 | 1575700 | sample | 1.0001269 | 0.0033308 |
| P14 clWT | C11 | 431330 | sample | 0.2737734 | -19.0551134 |
| P14 clWT | C12 | 1699500 | DMSO | 1.0787052 | 2.0651075 |
| P14 clWT | D01 | 1419400 | DMSO | 0.9009203 | -2.5997039 |
| P14 clWT | D02 | 1617500 | sample | 1.0266582 | 0.6994719 |

*(continued)*

| Plate | Well | Value | Treatment | Norm | Score |
|---|---|---|---|---|---|
| P14 clWT | D03 | 1522800 | sample | 0.9665503 | -0.8776707 |
| P14 clWT | D04 | 1646200 | sample | 1.0448746 | 1.1774444 |
| P14 clWT | D05 | 1556800 | sample | 0.9881308 | -0.3114315 |
| P14 clWT | D06 | 1546800 | sample | 0.9817836 | -0.4779725 |
| P14 clWT | D07 | 1627000 | sample | 1.0326880 | 0.8576858 |
| P14 clWT | D08 | 1493900 | sample | 0.9482069 | -1.3589740 |
| P14 clWT | D09 | 1606000 | sample | 1.0193589 | 0.5079498 |
| P14 clWT | D10 | 1561600 | sample | 0.9911774 | -0.2314919 |
| P14 clWT | D11 | 862540 | sample | 0.5474706 | -11.8737020 |
| P14 clWT | D12 | 1592900 | DMSO | 1.0110441 | 0.2897812 |
| P14 clWT | E01 | 1514500 | DMSO | 0.9612821 | -1.0158997 |
| P14 clWT | E02 | 1633400 | sample | 1.0367502 | 0.9642720 |
| P14 clWT | E03 | 1574400 | sample | 0.9993018 | -0.0183195 |
| P14 clWT | E04 | 1593300 | sample | 1.0112980 | 0.2964429 |
| P14 clWT | E05 | 1593600 | sample | 1.0114884 | 0.3014391 |
| P14 clWT | E06 | 1635600 | sample | 1.0381466 | 1.0009110 |
| P14 clWT | E07 | 1586800 | sample | 1.0071723 | 0.1881912 |
| P14 clWT | E08 | 1570500 | sample | 0.9968264 | -0.0832705 |
| P14 clWT | E09 | 1608800 | sample | 1.0211361 | 0.5545813 |
| P14 clWT | E10 | 1571800 | sample | 0.9976515 | -0.0616201 |
| P14 clWT | E11 | 1638800 | sample | 1.0401777 | 1.0542041 |
| P14 clWT | E12 | 1735600 | DMSO | 1.1016185 | 2.6663203 |
| P14 clWT | F01 | 1433600 | DMSO | 0.9099334 | -2.3632158 |
| P14 clWT | F02 | 1639800 | sample | 1.0408124 | 1.0708582 |
| P14 clWT | F03 | 1629300 | sample | 1.0341479 | 0.8959902 |
| P14 clWT | F04 | 1602400 | sample | 1.0170739 | 0.4479951 |
| P14 clWT | F05 | 1624000 | sample | 1.0307839 | 0.8077235 |
| P14 clWT | F06 | 1578900 | sample | 1.0021580 | 0.0566239 |
| P14 clWT | F07 | 1549000 | sample | 0.9831799 | -0.4413335 |
| P14 clWT | F08 | 1617400 | sample | 1.0265947 | 0.6978065 |
| P14 clWT | F09 | 1609100 | sample | 1.0213266 | 0.5595775 |
| P14 clWT | F10 | 1529300 | sample | 0.9706760 | -0.7694191 |
| P14 clWT | F11 | 1688400 | sample | 1.0716598 | 1.8802471 |
| P14 clWT | F12 | 552760 | SAL | 0.3508474 | -17.0328069 |
| P14 clWT | G01 | 1512300 | empty | 0.9598858 | -1.0525387 |
| P14 clWT | G02 | 1652000 | sample | 1.0485560 | 1.2740381 |
| P14 clWT | G03 | 1610700 | sample | 1.0223421 | 0.5862241 |
| P14 clWT | G04 | 1594800 | sample | 1.0122501 | 0.3214240 |
| P14 clWT | G05 | 1563300 | sample | 0.9922564 | -0.2031799 |
| P14 clWT | G06 | 1484900 | sample | 0.9424944 | -1.5088608 |
| P14 clWT | G07 | 1559200 | sample | 0.9896541 | -0.2714617 |
| P14 clWT | G08 | 1595200 | sample | 1.0125040 | 0.3280856 |
| P14 clWT | G09 | 1608700 | sample | 1.0210727 | 0.5529159 |
| P14 clWT | G10 | 1560900 | sample | 0.9907331 | -0.2431498 |
| P14 clWT | G11 | 630840 | sample | 0.4004062 | -15.7324553 |
| P14 clWT | G12 | 517240 | SAL | 0.3283021 | -17.6243603 |
| P14 clWT | H01 | 1240400 | empty | 0.7873056 | -5.5807865 |
| P14 clWT | H02 | 1625000 | sample | 1.0314186 | 0.8243776 |
| P14 clWT | H03 | 1495300 | sample | 0.9490955 | -1.3356582 |
| P14 clWT | H04 | 1535100 | sample | 0.9743573 | -0.6728254 |
| P14 clWT | H05 | 1620600 | sample | 1.0286258 | 0.7510996 |
| P14 clWT | H06 | 1630400 | sample | 1.0348461 | 0.9143097 |
| P14 clWT | H07 | 1560700 | sample | 0.9906062 | -0.2464806 |
| P14 clWT | H08 | 1530900 | sample | 0.9716915 | -0.7427725 |
| P14 clWT | H09 | 1575300 | sample | 0.9998731 | -0.0033308 |
| P14 clWT | H10 | 1622800 | sample | 1.0300222 | 0.7877386 |
| P14 clWT | H11 | 1668000 | sample | 1.0587115 | 1.5405036 |
| P14 clWT | H12 | 489180 | SAL | 0.3104919 | -18.0916741 |
| P15 clWT | A01 | 363050 | SAL | 0.2233535 | -17.1323367 |
| P15 clWT | A02 | 1677000 | sample | 1.0317143 | 0.6995976 |
| P15 clWT | A03 | 1630200 | sample | 1.0029223 | 0.0644634 |
| P15 clWT | A04 | 1563400 | sample | 0.9618260 | -0.8420956 |

*(continued)*

| Plate | Well | Value | Treatment | Norm | Score |
|---|---|---|---|---|---|
| P15 clWT | A05 | 1685800 | sample | 1.0371282 | 0.8190245 |
| P15 clWT | A06 | 1705800 | sample | 1.0494325 | 1.0904493 |
| P15 clWT | A07 | 1586100 | sample | 0.9757913 | -0.5340284 |
| P15 clWT | A08 | 1558500 | sample | 0.9588114 | -0.9085947 |
| P15 clWT | A09 | 1611500 | sample | 0.9914178 | -0.1893188 |
| P15 clWT | A10 | 1681300 | sample | 1.0343597 | 0.7579539 |
| P15 clWT | A11 | 1526700 | sample | 0.9392476 | -1.3401602 |
| P15 clWT | A12 | 1761500 | empty | 1.0836999 | 1.8463676 |
| P15 clWT | B01 | 447160 | SAL | 0.2750992 | -15.9908595 |
| P15 clWT | B02 | 1686000 | sample | 1.0372512 | 0.8217387 |
| P15 clWT | B03 | 1584900 | sample | 0.9750531 | -0.5503139 |
| P15 clWT | B04 | 1645800 | sample | 1.0125196 | 0.2761748 |
| P15 clWT | B05 | 1665600 | sample | 1.0247009 | 0.5448854 |
| P15 clWT | B06 | 1687300 | sample | 1.0380510 | 0.8393814 |
| P15 clWT | B07 | 1652300 | sample | 1.0165185 | 0.3643879 |
| P15 clWT | B08 | 1451700 | sample | 0.8931065 | -2.3580034 |
| P15 clWT | B09 | 1681700 | sample | 1.0346058 | 0.7633824 |
| P15 clWT | B10 | 1697700 | sample | 1.0444492 | 0.9805223 |
| P15 clWT | B11 | 1563600 | sample | 0.9619490 | -0.8393814 |
| P15 clWT | B12 | 1866800 | empty | 1.1484820 | 3.2754194 |
| P15 clWT | C01 | 495000 | SAL | 0.3045311 | -15.3416112 |
| P15 clWT | C02 | 1581700 | sample | 0.9730844 | -0.5937419 |
| P15 clWT | C03 | 1669800 | sample | 1.0272848 | 0.6018846 |
| P15 clWT | C04 | 1587000 | sample | 0.9763450 | -0.5218143 |
| P15 clWT | C05 | 1595800 | sample | 0.9817589 | -0.4023873 |
| P15 clWT | C06 | 1650400 | sample | 1.0153496 | 0.3386025 |
| P15 clWT | C07 | 1665400 | sample | 1.0245778 | 0.5421711 |
| P15 clWT | C08 | 1662000 | sample | 1.0224861 | 0.4960289 |
| P15 clWT | C09 | 295210 | sample | 0.1816174 | -18.0530098 |
| P15 clWT | C10 | 1559700 | sample | 0.9595497 | -0.8923092 |
| P15 clWT | C11 | 1636400 | sample | 1.0067366 | 0.1486051 |
| P15 clWT | C12 | 1771200 | DMSO | 1.0896675 | 1.9780086 |
| P15 clWT | D01 | 1625200 | DMSO | 0.9998462 | -0.0033928 |
| P15 clWT | D02 | 528280 | sample | 0.3250054 | -14.8899603 |
| P15 clWT | D03 | 1626600 | sample | 1.0007075 | 0.0156069 |
| P15 clWT | D04 | 1633800 | sample | 1.0051370 | 0.1133199 |
| P15 clWT | D05 | 1545400 | sample | 0.9507521 | -1.0863780 |
| P15 clWT | D06 | 1603600 | sample | 0.9865576 | -0.2965317 |
| P15 clWT | D07 | 1694500 | sample | 1.0424805 | 0.9370943 |
| P15 clWT | D08 | 1332600 | sample | 0.8198345 | -3.9743384 |
| P15 clWT | D09 | 1659500 | sample | 1.0209480 | 0.4621008 |
| P15 clWT | D10 | 1624300 | sample | 0.9992925 | -0.0156069 |
| P15 clWT | D11 | 1659200 | sample | 1.0207635 | 0.4580294 |
| P15 clWT | D12 | 1701400 | DMSO | 1.0467255 | 1.0307359 |
| P15 clWT | E01 | 1700300 | DMSO | 1.0460488 | 1.0158075 |
| P15 clWT | E02 | 1713900 | sample | 1.0544157 | 1.2003764 |
| P15 clWT | E03 | 1461000 | sample | 0.8988280 | -2.2317909 |
| P15 clWT | E04 | 1720200 | sample | 1.0582916 | 1.2858752 |
| P15 clWT | E05 | 1698400 | sample | 1.0448799 | 0.9900222 |
| P15 clWT | E06 | 1666600 | sample | 1.0253161 | 0.5584566 |
| P15 clWT | E07 | 1643100 | sample | 1.0108585 | 0.2395324 |
| P15 clWT | E08 | 1413300 | sample | 0.8694823 | -2.8791391 |
| P15 clWT | E09 | 1664400 | sample | 1.0239626 | 0.5285999 |
| P15 clWT | E10 | 1689000 | sample | 1.0390969 | 0.8624525 |
| P15 clWT | E11 | 1616200 | sample | 0.9943093 | -0.1255340 |
| P15 clWT | E12 | 1730300 | DMSO | 1.0645052 | 1.4229448 |
| P15 clWT | F01 | 1753900 | DMSO | 1.0790243 | 1.7432261 |
| P15 clWT | F02 | 873690 | sample | 0.5375065 | -10.2023174 |
| P15 clWT | F03 | 1516200 | sample | 0.9327878 | -1.4826583 |
| P15 clWT | F04 | 1336900 | sample | 0.8224799 | -3.9159821 |
| P15 clWT | F05 | 1653000 | sample | 1.0169492 | 0.3738877 |
| P15 clWT | F06 | 1645700 | sample | 1.0124581 | 0.2748177 |

*(continued)*

| Plate | Well | Value | Treatment | Norm | Score |
|---|---|---|---|---|---|
| P15 clWT | F07 | 828720 | sample | 0.5098404 | -10.8126162 |
| P15 clWT | F08 | 1647400 | sample | 1.0135040 | 0.2978888 |
| P15 clWT | F09 | 1657700 | sample | 1.0198407 | 0.4376726 |
| P15 clWT | F10 | 1617000 | sample | 0.9948014 | -0.1146770 |
| P15 clWT | F11 | 1597900 | sample | 0.9830508 | -0.3738877 |
| P15 clWT | F12 | 532440 | SAL | 0.3275647 | -14.8335039 |
| P15 clWT | G01 | 1619900 | empty | 0.9965856 | -0.0753204 |
| P15 clWT | G02 | 1737800 | sample | 1.0691193 | 1.5247291 |
| P15 clWT | G03 | 1633100 | sample | 1.0047064 | 0.1038200 |
| P15 clWT | G04 | 274850 | sample | 0.1690916 | -18.3293203 |
| P15 clWT | G05 | 1598800 | sample | 0.9836045 | -0.3616736 |
| P15 clWT | G06 | 1577600 | sample | 0.9705620 | -0.6493840 |
| P15 clWT | G07 | 1560900 | sample | 0.9602879 | -0.8760237 |
| P15 clWT | G08 | 867920 | sample | 0.5339568 | -10.2806234 |
| P15 clWT | G09 | 1512900 | sample | 0.9307576 | -1.5274434 |
| P15 clWT | G10 | 1669900 | sample | 1.0273463 | 0.6032417 |
| P15 clWT | G11 | 1655600 | sample | 1.0185487 | 0.4091730 |
| P15 clWT | G12 | 474750 | SAL | 0.2920730 | -15.6164289 |
| P15 clWT | H01 | 1562100 | empty | 0.9610262 | -0.8597382 |
| P15 clWT | H02 | 980150 | sample | 0.6030022 | -8.7575229 |
| P15 clWT | H03 | 1617400 | sample | 0.9950475 | -0.1092485 |
| P15 clWT | H04 | 1488100 | sample | 0.9155003 | -1.8640102 |
| P15 clWT | H05 | 1648800 | sample | 1.0143653 | 0.3168885 |
| P15 clWT | H06 | 1554400 | sample | 0.9562890 | -0.9642368 |
| P15 clWT | H07 | 1551800 | sample | 0.9546895 | -0.9995220 |
| P15 clWT | H08 | 1610600 | sample | 0.9908641 | -0.2015330 |
| P15 clWT | H09 | 1658000 | sample | 1.0200252 | 0.4417439 |
| P15 clWT | H10 | 1709400 | sample | 1.0516472 | 1.1393058 |
| P15 clWT | H11 | 1111000 | sample | 0.6835030 | -6.9817258 |
| P15 clWT | H12 | 425090 | SAL | 0.2615214 | -16.2903768 |
| P16 clWT | A01 | 363610 | SAL | 0.2224254 | -15.0284344 |
| P16 clWT | A02 | 1456300 | sample | 0.8908396 | -2.1097787 |
| P16 clWT | A03 | 1570100 | sample | 0.9604527 | -0.7643440 |
| P16 clWT | A04 | 1221800 | sample | 0.7473926 | -4.8822254 |
| P16 clWT | A05 | 1722800 | sample | 1.0538614 | 1.0409976 |
| P16 clWT | A06 | 1804000 | sample | 1.1035327 | 2.0010090 |
| P16 clWT | A07 | 1618900 | sample | 0.9903043 | -0.1873914 |
| P16 clWT | A08 | 1387500 | sample | 0.8487536 | -2.9231874 |
| P16 clWT | A09 | 1375500 | sample | 0.8414131 | -3.0650610 |
| P16 clWT | A10 | 1443500 | sample | 0.8830096 | -2.2611106 |
| P16 clWT | A11 | 1669100 | sample | 1.0210124 | 0.4061132 |
| P16 clWT | A12 | 1803600 | empty | 1.1032880 | 1.9962798 |
| P16 clWT | B01 | 446760 | SAL | 0.2732895 | -14.0453686 |
| P16 clWT | B02 | 1727500 | sample | 1.0567365 | 1.0965647 |
| P16 clWT | B03 | 1603100 | sample | 0.9806392 | -0.3741916 |
| P16 clWT | B04 | 1712700 | sample | 1.0476831 | 0.9215873 |
| P16 clWT | B05 | 106500 | sample | 0.0651476 | -18.0681946 |
| P16 clWT | B06 | 1743500 | sample | 1.0665239 | 1.2857295 |
| P16 clWT | B07 | 1672900 | sample | 1.0233369 | 0.4510398 |
| P16 clWT | B08 | 1682100 | sample | 1.0289647 | 0.5598096 |
| P16 clWT | B09 | 1555800 | sample | 0.9517052 | -0.9334101 |
| P16 clWT | B10 | 1671700 | sample | 1.0226028 | 0.4368525 |
| P16 clWT | B11 | 932470 | sample | 0.5704053 | -8.3029162 |
| P16 clWT | B12 | 1808500 | empty | 1.1062854 | 2.0542116 |
| P16 clWT | C01 | 467720 | SAL | 0.2861110 | -13.7975627 |
| P16 clWT | C02 | 1509300 | sample | 0.9232604 | -1.4831703 |
| P16 clWT | C03 | 1545000 | sample | 0.9450986 | -1.0610963 |
| P16 clWT | C04 | 1549300 | sample | 0.9477290 | -1.0102583 |
| P16 clWT | C05 | 1395100 | sample | 0.8534027 | -2.8333341 |
| P16 clWT | C06 | 1673300 | sample | 1.0235816 | 0.4557690 |
| P16 clWT | C07 | 1638000 | sample | 1.0019881 | 0.0384241 |

*(continued)*

| Plate | Well | Value | Treatment | Norm | Score |
|---|---|---|---|---|---|
| P16 clWT | C08 | 1608200 | sample | 0.9837590 | -0.3138953 |
| P16 clWT | C09 | 1632200 | sample | 0.9984401 | -0.0301481 |
| P16 clWT | C10 | 1720200 | sample | 1.0522710 | 1.0102583 |
| P16 clWT | C11 | 1719900 | sample | 1.0520875 | 1.0067114 |
| P16 clWT | C12 | 1839000 | DMSO | 1.1249427 | 2.4148070 |
| P16 clWT | D01 | 1830400 | DMSO | 1.1196819 | 2.3131309 |
| P16 clWT | D02 | 369440 | sample | 0.2259917 | -14.9595075 |
| P16 clWT | D03 | 1609600 | sample | 0.9846154 | -0.2973434 |
| P16 clWT | D04 | 1512300 | sample | 0.9250956 | -1.4477019 |
| P16 clWT | D05 | 1308700 | sample | 0.8005505 | -3.8548241 |
| P16 clWT | D06 | 1633800 | sample | 0.9994189 | -0.0112317 |
| P16 clWT | D07 | 1638700 | sample | 1.0024163 | 0.0467001 |
| P16 clWT | D08 | 1670100 | sample | 1.0216241 | 0.4179360 |
| P16 clWT | D09 | 1692300 | sample | 1.0352042 | 0.6804022 |
| P16 clWT | D10 | 1683300 | sample | 1.0296987 | 0.5739970 |
| P16 clWT | D11 | 1703500 | sample | 1.0420554 | 0.8128175 |
| P16 clWT | D12 | 1603500 | DMSO | 0.9808839 | -0.3694625 |
| P16 clWT | E01 | 1731600 | DMSO | 1.0592445 | 1.1450382 |
| P16 clWT | E02 | 1687000 | sample | 1.0319621 | 0.6177413 |
| P16 clWT | E03 | 1743300 | sample | 1.0664016 | 1.2833650 |
| P16 clWT | E04 | 1702700 | sample | 1.0415660 | 0.8033593 |
| P16 clWT | E05 | 1627900 | sample | 0.9958098 | -0.0809862 |
| P16 clWT | E06 | 1626700 | sample | 0.9950757 | -0.0951735 |
| P16 clWT | E07 | 1675900 | sample | 1.0251720 | 0.4865082 |
| P16 clWT | E08 | 1541600 | sample | 0.9430188 | -1.1012939 |
| P16 clWT | E09 | 1744700 | sample | 1.0672580 | 1.2999169 |
| P16 clWT | E10 | 1707500 | sample | 1.0445022 | 0.8601087 |
| P16 clWT | E11 | 1701200 | sample | 1.0406484 | 0.7856251 |
| P16 clWT | E12 | 1799700 | DMSO | 1.1009023 | 1.9501709 |
| P16 clWT | F01 | 1674500 | DMSO | 1.0243156 | 0.4699563 |
| P16 clWT | F02 | 1689700 | sample | 1.0336137 | 0.6496629 |
| P16 clWT | F03 | 1581400 | sample | 0.9673650 | -0.6307464 |
| P16 clWT | F04 | 1635700 | sample | 1.0005811 | 0.0112317 |
| P16 clWT | F05 | 1662300 | sample | 1.0168527 | 0.3257181 |
| P16 clWT | F06 | 1736700 | sample | 1.0623643 | 1.2053345 |
| P16 clWT | F07 | 1688600 | sample | 1.0329408 | 0.6366578 |
| P16 clWT | F08 | 1583000 | sample | 0.9683438 | -0.6118299 |
| P16 clWT | F09 | 1669400 | sample | 1.0211959 | 0.4096600 |
| P16 clWT | F10 | 1674300 | sample | 1.0241933 | 0.4675918 |
| P16 clWT | F11 | 1691500 | sample | 1.0347148 | 0.6709439 |
| P16 clWT | F12 | 488540 | SAL | 0.2988469 | -13.5514120 |
| P16 clWT | G01 | 1558700 | empty | 0.9534791 | -0.8991240 |
| P16 clWT | G02 | 242980 | sample | 0.1486343 | -16.4546188 |
| P16 clWT | G03 | 1660100 | sample | 1.0155070 | 0.2997080 |
| P16 clWT | G04 | 1479800 | sample | 0.9052149 | -1.8319429 |
| P16 clWT | G05 | 881720 | sample | 0.5393608 | -8.9029233 |
| P16 clWT | G06 | 1586300 | sample | 0.9703624 | -0.5728147 |
| P16 clWT | G07 | 1616100 | sample | 0.9885915 | -0.2204952 |
| P16 clWT | G08 | 1591400 | sample | 0.9734822 | -0.5125184 |
| P16 clWT | G09 | 1507700 | sample | 0.9222817 | -1.5020868 |
| P16 clWT | G10 | 1684700 | sample | 1.0305551 | 0.5905489 |
| P16 clWT | G11 | 1684000 | sample | 1.0301269 | 0.5822729 |
| P16 clWT | G12 | 479410 | SAL | 0.2932620 | -13.6593541 |
| P16 clWT | H01 | 1418300 | empty | 0.8675944 | -2.5590451 |
| P16 clWT | H02 | 1661300 | sample | 1.0162410 | 0.3138953 |
| P16 clWT | H03 | 437930 | sample | 0.2678881 | -14.1497639 |
| P16 clWT | H04 | 327070 | sample | 0.2000734 | -15.4604396 |
| P16 clWT | H05 | 1582800 | sample | 0.9682214 | -0.6141945 |
| P16 clWT | H06 | 1627300 | sample | 0.9954427 | -0.0880799 |
| P16 clWT | H07 | 1678100 | sample | 1.0265178 | 0.5125184 |
| P16 clWT | H08 | 1620600 | sample | 0.9913442 | -0.1672926 |
| P16 clWT | H09 | 1630700 | sample | 0.9975226 | -0.0478823 |

*(continued)*

| Plate | Well | Value | Treatment | Norm | Score |
|---|---|---|---|---|---|
| P16 clWT | H10 | 1692100 | sample | 1.0350818 | 0.6780376 |
| P16 clWT | H11 | 1699800 | sample | 1.0397920 | 0.7690732 |
| P16 clWT | H12 | 439480 | SAL | 0.2688362 | -14.1314386 |

#### clone 101

| Plate | Well | Value | Treatment | Norm | Score |
|---|---|---|---|---|---|
| P1 cl101 | A01 | 368330 | SAL | 0.2348295 | -12.5992774 |
| P1 cl101 | A02 | 1467900 | sample | 0.9358623 | -1.0560898 |
| P1 cl101 | A03 | 1504300 | sample | 0.9590692 | -0.6739659 |
| P1 cl101 | A04 | 1410100 | sample | 0.8990118 | -1.6628690 |
| P1 cl101 | A05 | 1482600 | sample | 0.9452343 | -0.9017705 |
| P1 cl101 | A06 | 1504200 | sample | 0.9590054 | -0.6750157 |
| P1 cl101 | A07 | 1630400 | sample | 1.0394645 | 0.6498207 |
| P1 cl101 | A08 | 1477800 | sample | 0.9421741 | -0.9521605 |
| P1 cl101 | A09 | 1543800 | sample | 0.9842525 | -0.2592984 |
| P1 cl101 | A10 | 1557200 | sample | 0.9927957 | -0.1186264 |
| P1 cl101 | A11 | 1557400 | sample | 0.9929232 | -0.1165268 |
| P1 cl101 | A12 | 1579200 | empty | 1.0068218 | 0.1123276 |
| P1 cl101 | B01 | 374600 | SAL | 0.2388269 | -12.5334555 |
| P1 cl101 | B02 | 1520700 | sample | 0.9695250 | -0.5018001 |
| P1 cl101 | B03 | 1549400 | sample | 0.9878228 | -0.2005101 |
| P1 cl101 | B04 | 1580200 | sample | 1.0074594 | 0.1228256 |
| P1 cl101 | B05 | 1666800 | sample | 1.0626713 | 1.0319446 |
| P1 cl101 | B06 | 1702100 | sample | 1.0851769 | 1.4025209 |
| P1 cl101 | B07 | 1600800 | sample | 1.0205929 | 0.3390825 |
| P1 cl101 | B08 | 1605500 | sample | 1.0235894 | 0.3884227 |
| P1 cl101 | B09 | 1357400 | sample | 0.8654128 | -2.2161089 |
| P1 cl101 | B10 | 1709800 | sample | 1.0900861 | 1.4833548 |
| P1 cl101 | B11 | 1760400 | sample | 1.1223462 | 2.0145491 |
| P1 cl101 | B12 | 1661500 | empty | 1.0592923 | 0.9763057 |
| P1 cl101 | C01 | 402690 | SAL | 0.2567357 | -12.2385692 |
| P1 cl101 | C02 | 1481300 | sample | 0.9444055 | -0.9154178 |
| P1 cl101 | C03 | 1556200 | sample | 0.9921581 | -0.1291243 |
| P1 cl101 | C04 | 1545200 | sample | 0.9851450 | -0.2446013 |
| P1 cl101 | C05 | 1649900 | sample | 1.0518967 | 0.8545299 |
| P1 cl101 | C06 | 1607300 | sample | 1.0247370 | 0.4073189 |
| P1 cl101 | C07 | 1648500 | sample | 1.0510041 | 0.8398329 |
| P1 cl101 | C08 | 1609800 | sample | 1.0263309 | 0.4335637 |
| P1 cl101 | C09 | 1605500 | sample | 1.0235894 | 0.3884227 |
| P1 cl101 | C10 | 1661600 | sample | 1.0593561 | 0.9773555 |
| P1 cl101 | C11 | 1669300 | sample | 1.0642652 | 1.0581894 |
| P1 cl101 | C12 | 1575400 | DMSO | 1.0043991 | 0.0724356 |
| P1 cl101 | D01 | 1451700 | DMSO | 0.9255339 | -1.2261560 |
| P1 cl101 | D02 | 1558400 | sample | 0.9935607 | -0.1060289 |
| P1 cl101 | D03 | 1598300 | sample | 1.0189990 | 0.3128377 |
| P1 cl101 | D04 | 1558600 | sample | 0.9936882 | -0.1039293 |
| P1 cl101 | D05 | 632360 | sample | 0.4031623 | -9.8275141 |
| P1 cl101 | D06 | 1584200 | sample | 1.0100096 | 0.1648172 |
| P1 cl101 | D07 | 1663400 | sample | 1.0605037 | 0.9962517 |
| P1 cl101 | D08 | 309720 | sample | 0.1974625 | -13.2145600 |
| P1 cl101 | D09 | 1653800 | sample | 1.0543832 | 0.8954718 |
| P1 cl101 | D10 | 1745100 | sample | 1.1125916 | 1.8539310 |
| P1 cl101 | D11 | 267020 | sample | 0.1702391 | -13.6628208 |
| P1 cl101 | D12 | 1561500 | DMSO | 0.9955371 | -0.0734854 |
| P1 cl101 | E01 | 1405700 | DMSO | 0.8962066 | -1.7090599 |
| P1 cl101 | E02 | 1574800 | sample | 1.0040166 | 0.0661368 |
| P1 cl101 | E03 | 1643400 | sample | 1.0477526 | 0.7862935 |
| P1 cl101 | E04 | 1564300 | sample | 0.9973223 | -0.0440912 |

*(continued)*

| Plate | Well | Value | Treatment | Norm | Score |
|---|---|---|---|---|---|
| P1 cl101 | E05 | 1616900 | sample | 1.0308575 | 0.5080989 |
| P1 cl101 | E06 | 1722700 | sample | 1.0983105 | 1.6187778 |
| P1 cl101 | E07 | 1716700 | sample | 1.0944852 | 1.5557904 |
| P1 cl101 | E08 | 1580900 | sample | 1.0079056 | 0.1301741 |
| P1 cl101 | E09 | 1535300 | sample | 0.9788333 | -0.3485306 |
| P1 cl101 | E10 | 1687400 | sample | 1.0758049 | 1.2482016 |
| P1 cl101 | E11 | 1641200 | sample | 1.0463500 | 0.7631981 |
| P1 cl101 | E12 | 1637200 | DMSO | 1.0437998 | 0.7212065 |
| P1 cl101 | F01 | 1432400 | DMSO | 0.9132292 | -1.4287656 |
| P1 cl101 | F02 | 1572700 | sample | 1.0026777 | 0.0440912 |
| P1 cl101 | F03 | 1572800 | sample | 1.0027415 | 0.0451410 |
| P1 cl101 | F04 | 1064400 | sample | 0.6786101 | -5.2919968 |
| P1 cl101 | F05 | 1616200 | sample | 1.0304112 | 0.5007503 |
| P1 cl101 | F06 | 1630400 | sample | 1.0394645 | 0.6498207 |
| P1 cl101 | F07 | 1695800 | sample | 1.0811603 | 1.3363840 |
| P1 cl101 | F08 | 1519800 | sample | 0.9689512 | -0.5112482 |
| P1 cl101 | F09 | 1451800 | sample | 0.9255977 | -1.2251062 |
| P1 cl101 | F10 | 1683400 | sample | 1.0732547 | 1.2062099 |
| P1 cl101 | F11 | 1649000 | sample | 1.0513229 | 0.8450818 |
| P1 cl101 | F12 | 427080 | SAL | 0.2722856 | -11.9825252 |
| P1 cl101 | G01 | 1384100 | empty | 0.8824354 | -1.9358147 |
| P1 cl101 | G02 | 1553600 | sample | 0.9905005 | -0.1564189 |
| P1 cl101 | G03 | 1582800 | sample | 1.0091170 | 0.1501201 |
| P1 cl101 | G04 | 1555300 | sample | 0.9915843 | -0.1385724 |
| P1 cl101 | G05 | 1308100 | sample | 0.8339815 | -2.7336559 |
| P1 cl101 | G06 | 1575700 | sample | 1.0045904 | 0.0755850 |
| P1 cl101 | G07 | 1546800 | sample | 0.9861651 | -0.2278047 |
| P1 cl101 | G08 | 1481800 | sample | 0.9447243 | -0.9101689 |
| P1 cl101 | G09 | 1584800 | sample | 1.0103921 | 0.1711159 |
| P1 cl101 | G10 | 1625800 | sample | 1.0365317 | 0.6015303 |
| P1 cl101 | G11 | 1604800 | sample | 1.0231431 | 0.3810742 |
| P1 cl101 | G12 | 451230 | SAL | 0.2876825 | -11.7290006 |
| P1 cl101 | H01 | 1285300 | empty | 0.8194453 | -2.9730083 |
| P1 cl101 | H02 | 1456200 | sample | 0.9284029 | -1.1789154 |
| P1 cl101 | H03 | 1371500 | sample | 0.8744023 | -2.0680884 |
| P1 cl101 | H04 | 1425900 | sample | 0.9090851 | -1.4970021 |
| P1 cl101 | H05 | 1380400 | sample | 0.8800765 | -1.9746570 |
| P1 cl101 | H06 | 1414900 | sample | 0.9020720 | -1.6124791 |
| P1 cl101 | H07 | 957300 | sample | 0.6103283 | -6.4163230 |
| P1 cl101 | H08 | 1544600 | sample | 0.9847625 | -0.2509001 |
| P1 cl101 | H09 | 890100 | sample | 0.5674849 | -7.1217826 |
| P1 cl101 | H10 | 1524300 | sample | 0.9718202 | -0.4640077 |
| P1 cl101 | H11 | 1507300 | sample | 0.9609818 | -0.6424721 |
| P1 cl101 | H12 | 400550 | SAL | 0.2553714 | -12.2610347 |
| P2 cl101 | A01 | 342790 | SAL | 0.2344344 | -10.3287510 |
| P2 cl101 | A02 | 1404800 | sample | 0.9607441 | -0.5296275 |
| P2 cl101 | A03 | 1317300 | sample | 0.9009027 | -1.3369865 |
| P2 cl101 | A04 | 1355400 | sample | 0.9269594 | -0.9854393 |
| P2 cl101 | A05 | 1437200 | sample | 0.9829025 | -0.2306740 |
| P2 cl101 | A06 | 1498800 | sample | 1.0250308 | 0.3377067 |
| P2 cl101 | A07 | 1452800 | sample | 0.9935713 | -0.0867334 |
| P2 cl101 | A08 | 1419700 | sample | 0.9709342 | -0.3921458 |
| P2 cl101 | A09 | 1392200 | sample | 0.9521269 | -0.6458872 |
| P2 cl101 | A10 | 1392200 | sample | 0.9521269 | -0.6458872 |
| P2 cl101 | A11 | 1439300 | sample | 0.9843387 | -0.2112974 |
| P2 cl101 | A12 | 1514300 | empty | 1.0356312 | 0.4807246 |
| P2 cl101 | B01 | 378690 | SAL | 0.2589865 | -9.9975032 |
| P2 cl101 | B02 | 1477000 | sample | 1.0101217 | 0.1365590 |
| P2 cl101 | B03 | 1501200 | sample | 1.0266721 | 0.3598514 |
| P2 cl101 | B04 | 1520900 | sample | 1.0401450 | 0.5416225 |
| P2 cl101 | B05 | 1496900 | sample | 1.0237314 | 0.3201755 |
| P2 cl101 | B06 | 1609500 | sample | 1.1007386 | 1.3591312 |

*(continued)*

| Plate | Well | Value | Treatment | Norm | Score |
|---|---|---|---|---|---|
| P2 cl101 | B07 | 1585300 | sample | 1.0841882 | 1.1358387 |
| P2 cl101 | B08 | 1224700 | sample | 0.8375735 | -2.1914029 |
| P2 cl101 | B09 | 1534100 | sample | 1.0491725 | 0.6634184 |
| P2 cl101 | B10 | 1646400 | sample | 1.1259746 | 1.6996060 |
| P2 cl101 | B11 | 517490 | sample | 0.3539119 | -8.7168012 |
| P2 cl101 | B12 | 1609100 | empty | 1.1004651 | 1.3554404 |
| P2 cl101 | C01 | 371680 | SAL | 0.2541923 | -10.0621842 |
| P2 cl101 | C02 | 1313600 | sample | 0.8983723 | -1.3711262 |
| P2 cl101 | C03 | 1454600 | sample | 0.9948024 | -0.0701249 |
| P2 cl101 | C04 | 1361700 | sample | 0.9312680 | -0.9273095 |
| P2 cl101 | C05 | 1518900 | sample | 1.0387772 | 0.5231686 |
| P2 cl101 | C06 | 1642500 | sample | 1.1233073 | 1.6636208 |
| P2 cl101 | C07 | 1630700 | sample | 1.1152373 | 1.5547427 |
| P2 cl101 | C08 | 1529100 | sample | 1.0457530 | 0.6172836 |
| P2 cl101 | C09 | 1469900 | sample | 1.0052660 | 0.0710476 |
| P2 cl101 | C10 | 1572000 | sample | 1.0750923 | 1.0131202 |
| P2 cl101 | C11 | 1612900 | sample | 1.1030639 | 1.3905028 |
| P2 cl101 | C12 | 1546300 | DMSO | 1.0575161 | 0.7759873 |
| P2 cl101 | D01 | 1330100 | DMSO | 0.9096567 | -1.2188814 |
| P2 cl101 | D02 | 1503300 | sample | 1.0281083 | 0.3792280 |
| P2 cl101 | D03 | 1469200 | sample | 1.0047873 | 0.0645887 |
| P2 cl101 | D04 | 1387900 | sample | 0.9491862 | -0.6855631 |
| P2 cl101 | D05 | 1480500 | sample | 1.0125154 | 0.1688534 |
| P2 cl101 | D06 | 1455200 | sample | 0.9952127 | -0.0645887 |
| P2 cl101 | D07 | 1615000 | sample | 1.1045001 | 1.4098795 |
| P2 cl101 | D08 | 1532000 | sample | 1.0477363 | 0.6440418 |
| P2 cl101 | D09 | 1591000 | sample | 1.0880864 | 1.1884324 |
| P2 cl101 | D10 | 1608500 | sample | 1.1000547 | 1.3499042 |
| P2 cl101 | D11 | 1680300 | sample | 1.1491588 | 2.0123999 |
| P2 cl101 | D12 | 1590100 | DMSO | 1.0874709 | 1.1801282 |
| P2 cl101 | E01 | 1407500 | DMSO | 0.9625906 | -0.5047147 |
| P2 cl101 | E02 | 1486600 | sample | 1.0166872 | 0.2251378 |
| P2 cl101 | E03 | 1520200 | sample | 1.0396663 | 0.5351637 |
| P2 cl101 | E04 | 1431400 | sample | 0.9789359 | -0.2841904 |
| P2 cl101 | E05 | 1493100 | sample | 1.0211325 | 0.2851131 |
| P2 cl101 | E06 | 600750 | sample | 0.4108535 | -7.9485645 |
| P2 cl101 | E07 | 1570600 | sample | 1.0741349 | 1.0002024 |
| P2 cl101 | E08 | 1569000 | sample | 1.0730406 | 0.9854393 |
| P2 cl101 | E09 | 1549200 | sample | 1.0594994 | 0.8027455 |
| P2 cl101 | E10 | 1585300 | sample | 1.0841882 | 1.1358387 |
| P2 cl101 | E11 | 1711300 | sample | 1.1703597 | 2.2984357 |
| P2 cl101 | E12 | 1587500 | DMSO | 1.0856928 | 1.1561381 |
| P2 cl101 | F01 | 1354000 | DMSO | 0.9260019 | -0.9983570 |
| P2 cl101 | F02 | 1511100 | sample | 1.0334428 | 0.4511983 |
| P2 cl101 | F03 | 1445700 | sample | 0.9887156 | -0.1522448 |
| P2 cl101 | F04 | 1177800 | sample | 0.8054986 | -2.6241474 |
| P2 cl101 | F05 | 1636200 | sample | 1.1189988 | 1.6054910 |
| P2 cl101 | F06 | 1593400 | sample | 1.0897278 | 1.2105771 |
| P2 cl101 | F07 | 258180 | sample | 0.1765696 | -11.1094441 |
| P2 cl101 | F08 | 1527600 | sample | 1.0447271 | 0.6034432 |
| P2 cl101 | F09 | 243350 | sample | 0.1664273 | -11.2462799 |
| P2 cl101 | F10 | 1488900 | sample | 1.0182602 | 0.2463598 |
| P2 cl101 | F11 | 1560300 | sample | 1.0670907 | 0.9051648 |
| P2 cl101 | F12 | 428490 | SAL | 0.2930447 | -9.5380006 |
| P2 cl101 | G01 | 1287600 | empty | 0.8805909 | -1.6110272 |
| P2 cl101 | G02 | 1405000 | sample | 0.9608809 | -0.5277821 |
| P2 cl101 | G03 | 1435800 | sample | 0.9819450 | -0.2435917 |
| P2 cl101 | G04 | 257790 | sample | 0.1763028 | -11.1130426 |
| P2 cl101 | G05 | 1373900 | sample | 0.9396115 | -0.8147405 |
| P2 cl101 | G06 | 1506200 | sample | 1.0300916 | 0.4059862 |
| P2 cl101 | G07 | 818910 | sample | 0.5600533 | -5.9356110 |

*(continued)*

| Plate | Well | Value | Treatment | Norm | Score |
|---|---|---|---|---|---|
| P2 cl101 | G08 | 1450500 | sample | 0.9919984 | -0.1079554 |
| P2 cl101 | G09 | 1485700 | sample | 1.0160717 | 0.2168336 |
| P2 cl101 | G10 | 767610 | sample | 0.5249692 | -6.4089540 |
| P2 cl101 | G11 | 1450600 | sample | 0.9920667 | -0.1070327 |
| P2 cl101 | G12 | 410120 | SAL | 0.2804815 | -9.7074998 |
| P2 cl101 | H01 | 1134700 | empty | 0.7760224 | -3.0218293 |
| P2 cl101 | H02 | 1404400 | sample | 0.9604705 | -0.5333183 |
| P2 cl101 | H03 | 996150 | sample | 0.6812680 | -4.3002246 |
| P2 cl101 | H04 | 1400700 | sample | 0.9579401 | -0.5674580 |
| P2 cl101 | H05 | 1397300 | sample | 0.9556148 | -0.5988297 |
| P2 cl101 | H06 | 818830 | sample | 0.5599986 | -5.9363491 |
| P2 cl101 | H07 | 1371000 | sample | 0.9376282 | -0.8414987 |
| P2 cl101 | H08 | 1359400 | sample | 0.9296950 | -0.9485315 |
| P2 cl101 | H09 | 1378700 | sample | 0.9428943 | -0.7704511 |
| P2 cl101 | H10 | 1404100 | sample | 0.9602654 | -0.5360864 |
| P2 cl101 | H11 | 1367800 | sample | 0.9354397 | -0.8710250 |
| P2 cl101 | H12 | 398130 | SAL | 0.2722815 | -9.8181311 |
| P3 cl101 | A01 | 389150 | SAL | 0.2525472 | -6.1875327 |
| P3 cl101 | A02 | 1551000 | sample | 1.0065546 | 0.0542601 |
| P3 cl101 | A03 | 1577000 | sample | 1.0234279 | 0.1939396 |
| P3 cl101 | A04 | 1627200 | sample | 1.0560062 | 0.4636285 |
| P3 cl101 | A05 | 1482800 | sample | 0.9622948 | -0.3121299 |
| P3 cl101 | A06 | 1527800 | sample | 0.9914985 | -0.0703770 |
| P3 cl101 | A07 | 1628600 | sample | 1.0569148 | 0.4711497 |
| P3 cl101 | A08 | 1540300 | sample | 0.9996106 | -0.0032234 |
| P3 cl101 | A09 | 1447600 | sample | 0.9394510 | -0.5012345 |
| P3 cl101 | A10 | 1539700 | sample | 0.9992212 | -0.0064467 |
| P3 cl101 | A11 | 1679900 | sample | 1.0902070 | 0.7467480 |
| P3 cl101 | A12 | 1608000 | empty | 1.0435460 | 0.3604805 |
| P3 cl101 | B01 | 375180 | SAL | 0.2434811 | -6.2625836 |
| P3 cl101 | B02 | 1685400 | sample | 1.0937764 | 0.7762956 |
| P3 cl101 | B03 | 1739700 | sample | 1.1290155 | 1.0680109 |
| P3 cl101 | B04 | 1591200 | sample | 1.0326433 | 0.2702261 |
| P3 cl101 | B05 | 1241500 | sample | 0.8056980 | -1.6084630 |
| P3 cl101 | B06 | 1697300 | sample | 1.1014991 | 0.8402258 |
| P3 cl101 | B07 | 1746300 | sample | 1.1332987 | 1.1034680 |
| P3 cl101 | B08 | 1702400 | sample | 1.1048089 | 0.8676245 |
| P3 cl101 | B09 | 1661100 | sample | 1.0780064 | 0.6457490 |
| P3 cl101 | B10 | 1760200 | sample | 1.1423194 | 1.1781428 |
| P3 cl101 | B11 | 1846800 | sample | 1.1985203 | 1.6433829 |
| P3 cl101 | B12 | 1814900 | empty | 1.1778182 | 1.4720069 |
| P3 cl101 | C01 | 366760 | SAL | 0.2380167 | -6.3078182 |
| P3 cl101 | C02 | 1650700 | sample | 1.0712571 | 0.5898772 |
| P3 cl101 | C03 | 1621900 | sample | 1.0525667 | 0.4351553 |
| P3 cl101 | C04 | 1729100 | sample | 1.1221364 | 1.0110646 |
| P3 cl101 | C05 | 1735300 | sample | 1.1261600 | 1.0443728 |
| P3 cl101 | C06 | 1795100 | sample | 1.1649685 | 1.3656356 |
| P3 cl101 | C07 | 1746900 | sample | 1.1336881 | 1.1066913 |
| P3 cl101 | C08 | 1747100 | sample | 1.1338179 | 1.1077658 |
| P3 cl101 | C09 | 1736200 | sample | 1.1267441 | 1.0492078 |
| P3 cl101 | C10 | 1756400 | sample | 1.1398533 | 1.1577281 |
| P3 cl101 | C11 | 1758100 | sample | 1.1409566 | 1.1668610 |
| P3 cl101 | C12 | 1578300 | DMSO | 1.0242715 | 0.2009236 |
| P3 cl101 | D01 | 1375200 | DMSO | 0.8924654 | -0.8901881 |
| P3 cl101 | D02 | 1595200 | sample | 1.0352391 | 0.2917152 |
| P3 cl101 | D03 | 1649600 | sample | 1.0705432 | 0.5839677 |
| P3 cl101 | D04 | 1439700 | sample | 0.9343241 | -0.5436755 |
| P3 cl101 | D05 | 1760400 | sample | 1.1424492 | 1.1792172 |
| P3 cl101 | D06 | 1668600 | sample | 1.0828736 | 0.6860412 |
| P3 cl101 | D07 | 1696600 | sample | 1.1010448 | 0.8364652 |
| P3 cl101 | D08 | 1664300 | sample | 1.0800831 | 0.6629403 |

*(continued)*

| Plate | Well | Value | Treatment | Norm | Score |
|---|---|---|---|---|---|
| P3 cl101 | D09 | 1659200 | sample | 1.0767733 | 0.6355417 |
| P3 cl101 | D10 | 1652700 | sample | 1.0725550 | 0.6006218 |
| P3 cl101 | D11 | 382490 | sample | 0.2482251 | -6.2233122 |
| P3 cl101 | D12 | 1684000 | DMSO | 1.0928678 | 0.7687744 |
| P3 cl101 | E01 | 1381200 | DMSO | 0.8963593 | -0.8579544 |
| P3 cl101 | E02 | 1512700 | sample | 0.9816990 | -0.1514985 |
| P3 cl101 | E03 | 1544300 | sample | 1.0022065 | 0.0182658 |
| P3 cl101 | E04 | 1496000 | sample | 0.9708612 | -0.2412157 |
| P3 cl101 | E05 | 1534300 | sample | 0.9957168 | -0.0354571 |
| P3 cl101 | E06 | 1576600 | sample | 1.0231683 | 0.1917907 |
| P3 cl101 | E07 | 1672300 | sample | 1.0852748 | 0.7059186 |
| P3 cl101 | E08 | 1649500 | sample | 1.0704783 | 0.5834305 |
| P3 cl101 | E09 | 1587300 | sample | 1.0301123 | 0.2492742 |
| P3 cl101 | E10 | 1592100 | sample | 1.0332273 | 0.2750611 |
| P3 cl101 | E11 | 685730 | sample | 0.4450191 | -4.5942195 |
| P3 cl101 | E12 | 1470900 | DMSO | 0.9545720 | -0.3760602 |
| P3 cl101 | F01 | 1327300 | DMSO | 0.8613797 | -1.1475207 |
| P3 cl101 | F02 | 1463100 | sample | 0.9495100 | -0.4179640 |
| P3 cl101 | F03 | 1435700 | sample | 0.9317282 | -0.5651647 |
| P3 cl101 | F04 | 1484800 | sample | 0.9635927 | -0.3013854 |
| P3 cl101 | F05 | 1513100 | sample | 0.9819586 | -0.1493496 |
| P3 cl101 | F06 | 1475600 | sample | 0.9576222 | -0.3508104 |
| P3 cl101 | F07 | 1509800 | sample | 0.9798170 | -0.1670782 |
| P3 cl101 | F08 | 547760 | sample | 0.3554806 | -5.3354341 |
| P3 cl101 | F09 | 1495400 | sample | 0.9704718 | -0.2444391 |
| P3 cl101 | F10 | 1600500 | sample | 1.0386787 | 0.3201884 |
| P3 cl101 | F11 | 1541500 | sample | 1.0003894 | 0.0032234 |
| P3 cl101 | F12 | 162120 | SAL | 0.1052112 | -7.4072033 |
| P3 cl101 | G01 | 1376900 | empty | 0.8935687 | -0.8810552 |
| P3 cl101 | G02 | 1371500 | sample | 0.8900642 | -0.9100656 |
| P3 cl101 | G03 | 734010 | sample | 0.4763515 | -4.3348455 |
| P3 cl101 | G04 | 1343800 | sample | 0.8720877 | -1.0588780 |
| P3 cl101 | G05 | 1403000 | sample | 0.9105068 | -0.7408385 |
| P3 cl101 | G06 | 1327500 | sample | 0.8615095 | -1.1464463 |
| P3 cl101 | G07 | 1338000 | sample | 0.8683237 | -1.0900372 |
| P3 cl101 | G08 | 1505800 | sample | 0.9772211 | -0.1885673 |
| P3 cl101 | G09 | 1444800 | sample | 0.9376339 | -0.5162769 |
| P3 cl101 | G10 | 1495400 | sample | 0.9704718 | -0.2444391 |
| P3 cl101 | G11 | 1484900 | sample | 0.9636576 | -0.3008481 |
| P3 cl101 | G12 | 284370 | SAL | 0.1845480 | -6.7504411 |
| P3 cl101 | H01 | 1022900 | empty | 0.6638328 | -2.7828452 |
| P3 cl101 | H02 | 1204000 | sample | 0.7813615 | -1.8099238 |
| P3 cl101 | H03 | 1222200 | sample | 0.7931728 | -1.7121482 |
| P3 cl101 | H04 | 1217600 | sample | 0.7901876 | -1.7368607 |
| P3 cl101 | H05 | 1186400 | sample | 0.7699396 | -1.9044761 |
| P3 cl101 | H06 | 760780 | sample | 0.4937244 | -4.1910293 |
| P3 cl101 | H07 | 763860 | sample | 0.4957233 | -4.1744827 |
| P3 cl101 | H08 | 1011100 | sample | 0.6561750 | -2.8462382 |
| P3 cl101 | H09 | 1274500 | sample | 0.8271140 | -1.4311775 |
| P3 cl101 | H10 | 1321200 | sample | 0.8574210 | -1.1802917 |
| P3 cl101 | H11 | 1323100 | sample | 0.8586540 | -1.1700843 |
| P3 cl101 | H12 | 310830 | SAL | 0.2017198 | -6.6082903 |
| P4 cl101 | A01 | 348140 | SAL | 0.2402374 | -7.4747969 |
| P4 cl101 | A02 | 1471000 | sample | 1.0150778 | 0.1483404 |
| P4 cl101 | A03 | 1347100 | sample | 0.9295794 | -0.6928212 |
| P4 cl101 | A04 | 1277600 | sample | 0.8816203 | -1.1646592 |
| P4 cl101 | A05 | 1391400 | sample | 0.9601491 | -0.3920668 |
| P4 cl101 | A06 | 1403600 | sample | 0.9685678 | -0.3092406 |
| P4 cl101 | A07 | 1470200 | sample | 1.0145258 | 0.1429092 |
| P4 cl101 | A08 | 616330 | sample | 0.4253045 | -5.6540452 |
| P4 cl101 | A09 | 1356900 | sample | 0.9363420 | -0.6262886 |
| P4 cl101 | A10 | 1325100 | sample | 0.9143981 | -0.8421800 |

*(continued)*

| Plate | Well | Value | Treatment | Norm | Score |
|---|---|---|---|---|---|
| P4 cl101 | A11 | 1355500 | sample | 0.9353759 | -0.6357933 |
| P4 cl101 | A12 | 1506400 | empty | 1.0395059 | 0.3886723 |
| P4 cl101 | B01 | 348980 | SAL | 0.2408170 | -7.4690941 |
| P4 cl101 | B02 | 1501500 | sample | 1.0361246 | 0.3554061 |
| P4 cl101 | B03 | 1330200 | sample | 0.9179174 | -0.8075559 |
| P4 cl101 | B04 | 1466500 | sample | 1.0119725 | 0.1177898 |
| P4 cl101 | B05 | 1276800 | sample | 0.8810682 | -1.1700904 |
| P4 cl101 | B06 | 1521800 | sample | 1.0501328 | 0.4932235 |
| P4 cl101 | B07 | 1305600 | sample | 0.9009419 | -0.9745662 |
| P4 cl101 | B08 | 1324200 | sample | 0.9137770 | -0.8482901 |
| P4 cl101 | B09 | 1578500 | sample | 1.0892592 | 0.8781618 |
| P4 cl101 | B10 | 1524400 | sample | 1.0519270 | 0.5108750 |
| P4 cl101 | B11 | 1598900 | sample | 1.1033364 | 1.0166582 |
| P4 cl101 | B12 | 1612900 | empty | 1.1129973 | 1.1117047 |
| P4 cl101 | C01 | 371830 | SAL | 0.2565849 | -7.3139646 |
| P4 cl101 | C02 | 1492000 | sample | 1.0295691 | 0.2909102 |
| P4 cl101 | C03 | 1477300 | sample | 1.0194252 | 0.1911114 |
| P4 cl101 | C04 | 1393000 | sample | 0.9612531 | -0.3812044 |
| P4 cl101 | C05 | 1432200 | sample | 0.9883035 | -0.1150742 |
| P4 cl101 | C06 | 1517800 | sample | 1.0473726 | 0.4660673 |
| P4 cl101 | C07 | 1571900 | sample | 1.0847048 | 0.8333542 |
| P4 cl101 | C08 | 998830 | sample | 0.6892523 | -3.0572388 |
| P4 cl101 | C09 | 1560200 | sample | 1.0766311 | 0.7539225 |
| P4 cl101 | C10 | 1426400 | sample | 0.9843011 | -0.1544506 |
| P4 cl101 | C11 | 1323100 | sample | 0.9130180 | -0.8557580 |
| P4 cl101 | C12 | 1586200 | DMSO | 1.0945727 | 0.9304374 |
| P4 cl101 | D01 | 1350900 | DMSO | 0.9322016 | -0.6670228 |
| P4 cl101 | D02 | 1481700 | sample | 1.0224614 | 0.2209831 |
| P4 cl101 | D03 | 1428100 | sample | 0.9854742 | -0.1429092 |
| P4 cl101 | D04 | 1454700 | sample | 1.0038298 | 0.0376792 |
| P4 cl101 | D05 | 1590700 | sample | 1.0976779 | 0.9609881 |
| P4 cl101 | D06 | 1566900 | sample | 1.0812545 | 0.7994090 |
| P4 cl101 | D07 | 1517900 | sample | 1.0474416 | 0.4667462 |
| P4 cl101 | D08 | 1560900 | sample | 1.0771142 | 0.7586748 |
| P4 cl101 | D09 | 1572900 | sample | 1.0853949 | 0.8401432 |
| P4 cl101 | D10 | 1643000 | sample | 1.1337681 | 1.3160547 |
| P4 cl101 | D11 | 1690100 | sample | 1.1662699 | 1.6358183 |
| P4 cl101 | D12 | 1488700 | DMSO | 1.0272919 | 0.2685064 |
| P4 cl101 | E01 | 1395600 | DMSO | 0.9630473 | -0.3635529 |
| P4 cl101 | E02 | 1572900 | sample | 1.0853949 | 0.8401432 |
| P4 cl101 | E03 | 1485700 | sample | 1.0252217 | 0.2481393 |
| P4 cl101 | E04 | 723170 | sample | 0.4990305 | -4.9287046 |
| P4 cl101 | E05 | 1616500 | sample | 1.1154815 | 1.1361452 |
| P4 cl101 | E06 | 1625800 | sample | 1.1218990 | 1.1992833 |
| P4 cl101 | E07 | 1182600 | sample | 0.8160646 | -1.8096176 |
| P4 cl101 | E08 | 1424700 | sample | 0.9831280 | -0.1659919 |
| P4 cl101 | E09 | 1492700 | sample | 1.0300521 | 0.2956625 |
| P4 cl101 | E10 | 1611700 | sample | 1.1121692 | 1.1035579 |
| P4 cl101 | E11 | 1606900 | sample | 1.1088569 | 1.0709705 |
| P4 cl101 | E12 | 1613700 | DMSO | 1.1135493 | 1.1171359 |
| P4 cl101 | F01 | 1369900 | DMSO | 0.9453128 | -0.5380311 |
| P4 cl101 | F02 | 1506300 | sample | 1.0394369 | 0.3879934 |
| P4 cl101 | F03 | 1443600 | sample | 0.9961702 | -0.0376792 |
| P4 cl101 | F04 | 1431400 | sample | 0.9877514 | -0.1205054 |
| P4 cl101 | F05 | 1375000 | sample | 0.9488321 | -0.5034070 |
| P4 cl101 | F06 | 1287600 | sample | 0.8885209 | -1.0967688 |
| P4 cl101 | F07 | 1525900 | sample | 1.0529621 | 0.5210585 |
| P4 cl101 | F08 | 934300 | sample | 0.6447228 | -3.4953354 |
| P4 cl101 | F09 | 1495400 | sample | 1.0319153 | 0.3139929 |
| P4 cl101 | F10 | 1511500 | sample | 1.0430252 | 0.4232964 |
| P4 cl101 | F11 | 1545800 | sample | 1.0666943 | 0.6561604 |
| P4 cl101 | F12 | 436820 | SAL | 0.3014319 | -6.8727451 |

*(continued)*

| Plate | Well | Value | Treatment | Norm | Score |
|---|---|---|---|---|---|
| P4 cl101 | G01 | 1343500 | empty | 0.9270952 | -0.7172617 |
| P4 cl101 | G02 | 1532500 | sample | 1.0575165 | 0.5658662 |
| P4 cl101 | G03 | 1513600 | sample | 1.0444743 | 0.4375534 |
| P4 cl101 | G04 | 1505700 | sample | 1.0390229 | 0.3839200 |
| P4 cl101 | G05 | 1440100 | sample | 0.9937550 | -0.0614408 |
| P4 cl101 | G06 | 1481600 | sample | 1.0223924 | 0.2203042 |
| P4 cl101 | G07 | 729880 | sample | 0.5036608 | -4.8831502 |
| P4 cl101 | G08 | 1478500 | sample | 1.0202533 | 0.1992582 |
| P4 cl101 | G09 | 726280 | sample | 0.5011766 | -4.9075907 |
| P4 cl101 | G10 | 693310 | sample | 0.4784253 | -5.1314252 |
| P4 cl101 | G11 | 1591600 | sample | 1.0982990 | 0.9670982 |
| P4 cl101 | G12 | 429200 | SAL | 0.2961736 | -6.9244776 |
| P4 cl101 | H01 | 1243800 | empty | 0.8582962 | -1.3941286 |
| P4 cl101 | H02 | 1131600 | sample | 0.7808715 | -2.1558585 |
| P4 cl101 | H03 | 1337500 | sample | 0.9229548 | -0.7579959 |
| P4 cl101 | H04 | 230550 | sample | 0.1590933 | -8.2731197 |
| P4 cl101 | H05 | 1436000 | sample | 0.9909257 | -0.0892758 |
| P4 cl101 | H06 | 1332100 | sample | 0.9192285 | -0.7946567 |
| P4 cl101 | H07 | 1429200 | sample | 0.9862333 | -0.1354413 |
| P4 cl101 | H08 | 1381400 | sample | 0.9532485 | -0.4599572 |
| P4 cl101 | H09 | 75570 | sample | 0.0521478 | -9.3252845 |
| P4 cl101 | H10 | 751380 | sample | 0.5184970 | -4.7371859 |
| P4 cl101 | H11 | 191850 | sample | 0.1323880 | -8.5358554 |
| P4 cl101 | H12 | 370980 | SAL | 0.2559983 | -7.3197353 |
| P5 cl101 | A01 | 307020 | SAL | 0.2102949 | -8.4069257 |
| P5 cl101 | A02 | 1368900 | sample | 0.9376349 | -0.6639177 |
| P5 cl101 | A03 | 1292300 | sample | 0.8851673 | -1.2224689 |
| P5 cl101 | A04 | 1311800 | sample | 0.8985239 | -1.0802790 |
| P5 cl101 | A05 | 1291100 | sample | 0.8843454 | -1.2312191 |
| P5 cl101 | A06 | 1537800 | sample | 1.0533237 | 0.5676660 |
| P5 cl101 | A07 | 1428500 | sample | 0.9784582 | -0.2293269 |
| P5 cl101 | A08 | 1351400 | sample | 0.9256481 | -0.7915240 |
| P5 cl101 | A09 | 1414700 | sample | 0.9690058 | -0.3299536 |
| P5 cl101 | A10 | 1332400 | sample | 0.9126340 | -0.9300681 |
| P5 cl101 | A11 | 1374000 | sample | 0.9411281 | -0.6267295 |
| P5 cl101 | A12 | 1440100 | empty | 0.9864036 | -0.1447421 |
| P5 cl101 | B01 | 331580 | SAL | 0.2271174 | -8.2278393 |
| P5 cl101 | B02 | 1470200 | sample | 1.0070208 | 0.0747409 |
| P5 cl101 | B03 | 1471400 | sample | 1.0078427 | 0.0834910 |
| P5 cl101 | B04 | 1453400 | sample | 0.9955135 | -0.0477612 |
| P5 cl101 | B05 | 1516200 | sample | 1.0385287 | 0.4101633 |
| P5 cl101 | B06 | 1585600 | sample | 1.0860646 | 0.9162137 |
| P5 cl101 | B07 | 1518900 | sample | 1.0403781 | 0.4298511 |
| P5 cl101 | B08 | 1382900 | sample | 0.9472242 | -0.5618326 |
| P5 cl101 | B09 | 1495000 | sample | 1.0240077 | 0.2555773 |
| P5 cl101 | B10 | 1579300 | sample | 1.0817494 | 0.8702754 |
| P5 cl101 | B11 | 1471400 | sample | 1.0078427 | 0.0834910 |
| P5 cl101 | B12 | 1671500 | empty | 1.1449022 | 1.5425786 |
| P5 cl101 | C01 | 365600 | SAL | 0.2504195 | -7.9797726 |
| P5 cl101 | C02 | 1415900 | sample | 0.9698277 | -0.3212034 |
| P5 cl101 | C03 | 1554800 | sample | 1.0649680 | 0.6916265 |
| P5 cl101 | C04 | 1359600 | sample | 0.9312648 | -0.7317313 |
| P5 cl101 | C05 | 1553900 | sample | 1.0643515 | 0.6850639 |
| P5 cl101 | C06 | 1579800 | sample | 1.0820919 | 0.8739213 |
| P5 cl101 | C07 | 1468800 | sample | 1.0060619 | 0.0645324 |
| P5 cl101 | C08 | 1503700 | sample | 1.0299668 | 0.3190159 |
| P5 cl101 | C09 | 1110000 | sample | 0.7603000 | -2.5517626 |
| P5 cl101 | C10 | 1578500 | sample | 1.0812014 | 0.8644419 |
| P5 cl101 | C11 | 1476200 | sample | 1.0111305 | 0.1184916 |
| P5 cl101 | C12 | 1574500 | DMSO | 1.0784616 | 0.8352748 |
| P5 cl101 | D01 | 1357400 | DMSO | 0.9297579 | -0.7477733 |

*(continued)*

| Plate | Well | Value | Treatment | Norm | Score |
|---|---|---|---|---|---|
| P5 cl101 | D02 | 1497700 | sample | 1.0258570 | 0.2752651 |
| P5 cl101 | D03 | 1446900 | sample | 0.9910613 | -0.0951579 |
| P5 cl101 | D04 | 1365100 | sample | 0.9350320 | -0.6916265 |
| P5 cl101 | D05 | 1543200 | sample | 1.0570225 | 0.6070417 |
| P5 cl101 | D06 | 672310 | sample | 0.4605021 | -5.7433070 |
| P5 cl101 | D07 | 1612200 | sample | 1.1042844 | 1.1101753 |
| P5 cl101 | D08 | 1574400 | sample | 1.0783931 | 0.8345456 |
| P5 cl101 | D09 | 1529400 | sample | 1.0475701 | 0.5064150 |
| P5 cl101 | D10 | 1468400 | sample | 1.0057879 | 0.0616156 |
| P5 cl101 | D11 | 1604000 | sample | 1.0986678 | 1.0503826 |
| P5 cl101 | D12 | 1504700 | DMSO | 1.0306517 | 0.3263077 |
| P5 cl101 | E01 | 1298900 | DMSO | 0.8896880 | -1.1743431 |
| P5 cl101 | E02 | 1537900 | sample | 1.0533922 | 0.5683952 |
| P5 cl101 | E03 | 1560300 | sample | 1.0687352 | 0.7317313 |
| P5 cl101 | E04 | 452580 | sample | 0.3099969 | -7.3455325 |
| P5 cl101 | E05 | 1191700 | sample | 0.8162608 | -1.9560232 |
| P5 cl101 | E06 | 1606100 | sample | 1.1001062 | 1.0656954 |
| P5 cl101 | E07 | 1543800 | sample | 1.0574335 | 0.6114168 |
| P5 cl101 | E08 | 1457900 | sample | 0.9985958 | -0.0149482 |
| P5 cl101 | E09 | 1359600 | sample | 0.9312648 | -0.7317313 |
| P5 cl101 | E10 | 1578900 | sample | 1.0814754 | 0.8673587 |
| P5 cl101 | E11 | 1527700 | sample | 1.0464057 | 0.4940189 |
| P5 cl101 | E12 | 1603100 | DMSO | 1.0980513 | 1.0438200 |
| P5 cl101 | F01 | 1327100 | DMSO | 0.9090037 | -0.9687146 |
| P5 cl101 | F02 | 1560000 | sample | 1.0685297 | 0.7295438 |
| P5 cl101 | F03 | 1429600 | sample | 0.9792116 | -0.2213059 |
| P5 cl101 | F04 | 130210 | sample | 0.0891880 | -9.6961875 |
| P5 cl101 | F05 | 1468900 | sample | 1.0061303 | 0.0652615 |
| P5 cl101 | F06 | 996490 | sample | 0.6825508 | -3.3794539 |
| P5 cl101 | F07 | 881780 | sample | 0.6039796 | -4.2158954 |
| P5 cl101 | F08 | 1478100 | sample | 1.0124319 | 0.1323460 |
| P5 cl101 | F09 | 1513400 | sample | 1.0366108 | 0.3897463 |
| P5 cl101 | F10 | 1618900 | sample | 1.1088736 | 1.1590303 |
| P5 cl101 | F11 | 1465000 | sample | 1.0034590 | 0.0368235 |
| P5 cl101 | F12 | 394790 | SAL | 0.2704134 | -7.7669252 |
| P5 cl101 | G01 | 1214400 | empty | 0.8318093 | -1.7904995 |
| P5 cl101 | G02 | 1470900 | sample | 1.0075003 | 0.0798451 |
| P5 cl101 | G03 | 1339800 | sample | 0.9177027 | -0.8761088 |
| P5 cl101 | G04 | 1349300 | sample | 0.9242097 | -0.8068368 |
| P5 cl101 | G05 | 1490700 | sample | 1.0210624 | 0.2242226 |
| P5 cl101 | G06 | 138460 | sample | 0.0948389 | -9.6360302 |
| P5 cl101 | G07 | 712250 | sample | 0.4878592 | -5.4520729 |
| P5 cl101 | G08 | 1072800 | sample | 0.7348197 | -2.8230173 |
| P5 cl101 | G09 | 1426000 | sample | 0.9767458 | -0.2475563 |
| P5 cl101 | G10 | 1202800 | sample | 0.8238638 | -1.8750843 |
| P5 cl101 | G11 | 1550700 | sample | 1.0621597 | 0.6617301 |
| P5 cl101 | G12 | 411110 | SAL | 0.2815918 | -7.6479231 |
| P5 cl101 | H01 | 1109300 | empty | 0.7598205 | -2.5568669 |
| P5 cl101 | H02 | 1370900 | sample | 0.9390048 | -0.6493341 |
| P5 cl101 | H03 | 1317000 | sample | 0.9020857 | -1.0423617 |
| P5 cl101 | H04 | 1381600 | sample | 0.9463338 | -0.5713119 |
| P5 cl101 | H05 | 1462000 | sample | 1.0014042 | 0.0149482 |
| P5 cl101 | H06 | 1397900 | sample | 0.9574985 | -0.4524557 |
| P5 cl101 | H07 | 726960 | sample | 0.4979349 | -5.3448106 |
| P5 cl101 | H08 | 104890 | sample | 0.0718449 | -9.8808157 |
| P5 cl101 | H09 | 1261100 | sample | 0.8637967 | -1.4499728 |
| P5 cl101 | H10 | 265010 | sample | 0.1815199 | -8.7132539 |
| P5 cl101 | H11 | 1496700 | sample | 1.0251721 | 0.2679734 |
| P5 cl101 | H12 | 349080 | SAL | 0.2391041 | -8.1002330 |
| P6 cl101 | A01 | 384300 | SAL | 0.2526627 | -11.6165704 |
| P6 cl101 | A02 | 1423500 | sample | 0.9358974 | -0.9964068 |
| P6 cl101 | A03 | 1467100 | sample | 0.9645628 | -0.5508341 |

*(continued)*

| Plate | Well | Value | Treatment | Norm | Score |
|---|---|---|---|---|---|
| P6 cl101 | A04 | 1480600 | sample | 0.9734385 | -0.4128701 |
| P6 cl101 | A05 | 1366300 | sample | 0.8982906 | -1.5809655 |
| P6 cl101 | A06 | 1583700 | sample | 1.0412229 | 0.6407662 |
| P6 cl101 | A07 | 1450300 | sample | 0.9535174 | -0.7225227 |
| P6 cl101 | A08 | 1439500 | sample | 0.9464168 | -0.8328939 |
| P6 cl101 | A09 | 1181500 | sample | 0.7767916 | -3.4695396 |
| P6 cl101 | A10 | 582680 | sample | 0.3830901 | -9.5892147 |
| P6 cl101 | A11 | 1230700 | sample | 0.8091387 | -2.9667374 |
| P6 cl101 | A12 | 1342200 | empty | 0.8824458 | -1.8272568 |
| P6 cl101 | B01 | 1527100 | SAL | 1.0040105 | 0.0623393 |
| P6 cl101 | B02 | 1656400 | sample | 1.0890204 | 1.3837280 |
| P6 cl101 | B03 | 1483500 | sample | 0.9753452 | -0.3832334 |
| P6 cl101 | B04 | 1537500 | sample | 1.0108481 | 0.1686227 |
| P6 cl101 | B05 | 1208800 | sample | 0.7947403 | -3.1905457 |
| P6 cl101 | B06 | 1598600 | sample | 1.0510191 | 0.7930376 |
| P6 cl101 | B07 | 1456000 | sample | 0.9572650 | -0.6642712 |
| P6 cl101 | B08 | 1448200 | sample | 0.9521368 | -0.7439837 |
| P6 cl101 | B09 | 1586700 | sample | 1.0431953 | 0.6714249 |
| P6 cl101 | B10 | 1690800 | sample | 1.1116371 | 1.7352808 |
| P6 cl101 | B11 | 1612100 | sample | 1.0598948 | 0.9310016 |
| P6 cl101 | B12 | 1646500 | empty | 1.0825115 | 1.2825544 |
| P6 cl101 | C01 | 1539200 | SAL | 1.0119658 | 0.1859959 |
| P6 cl101 | C02 | 1356100 | sample | 0.8915845 | -1.6852049 |
| P6 cl101 | C03 | 1459600 | sample | 0.9596318 | -0.6274808 |
| P6 cl101 | C04 | 1558400 | sample | 1.0245891 | 0.3822114 |
| P6 cl101 | C05 | 1471800 | sample | 0.9676529 | -0.5028022 |
| P6 cl101 | C06 | 1315400 | sample | 0.8648258 | -2.1011409 |
| P6 cl101 | C07 | 1566600 | sample | 1.0299803 | 0.4660118 |
| P6 cl101 | C08 | 1521000 | sample | 1.0000000 | 0.0000000 |
| P6 cl101 | C09 | 1463700 | sample | 0.9623274 | -0.5855806 |
| P6 cl101 | C10 | 1719600 | sample | 1.1305720 | 2.0296040 |
| P6 cl101 | C11 | 1731200 | sample | 1.1381986 | 2.1481509 |
| P6 cl101 | C12 | 1635900 | DMSO | 1.0755424 | 1.1742271 |
| P6 cl101 | D01 | 1415400 | DMSO | 0.9305720 | -1.0791852 |
| P6 cl101 | D02 | 1495800 | sample | 0.9834320 | -0.2575328 |
| P6 cl101 | D03 | 1587100 | sample | 1.0434583 | 0.6755127 |
| P6 cl101 | D04 | 1494500 | sample | 0.9825773 | -0.2708183 |
| P6 cl101 | D05 | 1535700 | sample | 1.0096647 | 0.1502275 |
| P6 cl101 | D06 | 1462900 | sample | 0.9618014 | -0.5937563 |
| P6 cl101 | D07 | 1589800 | sample | 1.0452334 | 0.7031055 |
| P6 cl101 | D08 | 1556400 | sample | 1.0232742 | 0.3617723 |
| P6 cl101 | D09 | 271260 | sample | 0.1783432 | -12.7717891 |
| P6 cl101 | D10 | 1585700 | sample | 1.0425378 | 0.6612053 |
| P6 cl101 | D11 | 1552000 | sample | 1.0203813 | 0.3168063 |
| P6 cl101 | D12 | 1503400 | DMSO | 0.9884287 | -0.1798642 |
| P6 cl101 | E01 | 1464900 | DMSO | 0.9631164 | -0.5733171 |
| P6 cl101 | E02 | 1527300 | sample | 1.0041420 | 0.0643832 |
| P6 cl101 | E03 | 1577200 | sample | 1.0369494 | 0.5743391 |
| P6 cl101 | E04 | 1448100 | sample | 0.9520710 | -0.7450057 |
| P6 cl101 | E05 | 1664400 | sample | 1.0942801 | 1.4654845 |
| P6 cl101 | E06 | 1611600 | sample | 1.0595661 | 0.9258919 |
| P6 cl101 | E07 | 1694700 | sample | 1.1142012 | 1.7751370 |
| P6 cl101 | E08 | 1455100 | sample | 0.9566732 | -0.6734688 |
| P6 cl101 | E09 | 855780 | sample | 0.5626430 | -6.7982537 |
| P6 cl101 | E10 | 1553300 | sample | 1.0212360 | 0.3300917 |
| P6 cl101 | E11 | 1610600 | sample | 1.0589086 | 0.9156723 |
| P6 cl101 | E12 | 1613000 | DMSO | 1.0604865 | 0.9401992 |
| P6 cl101 | F01 | 1386700 | DMSO | 0.9117028 | -1.3724865 |
| P6 cl101 | F02 | 1534800 | sample | 1.0090730 | 0.1410299 |
| P6 cl101 | F03 | 1545900 | sample | 1.0163708 | 0.2544670 |
| P6 cl101 | F04 | 1452400 | sample | 0.9548981 | -0.7010616 |
| P6 cl101 | F05 | 1616400 | sample | 1.0627219 | 0.9749457 |

*(continued)*

| Plate | Well | Value | Treatment | Norm | Score |
|---|---|---|---|---|---|
| P6 cl101 | F06 | 1527100 | sample | 1.0040105 | 0.0623393 |
| P6 cl101 | F07 | 1573400 | sample | 1.0344510 | 0.5355048 |
| P6 cl101 | F08 | 1683100 | sample | 1.1065746 | 1.6565902 |
| P6 cl101 | F09 | 1557000 | sample | 1.0236686 | 0.3679041 |
| P6 cl101 | F10 | 1593100 | sample | 1.0474030 | 0.7368301 |
| P6 cl101 | F11 | 1688800 | sample | 1.1103222 | 1.7148417 |
| P6 cl101 | F12 | 428060 | SAL | 0.2814333 | -11.1693626 |
| P6 cl101 | G01 | 1448200 | empty | 0.9521368 | -0.7439837 |
| P6 cl101 | G02 | 1461700 | sample | 0.9610125 | -0.6060197 |
| P6 cl101 | G03 | 1669900 | sample | 1.0978961 | 1.5216920 |
| P6 cl101 | G04 | 1447300 | sample | 0.9515450 | -0.7531813 |
| P6 cl101 | G05 | 1488600 | sample | 0.9786982 | -0.3311136 |
| P6 cl101 | G06 | 1521000 | sample | 1.0000000 | 0.0000000 |
| P6 cl101 | G07 | 1506900 | sample | 0.9907298 | -0.1440958 |
| P6 cl101 | G08 | 1461000 | sample | 0.9605523 | -0.6131734 |
| P6 cl101 | G09 | 1496400 | sample | 0.9838264 | -0.2514011 |
| P6 cl101 | G10 | 1637400 | sample | 1.0765286 | 1.1895564 |
| P6 cl101 | G11 | 1536500 | sample | 1.0101907 | 0.1584031 |
| P6 cl101 | G12 | 418910 | SAL | 0.2754175 | -11.2628715 |
| P6 cl101 | H01 | 1370700 | empty | 0.9011834 | -1.5359994 |
| P6 cl101 | H02 | 1541900 | sample | 1.0137410 | 0.2135887 |
| P6 cl101 | H03 | 1532700 | sample | 1.0076923 | 0.1195688 |
| P6 cl101 | H04 | 1521000 | sample | 1.0000000 | 0.0000000 |
| P6 cl101 | H05 | 1507700 | sample | 0.9912558 | -0.1359201 |
| P6 cl101 | H06 | 1426000 | sample | 0.9375411 | -0.9708579 |
| P6 cl101 | H07 | 1219500 | sample | 0.8017751 | -3.0811964 |
| P6 cl101 | H08 | 1196800 | sample | 0.7868508 | -3.3131804 |
| P6 cl101 | H09 | 669940 | sample | 0.4404602 | -8.6974561 |
| P6 cl101 | H10 | 1396300 | sample | 0.9180145 | -1.2743788 |
| P6 cl101 | H11 | 1402400 | sample | 0.9220250 | -1.2120395 |
| P6 cl101 | H12 | 360760 | SAL | 0.2371861 | -11.8571388 |
| P7 cl101 | A01 | 332820 | SAL | 0.2359672 | -9.0124176 |
| P7 cl101 | A02 | 1363100 | sample | 0.9664292 | -0.3959967 |
| P7 cl101 | A03 | 1369300 | sample | 0.9708249 | -0.3441450 |
| P7 cl101 | A04 | 1311100 | sample | 0.9295615 | -0.8308823 |
| P7 cl101 | A05 | 1301500 | sample | 0.9227551 | -0.9111689 |
| P7 cl101 | A06 | 1366000 | sample | 0.9684852 | -0.3717435 |
| P7 cl101 | A07 | 1354200 | sample | 0.9601191 | -0.4704291 |
| P7 cl101 | A08 | 184020 | sample | 0.1304690 | -10.2568593 |
| P7 cl101 | A09 | 1207400 | sample | 0.8560389 | -1.6981444 |
| P7 cl101 | A10 | 1354100 | sample | 0.9600482 | -0.4712654 |
| P7 cl101 | A11 | 1373000 | sample | 0.9734482 | -0.3132012 |
| P7 cl101 | A12 | 1418900 | empty | 1.0059910 | 0.0706689 |
| P7 cl101 | B01 | 334260 | SAL | 0.2369882 | -9.0003746 |
| P7 cl101 | B02 | 912270 | sample | 0.6467936 | -4.1663708 |
| P7 cl101 | B03 | 1413400 | sample | 1.0020915 | 0.0246714 |
| P7 cl101 | B04 | 1475300 | sample | 1.0459782 | 0.5423525 |
| P7 cl101 | B05 | 1419400 | sample | 1.0063455 | 0.0748505 |
| P7 cl101 | B06 | 1251300 | sample | 0.8871637 | -1.3310007 |
| P7 cl101 | B07 | 1372100 | sample | 0.9728101 | -0.3207281 |
| P7 cl101 | B08 | 1345500 | sample | 0.9539509 | -0.5431888 |
| P7 cl101 | B09 | 1227300 | sample | 0.8701478 | -1.5317171 |
| P7 cl101 | B10 | 1401500 | sample | 0.9936545 | -0.0748505 |
| P7 cl101 | B11 | 1448200 | sample | 1.0267645 | 0.3157102 |
| P7 cl101 | B12 | 1701100 | empty | 1.2060690 | 2.4307593 |
| P7 cl101 | C01 | 355980 | SAL | 0.2523875 | -8.8187262 |
| P7 cl101 | C02 | 1431500 | sample | 1.0149243 | 0.1760450 |
| P7 cl101 | C03 | 1345000 | sample | 0.9535964 | -0.5473704 |
| P7 cl101 | C04 | 1445900 | sample | 1.0251338 | 0.2964749 |
| P7 cl101 | C05 | 1453500 | sample | 1.0305222 | 0.3600351 |
| P7 cl101 | C06 | 1491100 | sample | 1.0571803 | 0.6744908 |

*(continued)*

| Plate | Well | Value | Treatment | Norm | Score |
|---|---|---|---|---|---|
| P7 cl101 | C07 | 1493400 | sample | 1.0588110 | 0.6937261 |
| P7 cl101 | C08 | 1457700 | sample | 1.0334999 | 0.3951604 |
| P7 cl101 | C09 | 1451100 | sample | 1.0288206 | 0.3399634 |
| P7 cl101 | C10 | 1163400 | sample | 0.8248431 | -2.0661245 |
| P7 cl101 | C11 | 1514100 | sample | 1.0734872 | 0.8668440 |
| P7 cl101 | C12 | 1500100 | DMSO | 1.0635613 | 0.7497594 |
| P7 cl101 | D01 | 1276000 | DMSO | 0.9046758 | -1.1244300 |
| P7 cl101 | D02 | 1131000 | sample | 0.8018717 | -2.3370917 |
| P7 cl101 | D03 | 1371300 | sample | 0.9722429 | -0.3274186 |
| P7 cl101 | D04 | 1289900 | sample | 0.9145308 | -1.0081818 |
| P7 cl101 | D05 | 1434600 | sample | 1.0171222 | 0.2019709 |
| P7 cl101 | D06 | 1468500 | sample | 1.0411571 | 0.4854828 |
| P7 cl101 | D07 | 833160 | sample | 0.5907051 | -4.8279823 |
| P7 cl101 | D08 | 1429800 | sample | 1.0137190 | 0.1618276 |
| P7 cl101 | D09 | 1582600 | sample | 1.1220532 | 1.4397221 |
| P7 cl101 | D10 | 1515200 | sample | 1.0742671 | 0.8760435 |
| P7 cl101 | D11 | 1547000 | sample | 1.0968131 | 1.1419927 |
| P7 cl101 | D12 | 1530500 | DMSO | 1.0851147 | 1.0040002 |
| P7 cl101 | E01 | 1366600 | DMSO | 0.9689106 | -0.3667256 |
| P7 cl101 | E02 | 1491100 | sample | 1.0571803 | 0.6744908 |
| P7 cl101 | E03 | 232890 | sample | 0.1651175 | -9.8481505 |
| P7 cl101 | E04 | 1325500 | sample | 0.9397710 | -0.7104524 |
| P7 cl101 | E05 | 1543600 | sample | 1.0944025 | 1.1135579 |
| P7 cl101 | E06 | 1556800 | sample | 1.1037612 | 1.2239519 |
| P7 cl101 | E07 | 1587500 | sample | 1.1255273 | 1.4807017 |
| P7 cl101 | E08 | 1465800 | sample | 1.0392428 | 0.4629022 |
| P7 cl101 | E09 | 1519400 | sample | 1.0772449 | 0.9111689 |
| P7 cl101 | E10 | 1622100 | sample | 1.1500585 | 1.7700678 |
| P7 cl101 | E11 | 1577500 | sample | 1.1184374 | 1.3970698 |
| P7 cl101 | E12 | 1549700 | DMSO | 1.0987274 | 1.1645733 |
| P7 cl101 | F01 | 1355200 | DMSO | 0.9608281 | -0.4620659 |
| P7 cl101 | F02 | 1430900 | sample | 1.0144989 | 0.1710271 |
| P7 cl101 | F03 | 1322800 | sample | 0.9378567 | -0.7330330 |
| P7 cl101 | F04 | 1368800 | sample | 0.9704704 | -0.3483266 |
| P7 cl101 | F05 | 1500300 | sample | 1.0637031 | 0.7514320 |
| P7 cl101 | F06 | 1543200 | sample | 1.0941189 | 1.1102126 |
| P7 cl101 | F07 | 1515100 | sample | 1.0741962 | 0.8752072 |
| P7 cl101 | F08 | 1544900 | sample | 1.0953242 | 1.1244300 |
| P7 cl101 | F09 | 1516000 | sample | 1.0748343 | 0.8827340 |
| P7 cl101 | F10 | 1354200 | sample | 0.9601191 | -0.4704291 |
| P7 cl101 | F11 | 1522200 | sample | 1.0792300 | 0.9345858 |
| P7 cl101 | F12 | 398260 | SAL | 0.2823638 | -8.4651308 |
| P7 cl101 | G01 | 1206400 | empty | 0.8553299 | -1.7065076 |
| P7 cl101 | G02 | 220080 | sample | 0.1560353 | -9.9552829 |
| P7 cl101 | G03 | 1369600 | sample | 0.9710376 | -0.3416361 |
| P7 cl101 | G04 | 1368000 | sample | 0.9699032 | -0.3550171 |
| P7 cl101 | G05 | 1509300 | sample | 1.0700840 | 0.8267007 |
| P7 cl101 | G06 | 1500100 | sample | 1.0635613 | 0.7497594 |
| P7 cl101 | G07 | 1472800 | sample | 1.0442057 | 0.5214445 |
| P7 cl101 | G08 | 1449800 | sample | 1.0278989 | 0.3290913 |
| P7 cl101 | G09 | 1412200 | sample | 1.0012407 | 0.0146356 |
| P7 cl101 | G10 | 1471800 | sample | 1.0434968 | 0.5130813 |
| P7 cl101 | G11 | 1559500 | sample | 1.1056755 | 1.2465325 |
| P7 cl101 | G12 | 397900 | SAL | 0.2821085 | -8.4681416 |
| P7 cl101 | H01 | 1148800 | empty | 0.8144918 | -2.1882270 |
| P7 cl101 | H02 | 1401300 | sample | 0.9935127 | -0.0765231 |
| P7 cl101 | H03 | 1305800 | sample | 0.9258038 | -0.8752072 |
| P7 cl101 | H04 | 1365500 | sample | 0.9681307 | -0.3759251 |
| P7 cl101 | H05 | 1408700 | sample | 0.9987593 | -0.0146356 |
| P7 cl101 | H06 | 1333600 | sample | 0.9455138 | -0.6427107 |
| P7 cl101 | H07 | 1307900 | sample | 0.9272927 | -0.8576445 |
| P7 cl101 | H08 | 1333800 | sample | 0.9456556 | -0.6410380 |

*(continued)*

| Plate | Well | Value | Treatment | Norm | Score |
|---|---|---|---|---|---|
| P7 cl101 | H09 | 1330400 | sample | 0.9432451 | -0.6694728 |
| P7 cl101 | H10 | 1310700 | sample | 0.9292779 | -0.8342276 |
| P7 cl101 | H11 | 1141500 | sample | 0.8093162 | -2.2492782 |
| P7 cl101 | H12 | 344830 | SAL | 0.2444823 | -8.9119757 |
| P8 cl101 | A01 | 354000 | SAL | 0.2360000 | -12.8399736 |
| P8 cl101 | A02 | 1437200 | sample | 0.9581333 | -0.7036216 |
| P8 cl101 | A03 | 1091500 | sample | 0.7276667 | -4.5769016 |
| P8 cl101 | A04 | 1380400 | sample | 0.9202667 | -1.3400182 |
| P8 cl101 | A05 | 1397700 | sample | 0.9318000 | -1.1461861 |
| P8 cl101 | A06 | 1440800 | sample | 0.9605333 | -0.6632866 |
| P8 cl101 | A07 | 1437800 | sample | 0.9585333 | -0.6968991 |
| P8 cl101 | A08 | 1041000 | sample | 0.6940000 | -5.1427119 |
| P8 cl101 | A09 | 1359100 | sample | 0.9060667 | -1.5786669 |
| P8 cl101 | A10 | 1306000 | sample | 0.8706667 | -2.1736081 |
| P8 cl101 | A11 | 1495200 | sample | 0.9968000 | -0.0537800 |
| P8 cl101 | A12 | 1548400 | empty | 1.0322667 | 0.5422816 |
| P8 cl101 | B01 | 384310 | SAL | 0.2562067 | -12.5003753 |
| P8 cl101 | B02 | 1444600 | sample | 0.9630667 | -0.6207108 |
| P8 cl101 | B03 | 1631800 | sample | 1.0878667 | 1.4767090 |
| P8 cl101 | B04 | 1476100 | sample | 0.9840667 | -0.2677796 |
| P8 cl101 | B05 | 1562600 | sample | 1.0417333 | 0.7013808 |
| P8 cl101 | B06 | 1553600 | sample | 1.0357333 | 0.6005433 |
| P8 cl101 | B07 | 1418300 | sample | 0.9455333 | -0.9153803 |
| P8 cl101 | B08 | 1437800 | sample | 0.9585333 | -0.6968991 |
| P8 cl101 | B09 | 1560500 | sample | 1.0403333 | 0.6778520 |
| P8 cl101 | B10 | 1550100 | sample | 1.0334000 | 0.5613287 |
| P8 cl101 | B11 | 1512500 | sample | 1.0083333 | 0.1400521 |
| P8 cl101 | B12 | 1645000 | empty | 1.0966667 | 1.6246040 |
| P8 cl101 | C01 | 402130 | SAL | 0.2680867 | -12.3007171 |
| P8 cl101 | C02 | 1437200 | sample | 0.9581333 | -0.7036216 |
| P8 cl101 | C03 | 1474700 | sample | 0.9831333 | -0.2834654 |
| P8 cl101 | C04 | 1324600 | sample | 0.8830667 | -1.9652106 |
| P8 cl101 | C05 | 1499500 | sample | 0.9996667 | -0.0056021 |
| P8 cl101 | C06 | 1552200 | sample | 1.0348000 | 0.5848574 |
| P8 cl101 | C07 | 1354000 | sample | 0.9026667 | -1.6358082 |
| P8 cl101 | C08 | 1543200 | sample | 1.0288000 | 0.4840199 |
| P8 cl101 | C09 | 1519500 | sample | 1.0130000 | 0.2184812 |
| P8 cl101 | C10 | 1092500 | sample | 0.7283333 | -4.5656974 |
| P8 cl101 | C11 | 1720700 | sample | 1.1471333 | 2.4727593 |
| P8 cl101 | C12 | 1650700 | DMSO | 1.1004667 | 1.6884677 |
| P8 cl101 | D01 | 1376100 | DMSO | 0.9174000 | -1.3881961 |
| P8 cl101 | D02 | 1518400 | sample | 1.0122667 | 0.2061566 |
| P8 cl101 | D03 | 1489800 | sample | 0.9932000 | -0.1142825 |
| P8 cl101 | D04 | 1520500 | sample | 1.0136667 | 0.2296854 |
| P8 cl101 | D05 | 1524900 | sample | 1.0166000 | 0.2789837 |
| P8 cl101 | D06 | 1561000 | sample | 1.0406667 | 0.6834541 |
| P8 cl101 | D07 | 1547300 | sample | 1.0315333 | 0.5299570 |
| P8 cl101 | D08 | 1559900 | sample | 1.0399333 | 0.6711295 |
| P8 cl101 | D09 | 1646500 | sample | 1.0976667 | 1.6414102 |
| P8 cl101 | D10 | 1504500 | sample | 1.0030000 | 0.0504187 |
| P8 cl101 | D11 | 1599500 | sample | 1.0663333 | 1.1148145 |
| P8 cl101 | D12 | 1576200 | DMSO | 1.0508000 | 0.8537574 |
| P8 cl101 | E01 | 1431800 | DMSO | 0.9545333 | -0.7641241 |
| P8 cl101 | E02 | 1474300 | sample | 0.9828667 | -0.2879471 |
| P8 cl101 | E03 | 1540600 | sample | 1.0270667 | 0.4548891 |
| P8 cl101 | E04 | 1492900 | sample | 0.9952667 | -0.0795496 |
| P8 cl101 | E05 | 1541900 | sample | 1.0279333 | 0.4694545 |
| P8 cl101 | E06 | 1541700 | sample | 1.0278000 | 0.4672137 |
| P8 cl101 | E07 | 1634100 | sample | 1.0894000 | 1.5024786 |
| P8 cl101 | E08 | 1485000 | sample | 0.9900000 | -0.1680625 |
| P8 cl101 | E09 | 821350 | sample | 0.5475667 | -7.6037069 |
| P8 cl101 | E10 | 1567800 | sample | 1.0452000 | 0.7596424 |

*(continued)*

| Plate | Well | Value | Treatment | Norm | Score |
|---|---|---|---|---|---|
| P8 cl101 | E11 | 1568400 | sample | 1.0456000 | 0.7663649 |
| P8 cl101 | E12 | 1580400 | DMSO | 1.0536000 | 0.9008149 |
| P8 cl101 | F01 | 1373400 | DMSO | 0.9156000 | -1.4184473 |
| P8 cl101 | F02 | 1499200 | sample | 0.9994667 | -0.0089633 |
| P8 cl101 | F03 | 1516800 | sample | 1.0112000 | 0.1882300 |
| P8 cl101 | F04 | 1542800 | sample | 1.0285333 | 0.4795383 |
| P8 cl101 | F05 | 1527800 | sample | 1.0185333 | 0.3114758 |
| P8 cl101 | F06 | 1512700 | sample | 1.0084667 | 0.1422929 |
| P8 cl101 | F07 | 1542800 | sample | 1.0285333 | 0.4795383 |
| P8 cl101 | F08 | 1583800 | sample | 1.0558667 | 0.9389091 |
| P8 cl101 | F09 | 1576000 | sample | 1.0506667 | 0.8515166 |
| P8 cl101 | F10 | 1619500 | sample | 1.0796667 | 1.3388978 |
| P8 cl101 | F11 | 1605000 | sample | 1.0700000 | 1.1764374 |
| P8 cl101 | F12 | 450180 | SAL | 0.3001200 | -11.7623570 |
| P8 cl101 | G01 | 1383200 | empty | 0.9221333 | -1.3086465 |
| P8 cl101 | G02 | 1508700 | sample | 1.0058000 | 0.0974762 |
| P8 cl101 | G03 | 1504100 | sample | 1.0027333 | 0.0459371 |
| P8 cl101 | G04 | 1458000 | sample | 0.9720000 | -0.4705749 |
| P8 cl101 | G05 | 1500500 | sample | 1.0003333 | 0.0056021 |
| P8 cl101 | G06 | 434260 | sample | 0.2895067 | -11.9407273 |
| P8 cl101 | G07 | 1459700 | sample | 0.9731333 | -0.4515279 |
| P8 cl101 | G08 | 1526100 | sample | 1.0174000 | 0.2924287 |
| P8 cl101 | G09 | 1506400 | sample | 1.0042667 | 0.0717067 |
| P8 cl101 | G10 | 1502800 | sample | 1.0018667 | 0.0313717 |
| P8 cl101 | G11 | 1302600 | sample | 0.8684000 | -2.2117023 |
| P8 cl101 | G12 | 379960 | SAL | 0.2533067 | -12.5491135 |
| P8 cl101 | H01 | 1226200 | empty | 0.8174667 | -3.0677005 |
| P8 cl101 | H02 | 1388800 | sample | 0.9258667 | -1.2459032 |
| P8 cl101 | H03 | 1401400 | sample | 0.9342667 | -1.1047307 |
| P8 cl101 | H04 | 1328700 | sample | 0.8858000 | -1.9192735 |
| P8 cl101 | H05 | 1473300 | sample | 0.9822000 | -0.2991512 |
| P8 cl101 | H06 | 1417700 | sample | 0.9451333 | -0.9221028 |
| P8 cl101 | H07 | 1380200 | sample | 0.9201333 | -1.3422590 |
| P8 cl101 | H08 | 1320000 | sample | 0.8800000 | -2.0167498 |
| P8 cl101 | H09 | 1268200 | sample | 0.8454667 | -2.5971255 |
| P8 cl101 | H10 | 1353800 | sample | 0.9025333 | -1.6380490 |
| P8 cl101 | H11 | 125370 | sample | 0.0835800 | -15.4015819 |
| P8 cl101 | H12 | 328270 | SAL | 0.2188467 | -13.1282568 |
| P9 cl101 | A01 | 370210 | SAL | 0.2405992 | -14.0990288 |
| P9 cl101 | A02 | 1550400 | sample | 1.0076038 | 0.1411725 |
| P9 cl101 | A03 | 639850 | sample | 0.4158380 | -10.8455460 |
| P9 cl101 | A04 | 1495500 | sample | 0.9719244 | -0.5212523 |
| P9 cl101 | A05 | 1498600 | sample | 0.9739390 | -0.4838476 |
| P9 cl101 | A06 | 1654200 | sample | 1.0750634 | 1.3936258 |
| P9 cl101 | A07 | 1417500 | sample | 0.9212322 | -1.4624021 |
| P9 cl101 | A08 | 1396300 | sample | 0.9074543 | -1.7182019 |
| P9 cl101 | A09 | 1537400 | sample | 0.9991551 | -0.0156858 |
| P9 cl101 | A10 | 1490800 | sample | 0.9688698 | -0.5779626 |
| P9 cl101 | A11 | 1336600 | sample | 0.8686554 | -2.4385435 |
| P9 cl101 | A12 | 1483600 | empty | 0.9641906 | -0.6648379 |
| P9 cl101 | B01 | 382840 | SAL | 0.2488074 | -13.9466349 |
| P9 cl101 | B02 | 1626200 | sample | 1.0568662 | 1.0557771 |
| P9 cl101 | B03 | 1592900 | sample | 1.0352245 | 0.6539785 |
| P9 cl101 | B04 | 1597100 | sample | 1.0379541 | 0.7046558 |
| P9 cl101 | B05 | 1618900 | sample | 1.0521219 | 0.9676952 |
| P9 cl101 | B06 | 1597700 | sample | 1.0383441 | 0.7118954 |
| P9 cl101 | B07 | 1500300 | sample | 0.9750439 | -0.4633353 |
| P9 cl101 | B08 | 1381500 | sample | 0.8978358 | -1.8967790 |
| P9 cl101 | B09 | 1621600 | sample | 1.0538766 | 1.0002734 |
| P9 cl101 | B10 | 1662100 | sample | 1.0801976 | 1.4889474 |
| P9 cl101 | B11 | 1318500 | sample | 0.8568922 | -2.6569386 |

*(continued)*

| Plate | Well | Value | Treatment | Norm | Score |
|---|---|---|---|---|---|
| P9 cl101 | B12 | 1673700 | empty | 1.0877364 | 1.6289133 |
| P9 cl101 | C01 | 416130 | SAL | 0.2704426 | -13.5449569 |
| P9 cl101 | C02 | 1568400 | sample | 1.0193020 | 0.3583609 |
| P9 cl101 | C03 | 1580000 | sample | 1.0268408 | 0.4983268 |
| P9 cl101 | C04 | 1455900 | sample | 0.9461883 | -0.9990668 |
| P9 cl101 | C05 | 1511700 | sample | 0.9824527 | -0.3257827 |
| P9 cl101 | C06 | 1520800 | sample | 0.9883668 | -0.2159818 |
| P9 cl101 | C07 | 1052000 | sample | 0.6836940 | -5.8725340 |
| P9 cl101 | C08 | 1543100 | sample | 1.0028596 | 0.0530905 |
| P9 cl101 | C09 | 1492900 | sample | 0.9702346 | -0.5526239 |
| P9 cl101 | C10 | 1667500 | sample | 1.0837070 | 1.5541039 |
| P9 cl101 | C11 | 1785900 | sample | 1.1606551 | 2.9827212 |
| P9 cl101 | C12 | 1653600 | DMSO | 1.0746734 | 1.3863862 |
| P9 cl101 | D01 | 1472500 | DMSO | 0.9569767 | -0.7987708 |
| P9 cl101 | D02 | 1480700 | sample | 0.9623058 | -0.6998294 |
| P9 cl101 | D03 | 1468100 | sample | 0.9541171 | -0.8518613 |
| P9 cl101 | D04 | 1125900 | sample | 0.7317216 | -4.9808548 |
| P9 cl101 | D05 | 1495100 | sample | 0.9716644 | -0.5260787 |
| P9 cl101 | D06 | 1532900 | sample | 0.9962306 | -0.0699829 |
| P9 cl101 | D07 | 1592500 | sample | 1.0349646 | 0.6491521 |
| P9 cl101 | D08 | 1573900 | sample | 1.0228765 | 0.4247241 |
| P9 cl101 | D09 | 1619200 | sample | 1.0523169 | 0.9713150 |
| P9 cl101 | D10 | 1581900 | sample | 1.0280756 | 0.5212523 |
| P9 cl101 | D11 | 1723800 | sample | 1.1202964 | 2.2334211 |
| P9 cl101 | D12 | 1578900 | DMSO | 1.0261260 | 0.4850542 |
| P9 cl101 | E01 | 1489000 | DMSO | 0.9677000 | -0.5996814 |
| P9 cl101 | E02 | 1635300 | sample | 1.0627803 | 1.1655779 |
| P9 cl101 | E03 | 1542100 | sample | 1.0022097 | 0.0410245 |
| P9 cl101 | E04 | 1547800 | sample | 1.0059141 | 0.1098008 |
| P9 cl101 | E05 | 1548900 | sample | 1.0066290 | 0.1230734 |
| P9 cl101 | E06 | 1570400 | sample | 1.0206018 | 0.3824930 |
| P9 cl101 | E07 | 1583900 | sample | 1.0293754 | 0.5453843 |
| P9 cl101 | E08 | 1501400 | sample | 0.9757588 | -0.4500627 |
| P9 cl101 | E09 | 1627700 | sample | 1.0578410 | 1.0738762 |
| P9 cl101 | E10 | 1678200 | sample | 1.0906609 | 1.6832104 |
| P9 cl101 | E11 | 1572700 | sample | 1.0220966 | 0.4102448 |
| P9 cl101 | E12 | 1691000 | DMSO | 1.0989797 | 1.8376555 |
| P9 cl101 | F01 | 1505700 | DMSO | 0.9785533 | -0.3981788 |
| P9 cl101 | F02 | 1550400 | sample | 1.0076038 | 0.1411725 |
| P9 cl101 | F03 | 1532000 | sample | 0.9956457 | -0.0808424 |
| P9 cl101 | F04 | 1411600 | sample | 0.9173978 | -1.5335917 |
| P9 cl101 | F05 | 1563600 | sample | 1.0161825 | 0.3004440 |
| P9 cl101 | F06 | 1581700 | sample | 1.0279457 | 0.5188390 |
| P9 cl101 | F07 | 1588900 | sample | 1.0326249 | 0.6057144 |
| P9 cl101 | F08 | 1562000 | sample | 1.0151427 | 0.2811384 |
| P9 cl101 | F09 | 1582300 | sample | 1.0283356 | 0.5260787 |
| P9 cl101 | F10 | 1594100 | sample | 1.0360044 | 0.6684577 |
| P9 cl101 | F11 | 1649800 | sample | 1.0722038 | 1.3405353 |
| P9 cl101 | F12 | 439940 | SAL | 0.2859167 | -13.2576649 |
| P9 cl101 | G01 | 1429300 | empty | 0.9289010 | -1.3200231 |
| P9 cl101 | G02 | 1466700 | sample | 0.9532073 | -0.8687538 |
| P9 cl101 | G03 | 1578100 | sample | 1.0256060 | 0.4754014 |
| P9 cl101 | G04 | 1540000 | sample | 1.0008449 | 0.0156858 |
| P9 cl101 | G05 | 1527000 | sample | 0.9923962 | -0.1411725 |
| P9 cl101 | G06 | 932140 | sample | 0.6057971 | -7.3187677 |
| P9 cl101 | G07 | 1359400 | sample | 0.8834731 | -2.1634382 |
| P9 cl101 | G08 | 1392800 | sample | 0.9051797 | -1.7604329 |
| P9 cl101 | G09 | 919700 | sample | 0.5977124 | -7.4688691 |
| P9 cl101 | G10 | 1595100 | sample | 1.0366543 | 0.6805238 |
| P9 cl101 | G11 | 1578200 | sample | 1.0256710 | 0.4766080 |
| P9 cl101 | G12 | 425820 | SAL | 0.2767401 | -13.4280371 |
| P9 cl101 | H01 | 1335100 | empty | 0.8676805 | -2.4566426 |

*(continued)*

| Plate | Well | Value | Treatment | Norm | Score |
|---|---|---|---|---|---|
| P9 cl101 | H02 | 1470900 | sample | 0.9559368 | -0.8180764 |
| P9 cl101 | H03 | 1508300 | sample | 0.9802431 | -0.3668071 |
| P9 cl101 | H04 | 1440200 | sample | 0.9359849 | -1.1885034 |
| P9 cl101 | H05 | 1466800 | sample | 0.9532722 | -0.8675471 |
| P9 cl101 | H06 | 1456800 | sample | 0.9467733 | -0.9882074 |
| P9 cl101 | H07 | 1177500 | sample | 0.7652564 | -4.3582480 |
| P9 cl101 | H08 | 1386700 | sample | 0.9012153 | -1.8340357 |
| P9 cl101 | H09 | 1454700 | sample | 0.9454085 | -1.0135460 |
| P9 cl101 | H10 | 1490200 | sample | 0.9684799 | -0.5852022 |
| P9 cl101 | H11 | 1483900 | sample | 0.9643855 | -0.6612181 |
| P9 cl101 | H12 | 408670 | SAL | 0.2655943 | -13.6349695 |
| P10 cl101 | A01 | 330830 | SAL | 0.2255300 | -9.3504419 |
| P10 cl101 | A02 | 1535300 | sample | 1.0466289 | 0.5629673 |
| P10 cl101 | A03 | 1466800 | sample | 0.9999318 | -0.0008231 |
| P10 cl101 | A04 | 1371400 | sample | 0.9348967 | -0.7860142 |
| P10 cl101 | A05 | 1383400 | sample | 0.9430772 | -0.6872481 |
| P10 cl101 | A06 | 555440 | sample | 0.3786489 | -7.5017858 |
| P10 cl101 | A07 | 1394400 | sample | 0.9505760 | -0.5967124 |
| P10 cl101 | A08 | 1240000 | sample | 0.8453201 | -1.8675040 |
| P10 cl101 | A09 | 1290600 | sample | 0.8798146 | -1.4510399 |
| P10 cl101 | A10 | 1350300 | sample | 0.9205126 | -0.9596781 |
| P10 cl101 | A11 | 1393100 | sample | 0.9496898 | -0.6074121 |
| P10 cl101 | A12 | 1341300 | empty | 0.9143773 | -1.0337528 |
| P10 cl101 | B01 | 361150 | SAL | 0.2461995 | -9.1008927 |
| P10 cl101 | B02 | 1614700 | sample | 1.1007567 | 1.2164702 |
| P10 cl101 | B03 | 1455500 | sample | 0.9922285 | -0.0938279 |
| P10 cl101 | B04 | 1511900 | sample | 1.0306769 | 0.3703732 |
| P10 cl101 | B05 | 1579300 | sample | 1.0766242 | 0.9251100 |
| P10 cl101 | B06 | 1568700 | sample | 1.0693981 | 0.8378665 |
| P10 cl101 | B07 | 1448700 | sample | 0.9875929 | -0.1497954 |
| P10 cl101 | B08 | 1434300 | sample | 0.9777763 | -0.2683148 |
| P10 cl101 | B09 | 1494700 | sample | 1.0189515 | 0.2288083 |
| P10 cl101 | B10 | 1498900 | sample | 1.0218147 | 0.2633765 |
| P10 cl101 | B11 | 1554500 | sample | 1.0597178 | 0.7209932 |
| P10 cl101 | B12 | 1622800 | empty | 1.1062785 | 1.2831374 |
| P10 cl101 | C01 | 360100 | SAL | 0.2454837 | -9.1095347 |
| P10 cl101 | C02 | 1430200 | sample | 0.9749813 | -0.3020599 |
| P10 cl101 | C03 | 1466400 | sample | 0.9996591 | -0.0041153 |
| P10 cl101 | C04 | 1396900 | sample | 0.9522803 | -0.5761361 |
| P10 cl101 | C05 | 1423500 | sample | 0.9704138 | -0.3572044 |
| P10 cl101 | C06 | 1510800 | sample | 1.0299271 | 0.3613196 |
| P10 cl101 | C07 | 1529300 | sample | 1.0425387 | 0.5135842 |
| P10 cl101 | C08 | 1531800 | sample | 1.0442430 | 0.5341605 |
| P10 cl101 | C09 | 1147300 | sample | 0.7821256 | -2.6304728 |
| P10 cl101 | C10 | 1547100 | sample | 1.0546731 | 0.6600874 |
| P10 cl101 | C11 | 467990 | sample | 0.3190333 | -8.2215444 |
| P10 cl101 | C12 | 1536500 | DMSO | 1.0474470 | 0.5728439 |
| P10 cl101 | D01 | 1575200 | DMSO | 1.0738292 | 0.8913648 |
| P10 cl101 | D02 | 1594500 | sample | 1.0869862 | 1.0502138 |
| P10 cl101 | D03 | 1572400 | sample | 1.0719204 | 0.8683194 |
| P10 cl101 | D04 | 1443700 | sample | 0.9841843 | -0.1909480 |
| P10 cl101 | D05 | 1437300 | sample | 0.9798214 | -0.2436233 |
| P10 cl101 | D06 | 1562000 | sample | 1.0648306 | 0.7827220 |
| P10 cl101 | D07 | 1322500 | sample | 0.9015611 | -1.1884865 |
| P10 cl101 | D08 | 1491400 | sample | 1.0167019 | 0.2016476 |
| P10 cl101 | D09 | 1567600 | sample | 1.0686482 | 0.8288129 |
| P10 cl101 | D10 | 1624700 | sample | 1.1075738 | 1.2987754 |
| P10 cl101 | D11 | 1535400 | sample | 1.0466971 | 0.5637903 |
| P10 cl101 | D12 | 1351700 | DMSO | 0.9214670 | -0.9481554 |
| P10 cl101 | E01 | 1558100 | DMSO | 1.0621719 | 0.7506230 |
| P10 cl101 | E02 | 1502900 | sample | 1.0245416 | 0.2962986 |
| P10 cl101 | E03 | 1605600 | sample | 1.0945531 | 1.1415725 |

*(continued)*

| Plate | Well | Value | Treatment | Norm | Score |
|---|---|---|---|---|---|
| P10 cl101 | E04 | 1463900 | sample | 0.9979549 | -0.0246915 |
| P10 cl101 | E05 | 1514700 | sample | 1.0325857 | 0.3934186 |
| P10 cl101 | E06 | 1554600 | sample | 1.0597859 | 0.7218162 |
| P10 cl101 | E07 | 1580100 | sample | 1.0771695 | 0.9316944 |
| P10 cl101 | E08 | 1427800 | sample | 0.9733451 | -0.3218132 |
| P10 cl101 | E09 | 1612900 | sample | 1.0995296 | 1.2016553 |
| P10 cl101 | E10 | 1576200 | sample | 1.0745109 | 0.8995954 |
| P10 cl101 | E11 | 1565800 | sample | 1.0674211 | 0.8139980 |
| P10 cl101 | E12 | 1701400 | DMSO | 1.1598609 | 1.9300559 |
| P10 cl101 | F01 | 1441900 | DMSO | 0.9829573 | -0.2057629 |
| P10 cl101 | F02 | 1569400 | sample | 1.0698752 | 0.8436279 |
| P10 cl101 | F03 | 1381800 | sample | 0.9419865 | -0.7004169 |
| P10 cl101 | F04 | 1484400 | sample | 1.0119299 | 0.1440340 |
| P10 cl101 | F05 | 1606400 | sample | 1.0950985 | 1.1481569 |
| P10 cl101 | F06 | 1552900 | sample | 1.0586270 | 0.7078243 |
| P10 cl101 | F07 | 1467800 | sample | 1.0006135 | 0.0074075 |
| P10 cl101 | F08 | 287330 | sample | 0.1958757 | -9.7084694 |
| P10 cl101 | F09 | 1351300 | sample | 0.9211944 | -0.9514476 |
| P10 cl101 | F10 | 1519200 | sample | 1.0356534 | 0.4304560 |
| P10 cl101 | F11 | 1529400 | sample | 1.0426069 | 0.5144072 |
| P10 cl101 | F12 | 385490 | SAL | 0.2627923 | -8.9005620 |
| P10 cl101 | G01 | 1449100 | empty | 0.9878656 | -0.1465032 |
| P10 cl101 | G02 | 1576200 | sample | 1.0745109 | 0.8995954 |
| P10 cl101 | G03 | 451280 | sample | 0.3076420 | -8.3590763 |
| P10 cl101 | G04 | 1456100 | sample | 0.9926375 | -0.0888896 |
| P10 cl101 | G05 | 1494500 | sample | 1.0188152 | 0.2271622 |
| P10 cl101 | G06 | 1496600 | sample | 1.0202468 | 0.2444463 |
| P10 cl101 | G07 | 1388400 | sample | 0.9464858 | -0.6460955 |
| P10 cl101 | G08 | 1242100 | sample | 0.8467517 | -1.8502199 |
| P10 cl101 | G09 | 1359300 | sample | 0.9266480 | -0.8856035 |
| P10 cl101 | G10 | 1200300 | sample | 0.8182562 | -2.1942555 |
| P10 cl101 | G11 | 241400 | sample | 0.1645647 | -10.0864970 |
| P10 cl101 | G12 | 364900 | SAL | 0.2487559 | -9.0700283 |
| P10 cl101 | H01 | 1317000 | empty | 0.8978117 | -1.2337543 |
| P10 cl101 | H02 | 1493000 | sample | 1.0177926 | 0.2148165 |
| P10 cl101 | H03 | 1386500 | sample | 0.9451905 | -0.6617335 |
| P10 cl101 | H04 | 1182300 | sample | 0.8059854 | -2.3424048 |
| P10 cl101 | H05 | 1497300 | sample | 1.0207240 | 0.2502077 |
| P10 cl101 | H06 | 1467000 | sample | 1.0000682 | 0.0008231 |
| P10 cl101 | H07 | 1366400 | sample | 0.9314882 | -0.8271668 |
| P10 cl101 | H08 | 1191900 | sample | 0.8125298 | -2.2633918 |
| P10 cl101 | H09 | 1325800 | sample | 0.9038108 | -1.1613258 |
| P10 cl101 | H10 | 1392200 | sample | 0.9490763 | -0.6148195 |
| P10 cl101 | H11 | 1408700 | sample | 0.9603245 | -0.4790160 |
| P10 cl101 | H12 | 344820 | SAL | 0.2350671 | -9.2352970 |
| P11 cl101 | A01 | 334680 | SAL | 0.2160341 | -14.1238365 |
| P11 cl101 | A02 | 1440700 | sample | 0.9299639 | -1.2617629 |
| P11 cl101 | A03 | 1535700 | sample | 0.9912858 | -0.1569935 |
| P11 cl101 | A04 | 1486600 | sample | 0.9595920 | -0.7279849 |
| P11 cl101 | A05 | 1492400 | sample | 0.9633359 | -0.6605358 |
| P11 cl101 | A06 | 1465900 | sample | 0.9462303 | -0.9687083 |
| P11 cl101 | A07 | 1230800 | sample | 0.7944746 | -3.7027217 |
| P11 cl101 | A08 | 1226400 | sample | 0.7916344 | -3.7538900 |
| P11 cl101 | A09 | 1279800 | sample | 0.8261038 | -3.1328933 |
| P11 cl101 | A10 | 1515500 | sample | 0.9782468 | -0.3919024 |
| P11 cl101 | A11 | 1432500 | sample | 0.9246708 | -1.3571219 |
| P11 cl101 | A12 | 1487500 | empty | 0.9601730 | -0.7175186 |
| P11 cl101 | B01 | 387340 | SAL | 0.2500258 | -13.5114454 |
| P11 cl101 | B02 | 67293 | sample | 0.0434373 | -17.2333203 |
| P11 cl101 | B03 | 1552400 | sample | 1.0020656 | 0.0372133 |
| P11 cl101 | B04 | 1600200 | sample | 1.0329202 | 0.5930867 |

*(continued)*

| Plate | Well | Value | Treatment | Norm | Score |
|---|---|---|---|---|---|
| P11 cl101 | B05 | 1706200 | sample | 1.1013426 | 1.8257767 |
| P11 cl101 | B06 | 1634900 | sample | 1.0553189 | 0.9966182 |
| P11 cl101 | B07 | 1514500 | sample | 0.9776013 | -0.4035315 |
| P11 cl101 | B08 | 1570500 | sample | 1.0137490 | 0.2477009 |
| P11 cl101 | B09 | 1538500 | sample | 0.9930932 | -0.1244319 |
| P11 cl101 | B10 | 1640700 | sample | 1.0590627 | 1.0640673 |
| P11 cl101 | B11 | 1587700 | sample | 1.0248515 | 0.4477223 |
| P11 cl101 | B12 | 1651700 | empty | 1.0661632 | 1.1919880 |
| P11 cl101 | C01 | 409510 | SAL | 0.2643364 | -13.2536271 |
| P11 cl101 | C02 | 1526200 | sample | 0.9851536 | -0.2674705 |
| P11 cl101 | C03 | 1659900 | sample | 1.0714562 | 1.2873470 |
| P11 cl101 | C04 | 1477600 | sample | 0.9537826 | -0.8326472 |
| P11 cl101 | C05 | 1635100 | sample | 1.0554480 | 0.9989441 |
| P11 cl101 | C06 | 1707500 | sample | 1.1021818 | 1.8408946 |
| P11 cl101 | C07 | 1682000 | sample | 1.0857217 | 1.5443513 |
| P11 cl101 | C08 | 1555000 | sample | 1.0037439 | 0.0674491 |
| P11 cl101 | C09 | 1595800 | sample | 1.0300800 | 0.5419184 |
| P11 cl101 | C10 | 1592300 | sample | 1.0278208 | 0.5012164 |
| P11 cl101 | C11 | 1302600 | sample | 0.8408211 | -2.8677486 |
| P11 cl101 | C12 | 1365900 | DMSO | 0.8816809 | -2.1316234 |
| P11 cl101 | D01 | 1561800 | DMSO | 1.0081332 | 0.1465273 |
| P11 cl101 | D02 | 1575900 | sample | 1.0172347 | 0.3104983 |
| P11 cl101 | D03 | 1527000 | sample | 0.9856700 | -0.2581672 |
| P11 cl101 | D04 | 1548800 | sample | 0.9997418 | -0.0046517 |
| P11 cl101 | D05 | 1574400 | sample | 1.0162665 | 0.2930546 |
| P11 cl101 | D06 | 1603100 | sample | 1.0347922 | 0.6268112 |
| P11 cl101 | D07 | 1516000 | sample | 0.9785696 | -0.3860878 |
| P11 cl101 | D08 | 1638800 | sample | 1.0578363 | 1.0419719 |
| P11 cl101 | D09 | 1631700 | sample | 1.0532533 | 0.9594050 |
| P11 cl101 | D10 | 113620 | sample | 0.0733411 | -16.6945766 |
| P11 cl101 | D11 | 1533300 | sample | 0.9897366 | -0.1849035 |
| P11 cl101 | D12 | 1546000 | DMSO | 0.9979344 | -0.0372133 |
| P11 cl101 | E01 | 1607700 | DMSO | 1.0377614 | 0.6803053 |
| P11 cl101 | E02 | 1590700 | sample | 1.0267880 | 0.4826098 |
| P11 cl101 | E03 | 1475400 | sample | 0.9523625 | -0.8582313 |
| P11 cl101 | E04 | 1677200 | sample | 1.0826233 | 1.4885313 |
| P11 cl101 | E05 | 1577800 | sample | 1.0184611 | 0.3325937 |
| P11 cl101 | E06 | 1611100 | sample | 1.0399561 | 0.7198444 |
| P11 cl101 | E07 | 1621600 | sample | 1.0467338 | 0.8419505 |
| P11 cl101 | E08 | 1548700 | sample | 0.9996773 | -0.0058146 |
| P11 cl101 | E09 | 1614600 | sample | 1.0422153 | 0.7605465 |
| P11 cl101 | E10 | 1641900 | sample | 1.0598373 | 1.0780223 |
| P11 cl101 | E11 | 1623700 | sample | 1.0480893 | 0.8663718 |
| P11 cl101 | E12 | 1656300 | DMSO | 1.0691325 | 1.2454821 |
| P11 cl101 | F01 | 1527600 | DMSO | 0.9860573 | -0.2511897 |
| P11 cl101 | F02 | 1504600 | sample | 0.9712109 | -0.5186601 |
| P11 cl101 | F03 | 1544500 | sample | 0.9969662 | -0.0546570 |
| P11 cl101 | F04 | 1566000 | sample | 1.0108443 | 0.1953697 |
| P11 cl101 | F05 | 1558000 | sample | 1.0056804 | 0.1023365 |
| P11 cl101 | F06 | 1651100 | sample | 1.0657759 | 1.1850105 |
| P11 cl101 | F07 | 1641400 | sample | 1.0595146 | 1.0722077 |
| P11 cl101 | F08 | 1549600 | sample | 1.0002582 | 0.0046517 |
| P11 cl101 | F09 | 1299700 | sample | 0.8389491 | -2.9014732 |
| P11 cl101 | F10 | 348810 | sample | 0.2251549 | -13.9595166 |
| P11 cl101 | F11 | 1590300 | sample | 1.0265298 | 0.4779581 |
| P11 cl101 | F12 | 464750 | SAL | 0.2999935 | -12.6112328 |
| P11 cl101 | G01 | 1561100 | empty | 1.0076814 | 0.1383869 |
| P11 cl101 | G02 | 1582900 | sample | 1.0217532 | 0.3919024 |
| P11 cl101 | G03 | 572910 | sample | 0.3698102 | -11.3534239 |
| P11 cl101 | G04 | 1209600 | sample | 0.7807901 | -3.9492597 |
| P11 cl101 | G05 | 1608400 | sample | 1.0382133 | 0.6884457 |
| P11 cl101 | G06 | 717280 | sample | 0.4630003 | -9.6745233 |

*(continued)*

| Plate | Well | Value | Treatment | Norm | Score |
|---|---|---|---|---|---|
| P11 cl101 | G07 | 1380700 | sample | 0.8912342 | -1.9595119 |
| P11 cl101 | G08 | 1575700 | sample | 1.0171056 | 0.3081725 |
| P11 cl101 | G09 | 1586500 | sample | 1.0240769 | 0.4337673 |
| P11 cl101 | G10 | 1561700 | sample | 1.0080687 | 0.1453644 |
| P11 cl101 | G11 | 1333900 | sample | 0.8610250 | -2.5037562 |
| P11 cl101 | G12 | 422050 | SAL | 0.2724309 | -13.1077976 |
| P11 cl101 | H01 | 1135600 | empty | 0.7330235 | -4.8098169 |
| P11 cl101 | H02 | 1506700 | sample | 0.9725665 | -0.4942389 |
| P11 cl101 | H03 | 1502700 | sample | 0.9699845 | -0.5407555 |
| P11 cl101 | H04 | 1529100 | sample | 0.9870256 | -0.2337459 |
| P11 cl101 | H05 | 1596800 | sample | 1.0307255 | 0.5535476 |
| P11 cl101 | H06 | 1467700 | sample | 0.9473922 | -0.9477758 |
| P11 cl101 | H07 | 1504900 | sample | 0.9714046 | -0.5151714 |
| P11 cl101 | H08 | 1512800 | sample | 0.9765040 | -0.4233011 |
| P11 cl101 | H09 | 1467400 | sample | 0.9471986 | -0.9512646 |
| P11 cl101 | H10 | 1554600 | sample | 1.0034857 | 0.0627974 |
| P11 cl101 | H11 | 1481900 | sample | 0.9565582 | -0.7826419 |
| P11 cl101 | H12 | 401040 | SAL | 0.2588691 | -13.3521260 |
| P12 cl101 | A01 | 324250 | SAL | 0.2148062 | -12.8321055 |
| P12 cl101 | A02 | 1509000 | sample | 0.9996688 | -0.0054132 |
| P12 cl101 | A03 | 1471700 | sample | 0.9749586 | -0.4092416 |
| P12 cl101 | A04 | 1404400 | sample | 0.9303743 | -1.1378648 |
| P12 cl101 | A05 | 1306400 | sample | 0.8654521 | -2.1988615 |
| P12 cl101 | A06 | 1386400 | sample | 0.9184498 | -1.3327418 |
| P12 cl101 | A07 | 1169800 | sample | 0.7749586 | -3.6777610 |
| P12 cl101 | A08 | 1321200 | sample | 0.8752567 | -2.0386294 |
| P12 cl101 | A09 | 1310300 | sample | 0.8680358 | -2.1566382 |
| P12 cl101 | A10 | 1362500 | sample | 0.9026168 | -1.5914951 |
| P12 cl101 | A11 | 1476200 | sample | 0.9779397 | -0.3605223 |
| P12 cl101 | A12 | 1536400 | empty | 1.0178205 | 0.2912328 |
| P12 cl101 | B01 | 385450 | SAL | 0.2553495 | -12.1695239 |
| P12 cl101 | B02 | 1591700 | sample | 1.0544551 | 0.8899380 |
| P12 cl101 | B03 | 731570 | sample | 0.4846439 | -8.4222568 |
| P12 cl101 | B04 | 1512100 | sample | 1.0017224 | 0.0281489 |
| P12 cl101 | B05 | 1543100 | sample | 1.0222590 | 0.3637703 |
| P12 cl101 | B06 | 1529000 | sample | 1.0129182 | 0.2111167 |
| P12 cl101 | B07 | 1488000 | sample | 0.9857569 | -0.2327697 |
| P12 cl101 | B08 | 1519400 | sample | 1.0065585 | 0.1071823 |
| P12 cl101 | B09 | 1544700 | sample | 1.0233190 | 0.3810927 |
| P12 cl101 | B10 | 1428800 | sample | 0.9465386 | -0.8736983 |
| P12 cl101 | B11 | 1571000 | sample | 1.0407420 | 0.6658296 |
| P12 cl101 | B12 | 1664800 | empty | 1.1028817 | 1.6813550 |
| P12 cl101 | C01 | 414250 | SAL | 0.2744286 | -11.8577208 |
| P12 cl101 | C02 | 1492300 | sample | 0.9886055 | -0.1862157 |
| P12 cl101 | C03 | 1454500 | sample | 0.9635641 | -0.5954573 |
| P12 cl101 | C04 | 1500900 | sample | 0.9943027 | -0.0931079 |
| P12 cl101 | C05 | 1543400 | sample | 1.0224578 | 0.3670182 |
| P12 cl101 | C06 | 1572600 | sample | 1.0418019 | 0.6831520 |
| P12 cl101 | C07 | 1381200 | sample | 0.9150050 | -1.3890396 |
| P12 cl101 | C08 | 1532800 | sample | 1.0154356 | 0.2522574 |
| P12 cl101 | C09 | 1574300 | sample | 1.0429281 | 0.7015570 |
| P12 cl101 | C10 | 1598800 | sample | 1.0591587 | 0.9668062 |
| P12 cl101 | C11 | 1525400 | sample | 1.0105333 | 0.1721413 |
| P12 cl101 | C12 | 1544900 | DMSO | 1.0234515 | 0.3832580 |
| P12 cl101 | D01 | 1517800 | DMSO | 1.0054985 | 0.0898599 |
| P12 cl101 | D02 | 1545600 | sample | 1.0239152 | 0.3908365 |
| P12 cl101 | D03 | 857440 | sample | 0.5680291 | -7.0595256 |
| P12 cl101 | D04 | 1461700 | sample | 0.9683339 | -0.5175066 |
| P12 cl101 | D05 | 1499800 | sample | 0.9935740 | -0.1050170 |
| P12 cl101 | D06 | 1517200 | sample | 1.0051010 | 0.0833640 |
| P12 cl101 | D07 | 1575600 | sample | 1.0437893 | 0.7156314 |
| P12 cl101 | D08 | 1564900 | sample | 1.0367009 | 0.5997879 |

*(continued)*

| Plate | Well | Value | Treatment | Norm | Score |
|---|---|---|---|---|---|
| P12 cl101 | D09 | 1593000 | sample | 1.0553163 | 0.9040125 |
| P12 cl101 | D10 | 1588300 | sample | 1.0522027 | 0.8531280 |
| P12 cl101 | D11 | 1002100 | sample | 0.6638622 | -5.4933645 |
| P12 cl101 | D12 | 1677300 | DMSO | 1.1111626 | 1.8166862 |
| P12 cl101 | E01 | 1620800 | DMSO | 1.0737330 | 1.2049891 |
| P12 cl101 | E02 | 1583100 | sample | 1.0487579 | 0.7968302 |
| P12 cl101 | E03 | 1524100 | sample | 1.0096721 | 0.1580669 |
| P12 cl101 | E04 | 1544100 | sample | 1.0229215 | 0.3745968 |
| P12 cl101 | E05 | 638410 | sample | 0.4229281 | -9.4308532 |
| P12 cl101 | E06 | 598400 | sample | 0.3964227 | -9.8640214 |
| P12 cl101 | E07 | 1555900 | sample | 1.0307387 | 0.5023495 |
| P12 cl101 | E08 | 1552100 | sample | 1.0282213 | 0.4612088 |
| P12 cl101 | E09 | 1628200 | sample | 1.0786353 | 1.2851052 |
| P12 cl101 | E10 | 1598100 | sample | 1.0586949 | 0.9592276 |
| P12 cl101 | E11 | 1391500 | sample | 0.9218284 | -1.2775266 |
| P12 cl101 | E12 | 1660200 | DMSO | 1.0998344 | 1.6315531 |
| P12 cl101 | F01 | 1623800 | DMSO | 1.0757204 | 1.2374686 |
| P12 cl101 | F02 | 1566100 | sample | 1.0374959 | 0.6127797 |
| P12 cl101 | F03 | 1053600 | sample | 0.6979795 | -4.9358000 |
| P12 cl101 | F04 | 1468600 | sample | 0.9729049 | -0.4428037 |
| P12 cl101 | F05 | 1577900 | sample | 1.0453130 | 0.7405324 |
| P12 cl101 | F06 | 1537900 | sample | 1.0188142 | 0.3074725 |
| P12 cl101 | F07 | 1525800 | sample | 1.0107983 | 0.1764719 |
| P12 cl101 | F08 | 1551000 | sample | 1.0274925 | 0.4492996 |
| P12 cl101 | F09 | 1540000 | sample | 1.0202054 | 0.3302082 |
| P12 cl101 | F10 | 1596000 | sample | 1.0573037 | 0.9364920 |
| P12 cl101 | F11 | 1595500 | sample | 1.0569725 | 0.9310787 |
| P12 cl101 | F12 | 415730 | SAL | 0.2754091 | -11.8416976 |
| P12 cl101 | G01 | 1459000 | empty | 0.9665452 | -0.5467381 |
| P12 cl101 | G02 | 1508900 | sample | 0.9996025 | -0.0064959 |
| P12 cl101 | G03 | 1492200 | sample | 0.9885393 | -0.1872984 |
| P12 cl101 | G04 | 1526600 | sample | 1.0113283 | 0.1851331 |
| P12 cl101 | G05 | 1510000 | sample | 1.0003312 | 0.0054132 |
| P12 cl101 | G06 | 1205200 | sample | 0.7984101 | -3.2945030 |
| P12 cl101 | G07 | 1473900 | sample | 0.9764160 | -0.3854233 |
| P12 cl101 | G08 | 1289500 | sample | 0.8542564 | -2.3818293 |
| P12 cl101 | G09 | 1575000 | sample | 1.0433919 | 0.7091355 |
| P12 cl101 | G10 | 1517200 | sample | 1.0051010 | 0.0833640 |
| P12 cl101 | G11 | 1519400 | sample | 1.0065585 | 0.1071823 |
| P12 cl101 | G12 | 394030 | SAL | 0.2610335 | -12.0766325 |
| P12 cl101 | H01 | 1365100 | empty | 0.9043392 | -1.5633462 |
| P12 cl101 | H02 | 1422900 | sample | 0.9426300 | -0.9375746 |
| P12 cl101 | H03 | 1406000 | sample | 0.9314342 | -1.1205424 |
| P12 cl101 | H04 | 1495500 | sample | 0.9907254 | -0.1515710 |
| P12 cl101 | H05 | 1351600 | sample | 0.8953958 | -1.7095039 |
| P12 cl101 | H06 | 1322400 | sample | 0.8760517 | -2.0256376 |
| P12 cl101 | H07 | 1288100 | sample | 0.8533289 | -2.3969864 |
| P12 cl101 | H08 | 1291500 | sample | 0.8555813 | -2.3601763 |
| P12 cl101 | H09 | 1423000 | sample | 0.9426963 | -0.9364920 |
| P12 cl101 | H10 | 1382900 | sample | 0.9161312 | -1.3706345 |
| P12 cl101 | H11 | 1540400 | sample | 1.0204704 | 0.3345388 |
| P12 cl101 | H12 | 379570 | SAL | 0.2514541 | -12.2331837 |
| P13 cl101 | A01 | 399840 | SAL | 0.2605500 | -10.4775515 |
| P13 cl101 | A02 | 1556000 | sample | 1.0139450 | 0.1975921 |
| P13 cl101 | A03 | 1542200 | sample | 1.0049524 | 0.0701729 |
| P13 cl101 | A04 | 1515000 | sample | 0.9872279 | -0.1809722 |
| P13 cl101 | A05 | 1433400 | sample | 0.9340545 | -0.9344075 |
| P13 cl101 | A06 | 333650 | sample | 0.2174182 | -11.0887020 |
| P13 cl101 | A07 | 1371700 | sample | 0.8938486 | -1.5041005 |
| P13 cl101 | A08 | 1370200 | sample | 0.8928711 | -1.5179505 |
| P13 cl101 | A09 | 1375000 | sample | 0.8959990 | -1.4736307 |

*(continued)*

| Plate | Well | Value | Treatment | Norm | Score |
|---|---|---|---|---|---|
| P13 cl101 | A10 | 1264700 | sample | 0.8241236 | -2.4920610 |
| P13 cl101 | A11 | 1079300 | sample | 0.7033103 | -4.2039102 |
| P13 cl101 | A12 | 1637400 | empty | 1.0669881 | 0.9491807 |
| P13 cl101 | B01 | 439860 | SAL | 0.2866284 | -10.1080358 |
| P13 cl101 | B02 | 1589800 | sample | 1.0359703 | 0.5096768 |
| P13 cl101 | B03 | 1548200 | sample | 1.0088622 | 0.1255725 |
| P13 cl101 | B04 | 1204100 | sample | 0.7846344 | -3.0515975 |
| P13 cl101 | B05 | 1575400 | sample | 1.0265867 | 0.3767176 |
| P13 cl101 | B06 | 1499500 | sample | 0.9771276 | -0.3240880 |
| P13 cl101 | B07 | 1511100 | sample | 0.9846866 | -0.2169820 |
| P13 cl101 | B08 | 1508200 | sample | 0.9827968 | -0.2437585 |
| P13 cl101 | B09 | 1548700 | sample | 1.0091881 | 0.1301892 |
| P13 cl101 | B10 | 1634700 | sample | 1.0652287 | 0.9242509 |
| P13 cl101 | B11 | 1510900 | sample | 0.9845562 | -0.2188286 |
| P13 cl101 | B12 | 1699500 | empty | 1.1074547 | 1.5225671 |
| P13 cl101 | C01 | 442610 | SAL | 0.2884204 | -10.0826443 |
| P13 cl101 | C02 | 1526700 | sample | 0.9948521 | -0.0729429 |
| P13 cl101 | C03 | 1691400 | sample | 1.1021765 | 1.4477776 |
| P13 cl101 | C04 | 1499900 | sample | 0.9773882 | -0.3203947 |
| P13 cl101 | C05 | 1370100 | sample | 0.8928059 | -1.5188738 |
| P13 cl101 | C06 | 1633100 | sample | 1.0641861 | 0.9094776 |
| P13 cl101 | C07 | 1540900 | sample | 1.0041053 | 0.0581696 |
| P13 cl101 | C08 | 1560300 | sample | 1.0167470 | 0.2372952 |
| P13 cl101 | C09 | 1585300 | sample | 1.0330379 | 0.4681271 |
| P13 cl101 | C10 | 1605100 | sample | 1.0459403 | 0.6509459 |
| P13 cl101 | C11 | 1572500 | sample | 1.0246970 | 0.3499411 |
| P13 cl101 | C12 | 1559300 | DMSO | 1.0160954 | 0.2280619 |
| P13 cl101 | D01 | 1517500 | DMSO | 0.9888570 | -0.1578890 |
| P13 cl101 | D02 | 1691600 | sample | 1.1023068 | 1.4496242 |
| P13 cl101 | D03 | 1619700 | sample | 1.0554542 | 0.7857517 |
| P13 cl101 | D04 | 1300900 | sample | 0.8477128 | -2.1578164 |
| P13 cl101 | D05 | 1606700 | sample | 1.0469829 | 0.6657191 |
| P13 cl101 | D06 | 1323800 | sample | 0.8626352 | -1.9463744 |
| P13 cl101 | D07 | 1608000 | sample | 1.0478301 | 0.6777224 |
| P13 cl101 | D08 | 1628800 | sample | 1.0613841 | 0.8697745 |
| P13 cl101 | D09 | 426400 | sample | 0.2778574 | -10.2323157 |
| P13 cl101 | D10 | 1283300 | sample | 0.8362440 | -2.3203221 |
| P13 cl101 | D11 | 1618800 | sample | 1.0548677 | 0.7774418 |
| P13 cl101 | D12 | 1526000 | DMSO | 0.9943959 | -0.0794062 |
| P13 cl101 | E01 | 1672100 | DMSO | 1.0895999 | 1.2695754 |
| P13 cl101 | E02 | 1664700 | sample | 1.0847778 | 1.2012491 |
| P13 cl101 | E03 | 1660200 | sample | 1.0818454 | 1.1596994 |
| P13 cl101 | E04 | 1589400 | sample | 1.0357096 | 0.5059835 |
| P13 cl101 | E05 | 1695100 | sample | 1.1045875 | 1.4819407 |
| P13 cl101 | E06 | 1634800 | sample | 1.0652939 | 0.9251742 |
| P13 cl101 | E07 | 1640400 | sample | 1.0689430 | 0.9768805 |
| P13 cl101 | E08 | 1492200 | sample | 0.9723707 | -0.3914909 |
| P13 cl101 | E09 | 1607300 | sample | 1.0473739 | 0.6712591 |
| P13 cl101 | E10 | 1574800 | sample | 1.0261958 | 0.3711777 |
| P13 cl101 | E11 | 1523800 | sample | 0.9929623 | -0.0997194 |
| P13 cl101 | E12 | 1617900 | DMSO | 1.0542812 | 0.7691318 |
| P13 cl101 | F01 | 1647400 | DMSO | 1.0735045 | 1.0415135 |
| P13 cl101 | F02 | 1694300 | sample | 1.1040662 | 1.4745541 |
| P13 cl101 | F03 | 1570900 | sample | 1.0236544 | 0.3351679 |
| P13 cl101 | F04 | 742530 | sample | 0.4838590 | -7.3134004 |
| P13 cl101 | F05 | 1287800 | sample | 0.8391763 | -2.2787723 |
| P13 cl101 | F06 | 1728800 | sample | 1.1265476 | 1.7931021 |
| P13 cl101 | F07 | 1571500 | sample | 1.0240454 | 0.3407079 |
| P13 cl101 | F08 | 1615600 | sample | 1.0527825 | 0.7478953 |
| P13 cl101 | F09 | 1537800 | sample | 1.0020852 | 0.0295465 |
| P13 cl101 | F10 | 1635500 | sample | 1.0657500 | 0.9316375 |
| P13 cl101 | F11 | 1652200 | sample | 1.0766323 | 1.0858332 |

*(continued)*

| Plate | Well | Value | Treatment | Norm | Score |
|---|---|---|---|---|---|
| P13 cl101 | F12 | 440540 | SAL | 0.2870715 | -10.1017572 |
| P13 cl101 | G01 | 1629400 | empty | 1.0617751 | 0.8753145 |
| P13 cl101 | G02 | 1528200 | sample | 0.9958295 | -0.0590930 |
| P13 cl101 | G03 | 1570100 | sample | 1.0231331 | 0.3277813 |
| P13 cl101 | G04 | 1531400 | sample | 0.9979148 | -0.0295465 |
| P13 cl101 | G05 | 1539800 | sample | 1.0033885 | 0.0480130 |
| P13 cl101 | G06 | 1488200 | sample | 0.9697641 | -0.4284240 |
| P13 cl101 | G07 | 1515800 | sample | 0.9877493 | -0.1735856 |
| P13 cl101 | G08 | 1507900 | sample | 0.9826013 | -0.2465285 |
| P13 cl101 | G09 | 1418000 | sample | 0.9240193 | -1.0765999 |
| P13 cl101 | G10 | 1504200 | sample | 0.9801903 | -0.2806916 |
| P13 cl101 | G11 | 1489700 | sample | 0.9707416 | -0.4145741 |
| P13 cl101 | G12 | 393830 | SAL | 0.2566337 | -10.5330434 |
| P13 cl101 | H01 | 1586700 | empty | 1.0339502 | 0.4810536 |
| P13 cl101 | H02 | 1469800 | sample | 0.9577740 | -0.5983162 |
| P13 cl101 | H03 | 1627900 | sample | 1.0607976 | 0.8614646 |
| P13 cl101 | H04 | 1609900 | sample | 1.0490682 | 0.6952656 |
| P13 cl101 | H05 | 1502500 | sample | 0.9790825 | -0.2963881 |
| P13 cl101 | H06 | 1475200 | sample | 0.9612928 | -0.5484566 |
| P13 cl101 | H07 | 1371800 | sample | 0.8939137 | -1.5031772 |
| P13 cl101 | H08 | 1305000 | sample | 0.8503845 | -2.1199600 |
| P13 cl101 | H09 | 1416900 | sample | 0.9233025 | -1.0867565 |
| P13 cl101 | H10 | 1473200 | sample | 0.9599896 | -0.5669231 |
| P13 cl101 | H11 | 117840 | sample | 0.0767887 | -13.0813351 |
| P13 cl101 | H12 | 341220 | SAL | 0.2223511 | -11.0188061 |
| P14 cl101 | A01 | 322770 | SAL | 0.2152589 | -8.8135457 |
| P14 cl101 | A02 | 1406000 | sample | 0.9376771 | -0.6999574 |
| P14 cl101 | A03 | 1445900 | sample | 0.9642869 | -0.4010992 |
| P14 cl101 | A04 | 1184400 | sample | 0.7898896 | -2.3597814 |
| P14 cl101 | A05 | 1338500 | sample | 0.8926606 | -1.2055446 |
| P14 cl101 | A06 | 1415800 | sample | 0.9442129 | -0.6265536 |
| P14 cl101 | A07 | 1407400 | sample | 0.9386108 | -0.6894711 |
| P14 cl101 | A08 | 1331900 | sample | 0.8882590 | -1.2549798 |
| P14 cl101 | A09 | 1003800 | sample | 0.6694455 | -3.7125080 |
| P14 cl101 | A10 | 1610700 | sample | 1.0741939 | 0.8332826 |
| P14 cl101 | A11 | 1538500 | sample | 1.0260429 | 0.2924916 |
| P14 cl101 | A12 | 1612500 | empty | 1.0753943 | 0.8467649 |
| P14 cl101 | B01 | 343850 | SAL | 0.2293174 | -8.6556527 |
| P14 cl101 | B02 | 1498800 | sample | 0.9995665 | -0.0048686 |
| P14 cl101 | B03 | 1407500 | sample | 0.9386775 | -0.6887221 |
| P14 cl101 | B04 | 1354800 | sample | 0.9035313 | -1.0834546 |
| P14 cl101 | B05 | 1589600 | sample | 1.0601220 | 0.6752398 |
| P14 cl101 | B06 | 1356000 | sample | 0.9043316 | -1.0744664 |
| P14 cl101 | B07 | 577060 | sample | 0.3848478 | -6.9088676 |
| P14 cl101 | B08 | 1463700 | sample | 0.9761579 | -0.2677740 |
| P14 cl101 | B09 | 1543900 | sample | 1.0296442 | 0.3329385 |
| P14 cl101 | B10 | 1615200 | sample | 1.0771950 | 0.8669884 |
| P14 cl101 | B11 | 1554300 | sample | 1.0365801 | 0.4108364 |
| P14 cl101 | B12 | 1684300 | empty | 1.1232785 | 1.3845599 |
| P14 cl101 | C01 | 359680 | SAL | 0.2398746 | -8.5370831 |
| P14 cl101 | C02 | 1409500 | sample | 0.9400113 | -0.6737417 |
| P14 cl101 | C03 | 1407200 | sample | 0.9384774 | -0.6909692 |
| P14 cl101 | C04 | 1366200 | sample | 0.9111341 | -0.9980666 |
| P14 cl101 | C05 | 1444900 | sample | 0.9636200 | -0.4085893 |
| P14 cl101 | C06 | 1554600 | sample | 1.0367802 | 0.4130835 |
| P14 cl101 | C07 | 1500100 | sample | 1.0004335 | 0.0048686 |
| P14 cl101 | C08 | 1535900 | sample | 1.0243089 | 0.2730171 |
| P14 cl101 | C09 | 803370 | sample | 0.5357765 | -5.2137649 |
| P14 cl101 | C10 | 1665700 | sample | 1.1108740 | 1.2452425 |
| P14 cl101 | C11 | 161600 | sample | 0.1077729 | -10.0207381 |
| P14 cl101 | C12 | 1641000 | DMSO | 1.0944013 | 1.0602351 |
| P14 cl101 | D01 | 1289100 | DMSO | 0.8597152 | -1.5755595 |

*(continued)*

| Plate | Well | Value | Treatment | Norm | Score |
|---|---|---|---|---|---|
| P14 cl101 | D02 | 1513000 | sample | 1.0090366 | 0.1014919 |
| P14 cl101 | D03 | 1440300 | sample | 0.9605522 | -0.4430442 |
| P14 cl101 | D04 | 1436600 | sample | 0.9580846 | -0.4707578 |
| P14 cl101 | D05 | 1457500 | sample | 0.9720231 | -0.3142131 |
| P14 cl101 | D06 | 1622100 | sample | 1.0817967 | 0.9186706 |
| P14 cl101 | D07 | 1559600 | sample | 1.0401147 | 0.4505344 |
| P14 cl101 | D08 | 1593800 | sample | 1.0629231 | 0.7066985 |
| P14 cl101 | D09 | 1639900 | sample | 1.0936677 | 1.0519959 |
| P14 cl101 | D10 | 1626500 | sample | 1.0847311 | 0.9516274 |
| P14 cl101 | D11 | 768730 | sample | 0.5126746 | -5.4732247 |
| P14 cl101 | D12 | 1547100 | DMSO | 1.0317783 | 0.3569071 |
| P14 cl101 | E01 | 1361000 | DMSO | 0.9076661 | -1.0370155 |
| P14 cl101 | E02 | 1481500 | sample | 0.9880289 | -0.1344487 |
| P14 cl101 | E03 | 1436600 | sample | 0.9580846 | -0.4707578 |
| P14 cl101 | E04 | 1456100 | sample | 0.9710894 | -0.3246993 |
| P14 cl101 | E05 | 1609000 | sample | 1.0730601 | 0.8205493 |
| P14 cl101 | E06 | 1623300 | sample | 1.0825970 | 0.9276589 |
| P14 cl101 | E07 | 1696600 | sample | 1.1314815 | 1.4766891 |
| P14 cl101 | E08 | 1587100 | sample | 1.0584548 | 0.6565143 |
| P14 cl101 | E09 | 1607000 | sample | 1.0717263 | 0.8055689 |
| P14 cl101 | E10 | 1652500 | sample | 1.1020708 | 1.1463721 |
| P14 cl101 | E11 | 1694600 | sample | 1.1301477 | 1.4617087 |
| P14 cl101 | E12 | 1743400 | DMSO | 1.1626930 | 1.8272295 |
| P14 cl101 | F01 | 1351400 | DMSO | 0.9012638 | -1.1089212 |
| P14 cl101 | F02 | 1542000 | sample | 1.0283771 | 0.3187072 |
| P14 cl101 | F03 | 1438800 | sample | 0.9595518 | -0.4542795 |
| P14 cl101 | F04 | 1433800 | sample | 0.9562173 | -0.4917304 |
| P14 cl101 | F05 | 1550100 | sample | 1.0337791 | 0.3793776 |
| P14 cl101 | F06 | 1647800 | sample | 1.0989363 | 1.1111683 |
| P14 cl101 | F07 | 1617900 | sample | 1.0789956 | 0.8872119 |
| P14 cl101 | F08 | 1593000 | sample | 1.0623895 | 0.7007064 |
| P14 cl101 | F09 | 1626900 | sample | 1.0849978 | 0.9546235 |
| P14 cl101 | F10 | 1567000 | sample | 1.0450499 | 0.5059617 |
| P14 cl101 | F11 | 1586000 | sample | 1.0577212 | 0.6482751 |
| P14 cl101 | F12 | 479320 | SAL | 0.3196639 | -7.6409579 |
| P14 cl101 | G01 | 1337800 | empty | 0.8921938 | -1.2107877 |
| P14 cl101 | G02 | 1555800 | sample | 1.0375804 | 0.4220717 |
| P14 cl101 | G03 | 1462700 | sample | 0.9754910 | -0.2752641 |
| P14 cl101 | G04 | 1434300 | sample | 0.9565507 | -0.4879853 |
| P14 cl101 | G05 | 1602900 | sample | 1.0689920 | 0.7748592 |
| P14 cl101 | G06 | 1504900 | sample | 1.0036347 | 0.0408215 |
| P14 cl101 | G07 | 1508100 | sample | 1.0057688 | 0.0647901 |
| P14 cl101 | G08 | 1514100 | sample | 1.0097702 | 0.1097311 |
| P14 cl101 | G09 | 1545200 | sample | 1.0305112 | 0.3426758 |
| P14 cl101 | G10 | 1530600 | sample | 1.0207743 | 0.2333191 |
| P14 cl101 | G11 | 590410 | sample | 0.3937510 | -6.8088737 |
| P14 cl101 | G12 | 430680 | SAL | 0.2872253 | -8.0052803 |
| P14 cl101 | H01 | 1241900 | empty | 0.8282370 | -1.9290960 |
| P14 cl101 | H02 | 1467000 | sample | 0.9783587 | -0.2430564 |
| P14 cl101 | H03 | 1342400 | sample | 0.8952616 | -1.1763329 |
| P14 cl101 | H04 | 1351100 | sample | 0.9010637 | -1.1111683 |
| P14 cl101 | H05 | 1435800 | sample | 0.9575511 | -0.4767500 |
| P14 cl101 | H06 | 1440700 | sample | 0.9608190 | -0.4400481 |
| P14 cl101 | H07 | 1414300 | sample | 0.9432125 | -0.6377889 |
| P14 cl101 | H08 | 1386300 | sample | 0.9245390 | -0.8475139 |
| P14 cl101 | H09 | 1442200 | sample | 0.9618193 | -0.4288128 |
| P14 cl101 | H10 | 1595000 | sample | 1.0637234 | 0.7156868 |
| P14 cl101 | H11 | 1614700 | sample | 1.0768615 | 0.8632433 |
| P14 cl101 | H12 | 383100 | SAL | 0.2554937 | -8.3616631 |
| P15 cl101 | A01 | 357900 | SAL | 0.2483175 | -9.7367527 |
| P15 cl101 | A02 | 1423400 | sample | 0.9875807 | -0.1608712 |

*(continued)*

| Plate | Well | Value | Treatment | Norm | Score |
|---|---|---|---|---|---|
| P15 cl101 | A03 | 1441200 | sample | 0.9999306 | -0.0008987 |
| P15 cl101 | A04 | 1294200 | sample | 0.8979394 | -1.3220199 |
| P15 cl101 | A05 | 1230700 | sample | 0.8538819 | -1.8927082 |
| P15 cl101 | A06 | 1351500 | sample | 0.9376951 | -0.8070522 |
| P15 cl101 | A07 | 1320600 | sample | 0.9162562 | -1.0847573 |
| P15 cl101 | A08 | 1316400 | sample | 0.9133421 | -1.1225036 |
| P15 cl101 | A09 | 1298200 | sample | 0.9007146 | -1.2860710 |
| P15 cl101 | A10 | 1378500 | sample | 0.9564282 | -0.5643973 |
| P15 cl101 | A11 | 1465800 | sample | 1.0169985 | 0.2201869 |
| P15 cl101 | A12 | 1365900 | empty | 0.9476861 | -0.6776363 |
| P15 cl101 | B01 | 392690 | SAL | 0.2724554 | -9.4240873 |
| P15 cl101 | B02 | 1472600 | sample | 1.0217165 | 0.2812999 |
| P15 cl101 | B03 | 1411500 | sample | 0.9793242 | -0.2678191 |
| P15 cl101 | B04 | 1410700 | sample | 0.9787692 | -0.2750089 |
| P15 cl101 | B05 | 1460900 | sample | 1.0135988 | 0.1761495 |
| P15 cl101 | B06 | 1566700 | sample | 1.0870048 | 1.1269972 |
| P15 cl101 | B07 | 1509600 | sample | 1.0473878 | 0.6138270 |
| P15 cl101 | B08 | 1302800 | sample | 0.9039062 | -1.2447298 |
| P15 cl101 | B09 | 1532100 | sample | 1.0629987 | 0.8160395 |
| P15 cl101 | B10 | 1468700 | sample | 1.0190106 | 0.2462498 |
| P15 cl101 | B11 | 1462900 | sample | 1.0149865 | 0.1941239 |
| P15 cl101 | B12 | 1607500 | empty | 1.1153126 | 1.4936757 |
| P15 cl101 | C01 | 391160 | SAL | 0.2713939 | -9.4378378 |
| P15 cl101 | C02 | 1463200 | sample | 1.0151946 | 0.1968201 |
| P15 cl101 | C03 | 1570200 | sample | 1.0894332 | 1.1584525 |
| P15 cl101 | C04 | 1399400 | sample | 0.9709290 | -0.3765645 |
| P15 cl101 | C05 | 1493600 | sample | 1.0362867 | 0.4700315 |
| P15 cl101 | C06 | 1554300 | sample | 1.0784014 | 1.0155557 |
| P15 cl101 | C07 | 1554000 | sample | 1.0781933 | 1.0128595 |
| P15 cl101 | C08 | 1507300 | sample | 1.0457920 | 0.5931564 |
| P15 cl101 | C09 | 294110 | sample | 0.2040588 | -10.3100474 |
| P15 cl101 | C10 | 1465300 | sample | 1.0166516 | 0.2156932 |
| P15 cl101 | C11 | 1530300 | sample | 1.0617498 | 0.7998625 |
| P15 cl101 | C12 | 1506200 | DMSO | 1.0450288 | 0.5832705 |
| P15 cl101 | D01 | 1419500 | DMSO | 0.9848748 | -0.1959214 |
| P15 cl101 | D02 | 540340 | sample | 0.3748977 | -8.0971245 |
| P15 cl101 | D03 | 1509700 | sample | 1.0474572 | 0.6147258 |
| P15 cl101 | D04 | 1511500 | sample | 1.0487060 | 0.6309027 |
| P15 cl101 | D05 | 1522500 | sample | 1.0563380 | 0.7297622 |
| P15 cl101 | D06 | 1540300 | sample | 1.0686880 | 0.8897346 |
| P15 cl101 | D07 | 1525800 | sample | 1.0586276 | 0.7594200 |
| P15 cl101 | D08 | 1488200 | sample | 1.0325401 | 0.4215006 |
| P15 cl101 | D09 | 1544000 | sample | 1.0712551 | 0.9229874 |
| P15 cl101 | D10 | 1498500 | sample | 1.0396864 | 0.5140689 |
| P15 cl101 | D11 | 1492200 | sample | 1.0353153 | 0.4574494 |
| P15 cl101 | D12 | 1438500 | DMSO | 0.9980573 | -0.0251642 |
| P15 cl101 | E01 | 1494600 | DMSO | 1.0369805 | 0.4790188 |
| P15 cl101 | E02 | 1478600 | sample | 1.0258794 | 0.3352233 |
| P15 cl101 | E03 | 1195400 | sample | 0.8293901 | -2.2099571 |
| P15 cl101 | E04 | 1628900 | sample | 1.1301603 | 1.6860022 |
| P15 cl101 | E05 | 1671400 | sample | 1.1596475 | 2.0679590 |
| P15 cl101 | E06 | 1599100 | sample | 1.1094845 | 1.4181831 |
| P15 cl101 | E07 | 1515600 | sample | 1.0515507 | 0.6677503 |
| P15 cl101 | E08 | 1386200 | sample | 0.9617706 | -0.4951957 |
| P15 cl101 | E09 | 1533800 | sample | 1.0641782 | 0.8313177 |
| P15 cl101 | E10 | 1507400 | sample | 1.0458614 | 0.5940552 |
| P15 cl101 | E11 | 1543900 | sample | 1.0711857 | 0.9220886 |
| P15 cl101 | E12 | 1541400 | DMSO | 1.0694512 | 0.8996206 |
| P15 cl101 | F01 | 1404700 | DMSO | 0.9746063 | -0.3289322 |
| P15 cl101 | F02 | 423430 | sample | 0.2937834 | -9.1478202 |
| P15 cl101 | F03 | 1348200 | sample | 0.9354055 | -0.8367101 |
| P15 cl101 | F04 | 1390700 | sample | 0.9648928 | -0.4547533 |

*(continued)*

| Plate | Well | Value | Treatment | Norm | Score |
|---|---|---|---|---|---|
| P15 cl101 | F05 | 1537300 | sample | 1.0666065 | 0.8627730 |
| P15 cl101 | F06 | 1517100 | sample | 1.0525914 | 0.6812312 |
| P15 cl101 | F07 | 765580 | sample | 0.5311732 | -6.0728434 |
| P15 cl101 | F08 | 1408200 | sample | 0.9770346 | -0.2974769 |
| P15 cl101 | F09 | 1479400 | sample | 1.0264345 | 0.3424130 |
| P15 cl101 | F10 | 1448000 | sample | 1.0046486 | 0.0602144 |
| P15 cl101 | F11 | 1386700 | sample | 0.9621175 | -0.4907021 |
| P15 cl101 | F12 | 426370 | SAL | 0.2958232 | -9.1213978 |
| P15 cl101 | G01 | 1346200 | empty | 0.9340179 | -0.8546845 |
| P15 cl101 | G02 | 1441400 | sample | 1.0000694 | 0.0008987 |
| P15 cl101 | G03 | 1466300 | sample | 1.0173455 | 0.2246805 |
| P15 cl101 | G04 | 240500 | sample | 0.1668632 | -10.7918522 |
| P15 cl101 | G05 | 1429900 | sample | 0.9920905 | -0.1024543 |
| P15 cl101 | G06 | 1373800 | sample | 0.9531673 | -0.6066373 |
| P15 cl101 | G07 | 1372500 | sample | 0.9522653 | -0.6183206 |
| P15 cl101 | G08 | 732330 | sample | 0.5081038 | -6.3716684 |
| P15 cl101 | G09 | 1410800 | sample | 0.9788385 | -0.2741102 |
| P15 cl101 | G10 | 1485100 | sample | 1.0303892 | 0.3936402 |
| P15 cl101 | G11 | 1410800 | sample | 0.9788385 | -0.2741102 |
| P15 cl101 | G12 | 363600 | SAL | 0.2522723 | -9.6855255 |
| P15 cl101 | H01 | 1339400 | empty | 0.9292999 | -0.9157976 |
| P15 cl101 | H02 | 902690 | sample | 0.6263026 | -4.8406058 |
| P15 cl101 | H03 | 1280600 | sample | 0.8885034 | -1.4442460 |
| P15 cl101 | H04 | 1348500 | sample | 0.9356137 | -0.8340139 |
| P15 cl101 | H05 | 1382300 | sample | 0.9590647 | -0.5302459 |
| P15 cl101 | H06 | 1323300 | sample | 0.9181295 | -1.0604918 |
| P15 cl101 | H07 | 1344800 | sample | 0.9330466 | -0.8672666 |
| P15 cl101 | H08 | 1303200 | sample | 0.9041837 | -1.2411349 |
| P15 cl101 | H09 | 1303700 | sample | 0.9045306 | -1.2366413 |
| P15 cl101 | H10 | 1387200 | sample | 0.9624644 | -0.4862085 |
| P15 cl101 | H11 | 1001400 | sample | 0.6947894 | -3.9534775 |
| P15 cl101 | H12 | 329280 | SAL | 0.2284604 | -9.9939669 |
| P16 cl101 | A01 | 362290 | SAL | 0.2463050 | -7.1011130 |
| P16 cl101 | A02 | 1284700 | sample | 0.8734108 | -1.1926893 |
| P16 cl101 | A03 | 1358200 | sample | 0.9233802 | -0.7218909 |
| P16 cl101 | A04 | 1255000 | sample | 0.8532191 | -1.3829302 |
| P16 cl101 | A05 | 1290100 | sample | 0.8770821 | -1.1581000 |
| P16 cl101 | A06 | 1343900 | sample | 0.9136583 | -0.8134884 |
| P16 cl101 | A07 | 1197000 | sample | 0.8137875 | -1.7544446 |
| P16 cl101 | A08 | 1082200 | sample | 0.7357400 | -2.4897869 |
| P16 cl101 | A09 | 1296900 | sample | 0.8817051 | -1.1145431 |
| P16 cl101 | A10 | 1279500 | sample | 0.8698756 | -1.2259974 |
| P16 cl101 | A11 | 1394400 | sample | 0.9479910 | -0.4900147 |
| P16 cl101 | A12 | 1433600 | empty | 0.9746414 | -0.2389222 |
| P16 cl101 | B01 | 389680 | SAL | 0.2649262 | -6.9256686 |
| P16 cl101 | B02 | 1517800 | sample | 1.0318852 | 0.3004142 |
| P16 cl101 | B03 | 1550900 | sample | 1.0543885 | 0.5124336 |
| P16 cl101 | B04 | 1532100 | sample | 1.0416072 | 0.3920117 |
| P16 cl101 | B05 | 96037 | sample | 0.0652913 | -8.8065754 |
| P16 cl101 | B06 | 1614100 | sample | 1.0973554 | 0.9172562 |
| P16 cl101 | B07 | 1507700 | sample | 1.0250187 | 0.2357195 |
| P16 cl101 | B08 | 1423400 | sample | 0.9677068 | -0.3042575 |
| P16 cl101 | B09 | 1430200 | sample | 0.9723299 | -0.2607006 |
| P16 cl101 | B10 | 1562900 | sample | 1.0625467 | 0.5892987 |
| P16 cl101 | B11 | 763220 | sample | 0.5188796 | -4.5329879 |
| P16 cl101 | B12 | 1726700 | empty | 1.1739071 | 1.6385065 |
| P16 cl101 | C01 | 411090 | SAL | 0.2794819 | -6.7885285 |
| P16 cl101 | C02 | 1331300 | sample | 0.9050921 | -0.8941967 |
| P16 cl101 | C03 | 1384800 | sample | 0.9414644 | -0.5515067 |
| P16 cl101 | C04 | 1350700 | sample | 0.9182813 | -0.7699315 |
| P16 cl101 | C05 | 1389500 | sample | 0.9446597 | -0.5214012 |
| P16 cl101 | C06 | 1550500 | sample | 1.0541165 | 0.5098715 |

*(continued)*

| Plate | Well | Value | Treatment | Norm | Score |
|---|---|---|---|---|---|
| P16 cl101 | C07 | 1484700 | sample | 1.0093820 | 0.0883948 |
| P16 cl101 | C08 | 1474700 | sample | 1.0025835 | 0.0243406 |
| P16 cl101 | C09 | 1489300 | sample | 1.0125093 | 0.1178597 |
| P16 cl101 | C10 | 1605500 | sample | 1.0915086 | 0.8621696 |
| P16 cl101 | C11 | 1624400 | sample | 1.1043579 | 0.9832320 |
| P16 cl101 | C12 | 1547500 | DMSO | 1.0520770 | 0.4906552 |
| P16 cl101 | D01 | 1456800 | DMSO | 0.9904140 | -0.0903164 |
| P16 cl101 | D02 | 329780 | sample | 0.2242029 | -7.3093532 |
| P16 cl101 | D03 | 1427500 | sample | 0.9704943 | -0.2779952 |
| P16 cl101 | D04 | 1396600 | sample | 0.9494867 | -0.4759227 |
| P16 cl101 | D05 | 1126300 | sample | 0.7657217 | -2.2073078 |
| P16 cl101 | D06 | 1512400 | sample | 1.0282140 | 0.2658249 |
| P16 cl101 | D07 | 1572000 | sample | 1.0687334 | 0.6475880 |
| P16 cl101 | D08 | 1588100 | sample | 1.0796791 | 0.7507153 |
| P16 cl101 | D09 | 1524800 | sample | 1.0366442 | 0.3452522 |
| P16 cl101 | D10 | 1591700 | sample | 1.0821266 | 0.7737748 |
| P16 cl101 | D11 | 1598100 | sample | 1.0864777 | 0.8147695 |
| P16 cl101 | D12 | 1538300 | DMSO | 1.0458223 | 0.4317253 |
| P16 cl101 | E01 | 1411000 | DMSO | 0.9592766 | -0.3836847 |
| P16 cl101 | E02 | 1543100 | sample | 1.0490856 | 0.4624713 |
| P16 cl101 | E03 | 1614600 | sample | 1.0976953 | 0.9204589 |
| P16 cl101 | E04 | 1544900 | sample | 1.0503093 | 0.4740011 |
| P16 cl101 | E05 | 1364100 | sample | 0.9273914 | -0.6840989 |
| P16 cl101 | E06 | 1630800 | sample | 1.1087090 | 1.0242267 |
| P16 cl101 | E07 | 1605800 | sample | 1.0917126 | 0.8640912 |
| P16 cl101 | E08 | 1477900 | sample | 1.0047590 | 0.0448379 |
| P16 cl101 | E09 | 1582300 | sample | 1.0757359 | 0.7135638 |
| P16 cl101 | E10 | 1587800 | sample | 1.0794752 | 0.7487936 |
| P16 cl101 | E11 | 1620100 | sample | 1.1014345 | 0.9556887 |
| P16 cl101 | E12 | 1539100 | DMSO | 1.0463662 | 0.4368497 |
| P16 cl101 | F01 | 1478200 | DMSO | 1.0049629 | 0.0467596 |
| P16 cl101 | F02 | 1471100 | sample | 1.0001360 | 0.0012811 |
| P16 cl101 | F03 | 1484800 | sample | 1.0094500 | 0.0890353 |
| P16 cl101 | F04 | 1516500 | sample | 1.0310014 | 0.2920872 |
| P16 cl101 | F05 | 1527100 | sample | 1.0382079 | 0.3599846 |
| P16 cl101 | F06 | 1603500 | sample | 1.0901489 | 0.8493587 |
| P16 cl101 | F07 | 1568900 | sample | 1.0666259 | 0.6277312 |
| P16 cl101 | F08 | 1441300 | sample | 0.9798763 | -0.1896004 |
| P16 cl101 | F09 | 1515400 | sample | 1.0302536 | 0.2850412 |
| P16 cl101 | F10 | 1627000 | sample | 1.1061255 | 0.9998861 |
| P16 cl101 | F11 | 1557400 | sample | 1.0588075 | 0.5540689 |
| P16 cl101 | F12 | 415520 | SAL | 0.2824937 | -6.7601525 |
| P16 cl101 | G01 | 1472300 | empty | 1.0009518 | 0.0089676 |
| P16 cl101 | G02 | 274550 | sample | 0.1866544 | -7.6631246 |
| P16 cl101 | G03 | 1582900 | sample | 1.0761439 | 0.7174071 |
| P16 cl101 | G04 | 1396100 | sample | 0.9491468 | -0.4791254 |
| P16 cl101 | G05 | 1174500 | sample | 0.7984907 | -1.8985666 |
| P16 cl101 | G06 | 1469500 | sample | 0.9990482 | -0.0089676 |
| P16 cl101 | G07 | 1473000 | sample | 1.0014277 | 0.0134514 |
| P16 cl101 | G08 | 1436600 | sample | 0.9766809 | -0.2197059 |
| P16 cl101 | G09 | 1308600 | sample | 0.8896594 | -1.0395997 |
| P16 cl101 | G10 | 1470700 | sample | 0.9998640 | -0.0012811 |
| P16 cl101 | G11 | 1474600 | sample | 1.0025155 | 0.0237001 |
| P16 cl101 | G12 | 381350 | SAL | 0.2592630 | -6.9790257 |
| P16 cl101 | H01 | 1270600 | empty | 0.8638249 | -1.2830057 |
| P16 cl101 | H02 | 1354100 | sample | 0.9205928 | -0.7481531 |
| P16 cl101 | H03 | 364620 | sample | 0.2478890 | -7.0861884 |
| P16 cl101 | H04 | 286860 | sample | 0.1950235 | -7.5842739 |
| P16 cl101 | H05 | 1385500 | sample | 0.9419403 | -0.5470229 |
| P16 cl101 | H06 | 1367100 | sample | 0.9294310 | -0.6648826 |
| P16 cl101 | H07 | 1348500 | sample | 0.9167856 | -0.7840234 |

*(continued)*

| Plate | Well | Value | Treatment | Norm | Score |
|---|---|---|---|---|---|
| P16 cl101 | H08 | 1251300 | sample | 0.8507037 | -1.4066303 |
| P16 cl101 | H09 | 1268000 | sample | 0.8620572 | -1.2996598 |
| P16 cl101 | H10 | 1485300 | sample | 1.0097899 | 0.0922381 |
| P16 cl101 | H11 | 1440900 | sample | 0.9796043 | -0.1921626 |
| P16 cl101 | H12 | 328930 | SAL | 0.2236250 | -7.3147978 |

#### clone 119

| Plate | Well | Value | Treatment | Norm | Score |
|---|---|---|---|---|---|
| P1 cl119 | A01 | 335350 | SAL | 0.2257489 | -8.0058364 |
| P1 cl119 | A02 | 1286700 | sample | 0.8661730 | -1.3837850 |
| P1 cl119 | A03 | 1101200 | sample | 0.7412992 | -2.6749928 |
| P1 cl119 | A04 | 1134800 | sample | 0.7639179 | -2.4411136 |
| P1 cl119 | A05 | 1052900 | sample | 0.7087849 | -3.0111940 |
| P1 cl119 | A06 | 1135600 | sample | 0.7644564 | -2.4355451 |
| P1 cl119 | A07 | 1180200 | sample | 0.7944800 | -2.1250983 |
| P1 cl119 | A08 | 1069500 | sample | 0.7199596 | -2.8956466 |
| P1 cl119 | A09 | 1189700 | sample | 0.8008751 | -2.0589718 |
| P1 cl119 | A10 | 1167800 | sample | 0.7861326 | -2.2114109 |
| P1 cl119 | A11 | 1052800 | sample | 0.7087176 | -3.0118901 |
| P1 cl119 | A12 | 1000900 | empty | 0.6737799 | -3.3731499 |
| P1 cl119 | B01 | 366210 | SAL | 0.2465231 | -7.7910295 |
| P1 cl119 | B02 | 1419800 | sample | 0.9557725 | -0.4573173 |
| P1 cl119 | B03 | 1500100 | sample | 1.0098283 | 0.1016261 |
| P1 cl119 | B04 | 1497400 | sample | 1.0080108 | 0.0828322 |
| P1 cl119 | B05 | 1515200 | sample | 1.0199933 | 0.2067325 |
| P1 cl119 | B06 | 1405800 | sample | 0.9463480 | -0.5547669 |
| P1 cl119 | B07 | 1467100 | sample | 0.9876136 | -0.1280767 |
| P1 cl119 | B08 | 1506600 | sample | 1.0142040 | 0.1468705 |
| P1 cl119 | B09 | 1024100 | sample | 0.6893975 | -3.2116619 |
| P1 cl119 | B10 | 1591300 | sample | 1.0712218 | 0.7364409 |
| P1 cl119 | B11 | 1542600 | sample | 1.0384382 | 0.3974553 |
| P1 cl119 | B12 | 1405600 | empty | 0.9462134 | -0.5561590 |
| P1 cl119 | C01 | 362090 | SAL | 0.2437496 | -7.8197076 |
| P1 cl119 | C02 | 1502900 | sample | 1.0117132 | 0.1211160 |
| P1 cl119 | C03 | 1556100 | sample | 1.0475261 | 0.4914246 |
| P1 cl119 | C04 | 1485300 | sample | 0.9998654 | -0.0013921 |
| P1 cl119 | C05 | 1515700 | sample | 1.0203299 | 0.2102128 |
| P1 cl119 | C06 | 1414200 | sample | 0.9520027 | -0.4962971 |
| P1 cl119 | C07 | 1487500 | sample | 1.0013463 | 0.0139214 |
| P1 cl119 | C08 | 1530700 | sample | 1.0304275 | 0.3146231 |
| P1 cl119 | C09 | 1582000 | sample | 1.0649613 | 0.6717065 |
| P1 cl119 | C10 | 1569400 | sample | 1.0564793 | 0.5840018 |
| P1 cl119 | C11 | 1537500 | sample | 1.0350050 | 0.3619558 |
| P1 cl119 | C12 | 1508800 | DMSO | 1.0156850 | 0.1621841 |
| P1 cl119 | D01 | 1395600 | DMSO | 0.9394817 | -0.6257659 |
| P1 cl119 | D02 | 1475100 | sample | 0.9929990 | -0.0723912 |
| P1 cl119 | D03 | 1490100 | sample | 1.0030966 | 0.0320192 |
| P1 cl119 | D04 | 1506500 | sample | 1.0141367 | 0.1461745 |
| P1 cl119 | D05 | 685490 | sample | 0.4614541 | -5.5686208 |
| P1 cl119 | D06 | 1504900 | sample | 1.0130596 | 0.1350374 |
| P1 cl119 | D07 | 1529700 | sample | 1.0297543 | 0.3076625 |
| P1 cl119 | D08 | 336030 | sample | 0.2262067 | -8.0011031 |
| P1 cl119 | D09 | 1564000 | sample | 1.0528442 | 0.5464141 |
| P1 cl119 | D10 | 1639200 | sample | 1.1034668 | 1.0698579 |
| P1 cl119 | D11 | 280100 | sample | 0.1885560 | -8.3904145 |
| P1 cl119 | D12 | 1434700 | DMSO | 0.9658028 | -0.3536030 |
| P1 cl119 | E01 | 1272200 | DMSO | 0.8564120 | -1.4847150 |
| P1 cl119 | E02 | 1601500 | sample | 1.0780882 | 0.8074399 |
| P1 cl119 | E03 | 1724200 | sample | 1.1606866 | 1.6615165 |

*(continued)*

| Plate | Well | Value | Treatment | Norm | Score |
|---|---|---|---|---|---|
| P1 cl119 | E04 | 1596500 | sample | 1.0747223 | 0.7726365 |
| P1 cl119 | E05 | 1646300 | sample | 1.1082464 | 1.1192788 |
| P1 cl119 | E06 | 1634200 | sample | 1.1001010 | 1.0350544 |
| P1 cl119 | E07 | 1582800 | sample | 1.0654998 | 0.6772750 |
| P1 cl119 | E08 | 1597600 | sample | 1.0754628 | 0.7802932 |
| P1 cl119 | E09 | 1493800 | sample | 1.0055873 | 0.0577737 |
| P1 cl119 | E10 | 1639100 | sample | 1.1033995 | 1.0691618 |
| P1 cl119 | E11 | 1646600 | sample | 1.1084483 | 1.1213670 |
| P1 cl119 | E12 | 1477100 | DMSO | 0.9943453 | -0.0584698 |
| P1 cl119 | F01 | 1397600 | DMSO | 0.9408280 | -0.6118446 |
| P1 cl119 | F02 | 1531900 | sample | 1.0312353 | 0.3229760 |
| P1 cl119 | F03 | 1559400 | sample | 1.0497476 | 0.5143949 |
| P1 cl119 | F04 | 1116200 | sample | 0.7513968 | -2.5705824 |
| P1 cl119 | F05 | 1527900 | sample | 1.0285426 | 0.2951332 |
| P1 cl119 | F06 | 1562800 | sample | 1.0520364 | 0.5380613 |
| P1 cl119 | F07 | 1624200 | sample | 1.0933692 | 0.9654476 |
| P1 cl119 | F08 | 1415400 | sample | 0.9528105 | -0.4879443 |
| P1 cl119 | F09 | 1250200 | sample | 0.8416022 | -1.6378501 |
| P1 cl119 | F10 | 1552200 | sample | 1.0449007 | 0.4642780 |
| P1 cl119 | F11 | 1520800 | sample | 1.0237630 | 0.2457123 |
| P1 cl119 | F12 | 419480 | SAL | 0.2823830 | -7.4202336 |
| P1 cl119 | G01 | 1247800 | empty | 0.8399865 | -1.6545558 |
| P1 cl119 | G02 | 1492800 | sample | 1.0049142 | 0.0508130 |
| P1 cl119 | G03 | 1468100 | sample | 0.9882868 | -0.1211160 |
| P1 cl119 | G04 | 1412100 | sample | 0.9505890 | -0.5109146 |
| P1 cl119 | G05 | 1280200 | sample | 0.8617974 | -1.4290294 |
| P1 cl119 | G06 | 1432300 | sample | 0.9641871 | -0.3703087 |
| P1 cl119 | G07 | 1498300 | sample | 1.0086166 | 0.0890968 |
| P1 cl119 | G08 | 1285400 | sample | 0.8652979 | -1.3928339 |
| P1 cl119 | G09 | 1484700 | sample | 0.9994615 | -0.0055686 |
| P1 cl119 | G10 | 1485700 | sample | 1.0001346 | 0.0013921 |
| P1 cl119 | G11 | 1438200 | sample | 0.9681589 | -0.3292406 |
| P1 cl119 | G12 | 363780 | SAL | 0.2448872 | -7.8079440 |
| P1 cl119 | H01 | 1149100 | empty | 0.7735443 | -2.3415758 |
| P1 cl119 | H02 | 1304000 | sample | 0.8778189 | -1.2633650 |
| P1 cl119 | H03 | 1155500 | sample | 0.7778526 | -2.2970274 |
| P1 cl119 | H04 | 1345400 | sample | 0.9056883 | -0.9751925 |
| P1 cl119 | H05 | 1225100 | sample | 0.8247055 | -1.8125634 |
| P1 cl119 | H06 | 1298500 | sample | 0.8741165 | -1.3016488 |
| P1 cl119 | H07 | 913400 | sample | 0.6148771 | -3.9822101 |
| P1 cl119 | H08 | 1375100 | sample | 0.9256816 | -0.7684601 |
| P1 cl119 | H09 | 781500 | sample | 0.5260855 | -4.9003250 |
| P1 cl119 | H10 | 1287200 | sample | 0.8665096 | -1.3803046 |
| P1 cl119 | H11 | 1143400 | sample | 0.7697072 | -2.3812517 |
| P1 cl119 | H12 | 316010 | SAL | 0.2127297 | -8.1404561 |
| P2 cl119 | A01 | 245420 | SAL | 0.1822313 | -4.7724825 |
| P2 cl119 | A02 | 998960 | sample | 0.7417561 | -1.5071066 |
| P2 cl119 | A03 | 1009700 | sample | 0.7497308 | -1.4605661 |
| P2 cl119 | A04 | 1071000 | sample | 0.7952478 | -1.1949298 |
| P2 cl119 | A05 | 1138700 | sample | 0.8455170 | -0.9015599 |
| P2 cl119 | A06 | 1194000 | sample | 0.8865788 | -0.6619240 |
| P2 cl119 | A07 | 1064800 | sample | 0.7906441 | -1.2217968 |
| P2 cl119 | A08 | 1100100 | sample | 0.8168554 | -1.0688284 |
| P2 cl119 | A09 | 1160400 | sample | 0.8616298 | -0.8075256 |
| P2 cl119 | A10 | 1063900 | sample | 0.7899759 | -1.2256968 |
| P2 cl119 | A11 | 959520 | sample | 0.7124708 | -1.6780151 |
| P2 cl119 | A12 | 1013600 | empty | 0.7526267 | -1.4436659 |
| P2 cl119 | B01 | 284680 | SAL | 0.2113830 | -4.6023540 |
| P2 cl119 | B02 | 1343800 | sample | 0.9978095 | -0.0127835 |
| P2 cl119 | B03 | 1291300 | sample | 0.9588268 | -0.2402860 |
| P2 cl119 | B04 | 1370800 | sample | 1.0178578 | 0.1042178 |
| P2 cl119 | B05 | 1428800 | sample | 1.0609244 | 0.3555539 |

*(continued)*

| Plate | Well | Value | Treatment | Norm | Score |
|---|---|---|---|---|---|
| P2 cl119 | B06 | 1459400 | sample | 1.0836458 | 0.4881554 |
| P2 cl119 | B07 | 1354400 | sample | 1.0056803 | 0.0331504 |
| P2 cl119 | B08 | 958020 | sample | 0.7113570 | -1.6845152 |
| P2 cl119 | B09 | 1443900 | sample | 1.0721366 | 0.4209880 |
| P2 cl119 | B10 | 1556200 | sample | 1.1555226 | 0.9076267 |
| P2 cl119 | B11 | 487020 | sample | 0.3616261 | -3.7255377 |
| P2 cl119 | B12 | 1238500 | empty | 0.9196213 | -0.4690885 |
| P2 cl119 | C01 | 286000 | SAL | 0.2123631 | -4.5966339 |
| P2 cl119 | C02 | 1182000 | sample | 0.8776685 | -0.7139245 |
| P2 cl119 | C03 | 1409300 | sample | 1.0464451 | 0.2710530 |
| P2 cl119 | C04 | 1315700 | sample | 0.9769445 | -0.1345515 |
| P2 cl119 | C05 | 1522800 | sample | 1.1307221 | 0.7628917 |
| P2 cl119 | C06 | 1510600 | sample | 1.1216633 | 0.7100245 |
| P2 cl119 | C07 | 1549300 | sample | 1.1503991 | 0.8777263 |
| P2 cl119 | C08 | 1448600 | sample | 1.0756265 | 0.4413549 |
| P2 cl119 | C09 | 1501300 | sample | 1.1147578 | 0.6697240 |
| P2 cl119 | C10 | 1493500 | sample | 1.1089660 | 0.6359237 |
| P2 cl119 | C11 | 1524600 | sample | 1.1320587 | 0.7706918 |
| P2 cl119 | C12 | 1300800 | DMSO | 0.9658808 | -0.1991189 |
| P2 cl119 | D01 | 828760 | DMSO | 0.6153778 | -2.2446480 |
| P2 cl119 | D02 | 1366500 | sample | 1.0146649 | 0.0855843 |
| P2 cl119 | D03 | 1465100 | sample | 1.0878782 | 0.5128556 |
| P2 cl119 | D04 | 1298400 | sample | 0.9640988 | -0.2095190 |
| P2 cl119 | D05 | 1466300 | sample | 1.0887693 | 0.5180557 |
| P2 cl119 | D06 | 1441600 | sample | 1.0704288 | 0.4110212 |
| P2 cl119 | D07 | 1576100 | sample | 1.1702989 | 0.9938609 |
| P2 cl119 | D08 | 1489100 | sample | 1.1056989 | 0.6168568 |
| P2 cl119 | D09 | 1580000 | sample | 1.1731947 | 1.0107611 |
| P2 cl119 | D10 | 1586800 | sample | 1.1782439 | 1.0402281 |
| P2 cl119 | D11 | 1538700 | sample | 1.1425283 | 0.8317925 |
| P2 cl119 | D12 | 1414600 | DMSO | 1.0503805 | 0.2940199 |
| P2 cl119 | E01 | 993990 | DMSO | 0.7380657 | -1.5286435 |
| P2 cl119 | E02 | 1393000 | sample | 1.0343419 | 0.2004189 |
| P2 cl119 | E03 | 1462100 | sample | 1.0856506 | 0.4998555 |
| P2 cl119 | E04 | 1282300 | sample | 0.9521441 | -0.2792864 |
| P2 cl119 | E05 | 1388800 | sample | 1.0312233 | 0.1822187 |
| P2 cl119 | E06 | 774630 | sample | 0.5751847 | -2.4792140 |
| P2 cl119 | E07 | 1232700 | sample | 0.9153146 | -0.4942221 |
| P2 cl119 | E08 | 1408300 | sample | 1.0457026 | 0.2667196 |
| P2 cl119 | E09 | 1569900 | sample | 1.1656952 | 0.9669940 |
| P2 cl119 | E10 | 1600500 | sample | 1.1884166 | 1.0995954 |
| P2 cl119 | E11 | 1602800 | sample | 1.1901244 | 1.1095622 |
| P2 cl119 | E12 | 1466900 | DMSO | 1.0892148 | 0.5206557 |
| P2 cl119 | F01 | 1005800 | DMSO | 0.7468350 | -1.4774663 |
| P2 cl119 | F02 | 1349700 | sample | 1.0021905 | 0.0127835 |
| P2 cl119 | F03 | 1429100 | sample | 1.0611472 | 0.3568539 |
| P2 cl119 | F04 | 1120300 | sample | 0.8318545 | -0.9812941 |
| P2 cl119 | F05 | 1573500 | sample | 1.1683683 | 0.9825942 |
| P2 cl119 | F06 | 1551300 | sample | 1.1518842 | 0.8863931 |
| P2 cl119 | F07 | 180000 | sample | 0.1336551 | -5.0559723 |
| P2 cl119 | F08 | 1556800 | sample | 1.1559681 | 0.9102267 |
| P2 cl119 | F09 | 239170 | sample | 0.1775905 | -4.7995662 |
| P2 cl119 | F10 | 1501000 | sample | 1.1145350 | 0.6684240 |
| P2 cl119 | F11 | 1621700 | sample | 1.2041582 | 1.1914631 |
| P2 cl119 | F12 | 388730 | SAL | 0.2886430 | -4.1514657 |
| P2 cl119 | G01 | 1047900 | empty | 0.7780954 | -1.2950309 |
| P2 cl119 | G02 | 1236900 | sample | 0.9184333 | -0.4760219 |
| P2 cl119 | G03 | 1258700 | sample | 0.9346204 | -0.3815542 |
| P2 cl119 | G04 | 371830 | sample | 0.2760943 | -4.2246998 |
| P2 cl119 | G05 | 1224200 | sample | 0.9090032 | -0.5310558 |
| P2 cl119 | G06 | 1467100 | sample | 1.0893633 | 0.5215224 |
| P2 cl119 | G07 | 847420 | sample | 0.6292333 | -2.1637872 |

*(continued)*

| Plate | Well | Value | Treatment | Norm | Score |
|---|---|---|---|---|---|
| P2 cl119 | G08 | 1451200 | sample | 1.0775571 | 0.4526217 |
| P2 cl119 | G09 | 1440100 | sample | 1.0693150 | 0.4045211 |
| P2 cl119 | G10 | 856750 | sample | 0.6361611 | -2.1233567 |
| P2 cl119 | G11 | 1342200 | sample | 0.9966215 | -0.0197169 |
| P2 cl119 | G12 | 345290 | SAL | 0.2563876 | -4.3397078 |
| P2 cl119 | H01 | 911730 | empty | 0.6769853 | -1.8851074 |
| P2 cl119 | H02 | 1160300 | sample | 0.8615556 | -0.8079589 |
| P2 cl119 | H03 | 706480 | sample | 0.5245814 | -2.7745339 |
| P2 cl119 | H04 | 1276000 | sample | 0.9474661 | -0.3065867 |
| P2 cl119 | H05 | 1299400 | sample | 0.9648413 | -0.2051856 |
| P2 cl119 | H06 | 780230 | sample | 0.5793429 | -2.4549470 |
| P2 cl119 | H07 | 1281100 | sample | 0.9512530 | -0.2844865 |
| P2 cl119 | H08 | 1293400 | sample | 0.9603861 | -0.2311859 |
| P2 cl119 | H09 | 1260200 | sample | 0.9357342 | -0.3750541 |
| P2 cl119 | H10 | 1190000 | sample | 0.8836087 | -0.6792575 |
| P2 cl119 | H11 | 996770 | sample | 0.7401299 | -1.5165967 |
| P2 cl119 | H12 | 276130 | SAL | 0.2050343 | -4.6394044 |
| P3 cl119 | A01 | 210680 | SAL | 0.1293746 | -6.1814658 |
| P3 cl119 | A02 | 1017800 | sample | 0.6250115 | -2.6624291 |
| P3 cl119 | A03 | 1424200 | sample | 0.8745740 | -0.8905284 |
| P3 cl119 | A04 | 1399300 | sample | 0.8592834 | -0.9990922 |
| P3 cl119 | A05 | 1198900 | sample | 0.7362216 | -1.8728346 |
| P3 cl119 | A06 | 1250100 | sample | 0.7676625 | -1.6496030 |
| P3 cl119 | A07 | 1301700 | sample | 0.7993491 | -1.4246274 |
| P3 cl119 | A08 | 1241000 | sample | 0.7620744 | -1.6892789 |
| P3 cl119 | A09 | 1160700 | sample | 0.7127637 | -2.0393862 |
| P3 cl119 | A10 | 1178700 | sample | 0.7238171 | -1.9609064 |
| P3 cl119 | A11 | 1186500 | sample | 0.7286070 | -1.9268985 |
| P3 cl119 | A12 | 1309700 | empty | 0.8042617 | -1.3897474 |
| P3 cl119 | B01 | 280620 | SAL | 0.1723234 | -5.8765280 |
| P3 cl119 | B02 | 1366800 | sample | 0.8393257 | -1.1407919 |
| P3 cl119 | B03 | 1692200 | sample | 1.0391477 | 0.2779495 |
| P3 cl119 | B04 | 1550600 | sample | 0.9521938 | -0.3394254 |
| P3 cl119 | B05 | 1062700 | sample | 0.6525837 | -2.4666655 |
| P3 cl119 | B06 | 1756700 | sample | 1.0787559 | 0.5591690 |
| P3 cl119 | B07 | 1498400 | sample | 0.9201388 | -0.5670170 |
| P3 cl119 | B08 | 1558000 | sample | 0.9567380 | -0.3071614 |
| P3 cl119 | B09 | 1476000 | sample | 0.9063834 | -0.6646808 |
| P3 cl119 | B10 | 1687300 | sample | 1.0361387 | 0.2565855 |
| P3 cl119 | B11 | 1557400 | sample | 0.9563696 | -0.3097774 |
| P3 cl119 | B12 | 1467400 | empty | 0.9011023 | -0.7021767 |
| P3 cl119 | C01 | 252530 | SAL | 0.1550738 | -5.9990002 |
| P3 cl119 | C02 | 1557200 | sample | 0.9562467 | -0.3106494 |
| P3 cl119 | C03 | 1763000 | sample | 1.0826246 | 0.5866369 |
| P3 cl119 | C04 | 1786600 | sample | 1.0971169 | 0.6895327 |
| P3 cl119 | C05 | 1828800 | sample | 1.1230311 | 0.8735244 |
| P3 cl119 | C06 | 1846400 | sample | 1.1338389 | 0.9502603 |
| P3 cl119 | C07 | 1703800 | sample | 1.0462710 | 0.3285254 |
| P3 cl119 | C08 | 1686800 | sample | 1.0358316 | 0.2544055 |
| P3 cl119 | C09 | 1743400 | sample | 1.0705886 | 0.5011811 |
| P3 cl119 | C10 | 1766900 | sample | 1.0850195 | 0.6036409 |
| P3 cl119 | C11 | 1728600 | sample | 1.0615002 | 0.4366532 |
| P3 cl119 | C12 | 1479400 | DMSO | 0.9084712 | -0.6498568 |
| P3 cl119 | D01 | 948660 | DMSO | 0.5825540 | -2.9638789 |
| P3 cl119 | D02 | 1498400 | sample | 0.9201388 | -0.5670170 |
| P3 cl119 | D03 | 1896800 | sample | 1.1647886 | 1.1700038 |
| P3 cl119 | D04 | 1829600 | sample | 1.1235224 | 0.8770124 |
| P3 cl119 | D05 | 1834400 | sample | 1.1264700 | 0.8979403 |
| P3 cl119 | D06 | 1598800 | sample | 0.9817925 | -0.1292738 |
| P3 cl119 | D07 | 1725200 | sample | 1.0594123 | 0.4218292 |
| P3 cl119 | D08 | 1654300 | sample | 1.0158740 | 0.1127058 |

*(continued)*

| Plate | Well | Value | Treatment | Norm | Score |
|---|---|---|---|---|---|
| P3 cl119 | D09 | 1658800 | sample | 1.0186374 | 0.1323258 |
| P3 cl119 | D10 | 1701400 | sample | 1.0447972 | 0.3180614 |
| P3 cl119 | D11 | 318990 | sample | 0.1958857 | -5.7092351 |
| P3 cl119 | D12 | 1594100 | DMSO | 0.9789063 | -0.1497657 |
| P3 cl119 | E01 | 899670 | DMSO | 0.5524701 | -3.1774750 |
| P3 cl119 | E02 | 1579400 | sample | 0.9698793 | -0.2138576 |
| P3 cl119 | E03 | 1953000 | sample | 1.1992999 | 1.4150354 |
| P3 cl119 | E04 | 1898100 | sample | 1.1655869 | 1.1756718 |
| P3 cl119 | E05 | 1917500 | sample | 1.1775001 | 1.2602557 |
| P3 cl119 | E06 | 1897200 | sample | 1.1650342 | 1.1717478 |
| P3 cl119 | E07 | 1799500 | sample | 1.1050385 | 0.7457766 |
| P3 cl119 | E08 | 1680100 | sample | 1.0317173 | 0.2251936 |
| P3 cl119 | E09 | 1672800 | sample | 1.0272345 | 0.1933656 |
| P3 cl119 | E10 | 1803900 | sample | 1.1077405 | 0.7649606 |
| P3 cl119 | E11 | 811840 | sample | 0.4985354 | -3.5604131 |
| P3 cl119 | E12 | 1656400 | DMSO | 1.0171636 | 0.1218618 |
| P3 cl119 | F01 | 892440 | DMSO | 0.5480303 | -3.2089977 |
| P3 cl119 | F02 | 1626100 | sample | 0.9985569 | -0.0102460 |
| P3 cl119 | F03 | 1917000 | sample | 1.1771930 | 1.2580757 |
| P3 cl119 | F04 | 1785900 | sample | 1.0966870 | 0.6864807 |
| P3 cl119 | F05 | 1786200 | sample | 1.0968713 | 0.6877887 |
| P3 cl119 | F06 | 1816600 | sample | 1.1155393 | 0.8203325 |
| P3 cl119 | F07 | 1803300 | sample | 1.1073720 | 0.7623446 |
| P3 cl119 | F08 | 660820 | sample | 0.4057969 | -4.2188590 |
| P3 cl119 | F09 | 1635800 | sample | 1.0045135 | 0.0320459 |
| P3 cl119 | F10 | 1853600 | sample | 1.1382603 | 0.9816522 |
| P3 cl119 | F11 | 1782700 | sample | 1.0947220 | 0.6725288 |
| P3 cl119 | F12 | 400000 | SAL | 0.2456323 | -5.3560321 |
| P3 cl119 | G01 | 1062800 | empty | 0.6526452 | -2.4662295 |
| P3 cl119 | G02 | 1494200 | sample | 0.9175596 | -0.5853289 |
| P3 cl119 | G03 | 1088200 | sample | 0.6682428 | -2.3554857 |
| P3 cl119 | G04 | 1536500 | sample | 0.9435353 | -0.4009013 |
| P3 cl119 | G05 | 1777900 | sample | 1.0917744 | 0.6516008 |
| P3 cl119 | G06 | 1630800 | sample | 1.0014431 | 0.0102460 |
| P3 cl119 | G07 | 1585900 | sample | 0.9738709 | -0.1855177 |
| P3 cl119 | G08 | 1755900 | sample | 1.0782646 | 0.5556810 |
| P3 cl119 | G09 | 1665300 | sample | 1.0226289 | 0.1606657 |
| P3 cl119 | G10 | 1570400 | sample | 0.9643526 | -0.2530975 |
| P3 cl119 | G11 | 1612300 | sample | 0.9900826 | -0.0704139 |
| P3 cl119 | G12 | 378600 | SAL | 0.2324910 | -5.4493360 |
| P3 cl119 | H01 | 902670 | empty | 0.5543124 | -3.1643950 |
| P3 cl119 | H02 | 1419400 | sample | 0.8716264 | -0.9114563 |
| P3 cl119 | H03 | 1681700 | sample | 1.0326998 | 0.2321696 |
| P3 cl119 | H04 | 1552900 | sample | 0.9536062 | -0.3293974 |
| P3 cl119 | H05 | 1572000 | sample | 0.9653351 | -0.2461215 |
| P3 cl119 | H06 | 772100 | sample | 0.4741318 | -3.7336791 |
| P3 cl119 | H07 | 973060 | sample | 0.5975375 | -2.8574951 |
| P3 cl119 | H08 | 1473300 | sample | 0.9047254 | -0.6764528 |
| P3 cl119 | H09 | 1472200 | sample | 0.9040499 | -0.6812487 |
| P3 cl119 | H10 | 1548600 | sample | 0.9509656 | -0.3481454 |
| P3 cl119 | H11 | 1439400 | sample | 0.8839080 | -0.8242565 |
| P3 cl119 | H12 | 292860 | SAL | 0.1798397 | -5.8231617 |
| P4 cl119 | A01 | 269000 | SAL | 0.2051165 | -4.9015189 |
| P4 cl119 | A02 | 1147100 | sample | 0.8746807 | -0.7727609 |
| P4 cl119 | A03 | 1151000 | sample | 0.8776545 | -0.7544234 |
| P4 cl119 | A04 | 954800 | sample | 0.7280491 | -1.6769406 |
| P4 cl119 | A05 | 1108300 | sample | 0.8450951 | -0.9551955 |
| P4 cl119 | A06 | 1127500 | sample | 0.8597354 | -0.8649186 |
| P4 cl119 | A07 | 1108500 | sample | 0.8452476 | -0.9542551 |
| P4 cl119 | A08 | 442120 | sample | 0.3371230 | -4.0875221 |
| P4 cl119 | A09 | 1020900 | sample | 0.7784513 | -1.3661435 |
| P4 cl119 | A10 | 1060000 | sample | 0.8082657 | -1.1822984 |

*(continued)*

| Plate | Well | Value | Treatment | Norm | Score |
|---|---|---|---|---|---|
| P4 cl119 | A11 | 921290 | sample | 0.7024972 | -1.8345020 |
| P4 cl119 | A12 | 999220 | empty | 0.7619200 | -1.4680812 |
| P4 cl119 | B01 | 319100 | SAL | 0.2433185 | -4.6659526 |
| P4 cl119 | B02 | 1274700 | sample | 0.9719776 | -0.1727956 |
| P4 cl119 | B03 | 1267600 | sample | 0.9665637 | -0.2061793 |
| P4 cl119 | B04 | 1308200 | sample | 0.9975218 | -0.0152812 |
| P4 cl119 | B05 | 1054800 | sample | 0.8043006 | -1.2067484 |
| P4 cl119 | B06 | 1435400 | sample | 1.0945137 | 0.5828033 |
| P4 cl119 | B07 | 1231800 | sample | 0.9392657 | -0.3745081 |
| P4 cl119 | B08 | 1106400 | sample | 0.8436463 | -0.9641292 |
| P4 cl119 | B09 | 1292100 | sample | 0.9852453 | -0.0909822 |
| P4 cl119 | B10 | 1359100 | sample | 1.0363338 | 0.2240466 |
| P4 cl119 | B11 | 1319600 | sample | 1.0062145 | 0.0383207 |
| P4 cl119 | B12 | 1239000 | empty | 0.9447558 | -0.3406543 |
| P4 cl119 | C01 | 325300 | SAL | 0.2480461 | -4.6368007 |
| P4 cl119 | C02 | 1296100 | sample | 0.9882954 | -0.0721745 |
| P4 cl119 | C03 | 1328900 | sample | 1.0133059 | 0.0820485 |
| P4 cl119 | C04 | 1258100 | sample | 0.9593198 | -0.2508476 |
| P4 cl119 | C05 | 1392700 | sample | 1.0619543 | 0.3820312 |
| P4 cl119 | C06 | 1513800 | sample | 1.1542949 | 0.9514340 |
| P4 cl119 | C07 | 1432600 | sample | 1.0923787 | 0.5696379 |
| P4 cl119 | C08 | 886570 | sample | 0.6760227 | -1.9977528 |
| P4 cl119 | C09 | 1464600 | sample | 1.1167791 | 0.7200994 |
| P4 cl119 | C10 | 1335200 | sample | 1.0181097 | 0.1116707 |
| P4 cl119 | C11 | 1221400 | sample | 0.9313355 | -0.4234081 |
| P4 cl119 | C12 | 1297300 | DMSO | 0.9892104 | -0.0665322 |
| P4 cl119 | D01 | 1078600 | DMSO | 0.8224484 | -1.0948426 |
| P4 cl119 | D02 | 1234300 | sample | 0.9411720 | -0.3627533 |
| P4 cl119 | D03 | 1466600 | sample | 1.1183042 | 0.7295033 |
| P4 cl119 | D04 | 1385400 | sample | 1.0563880 | 0.3477072 |
| P4 cl119 | D05 | 1427700 | sample | 1.0886423 | 0.5465985 |
| P4 cl119 | D06 | 1503900 | sample | 1.1467460 | 0.9048850 |
| P4 cl119 | D07 | 1481900 | sample | 1.1299706 | 0.8014427 |
| P4 cl119 | D08 | 1457800 | sample | 1.1115940 | 0.6881263 |
| P4 cl119 | D09 | 1343200 | sample | 1.0242098 | 0.1492860 |
| P4 cl119 | D10 | 1444900 | sample | 1.1017576 | 0.6274715 |
| P4 cl119 | D11 | 1427000 | sample | 1.0881086 | 0.5433071 |
| P4 cl119 | D12 | 1268000 | DMSO | 0.9668687 | -0.2042985 |
| P4 cl119 | E01 | 1013500 | DMSO | 0.7728087 | -1.4009378 |
| P4 cl119 | E02 | 1290000 | sample | 0.9836441 | -0.1008562 |
| P4 cl119 | E03 | 1511600 | sample | 1.1526173 | 0.9410898 |
| P4 cl119 | E04 | 778280 | sample | 0.5934500 | -2.5069239 |
| P4 cl119 | E05 | 1499500 | sample | 1.1433909 | 0.8841965 |
| P4 cl119 | E06 | 1567600 | sample | 1.1953182 | 1.2043974 |
| P4 cl119 | E07 | 1011500 | sample | 0.7712837 | -1.4103416 |
| P4 cl119 | E08 | 1314700 | sample | 1.0024782 | 0.0152812 |
| P4 cl119 | E09 | 1412300 | sample | 1.0768996 | 0.4741889 |
| P4 cl119 | E10 | 1558700 | sample | 1.1885318 | 1.1625503 |
| P4 cl119 | E11 | 1545800 | sample | 1.1786953 | 1.1018955 |
| P4 cl119 | E12 | 1521300 | DMSO | 1.1600137 | 0.9866984 |
| P4 cl119 | F01 | 839090 | DMSO | 0.6398185 | -2.2210000 |
| P4 cl119 | F02 | 1286000 | sample | 0.9805940 | -0.1196639 |
| P4 cl119 | F03 | 1445000 | sample | 1.1018338 | 0.6279417 |
| P4 cl119 | F04 | 1401500 | sample | 1.0686645 | 0.4234081 |
| P4 cl119 | F05 | 1293400 | sample | 0.9862366 | -0.0848697 |
| P4 cl119 | F06 | 1149500 | sample | 0.8765107 | -0.7614763 |
| P4 cl119 | F07 | 1530300 | sample | 1.1668764 | 1.0290157 |
| P4 cl119 | F08 | 892770 | sample | 0.6807503 | -1.9686008 |
| P4 cl119 | F09 | 1465300 | sample | 1.1173129 | 0.7233908 |
| P4 cl119 | F10 | 1405200 | sample | 1.0714858 | 0.4408052 |
| P4 cl119 | F11 | 1452000 | sample | 1.1071715 | 0.6608552 |
| P4 cl119 | F12 | 401060 | SAL | 0.3058142 | -4.2805831 |

*(continued)*

| Plate | Well | Value | Treatment | Norm | Score |
|---|---|---|---|---|---|
| P4 cl119 | G01 | 927430 | empty | 0.7071791 | -1.8056322 |
| P4 cl119 | G02 | 1337900 | sample | 1.0201685 | 0.1243658 |
| P4 cl119 | G03 | 1466500 | sample | 1.1182279 | 0.7290331 |
| P4 cl119 | G04 | 1392200 | sample | 1.0615731 | 0.3796802 |
| P4 cl119 | G05 | 1265200 | sample | 0.9647337 | -0.2174639 |
| P4 cl119 | G06 | 1410900 | sample | 1.0758321 | 0.4676062 |
| P4 cl119 | G07 | 781340 | sample | 0.5957833 | -2.4925361 |
| P4 cl119 | G08 | 1493500 | sample | 1.1388158 | 0.8559850 |
| P4 cl119 | G09 | 790110 | sample | 0.6024705 | -2.4513002 |
| P4 cl119 | G10 | 664060 | sample | 0.5063556 | -3.0439775 |
| P4 cl119 | G11 | 1436800 | sample | 1.0955812 | 0.5893860 |
| P4 cl119 | G12 | 364710 | SAL | 0.2780968 | -4.4514980 |
| P4 cl119 | H01 | 971800 | empty | 0.7410119 | -1.5970079 |
| P4 cl119 | H02 | 919960 | sample | 0.7014831 | -1.8407556 |
| P4 cl119 | H03 | 1172600 | sample | 0.8941248 | -0.6528619 |
| P4 cl119 | H04 | 338140 | sample | 0.2578367 | -4.5764280 |
| P4 cl119 | H05 | 1315500 | sample | 1.0030882 | 0.0190428 |
| P4 cl119 | H06 | 1339000 | sample | 1.0210073 | 0.1295380 |
| P4 cl119 | H07 | 1363100 | sample | 1.0393839 | 0.2428543 |
| P4 cl119 | H08 | 1270300 | sample | 0.9686225 | -0.1934841 |
| P4 cl119 | H09 | 63170 | sample | 0.0481681 | -5.8693156 |
| P4 cl119 | H10 | 657500 | sample | 0.5013535 | -3.0748221 |
| P4 cl119 | H11 | 297450 | sample | 0.2268100 | -4.7677493 |
| P4 cl119 | H12 | 313010 | SAL | 0.2386747 | -4.6945873 |
| P5 cl119 | A01 | 274460 | SAL | 0.2196383 | -4.5930371 |
| P5 cl119 | A02 | 1066900 | sample | 0.8537932 | -0.8605409 |
| P5 cl119 | A03 | 1085000 | sample | 0.8682778 | -0.7752876 |
| P5 cl119 | A04 | 1057500 | sample | 0.8462708 | -0.9048162 |
| P5 cl119 | A05 | 1102500 | sample | 0.8822823 | -0.6928603 |
| P5 cl119 | A06 | 1217400 | sample | 0.9742318 | -0.1516662 |
| P5 cl119 | A07 | 1047000 | sample | 0.8378681 | -0.9542725 |
| P5 cl119 | A08 | 1004500 | sample | 0.8038572 | -1.1544531 |
| P5 cl119 | A09 | 913510 | sample | 0.7310419 | -1.5830279 |
| P5 cl119 | A10 | 970800 | sample | 0.7768886 | -1.3131845 |
| P5 cl119 | A11 | 894780 | sample | 0.7160531 | -1.6712487 |
| P5 cl119 | A12 | 857390 | empty | 0.6861316 | -1.8473605 |
| P5 cl119 | B01 | 374410 | SAL | 0.2996239 | -4.1222596 |
| P5 cl119 | B02 | 1239900 | sample | 0.9922375 | -0.0456883 |
| P5 cl119 | B03 | 1297100 | sample | 1.0380122 | 0.2237312 |
| P5 cl119 | B04 | 1226600 | sample | 0.9815941 | -0.1083330 |
| P5 cl119 | B05 | 1270400 | sample | 1.0166453 | 0.0979707 |
| P5 cl119 | B06 | 1215600 | sample | 0.9727913 | -0.1601445 |
| P5 cl119 | B07 | 1163800 | sample | 0.9313380 | -0.4041292 |
| P5 cl119 | B08 | 1168100 | sample | 0.9347791 | -0.3838757 |
| P5 cl119 | B09 | 1157800 | sample | 0.9265365 | -0.4323900 |
| P5 cl119 | B10 | 1380100 | sample | 1.1044334 | 0.6146721 |
| P5 cl119 | B11 | 1240800 | sample | 0.9929577 | -0.0414492 |
| P5 cl119 | B12 | 1190000 | empty | 0.9523047 | -0.2807238 |
| P5 cl119 | C01 | 343140 | SAL | 0.2745999 | -4.2695453 |
| P5 cl119 | C02 | 1250400 | sample | 1.0006402 | 0.0037681 |
| P5 cl119 | C03 | 1334900 | sample | 1.0682618 | 0.4017742 |
| P5 cl119 | C04 | 1218700 | sample | 0.9752721 | -0.1455430 |
| P5 cl119 | C05 | 1305600 | sample | 1.0448143 | 0.2637673 |
| P5 cl119 | C06 | 1338500 | sample | 1.0711428 | 0.4187306 |
| P5 cl119 | C07 | 1136700 | sample | 0.9096511 | -0.5317738 |
| P5 cl119 | C08 | 1373500 | sample | 1.0991517 | 0.5835852 |
| P5 cl119 | C09 | 1042700 | sample | 0.8344270 | -0.9745261 |
| P5 cl119 | C10 | 1353700 | sample | 1.0833067 | 0.4903246 |
| P5 cl119 | C11 | 1343600 | sample | 1.0752241 | 0.4427523 |
| P5 cl119 | C12 | 1500600 | DMSO | 1.2008643 | 1.1822429 |
| P5 cl119 | D01 | 1012000 | DMSO | 0.8098592 | -1.1191271 |

*(continued)*

| Plate | Well | Value | Treatment | Norm | Score |
|---|---|---|---|---|---|
| P5 cl119 | D02 | 1385700 | sample | 1.1089149 | 0.6410488 |
| P5 cl119 | D03 | 1346100 | sample | 1.0772247 | 0.4545276 |
| P5 cl119 | D04 | 1248800 | sample | 0.9993598 | -0.0037681 |
| P5 cl119 | D05 | 1356300 | sample | 1.0853873 | 0.5025710 |
| P5 cl119 | D06 | 671630 | sample | 0.5374760 | -2.7223144 |
| P5 cl119 | D07 | 1473900 | sample | 1.1794974 | 1.0564824 |
| P5 cl119 | D08 | 1442800 | sample | 1.1546095 | 0.9099973 |
| P5 cl119 | D09 | 1396100 | sample | 1.1172375 | 0.6900342 |
| P5 cl119 | D10 | 1399700 | sample | 1.1201184 | 0.7069907 |
| P5 cl119 | D11 | 1517900 | sample | 1.2147087 | 1.2637281 |
| P5 cl119 | D12 | 1393600 | DMSO | 1.1152369 | 0.6782589 |
| P5 cl119 | E01 | 1007100 | DMSO | 0.8059379 | -1.1422068 |
| P5 cl119 | E02 | 1320500 | sample | 1.0567382 | 0.3339483 |
| P5 cl119 | E03 | 1362200 | sample | 1.0901088 | 0.5303608 |
| P5 cl119 | E04 | 516210 | sample | 0.4131002 | -3.4543630 |
| P5 cl119 | E05 | 1138700 | sample | 0.9112516 | -0.5223535 |
| P5 cl119 | E06 | 1541900 | sample | 1.2339149 | 1.3767713 |
| P5 cl119 | E07 | 1464000 | sample | 1.1715749 | 1.0098521 |
| P5 cl119 | E08 | 1489200 | sample | 1.1917414 | 1.1285474 |
| P5 cl119 | E09 | 1328100 | sample | 1.0628201 | 0.3697453 |
| P5 cl119 | E10 | 1603800 | sample | 1.2834507 | 1.6683284 |
| P5 cl119 | E11 | 1523000 | sample | 1.2187900 | 1.2877498 |
| P5 cl119 | E12 | 1498700 | DMSO | 1.1993438 | 1.1732936 |
| P5 cl119 | F01 | 1006800 | DMSO | 0.8056978 | -1.1436198 |
| P5 cl119 | F02 | 1389500 | sample | 1.1119558 | 0.6589473 |
| P5 cl119 | F03 | 1328800 | sample | 1.0633803 | 0.3730424 |
| P5 cl119 | F04 | 128190 | sample | 0.1025848 | -5.2819880 |
| P5 cl119 | F05 | 1451200 | sample | 1.1613316 | 0.9495624 |
| P5 cl119 | F06 | 976040 | sample | 0.7810819 | -1.2885034 |
| P5 cl119 | F07 | 841010 | sample | 0.6730234 | -1.9245124 |
| P5 cl119 | F08 | 1420700 | sample | 1.1369238 | 0.8059034 |
| P5 cl119 | F09 | 1498700 | sample | 1.1993438 | 1.1732936 |
| P5 cl119 | F10 | 1625500 | sample | 1.3008163 | 1.7705382 |
| P5 cl119 | F11 | 1473700 | sample | 1.1793374 | 1.0555404 |
| P5 cl119 | F12 | 410120 | SAL | 0.3282010 | -3.9540608 |
| P5 cl119 | G01 | 990750 | empty | 0.7928537 | -1.2192174 |
| P5 cl119 | G02 | 1158300 | sample | 0.9269366 | -0.4300350 |
| P5 cl119 | G03 | 1186600 | sample | 0.9495839 | -0.2967383 |
| P5 cl119 | G04 | 1209400 | sample | 0.9678297 | -0.1893473 |
| P5 cl119 | G05 | 1479500 | sample | 1.1839789 | 1.0828591 |
| P5 cl119 | G06 | 138770 | sample | 0.1110515 | -5.2321548 |
| P5 cl119 | G07 | 674400 | sample | 0.5396927 | -2.7092674 |
| P5 cl119 | G08 | 1046800 | sample | 0.8377081 | -0.9552146 |
| P5 cl119 | G09 | 1389400 | sample | 1.1118758 | 0.6584763 |
| P5 cl119 | G10 | 1098100 | sample | 0.8787612 | -0.7135848 |
| P5 cl119 | G11 | 1459700 | sample | 1.1681338 | 0.9895985 |
| P5 cl119 | G12 | 403170 | SAL | 0.3226392 | -3.9867962 |
| P5 cl119 | H01 | 912400 | empty | 0.7301536 | -1.5882562 |
| P5 cl119 | H02 | 1168700 | sample | 0.9352593 | -0.3810496 |
| P5 cl119 | H03 | 1122500 | sample | 0.8982875 | -0.5986577 |
| P5 cl119 | H04 | 1223300 | sample | 0.9789533 | -0.1238764 |
| P5 cl119 | H05 | 1370000 | sample | 1.0963508 | 0.5670998 |
| P5 cl119 | H06 | 1292200 | sample | 1.0340909 | 0.2006516 |
| P5 cl119 | H07 | 601110 | sample | 0.4810419 | -3.0544729 |
| P5 cl119 | H08 | 126250 | sample | 0.1010323 | -5.2911257 |
| P5 cl119 | H09 | 1310400 | sample | 1.0486556 | 0.2863760 |
| P5 cl119 | H10 | 386800 | sample | 0.3095391 | -4.0639010 |
| P5 cl119 | H11 | 1437200 | sample | 1.1501280 | 0.8836206 |
| P5 cl119 | H12 | 358510 | SAL | 0.2868998 | -4.1971506 |
| P6 cl119 | A01 | 213740 | SAL | 0.1520199 | -5.8463712 |
| P6 cl119 | A02 | 1166100 | sample | 0.8293741 | -1.1763747 |
| P6 cl119 | A03 | 895750 | sample | 0.6370910 | -2.5020640 |

*(continued)*

| Plate | Well | Value | Treatment | Norm | Score |
|---|---|---|---|---|---|
| P6 cl119 | A04 | 883290 | sample | 0.6282290 | -2.5631630 |
| P6 cl119 | A05 | 734660 | sample | 0.5225178 | -3.2919857 |
| P6 cl119 | A06 | 1267500 | sample | 0.9014936 | -0.6791492 |
| P6 cl119 | A07 | 1240500 | sample | 0.8822902 | -0.8115465 |
| P6 cl119 | A08 | 1167500 | sample | 0.8303698 | -1.1695096 |
| P6 cl119 | A09 | 1047900 | sample | 0.7453058 | -1.7559807 |
| P6 cl119 | A10 | 603040 | sample | 0.4289047 | -3.9373980 |
| P6 cl119 | A11 | 858740 | sample | 0.6107681 | -2.6835464 |
| P6 cl119 | A12 | 979830 | empty | 0.6968919 | -2.0897690 |
| P6 cl119 | B01 | 971380 | SAL | 0.6908819 | -2.1312045 |
| P6 cl119 | B02 | 1318700 | sample | 0.9379090 | -0.4280846 |
| P6 cl119 | B03 | 1364100 | sample | 0.9701991 | -0.2054610 |
| P6 cl119 | B04 | 1327600 | sample | 0.9442390 | -0.3844426 |
| P6 cl119 | B05 | 925250 | sample | 0.6580725 | -2.3574077 |
| P6 cl119 | B06 | 1493200 | sample | 1.0620199 | 0.4275943 |
| P6 cl119 | B07 | 1508200 | sample | 1.0726885 | 0.5011484 |
| P6 cl119 | B08 | 1466600 | sample | 1.0431010 | 0.2971584 |
| P6 cl119 | B09 | 1488900 | sample | 1.0589616 | 0.4065088 |
| P6 cl119 | B10 | 1538900 | sample | 1.0945235 | 0.6516890 |
| P6 cl119 | B11 | 1590700 | sample | 1.1313656 | 0.9056957 |
| P6 cl119 | B12 | 1350600 | empty | 0.9605974 | -0.2716597 |
| P6 cl119 | C01 | 973120 | SAL | 0.6921195 | -2.1226722 |
| P6 cl119 | C02 | 1165600 | sample | 0.8290185 | -1.1788265 |
| P6 cl119 | C03 | 1275200 | sample | 0.9069701 | -0.6413914 |
| P6 cl119 | C04 | 1269400 | sample | 0.9028450 | -0.6698323 |
| P6 cl119 | C05 | 1345000 | sample | 0.9566145 | -0.2991199 |
| P6 cl119 | C06 | 1336600 | sample | 0.9506401 | -0.3403101 |
| P6 cl119 | C07 | 1553600 | sample | 1.1049787 | 0.7237720 |
| P6 cl119 | C08 | 1574600 | sample | 1.1199147 | 0.8267477 |
| P6 cl119 | C09 | 1484300 | sample | 1.0556899 | 0.3839522 |
| P6 cl119 | C10 | 1680400 | sample | 1.1951636 | 1.3455490 |
| P6 cl119 | C11 | 1697500 | sample | 1.2073257 | 1.4294006 |
| P6 cl119 | C12 | 1619700 | DMSO | 1.1519915 | 1.0479002 |
| P6 cl119 | D01 | 907830 | DMSO | 0.6456828 | -2.4428285 |
| P6 cl119 | D02 | 1400500 | sample | 0.9960882 | -0.0269698 |
| P6 cl119 | D03 | 1366100 | sample | 0.9716216 | -0.1956538 |
| P6 cl119 | D04 | 1117300 | sample | 0.7946657 | -1.4156705 |
| P6 cl119 | D05 | 1611000 | sample | 1.1458037 | 1.0052389 |
| P6 cl119 | D06 | 1554600 | sample | 1.1056899 | 0.7286756 |
| P6 cl119 | D07 | 1661800 | sample | 1.1819346 | 1.2543420 |
| P6 cl119 | D08 | 1508400 | sample | 1.0728307 | 0.5021291 |
| P6 cl119 | D09 | 249070 | sample | 0.1771479 | -5.6731268 |
| P6 cl119 | D10 | 1627200 | sample | 1.1573257 | 1.0846773 |
| P6 cl119 | D11 | 1500300 | sample | 1.0670697 | 0.4624099 |
| P6 cl119 | D12 | 1547500 | DMSO | 1.1006401 | 0.6938600 |
| P6 cl119 | E01 | 853540 | DMSO | 0.6070697 | -2.7090452 |
| P6 cl119 | E02 | 1270600 | sample | 0.9036984 | -0.6639480 |
| P6 cl119 | E03 | 1390400 | sample | 0.9889047 | -0.0764962 |
| P6 cl119 | E04 | 1064500 | sample | 0.7571124 | -1.6745808 |
| P6 cl119 | E05 | 1499400 | sample | 1.0664296 | 0.4579966 |
| P6 cl119 | E06 | 1605200 | sample | 1.1416785 | 0.9767980 |
| P6 cl119 | E07 | 1730600 | sample | 1.2308677 | 1.5917099 |
| P6 cl119 | E08 | 1614600 | sample | 1.1483642 | 1.0228918 |
| P6 cl119 | E09 | 766240 | sample | 0.5449787 | -3.1371298 |
| P6 cl119 | E10 | 1607800 | sample | 1.1435277 | 0.9895473 |
| P6 cl119 | E11 | 1669300 | sample | 1.1872688 | 1.2911190 |
| P6 cl119 | E12 | 1545700 | DMSO | 1.0993599 | 0.6850335 |
| P6 cl119 | F01 | 879830 | DMSO | 0.6257681 | -2.5801294 |
| P6 cl119 | F02 | 1381700 | sample | 0.9827169 | -0.1191576 |
| P6 cl119 | F03 | 1335800 | sample | 0.9500711 | -0.3442330 |
| P6 cl119 | F04 | 1230900 | sample | 0.8754623 | -0.8586211 |
| P6 cl119 | F05 | 1461400 | sample | 1.0394026 | 0.2716597 |

*(continued)*

| Plate | Well | Value | Treatment | Norm | Score |
|---|---|---|---|---|---|
| P6 cl119 | F06 | 1495800 | sample | 1.0638691 | 0.4403437 |
| P6 cl119 | F07 | 1528200 | sample | 1.0869132 | 0.5992204 |
| P6 cl119 | F08 | 1628300 | sample | 1.1581081 | 1.0900712 |
| P6 cl119 | F09 | 1538000 | sample | 1.0938834 | 0.6472758 |
| P6 cl119 | F10 | 1627100 | sample | 1.1572546 | 1.0841869 |
| P6 cl119 | F11 | 1580700 | sample | 1.1242532 | 0.8566597 |
| P6 cl119 | F12 | 435510 | SAL | 0.3097511 | -4.7588989 |
| P6 cl119 | G01 | 847410 | empty | 0.6027098 | -2.7391043 |
| P6 cl119 | G02 | 1362100 | sample | 0.9687767 | -0.2152682 |
| P6 cl119 | G03 | 1500800 | sample | 1.0674253 | 0.4648617 |
| P6 cl119 | G04 | 1292600 | sample | 0.9193457 | -0.5560687 |
| P6 cl119 | G05 | 1413600 | sample | 1.0054054 | 0.0372674 |
| P6 cl119 | G06 | 1447100 | sample | 1.0292319 | 0.2015381 |
| P6 cl119 | G07 | 1607000 | sample | 1.1429587 | 0.9856244 |
| P6 cl119 | G08 | 1480300 | sample | 1.0528450 | 0.3643378 |
| P6 cl119 | G09 | 1594500 | sample | 1.1340683 | 0.9243294 |
| P6 cl119 | G10 | 1656700 | sample | 1.1783073 | 1.2293336 |
| P6 cl119 | G11 | 1535100 | sample | 1.0918208 | 0.6330553 |
| P6 cl119 | G12 | 408800 | SAL | 0.2907539 | -4.8898741 |
| P6 cl119 | H01 | 736420 | empty | 0.5237696 | -3.2833553 |
| P6 cl119 | H02 | 1385100 | sample | 0.9851351 | -0.1024853 |
| P6 cl119 | H03 | 1279200 | sample | 0.9098151 | -0.6217770 |
| P6 cl119 | H04 | 1301400 | sample | 0.9256046 | -0.5129170 |
| P6 cl119 | H05 | 1473300 | sample | 1.0478663 | 0.3300126 |
| P6 cl119 | H06 | 1411500 | sample | 1.0039118 | 0.0269698 |
| P6 cl119 | H07 | 1193000 | sample | 0.8485064 | -1.0444677 |
| P6 cl119 | H08 | 1134600 | sample | 0.8069701 | -1.3308382 |
| P6 cl119 | H09 | 553370 | sample | 0.3935775 | -4.1809601 |
| P6 cl119 | H10 | 1366200 | sample | 0.9716927 | -0.1951634 |
| P6 cl119 | H11 | 1361800 | sample | 0.9685633 | -0.2167393 |
| P6 cl119 | H12 | 343120 | SAL | 0.2440398 | -5.2119428 |
| P7 cl119 | A01 | 274140 | SAL | 0.2019076 | -5.6504440 |
| P7 cl119 | A02 | 1133600 | sample | 0.8349107 | -1.1688218 |
| P7 cl119 | A03 | 1142000 | sample | 0.8410974 | -1.1250203 |
| P7 cl119 | A04 | 1132800 | sample | 0.8343215 | -1.1729934 |
| P7 cl119 | A05 | 1064800 | sample | 0.7842386 | -1.5275769 |
| P7 cl119 | A06 | 1019900 | sample | 0.7511692 | -1.7617062 |
| P7 cl119 | A07 | 950140 | sample | 0.6997901 | -2.1254672 |
| P7 cl119 | A08 | 136090 | sample | 0.1002320 | -6.3703006 |
| P7 cl119 | A09 | 951050 | sample | 0.7004603 | -2.1207220 |
| P7 cl119 | A10 | 1046700 | sample | 0.7709078 | -1.6219586 |
| P7 cl119 | A11 | 996660 | sample | 0.7340527 | -1.8828904 |
| P7 cl119 | A12 | 991320 | empty | 0.7301197 | -1.9107356 |
| P7 cl119 | B01 | 323190 | SAL | 0.2380335 | -5.3946746 |
| P7 cl119 | B02 | 891440 | sample | 0.6565568 | -2.4315561 |
| P7 cl119 | B03 | 1374600 | sample | 1.0124102 | 0.0878637 |
| P7 cl119 | B04 | 1356900 | sample | 0.9993740 | -0.0044323 |
| P7 cl119 | B05 | 1348600 | sample | 0.9932609 | -0.0477123 |
| P7 cl119 | B06 | 1185800 | sample | 0.8733567 | -0.8966269 |
| P7 cl119 | B07 | 1234300 | sample | 0.9090775 | -0.6437254 |
| P7 cl119 | B08 | 1271800 | sample | 0.9366967 | -0.4481831 |
| P7 cl119 | B09 | 1073200 | sample | 0.7904253 | -1.4837754 |
| P7 cl119 | B10 | 1115700 | sample | 0.8217271 | -1.2621607 |
| P7 cl119 | B11 | 1429400 | sample | 1.0527711 | 0.3736163 |
| P7 cl119 | B12 | 1209100 | empty | 0.8905174 | -0.7751299 |
| P7 cl119 | C01 | 344110 | SAL | 0.2534414 | -5.2855880 |
| P7 cl119 | C02 | 1304900 | sample | 0.9610753 | -0.2755844 |
| P7 cl119 | C03 | 1267300 | sample | 0.9333824 | -0.4716482 |
| P7 cl119 | C04 | 1380500 | sample | 1.0167557 | 0.1186290 |
| P7 cl119 | C05 | 1283900 | sample | 0.9456085 | -0.3850881 |
| P7 cl119 | C06 | 1502600 | sample | 1.1066839 | 0.7553149 |

*(continued)*

| Plate | Well | Value | Treatment | Norm | Score |
|---|---|---|---|---|---|
| P7 cl119 | C07 | 1420200 | sample | 1.0459952 | 0.3256432 |
| P7 cl119 | C08 | 1396500 | sample | 1.0285399 | 0.2020604 |
| P7 cl119 | C09 | 1446000 | sample | 1.0649972 | 0.4601763 |
| P7 cl119 | C10 | 1118000 | sample | 0.8234211 | -1.2501674 |
| P7 cl119 | C11 | 1525400 | sample | 1.1234763 | 0.8742047 |
| P7 cl119 | C12 | 1476500 | DMSO | 1.0874609 | 0.6192175 |
| P7 cl119 | D01 | 1247600 | DMSO | 0.9188731 | -0.5743731 |
| P7 cl119 | D02 | 1055400 | sample | 0.7773154 | -1.5765928 |
| P7 cl119 | D03 | 1382500 | sample | 1.0182287 | 0.1290580 |
| P7 cl119 | D04 | 1154100 | sample | 0.8500092 | -1.0619253 |
| P7 cl119 | D05 | 1510600 | sample | 1.1125760 | 0.7970306 |
| P7 cl119 | D06 | 1499900 | sample | 1.1046953 | 0.7412359 |
| P7 cl119 | D07 | 755670 | sample | 0.5565605 | -3.1395237 |
| P7 cl119 | D08 | 1460100 | sample | 1.0753821 | 0.5337003 |
| P7 cl119 | D09 | 1426000 | sample | 1.0502670 | 0.3558871 |
| P7 cl119 | D10 | 1517000 | sample | 1.1172896 | 0.8304032 |
| P7 cl119 | D11 | 1499000 | sample | 1.1040324 | 0.7365429 |
| P7 cl119 | D12 | 1374300 | DMSO | 1.0121893 | 0.0862994 |
| P7 cl119 | E01 | 1170800 | DMSO | 0.8623090 | -0.9748438 |
| P7 cl119 | E02 | 1409800 | sample | 1.0383355 | 0.2714128 |
| P7 cl119 | E03 | 252610 | sample | 0.1860505 | -5.7627114 |
| P7 cl119 | E04 | 1423100 | sample | 1.0481311 | 0.3407651 |
| P7 cl119 | E05 | 1507900 | sample | 1.1105874 | 0.7829516 |
| P7 cl119 | E06 | 1570500 | sample | 1.1566931 | 1.1093770 |
| P7 cl119 | E07 | 1604700 | sample | 1.1818818 | 1.2877116 |
| P7 cl119 | E08 | 1484400 | sample | 1.0932793 | 0.6604117 |
| P7 cl119 | E09 | 1594900 | sample | 1.1746640 | 1.2366098 |
| P7 cl119 | E10 | 1638900 | sample | 1.2070705 | 1.4660462 |
| P7 cl119 | E11 | 1567100 | sample | 1.1541889 | 1.0916478 |
| P7 cl119 | E12 | 1542900 | DMSO | 1.1363653 | 0.9654578 |
| P7 cl119 | F01 | 1113900 | DMSO | 0.8204014 | -1.2715467 |
| P7 cl119 | F02 | 1252500 | sample | 0.9224820 | -0.5488222 |
| P7 cl119 | F03 | 1312600 | sample | 0.9667465 | -0.2354330 |
| P7 cl119 | F04 | 1433700 | sample | 1.0559381 | 0.3960384 |
| P7 cl119 | F05 | 1450100 | sample | 1.0680169 | 0.4815556 |
| P7 cl119 | F06 | 1472300 | sample | 1.0843675 | 0.5973167 |
| P7 cl119 | F07 | 1461200 | sample | 1.0761922 | 0.5394362 |
| P7 cl119 | F08 | 1429400 | sample | 1.0527711 | 0.3736163 |
| P7 cl119 | F09 | 1489800 | sample | 1.0972565 | 0.6885698 |
| P7 cl119 | F10 | 1292600 | sample | 0.9520162 | -0.3397222 |
| P7 cl119 | F11 | 1547000 | sample | 1.1393850 | 0.9868371 |
| P7 cl119 | F12 | 375940 | SAL | 0.2768846 | -5.1196117 |
| P7 cl119 | G01 | 1108700 | empty | 0.8165715 | -1.2986620 |
| P7 cl119 | G02 | 151550 | sample | 0.1116185 | -6.2896850 |
| P7 cl119 | G03 | 1278000 | sample | 0.9412631 | -0.4158534 |
| P7 cl119 | G04 | 1435100 | sample | 1.0569693 | 0.4033387 |
| P7 cl119 | G05 | 1424200 | sample | 1.0489413 | 0.3465011 |
| P7 cl119 | G06 | 1412600 | sample | 1.0403977 | 0.2860133 |
| P7 cl119 | G07 | 1391500 | sample | 1.0248573 | 0.1759881 |
| P7 cl119 | G08 | 1441600 | sample | 1.0617566 | 0.4372327 |
| P7 cl119 | G09 | 1366500 | sample | 1.0064445 | 0.0456265 |
| P7 cl119 | G10 | 1524400 | sample | 1.1227398 | 0.8689902 |
| P7 cl119 | G11 | 1478500 | sample | 1.0889339 | 0.6296464 |
| P7 cl119 | G12 | 404660 | SAL | 0.2980372 | -4.9698523 |
| P7 cl119 | H01 | 1020300 | empty | 0.7514638 | -1.7596205 |
| P7 cl119 | H02 | 1071800 | sample | 0.7893942 | -1.4910756 |
| P7 cl119 | H03 | 1172800 | sample | 0.8637820 | -0.9644149 |
| P7 cl119 | H04 | 1213000 | sample | 0.8933898 | -0.7547935 |
| P7 cl119 | H05 | 1355700 | sample | 0.9984901 | -0.0106896 |
| P7 cl119 | H06 | 1134600 | sample | 0.8356472 | -1.1636074 |
| P7 cl119 | H07 | 1310700 | sample | 0.9653471 | -0.2453405 |
| P7 cl119 | H08 | 1241600 | sample | 0.9144541 | -0.6056599 |

*(continued)*

| Plate | Well | Value | Treatment | Norm | Score |
|---|---|---|---|---|---|
| P7 cl119 | H09 | 1358600 | sample | 1.0006260 | 0.0044323 |
| P7 cl119 | H10 | 1325300 | sample | 0.9761002 | -0.1692093 |
| P7 cl119 | H11 | 996040 | sample | 0.7335960 | -1.8861233 |
| P7 cl119 | H12 | 325590 | SAL | 0.2398011 | -5.3821599 |
| P8 cl119 | A01 | 260080 | SAL | 0.1942418 | -4.9637643 |
| P8 cl119 | A02 | 1105100 | sample | 0.8253482 | -1.0759186 |
| P8 cl119 | A03 | 882870 | sample | 0.6593749 | -2.0983748 |
| P8 cl119 | A04 | 1028600 | sample | 0.7682139 | -1.4278868 |
| P8 cl119 | A05 | 1065200 | sample | 0.7955488 | -1.2594942 |
| P8 cl119 | A06 | 1163200 | sample | 0.8687404 | -0.8086068 |
| P8 cl119 | A07 | 1099100 | sample | 0.8208671 | -1.1035239 |
| P8 cl119 | A08 | 798900 | sample | 0.5966616 | -2.4847117 |
| P8 cl119 | A09 | 1037700 | sample | 0.7750103 | -1.3860187 |
| P8 cl119 | A10 | 1100800 | sample | 0.8221367 | -1.0957024 |
| P8 cl119 | A11 | 1049300 | sample | 0.7836738 | -1.3326484 |
| P8 cl119 | A12 | 961190 | empty | 0.7178685 | -1.7380329 |
| P8 cl119 | B01 | 297770 | SAL | 0.2223907 | -4.7903567 |
| P8 cl119 | B02 | 1106900 | sample | 0.8266926 | -1.0676370 |
| P8 cl119 | B03 | 1178700 | sample | 0.8803167 | -0.7372929 |
| P8 cl119 | B04 | 1289500 | sample | 0.9630681 | -0.2275141 |
| P8 cl119 | B05 | 1261300 | sample | 0.9420068 | -0.3572593 |
| P8 cl119 | B06 | 1286400 | sample | 0.9607528 | -0.2417769 |
| P8 cl119 | B07 | 1137300 | sample | 0.8493969 | -0.9277699 |
| P8 cl119 | B08 | 1092900 | sample | 0.8162366 | -1.1320495 |
| P8 cl119 | B09 | 1315300 | sample | 0.9823369 | -0.1088111 |
| P8 cl119 | B10 | 1329700 | sample | 0.9930916 | -0.0425583 |
| P8 cl119 | B11 | 1473500 | sample | 1.1004892 | 0.6190500 |
| P8 cl119 | B12 | 1205900 | empty | 0.9006311 | -0.6121487 |
| P8 cl119 | C01 | 313470 | SAL | 0.2341163 | -4.7181227 |
| P8 cl119 | C02 | 1241300 | sample | 0.9270697 | -0.4492771 |
| P8 cl119 | C03 | 1316800 | sample | 0.9834572 | -0.1019098 |
| P8 cl119 | C04 | 1094700 | sample | 0.8175809 | -1.1237679 |
| P8 cl119 | C05 | 1348200 | sample | 1.0069084 | 0.0425583 |
| P8 cl119 | C06 | 1364000 | sample | 1.0187087 | 0.1152523 |
| P8 cl119 | C07 | 1167000 | sample | 0.8715785 | -0.7911234 |
| P8 cl119 | C08 | 1320500 | sample | 0.9862205 | -0.0848865 |
| P8 cl119 | C09 | 1381100 | sample | 1.0314799 | 0.1939276 |
| P8 cl119 | C10 | 796940 | sample | 0.5951977 | -2.4937294 |
| P8 cl119 | C11 | 1566000 | sample | 1.1695732 | 1.0446325 |
| P8 cl119 | C12 | 1382200 | DMSO | 1.0323014 | 0.1989886 |
| P8 cl119 | D01 | 1070300 | DMSO | 0.7993577 | -1.2360296 |
| P8 cl119 | D02 | 1237100 | sample | 0.9239329 | -0.4686008 |
| P8 cl119 | D03 | 1422700 | sample | 1.0625490 | 0.3853247 |
| P8 cl119 | D04 | 1186200 | sample | 0.8859181 | -0.7027862 |
| P8 cl119 | D05 | 1387800 | sample | 1.0364838 | 0.2247536 |
| P8 cl119 | D06 | 1470000 | sample | 1.0978752 | 0.6029469 |
| P8 cl119 | D07 | 1468700 | sample | 1.0969043 | 0.5969657 |
| P8 cl119 | D08 | 1407600 | sample | 1.0512715 | 0.3158512 |
| P8 cl119 | D09 | 1445000 | sample | 1.0792039 | 0.4879246 |
| P8 cl119 | D10 | 1531300 | sample | 1.1436573 | 0.8849816 |
| P8 cl119 | D11 | 1560500 | sample | 1.1654655 | 1.0193276 |
| P8 cl119 | D12 | 1415300 | DMSO | 1.0570223 | 0.3512781 |
| P8 cl119 | E01 | 1060800 | DMSO | 0.7922626 | -1.2797381 |
| P8 cl119 | E02 | 1380900 | sample | 1.0313305 | 0.1930074 |
| P8 cl119 | E03 | 1428000 | sample | 1.0665073 | 0.4097094 |
| P8 cl119 | E04 | 1494400 | sample | 1.1160984 | 0.7152087 |
| P8 cl119 | E05 | 1480800 | sample | 1.1059412 | 0.6526365 |
| P8 cl119 | E06 | 1568400 | sample | 1.1713656 | 1.0556747 |
| P8 cl119 | E07 | 1585900 | sample | 1.1844356 | 1.1361903 |
| P8 cl119 | E08 | 1537300 | sample | 1.1481385 | 0.9125869 |
| P8 cl119 | E09 | 797970 | sample | 0.5959670 | -2.4889905 |
| P8 cl119 | E10 | 1581300 | sample | 1.1810000 | 1.1150262 |

*(continued)*

| Plate | Well | Value | Treatment | Norm | Score |
|---|---|---|---|---|---|
| P8 cl119 | E11 | 1555100 | sample | 1.1614325 | 0.9944828 |
| P8 cl119 | E12 | 1486100 | DMSO | 1.1098995 | 0.6770213 |
| P8 cl119 | F01 | 1060500 | DMSO | 0.7920385 | -1.2811184 |
| P8 cl119 | F02 | 1361100 | sample | 1.0165428 | 0.1019098 |
| P8 cl119 | F03 | 1391900 | sample | 1.0395459 | 0.2436172 |
| P8 cl119 | F04 | 1406300 | sample | 1.0503006 | 0.3098701 |
| P8 cl119 | F05 | 1578000 | sample | 1.1785354 | 1.0998432 |
| P8 cl119 | F06 | 1489000 | sample | 1.1120654 | 0.6903638 |
| P8 cl119 | F07 | 1488400 | sample | 1.1116173 | 0.6876033 |
| P8 cl119 | F08 | 1477900 | sample | 1.1037753 | 0.6392939 |
| P8 cl119 | F09 | 1486600 | sample | 1.1102730 | 0.6793217 |
| P8 cl119 | F10 | 1516100 | sample | 1.1323052 | 0.8150480 |
| P8 cl119 | F11 | 1498000 | sample | 1.1187871 | 0.7317719 |
| P8 cl119 | F12 | 373510 | SAL | 0.2789574 | -4.4418851 |
| P8 cl119 | G01 | 1121500 | empty | 0.8375966 | -1.0004640 |
| P8 cl119 | G02 | 1417000 | sample | 1.0582919 | 0.3590996 |
| P8 cl119 | G03 | 1372900 | sample | 1.0253557 | 0.1562003 |
| P8 cl119 | G04 | 1398900 | sample | 1.0447739 | 0.2758235 |
| P8 cl119 | G05 | 1449000 | sample | 1.0821913 | 0.5063282 |
| P8 cl119 | G06 | 318060 | sample | 0.2375443 | -4.6970046 |
| P8 cl119 | G07 | 1491700 | sample | 1.1140819 | 0.7027862 |
| P8 cl119 | G08 | 1518400 | sample | 1.1340229 | 0.8256301 |
| P8 cl119 | G09 | 1490000 | sample | 1.1128123 | 0.6949647 |
| P8 cl119 | G10 | 1524600 | sample | 1.1386534 | 0.8541556 |
| P8 cl119 | G11 | 1292200 | sample | 0.9650846 | -0.2150917 |
| P8 cl119 | G12 | 334150 | SAL | 0.2495612 | -4.6229762 |
| P8 cl119 | H01 | 1051600 | empty | 0.7853915 | -1.3220663 |
| P8 cl119 | H02 | 1253100 | sample | 0.9358826 | -0.3949866 |
| P8 cl119 | H03 | 1193400 | sample | 0.8912954 | -0.6696598 |
| P8 cl119 | H04 | 1239200 | sample | 0.9255013 | -0.4589390 |
| P8 cl119 | H05 | 1258000 | sample | 0.9395422 | -0.3724422 |
| P8 cl119 | H06 | 1258300 | sample | 0.9397662 | -0.3710619 |
| P8 cl119 | H07 | 1255900 | sample | 0.9379738 | -0.3821041 |
| P8 cl119 | H08 | 1289100 | sample | 0.9627693 | -0.2293545 |
| P8 cl119 | H09 | 1216800 | sample | 0.9087718 | -0.5619990 |
| P8 cl119 | H10 | 1241800 | sample | 0.9274431 | -0.4469767 |
| P8 cl119 | H11 | 141100 | sample | 0.1053811 | -5.5111784 |
| P8 cl119 | H12 | 303300 | SAL | 0.2265208 | -4.7649137 |
| P9 cl119 | A01 | 315600 | SAL | 0.2118192 | -8.1869584 |
| P9 cl119 | A02 | 1371800 | sample | 0.9207020 | -0.8236804 |
| P9 cl119 | A03 | 691410 | sample | 0.4640491 | -5.5670062 |
| P9 cl119 | A04 | 1142600 | sample | 0.7668714 | -2.4215438 |
| P9 cl119 | A05 | 1253100 | sample | 0.8410349 | -1.6511952 |
| P9 cl119 | A06 | 1327300 | sample | 0.8908353 | -1.1339113 |
| P9 cl119 | A07 | 985120 | sample | 0.6611765 | -3.5194126 |
| P9 cl119 | A08 | 1196600 | sample | 0.8031142 | -2.0450839 |
| P9 cl119 | A09 | 1185600 | sample | 0.7957314 | -2.1217702 |
| P9 cl119 | A10 | 1345300 | sample | 0.9029162 | -1.0084247 |
| P9 cl119 | A11 | 1044000 | sample | 0.7006947 | -3.1089318 |
| P9 cl119 | A12 | 1165000 | empty | 0.7819054 | -2.2653827 |
| P9 cl119 | B01 | 362980 | SAL | 0.2436189 | -7.8566496 |
| P9 cl119 | B02 | 1512100 | sample | 1.0148663 | 0.1544183 |
| P9 cl119 | B03 | 1548100 | sample | 1.0390282 | 0.4053916 |
| P9 cl119 | B04 | 1456600 | sample | 0.9776167 | -0.2324989 |
| P9 cl119 | B05 | 1578600 | sample | 1.0594986 | 0.6180218 |
| P9 cl119 | B06 | 1310100 | sample | 0.8792913 | -1.2538208 |
| P9 cl119 | B07 | 1381600 | sample | 0.9272794 | -0.7553599 |
| P9 cl119 | B08 | 1166100 | sample | 0.7826437 | -2.2577140 |
| P9 cl119 | B09 | 1438100 | sample | 0.9652002 | -0.3614713 |
| P9 cl119 | B10 | 1528500 | sample | 1.0258734 | 0.2687506 |
| P9 cl119 | B11 | 1173900 | sample | 0.7878788 | -2.2033365 |

*(continued)*

| Plate | Well | Value | Treatment | Norm | Score |
|---|---|---|---|---|---|
| P9 cl119 | B12 | 1540200 | empty | 1.0337260 | 0.3503169 |
| P9 cl119 | C01 | 374640 | SAL | 0.2514447 | -7.7753622 |
| P9 cl119 | C02 | 1484500 | sample | 0.9963422 | -0.0379946 |
| P9 cl119 | C03 | 1600000 | sample | 1.0738615 | 0.7672115 |
| P9 cl119 | C04 | 1577500 | sample | 1.0587604 | 0.6103531 |
| P9 cl119 | C05 | 1487600 | sample | 0.9984228 | -0.0163830 |
| P9 cl119 | C06 | 1471400 | sample | 0.9875499 | -0.1293210 |
| P9 cl119 | C07 | 788650 | sample | 0.5293131 | -4.8890994 |
| P9 cl119 | C08 | 1457600 | sample | 0.9782879 | -0.2255274 |
| P9 cl119 | C09 | 1494400 | sample | 1.0029867 | 0.0310231 |
| P9 cl119 | C10 | 1542600 | sample | 1.0353368 | 0.3670485 |
| P9 cl119 | C11 | 1705900 | sample | 1.1449377 | 1.5054913 |
| P9 cl119 | C12 | 1595100 | DMSO | 1.0705728 | 0.7330512 |
| P9 cl119 | D01 | 1337700 | DMSO | 0.8978154 | -1.0614079 |
| P9 cl119 | D02 | 1370300 | sample | 0.9196953 | -0.8341377 |
| P9 cl119 | D03 | 1499600 | sample | 1.0064767 | 0.0672748 |
| P9 cl119 | D04 | 1048500 | sample | 0.7037149 | -3.0775602 |
| P9 cl119 | D05 | 1540800 | sample | 1.0341287 | 0.3544998 |
| P9 cl119 | D06 | 1597800 | sample | 1.0723850 | 0.7518742 |
| P9 cl119 | D07 | 1587800 | sample | 1.0656733 | 0.6821594 |
| P9 cl119 | D08 | 1583300 | sample | 1.0626531 | 0.6507877 |
| P9 cl119 | D09 | 1600700 | sample | 1.0743314 | 0.7720915 |
| P9 cl119 | D10 | 1662800 | sample | 1.1160106 | 1.2050204 |
| P9 cl119 | D11 | 1694000 | sample | 1.1369509 | 1.4225306 |
| P9 cl119 | D12 | 1559200 | DMSO | 1.0464781 | 0.4827750 |
| P9 cl119 | E01 | 1421800 | DMSO | 0.9542602 | -0.4751064 |
| P9 cl119 | E02 | 1595900 | sample | 1.0711098 | 0.7386284 |
| P9 cl119 | E03 | 1584200 | sample | 1.0632572 | 0.6570621 |
| P9 cl119 | E04 | 1531600 | sample | 1.0279540 | 0.2903622 |
| P9 cl119 | E05 | 1559200 | sample | 1.0464781 | 0.4827750 |
| P9 cl119 | E06 | 1577600 | sample | 1.0588275 | 0.6110503 |
| P9 cl119 | E07 | 1628500 | sample | 1.0929897 | 0.9658987 |
| P9 cl119 | E08 | 1581300 | sample | 1.0613108 | 0.6368448 |
| P9 cl119 | E09 | 1751000 | sample | 1.1752072 | 1.8199050 |
| P9 cl119 | E10 | 1708900 | sample | 1.1469512 | 1.5264057 |
| P9 cl119 | E11 | 1534500 | sample | 1.0299003 | 0.3105795 |
| P9 cl119 | E12 | 1754900 | DMSO | 1.1778248 | 1.8470938 |
| P9 cl119 | F01 | 1236700 | DMSO | 0.8300279 | -1.7655275 |
| P9 cl119 | F02 | 1488900 | sample | 0.9992953 | -0.0073201 |
| P9 cl119 | F03 | 1432500 | sample | 0.9614417 | -0.4005116 |
| P9 cl119 | F04 | 1369800 | sample | 0.9193597 | -0.8376234 |
| P9 cl119 | F05 | 1502100 | sample | 1.0081546 | 0.0847035 |
| P9 cl119 | F06 | 1583200 | sample | 1.0625860 | 0.6500906 |
| P9 cl119 | F07 | 1601500 | sample | 1.0748683 | 0.7776687 |
| P9 cl119 | F08 | 1602200 | sample | 1.0753381 | 0.7825487 |
| P9 cl119 | F09 | 1616200 | sample | 1.0847344 | 0.8801494 |
| P9 cl119 | F10 | 1565300 | sample | 1.0505722 | 0.5253011 |
| P9 cl119 | F11 | 1674200 | sample | 1.1236619 | 1.2844953 |
| P9 cl119 | F12 | 464190 | SAL | 0.3115474 | -7.1510661 |
| P9 cl119 | G01 | 1275100 | empty | 0.8558005 | -1.4978226 |
| P9 cl119 | G02 | 1430300 | sample | 0.9599651 | -0.4158488 |
| P9 cl119 | G03 | 1522300 | sample | 1.0217121 | 0.2255274 |
| P9 cl119 | G04 | 1519900 | sample | 1.0201013 | 0.2087958 |
| P9 cl119 | G05 | 1610100 | sample | 1.0806403 | 0.8376234 |
| P9 cl119 | G06 | 985740 | sample | 0.6615927 | -3.5150903 |
| P9 cl119 | G07 | 1283200 | sample | 0.8612370 | -1.4413536 |
| P9 cl119 | G08 | 1398600 | sample | 0.9386892 | -0.6368448 |
| P9 cl119 | G09 | 963990 | sample | 0.6469949 | -3.6667200 |
| P9 cl119 | G10 | 1618000 | sample | 1.0859425 | 0.8926981 |
| P9 cl119 | G11 | 1709000 | sample | 1.1470184 | 1.5271029 |
| P9 cl119 | G12 | 431960 | SAL | 0.2899158 | -7.3757569 |
| P9 cl119 | H01 | 1327200 | empty | 0.8907681 | -1.1346085 |

*(continued)*

| Plate | Well | Value | Treatment | Norm | Score |
|---|---|---|---|---|---|
| P9 cl119 | H02 | 1394300 | sample | 0.9358032 | -0.6668221 |
| P9 cl119 | H03 | 1414500 | sample | 0.9493607 | -0.5259982 |
| P9 cl119 | H04 | 1430500 | sample | 0.9600993 | -0.4144545 |
| P9 cl119 | H05 | 1458600 | sample | 0.9789590 | -0.2185559 |
| P9 cl119 | H06 | 1455200 | sample | 0.9766771 | -0.2422590 |
| P9 cl119 | H07 | 1144800 | sample | 0.7683479 | -2.4062066 |
| P9 cl119 | H08 | 1400100 | sample | 0.9396960 | -0.6263875 |
| P9 cl119 | H09 | 1438500 | sample | 0.9654686 | -0.3586827 |
| P9 cl119 | H10 | 1491000 | sample | 1.0007047 | 0.0073201 |
| P9 cl119 | H11 | 1479100 | sample | 0.9927179 | -0.0756406 |
| P9 cl119 | H12 | 390850 | SAL | 0.2623242 | -7.6623545 |
| P10 cl119 | A01 | 256910 | SAL | 0.1733126 | -5.8330837 |
| P10 cl119 | A02 | 1159500 | sample | 0.7822039 | -1.5367632 |
| P10 cl119 | A03 | 1356600 | sample | 0.9151685 | -0.5985689 |
| P10 cl119 | A04 | 1224400 | sample | 0.8259858 | -1.2278397 |
| P10 cl119 | A05 | 1113600 | sample | 0.7512396 | -1.7552468 |
| P10 cl119 | A06 | 371960 | sample | 0.2509259 | -5.2854467 |
| P10 cl119 | A07 | 1185200 | sample | 0.7995413 | -1.4144314 |
| P10 cl119 | A08 | 1212900 | sample | 0.8182278 | -1.2825796 |
| P10 cl119 | A09 | 1349600 | sample | 0.9104463 | -0.6318888 |
| P10 cl119 | A10 | 1277800 | sample | 0.8620096 | -0.9736562 |
| P10 cl119 | A11 | 1342200 | sample | 0.9054542 | -0.6671128 |
| P10 cl119 | A12 | 1289200 | empty | 0.8697001 | -0.9193923 |
| P10 cl119 | B01 | 329150 | SAL | 0.2220461 | -5.4892219 |
| P10 cl119 | B02 | 1341900 | sample | 0.9052518 | -0.6685408 |
| P10 cl119 | B03 | 1581900 | sample | 1.0671569 | 0.4738571 |
| P10 cl119 | B04 | 1672300 | sample | 1.1281411 | 0.9041603 |
| P10 cl119 | B05 | 1566600 | sample | 1.0568354 | 0.4010293 |
| P10 cl119 | B06 | 1496500 | sample | 1.0095457 | 0.0673539 |
| P10 cl119 | B07 | 1480400 | sample | 0.9986845 | -0.0092820 |
| P10 cl119 | B08 | 1406000 | sample | 0.9484939 | -0.3634253 |
| P10 cl119 | B09 | 1481300 | sample | 0.9992917 | -0.0049980 |
| P10 cl119 | B10 | 1548900 | sample | 1.0448949 | 0.3167774 |
| P10 cl119 | B11 | 1558900 | sample | 1.0516410 | 0.3643773 |
| P10 cl119 | B12 | 1594800 | empty | 1.0758593 | 0.5352610 |
| P10 cl119 | C01 | 314420 | SAL | 0.2121092 | -5.5593366 |
| P10 cl119 | C02 | 1365500 | sample | 0.9211725 | -0.5562050 |
| P10 cl119 | C03 | 1599800 | sample | 1.0792323 | 0.5590610 |
| P10 cl119 | C04 | 1516300 | sample | 1.0229028 | 0.1616017 |
| P10 cl119 | C05 | 1446400 | sample | 0.9757480 | -0.1711217 |
| P10 cl119 | C06 | 1563900 | sample | 1.0550140 | 0.3881773 |
| P10 cl119 | C07 | 1527900 | sample | 1.0307282 | 0.2168176 |
| P10 cl119 | C08 | 1635300 | sample | 1.1031808 | 0.7280407 |
| P10 cl119 | C09 | 1154400 | sample | 0.7787634 | -1.5610391 |
| P10 cl119 | C10 | 1521600 | sample | 1.0264782 | 0.1868297 |
| P10 cl119 | C11 | 523250 | sample | 0.3529868 | -4.5653076 |
| P10 cl119 | C12 | 1639100 | DMSO | 1.1057443 | 0.7461286 |
| P10 cl119 | D01 | 1131300 | DMSO | 0.7631801 | -1.6709949 |
| P10 cl119 | D02 | 1317300 | sample | 0.8886565 | -0.7856366 |
| P10 cl119 | D03 | 1518900 | sample | 1.0246568 | 0.1739777 |
| P10 cl119 | D04 | 1436700 | sample | 0.9692043 | -0.2172936 |
| P10 cl119 | D05 | 1570900 | sample | 1.0597362 | 0.4214972 |
| P10 cl119 | D06 | 1664600 | sample | 1.1229467 | 0.8675084 |
| P10 cl119 | D07 | 1392900 | sample | 0.9396566 | -0.4257812 |
| P10 cl119 | D08 | 1622600 | sample | 1.0946133 | 0.6675888 |
| P10 cl119 | D09 | 1612400 | sample | 1.0877323 | 0.6190369 |
| P10 cl119 | D10 | 1668900 | sample | 1.1258475 | 0.8879764 |
| P10 cl119 | D11 | 1622900 | sample | 1.0948157 | 0.6690168 |
| P10 cl119 | D12 | 1649900 | DMSO | 1.1130300 | 0.7975365 |
| P10 cl119 | E01 | 1232500 | DMSO | 0.8314501 | -1.1892838 |
| P10 cl119 | E02 | 1299400 | sample | 0.8765811 | -0.8708404 |
| P10 cl119 | E03 | 1566700 | sample | 1.0569029 | 0.4015053 |

*(continued)*

| Plate | Well | Value | Treatment | Norm | Score |
|---|---|---|---|---|---|
| P10 cl119 | E04 | 1553300 | sample | 1.0478632 | 0.3377214 |
| P10 cl119 | E05 | 1626100 | sample | 1.0969744 | 0.6842487 |
| P10 cl119 | E06 | 1667200 | sample | 1.1247006 | 0.8798844 |
| P10 cl119 | E07 | 1592500 | sample | 1.0743077 | 0.5243130 |
| P10 cl119 | E08 | 1614900 | sample | 1.0894188 | 0.6309368 |
| P10 cl119 | E09 | 1666000 | sample | 1.1238911 | 0.8741724 |
| P10 cl119 | E10 | 1707100 | sample | 1.1516174 | 1.0698080 |
| P10 cl119 | E11 | 1659200 | sample | 1.1193038 | 0.8418045 |
| P10 cl119 | E12 | 1651000 | DMSO | 1.1137721 | 0.8027725 |
| P10 cl119 | F01 | 1125200 | DMSO | 0.7590650 | -1.7000309 |
| P10 cl119 | F02 | 1378400 | sample | 0.9298749 | -0.4948011 |
| P10 cl119 | F03 | 978330 | sample | 0.6599858 | -2.3991308 |
| P10 cl119 | F04 | 1134900 | sample | 0.7656087 | -1.6538590 |
| P10 cl119 | F05 | 1684200 | sample | 1.1361689 | 0.9608042 |
| P10 cl119 | F06 | 1626800 | sample | 1.0974466 | 0.6875807 |
| P10 cl119 | F07 | 1616600 | sample | 1.0905657 | 0.6390288 |
| P10 cl119 | F08 | 364650 | sample | 0.2459945 | -5.3202422 |
| P10 cl119 | F09 | 1307900 | sample | 0.8823152 | -0.8303805 |
| P10 cl119 | F10 | 1640200 | sample | 1.1064863 | 0.7513646 |
| P10 cl119 | F11 | 1635700 | sample | 1.1034506 | 0.7299447 |
| P10 cl119 | F12 | 421620 | SAL | 0.2844268 | -5.0490655 |
| P10 cl119 | G01 | 1026500 | empty | 0.6924815 | -2.1698420 |
| P10 cl119 | G02 | 1298700 | sample | 0.8761089 | -0.8741724 |
| P10 cl119 | G03 | 440430 | sample | 0.2971161 | -4.9595301 |
| P10 cl119 | G04 | 907790 | sample | 0.6123992 | -2.7349006 |
| P10 cl119 | G05 | 1650300 | sample | 1.1132998 | 0.7994405 |
| P10 cl119 | G06 | 1641800 | sample | 1.1075657 | 0.7589806 |
| P10 cl119 | G07 | 1610100 | sample | 1.0861807 | 0.6080889 |
| P10 cl119 | G08 | 1398700 | sample | 0.9435693 | -0.3981733 |
| P10 cl119 | G09 | 1625200 | sample | 1.0963673 | 0.6799647 |
| P10 cl119 | G10 | 1361400 | sample | 0.9184066 | -0.5757209 |
| P10 cl119 | G11 | 282230 | sample | 0.1903936 | -5.7125607 |
| P10 cl119 | G12 | 374760 | SAL | 0.2528148 | -5.2721187 |
| P10 cl119 | H01 | 913260 | empty | 0.6160893 | -2.7088634 |
| P10 cl119 | H02 | 1326800 | sample | 0.8950653 | -0.7404166 |
| P10 cl119 | H03 | 1319300 | sample | 0.8900057 | -0.7761166 |
| P10 cl119 | H04 | 929520 | sample | 0.6270584 | -2.6314660 |
| P10 cl119 | H05 | 1677000 | sample | 1.1313118 | 0.9265323 |
| P10 cl119 | H06 | 1500000 | sample | 1.0119068 | 0.0840138 |
| P10 cl119 | H07 | 1483400 | sample | 1.0007083 | 0.0049980 |
| P10 cl119 | H08 | 1137200 | sample | 0.7671603 | -1.6429110 |
| P10 cl119 | H09 | 1410900 | sample | 0.9517995 | -0.3401014 |
| P10 cl119 | H10 | 1415200 | sample | 0.9547003 | -0.3196334 |
| P10 cl119 | H11 | 1467700 | sample | 0.9901170 | -0.0697339 |
| P10 cl119 | H12 | 331710 | SAL | 0.2237731 | -5.4770363 |
| P11 cl119 | A01 | 329320 | SAL | 0.2206130 | -8.3039448 |
| P11 cl119 | A02 | 1431700 | sample | 0.9591023 | -0.4357424 |
| P11 cl119 | A03 | 1288800 | sample | 0.8633730 | -1.4556867 |
| P11 cl119 | A04 | 1230200 | sample | 0.8241166 | -1.8739423 |
| P11 cl119 | A05 | 1126700 | sample | 0.7547814 | -2.6126703 |
| P11 cl119 | A06 | 1326600 | sample | 0.8886954 | -1.1858904 |
| P11 cl119 | A07 | 1111100 | sample | 0.7443309 | -2.7240148 |
| P11 cl119 | A08 | 1037900 | sample | 0.6952939 | -3.2464775 |
| P11 cl119 | A09 | 1296000 | sample | 0.8681963 | -1.4042969 |
| P11 cl119 | A10 | 1470700 | sample | 0.9852286 | -0.1573812 |
| P11 cl119 | A11 | 1242700 | sample | 0.8324904 | -1.7847240 |
| P11 cl119 | A12 | 1310500 | empty | 0.8779099 | -1.3008036 |
| P11 cl119 | B01 | 349090 | SAL | 0.2338570 | -8.1628371 |
| P11 cl119 | B02 | 63685 | sample | 0.0426629 | -10.1999062 |
| P11 cl119 | B03 | 1511200 | sample | 1.0123597 | 0.1316863 |
| P11 cl119 | B04 | 1470000 | sample | 0.9847597 | -0.1623774 |

*(continued)*

| Plate | Well | Value | Treatment | Norm | Score |
|---|---|---|---|---|---|
| P11 cl119 | B05 | 1459800 | sample | 0.9779266 | -0.2351796 |
| P11 cl119 | B06 | 1546800 | sample | 1.0362083 | 0.3857802 |
| P11 cl119 | B07 | 1335700 | sample | 0.8947915 | -1.1209394 |
| P11 cl119 | B08 | 1367400 | sample | 0.9160275 | -0.8946817 |
| P11 cl119 | B09 | 1498900 | sample | 1.0041199 | 0.0438954 |
| P11 cl119 | B10 | 1480500 | sample | 0.9917937 | -0.0874340 |
| P11 cl119 | B11 | 1621400 | sample | 1.0861832 | 0.9182353 |
| P11 cl119 | B12 | 1602800 | empty | 1.0737230 | 0.7854784 |
| P11 cl119 | C01 | 373610 | SAL | 0.2502830 | -7.9878263 |
| P11 cl119 | C02 | 1445600 | sample | 0.9684140 | -0.3365316 |
| P11 cl119 | C03 | 1486600 | sample | 0.9958801 | -0.0438954 |
| P11 cl119 | C04 | 1388800 | sample | 0.9303634 | -0.7419398 |
| P11 cl119 | C05 | 1605600 | sample | 1.0755987 | 0.8054633 |
| P11 cl119 | C06 | 1536100 | sample | 1.0290404 | 0.3094093 |
| P11 cl119 | C07 | 1499100 | sample | 1.0042539 | 0.0453229 |
| P11 cl119 | C08 | 1573700 | sample | 1.0542288 | 0.5777781 |
| P11 cl119 | C09 | 1645500 | sample | 1.1023279 | 1.0902483 |
| P11 cl119 | C10 | 1630100 | sample | 1.0920114 | 0.9803313 |
| P11 cl119 | C11 | 1144200 | sample | 0.7665048 | -2.4877646 |
| P11 cl119 | C12 | 1595500 | DMSO | 1.0688327 | 0.7333749 |
| P11 cl119 | D01 | 1409700 | DMSO | 0.9443644 | -0.5927667 |
| P11 cl119 | D02 | 1438700 | sample | 0.9637917 | -0.3857802 |
| P11 cl119 | D03 | 1530800 | sample | 1.0254899 | 0.2715807 |
| P11 cl119 | D04 | 1462000 | sample | 0.9794004 | -0.2194772 |
| P11 cl119 | D05 | 1536200 | sample | 1.0291074 | 0.3101230 |
| P11 cl119 | D06 | 1630700 | sample | 1.0924133 | 0.9846138 |
| P11 cl119 | D07 | 1482100 | sample | 0.9928655 | -0.0760140 |
| P11 cl119 | D08 | 1586800 | sample | 1.0630045 | 0.6712789 |
| P11 cl119 | D09 | 1662900 | sample | 1.1139843 | 1.2144402 |
| P11 cl119 | D10 | 1680100 | sample | 1.1255066 | 1.3372047 |
| P11 cl119 | D11 | 1340200 | sample | 0.8978061 | -1.0888208 |
| P11 cl119 | D12 | 1566500 | DMSO | 1.0494055 | 0.5263883 |
| P11 cl119 | E01 | 1293000 | DMSO | 0.8661866 | -1.4257093 |
| P11 cl119 | E02 | 1454800 | sample | 0.9745771 | -0.2708669 |
| P11 cl119 | E03 | 1388900 | sample | 0.9304304 | -0.7412261 |
| P11 cl119 | E04 | 1716000 | sample | 1.1495562 | 1.5934398 |
| P11 cl119 | E05 | 1573600 | sample | 1.0541618 | 0.5770643 |
| P11 cl119 | E06 | 1699100 | sample | 1.1382348 | 1.4728166 |
| P11 cl119 | E07 | 1698800 | sample | 1.1380338 | 1.4706754 |
| P11 cl119 | E08 | 1670100 | sample | 1.1188076 | 1.2658300 |
| P11 cl119 | E09 | 1685900 | sample | 1.1293921 | 1.3786020 |
| P11 cl119 | E10 | 1704100 | sample | 1.1415843 | 1.5085039 |
| P11 cl119 | E11 | 1674500 | sample | 1.1217551 | 1.2972349 |
| P11 cl119 | E12 | 1766900 | DMSO | 1.1836543 | 1.9567369 |
| P11 cl119 | F01 | 1358100 | DMSO | 0.9097974 | -0.9610601 |
| P11 cl119 | F02 | 1397800 | sample | 0.9363926 | -0.6777026 |
| P11 cl119 | F03 | 1471000 | sample | 0.9854296 | -0.1552399 |
| P11 cl119 | F04 | 1574500 | sample | 1.0547647 | 0.5834880 |
| P11 cl119 | F05 | 1585800 | sample | 1.0623346 | 0.6641414 |
| P11 cl119 | F06 | 1601100 | sample | 1.0725842 | 0.7733447 |
| P11 cl119 | F07 | 1568300 | sample | 1.0506113 | 0.5392357 |
| P11 cl119 | F08 | 1602900 | sample | 1.0737900 | 0.7861921 |
| P11 cl119 | F09 | 1407400 | sample | 0.9428236 | -0.6091829 |
| P11 cl119 | F10 | 446240 | sample | 0.2989382 | -7.4694320 |
| P11 cl119 | F11 | 1733500 | sample | 1.1612795 | 1.7183455 |
| P11 cl119 | F12 | 440560 | SAL | 0.2951331 | -7.5099728 |
| P11 cl119 | G01 | 1341500 | empty | 0.8986769 | -1.0795421 |
| P11 cl119 | G02 | 1517900 | sample | 1.0168481 | 0.1795073 |
| P11 cl119 | G03 | 591630 | sample | 0.3963356 | -6.4317155 |
| P11 cl119 | G04 | 1469100 | sample | 0.9841568 | -0.1688011 |
| P11 cl119 | G05 | 1552300 | sample | 1.0398928 | 0.4250362 |
| P11 cl119 | G06 | 744800 | sample | 0.4989449 | -5.3384695 |

(continued)

| Plate | Well | Value | Treatment | Norm | Score |
|---|---|---|---|---|---|
| P11 cl119 | G07 | 1473300 | sample | 0.9869704 | -0.1388238 |
| P11 cl119 | G08 | 1531400 | sample | 1.0258918 | 0.2758632 |
| P11 cl119 | G09 | 1628100 | sample | 1.0906716 | 0.9660563 |
| P11 cl119 | G10 | 1614400 | sample | 1.0814939 | 0.8682730 |
| P11 cl119 | G11 | 1358200 | sample | 0.9098643 | -0.9603464 |
| P11 cl119 | G12 | 438760 | SAL | 0.2939273 | -7.5228203 |
| P11 cl119 | H01 | 1353800 | empty | 0.9069168 | -0.9917512 |
| P11 cl119 | H02 | 1425600 | sample | 0.9550159 | -0.4792810 |
| P11 cl119 | H03 | 1551800 | sample | 1.0395579 | 0.4214675 |
| P11 cl119 | H04 | 1581000 | sample | 1.0591191 | 0.6298816 |
| P11 cl119 | H05 | 1476300 | sample | 0.9889801 | -0.1174114 |
| P11 cl119 | H06 | 1423200 | sample | 0.9534081 | -0.4964109 |
| P11 cl119 | H07 | 1365700 | sample | 0.9148886 | -0.9068154 |
| P11 cl119 | H08 | 1456800 | sample | 0.9759169 | -0.2565920 |
| P11 cl119 | H09 | 1500900 | sample | 1.0054597 | 0.0581704 |
| P11 cl119 | H10 | 1548400 | sample | 1.0372802 | 0.3972001 |
| P11 cl119 | H11 | 1536300 | sample | 1.0291743 | 0.3108367 |
| P11 cl119 | H12 | 351630 | SAL | 0.2355585 | -8.1447079 |
| P12 cl119 | A01 | 290140 | SAL | 0.1978789 | -7.0954859 |
| P12 cl119 | A02 | 1384800 | sample | 0.9444501 | -0.4913888 |
| P12 cl119 | A03 | 1365500 | sample | 0.9312873 | -0.6078260 |
| P12 cl119 | A04 | 1327900 | sample | 0.9056436 | -0.8346672 |
| P12 cl119 | A05 | 1146800 | sample | 0.7821313 | -1.9272457 |
| P12 cl119 | A06 | 1200600 | sample | 0.8188235 | -1.6026697 |
| P12 cl119 | A07 | 940610 | sample | 0.6415072 | -3.1711925 |
| P12 cl119 | A08 | 1258200 | sample | 0.8581074 | -1.2551682 |
| P12 cl119 | A09 | 1160100 | sample | 0.7912020 | -1.8470067 |
| P12 cl119 | A10 | 1263700 | sample | 0.8618585 | -1.2219866 |
| P12 cl119 | A11 | 1346200 | sample | 0.9181245 | -0.7242631 |
| P12 cl119 | A12 | 1226200 | empty | 0.8362830 | -1.4482246 |
| P12 cl119 | B01 | 341500 | SAL | 0.2329071 | -6.7856304 |
| P12 cl119 | B02 | 1377600 | sample | 0.9395396 | -0.5348265 |
| P12 cl119 | B03 | 799290 | sample | 0.5451253 | -4.0237778 |
| P12 cl119 | B04 | 1441300 | sample | 0.9829838 | -0.1505237 |
| P12 cl119 | B05 | 1493100 | sample | 1.0183120 | 0.1619864 |
| P12 cl119 | B06 | 1507300 | sample | 1.0279966 | 0.2476551 |
| P12 cl119 | B07 | 1448800 | sample | 0.9880989 | -0.1052761 |
| P12 cl119 | B08 | 1377300 | sample | 0.9393350 | -0.5366364 |
| P12 cl119 | B09 | 1486800 | sample | 1.0140153 | 0.1239784 |
| P12 cl119 | B10 | 1360800 | sample | 0.9280818 | -0.6361811 |
| P12 cl119 | B11 | 1551400 | sample | 1.0580733 | 0.5137110 |
| P12 cl119 | B12 | 1515800 | empty | 1.0337937 | 0.2989358 |
| P12 cl119 | C01 | 371490 | SAL | 0.2533606 | -6.6047004 |
| P12 cl119 | C02 | 1402000 | sample | 0.9561807 | -0.3876210 |
| P12 cl119 | C03 | 1552400 | sample | 1.0587553 | 0.5197440 |
| P12 cl119 | C04 | 1460500 | sample | 0.9960784 | -0.0346898 |
| P12 cl119 | C05 | 1468200 | sample | 1.0013299 | 0.0117644 |
| P12 cl119 | C06 | 1507100 | sample | 1.0278602 | 0.2464485 |
| P12 cl119 | C07 | 1003700 | sample | 0.6845354 | -2.7905698 |
| P12 cl119 | C08 | 1532900 | sample | 1.0454561 | 0.4021003 |
| P12 cl119 | C09 | 1546000 | sample | 1.0543905 | 0.4811327 |
| P12 cl119 | C10 | 1614500 | sample | 1.1011083 | 0.8943941 |
| P12 cl119 | C11 | 1548100 | sample | 1.0558227 | 0.4938020 |
| P12 cl119 | C12 | 1515900 | DMSO | 1.0338619 | 0.2995391 |
| P12 cl119 | D01 | 1281700 | DMSO | 0.8741347 | -1.1133924 |
| P12 cl119 | D02 | 1495400 | sample | 1.0198806 | 0.1758623 |
| P12 cl119 | D03 | 696780 | sample | 0.4752123 | -4.6422219 |
| P12 cl119 | D04 | 1423300 | sample | 0.9707076 | -0.2591179 |
| P12 cl119 | D05 | 1414300 | sample | 0.9645695 | -0.3134150 |
| P12 cl119 | D06 | 1602000 | sample | 1.0925831 | 0.8189814 |
| P12 cl119 | D07 | 1530600 | sample | 1.0438875 | 0.3882243 |
| P12 cl119 | D08 | 1634000 | sample | 1.1144075 | 1.0120378 |

*(continued)*

| Plate | Well | Value | Treatment | Norm | Score |
|---|---|---|---|---|---|
| P12 cl119 | D09 | 1547400 | sample | 1.0553453 | 0.4895789 |
| P12 cl119 | D10 | 1577500 | sample | 1.0758738 | 0.6711726 |
| P12 cl119 | D11 | 1464300 | sample | 0.9986701 | -0.0117644 |
| P12 cl119 | D12 | 1685500 | DMSO | 1.1495311 | 1.3227379 |
| P12 cl119 | E01 | 1379100 | DMSO | 0.9405627 | -0.5257770 |
| P12 cl119 | E02 | 1588700 | sample | 1.0835124 | 0.7387423 |
| P12 cl119 | E03 | 1611300 | sample | 1.0989258 | 0.8750884 |
| P12 cl119 | E04 | 1626200 | sample | 1.1090878 | 0.9649803 |
| P12 cl119 | E05 | 599060 | sample | 0.4085661 | -5.2317678 |
| P12 cl119 | E06 | 642740 | sample | 0.4383564 | -4.9682458 |
| P12 cl119 | E07 | 1576300 | sample | 1.0750554 | 0.6639330 |
| P12 cl119 | E08 | 1680300 | sample | 1.1459847 | 1.2913663 |
| P12 cl119 | E09 | 1711100 | sample | 1.1669906 | 1.4771830 |
| P12 cl119 | E10 | 1674500 | sample | 1.1420290 | 1.2563748 |
| P12 cl119 | E11 | 1359200 | sample | 0.9269906 | -0.6458340 |
| P12 cl119 | E12 | 1595100 | DMSO | 1.0878772 | 0.7773536 |
| P12 cl119 | F01 | 1419700 | DMSO | 0.9682523 | -0.2808367 |
| P12 cl119 | F02 | 1593700 | sample | 1.0869224 | 0.7689074 |
| P12 cl119 | F03 | 1104300 | sample | 0.7531458 | -2.1836488 |
| P12 cl119 | F04 | 1547200 | sample | 1.0552089 | 0.4883723 |
| P12 cl119 | F05 | 1621100 | sample | 1.1056095 | 0.9342119 |
| P12 cl119 | F06 | 1672200 | sample | 1.1404604 | 1.2424989 |
| P12 cl119 | F07 | 1656900 | sample | 1.1300256 | 1.1501938 |
| P12 cl119 | F08 | 1630500 | sample | 1.1120205 | 0.9909222 |
| P12 cl119 | F09 | 1611900 | sample | 1.0993350 | 0.8787082 |
| P12 cl119 | F10 | 1675400 | sample | 1.1426428 | 1.2618045 |
| P12 cl119 | F11 | 1610000 | sample | 1.0980392 | 0.8672455 |
| P12 cl119 | F12 | 426460 | SAL | 0.2908508 | -6.2730657 |
| P12 cl119 | G01 | 1322000 | empty | 0.9016198 | -0.8702620 |
| P12 cl119 | G02 | 1461300 | sample | 0.9966240 | -0.0298634 |
| P12 cl119 | G03 | 1490200 | sample | 1.0163342 | 0.1444906 |
| P12 cl119 | G04 | 1497500 | sample | 1.0213129 | 0.1885316 |
| P12 cl119 | G05 | 1524600 | sample | 1.0397954 | 0.3520263 |
| P12 cl119 | G06 | 1135600 | sample | 0.7744928 | -1.9948155 |
| P12 cl119 | G07 | 1578600 | sample | 1.0766240 | 0.6778089 |
| P12 cl119 | G08 | 1319600 | sample | 0.8999829 | -0.8847412 |
| P12 cl119 | G09 | 1627800 | sample | 1.1101790 | 0.9746331 |
| P12 cl119 | G10 | 1582200 | sample | 1.0790793 | 0.6995278 |
| P12 cl119 | G11 | 1530400 | sample | 1.0437511 | 0.3870177 |
| P12 cl119 | G12 | 404340 | SAL | 0.2757647 | -6.4065159 |
| P12 cl119 | H01 | 1301700 | empty | 0.8877749 | -0.9927322 |
| P12 cl119 | H02 | 1371700 | sample | 0.9355158 | -0.5704213 |
| P12 cl119 | H03 | 1197700 | sample | 0.8168457 | -1.6201654 |
| P12 cl119 | H04 | 1397100 | sample | 0.9528389 | -0.4171828 |
| P12 cl119 | H05 | 1356300 | sample | 0.9250128 | -0.6633297 |
| P12 cl119 | H06 | 1243300 | sample | 0.8479454 | -1.3450601 |
| P12 cl119 | H07 | 1374900 | sample | 0.9376982 | -0.5511157 |
| P12 cl119 | H08 | 1359700 | sample | 0.9273316 | -0.6428174 |
| P12 cl119 | H09 | 1111900 | sample | 0.7583291 | -2.1377979 |
| P12 cl119 | H10 | 1337200 | sample | 0.9119864 | -0.7785602 |
| P12 cl119 | H11 | 1381800 | sample | 0.9424041 | -0.5094879 |
| P12 cl119 | H12 | 336050 | SAL | 0.2291901 | -6.8185103 |
| P13 cl119 | A01 | 167030 | SAL | 0.1100764 | -7.9442834 |
| P13 cl119 | A02 | 1298400 | sample | 0.8556742 | -1.2883862 |
| P13 cl119 | A03 | 1191000 | sample | 0.7848952 | -1.9202249 |
| P13 cl119 | A04 | 1324300 | sample | 0.8727428 | -1.1360154 |
| P13 cl119 | A05 | 1045000 | sample | 0.6886780 | -2.7791490 |
| P13 cl119 | A06 | 363830 | sample | 0.2397720 | -6.7865007 |
| P13 cl119 | A07 | 1332500 | sample | 0.8781468 | -1.0877745 |
| P13 cl119 | A08 | 1345000 | sample | 0.8863846 | -1.0142364 |
| P13 cl119 | A09 | 1352800 | sample | 0.8915250 | -0.9683487 |

*(continued)*

| Plate | Well | Value | Treatment | Norm | Score |
|---|---|---|---|---|---|
| P13 cl119 | A10 | 913440 | sample | 0.6019771 | -3.5531220 |
| P13 cl119 | A11 | 1045500 | sample | 0.6890075 | -2.7762075 |
| P13 cl119 | A12 | 1465100 | empty | 0.9655331 | -0.3076831 |
| P13 cl119 | B01 | 332160 | SAL | 0.2189008 | -6.9728166 |
| P13 cl119 | B02 | 1423000 | sample | 0.9377883 | -0.5553592 |
| P13 cl119 | B03 | 1562400 | sample | 1.0296560 | 0.2647369 |
| P13 cl119 | B04 | 1172600 | sample | 0.7727692 | -2.0284729 |
| P13 cl119 | B05 | 1520500 | sample | 1.0020430 | 0.0182374 |
| P13 cl119 | B06 | 1466500 | sample | 0.9664558 | -0.2994468 |
| P13 cl119 | B07 | 1495400 | sample | 0.9855015 | -0.1294269 |
| P13 cl119 | B08 | 1482200 | sample | 0.9768024 | -0.2070831 |
| P13 cl119 | B09 | 1529700 | sample | 1.0081060 | 0.0723614 |
| P13 cl119 | B10 | 1598400 | sample | 1.0533808 | 0.4765264 |
| P13 cl119 | B11 | 1554400 | sample | 1.0243838 | 0.2176726 |
| P13 cl119 | B12 | 1655400 | empty | 1.0909450 | 0.8118598 |
| P13 cl119 | C01 | 356170 | SAL | 0.2347239 | -6.8315648 |
| P13 cl119 | C02 | 1383500 | sample | 0.9117570 | -0.7877393 |
| P13 cl119 | C03 | 1556600 | sample | 1.0258337 | 0.2306152 |
| P13 cl119 | C04 | 1400400 | sample | 0.9228944 | -0.6883159 |
| P13 cl119 | C05 | 1252100 | sample | 0.8251615 | -1.5607710 |
| P13 cl119 | C06 | 1601300 | sample | 1.0552919 | 0.4935872 |
| P13 cl119 | C07 | 1666700 | sample | 1.0983920 | 0.8783382 |
| P13 cl119 | C08 | 1580000 | sample | 1.0412548 | 0.3682784 |
| P13 cl119 | C09 | 1531300 | sample | 1.0091604 | 0.0817743 |
| P13 cl119 | C10 | 1357100 | sample | 0.8943588 | -0.9430516 |
| P13 cl119 | C11 | 1699500 | sample | 1.1200079 | 1.0713019 |
| P13 cl119 | C12 | 1658500 | DMSO | 1.0929880 | 0.8300972 |
| P13 cl119 | D01 | 1284300 | DMSO | 0.8463820 | -1.3713371 |
| P13 cl119 | D02 | 1601600 | sample | 1.0554897 | 0.4953521 |
| P13 cl119 | D03 | 1365300 | sample | 0.8997628 | -0.8948107 |
| P13 cl119 | D04 | 1142300 | sample | 0.7528008 | -2.2067290 |
| P13 cl119 | D05 | 1589600 | sample | 1.0475814 | 0.4247556 |
| P13 cl119 | D06 | 1341900 | sample | 0.8843416 | -1.0324739 |
| P13 cl119 | D07 | 1458500 | sample | 0.9611836 | -0.3465112 |
| P13 cl119 | D08 | 1602100 | sample | 1.0558192 | 0.4982937 |
| P13 cl119 | D09 | 585930 | sample | 0.3861408 | -5.4798771 |
| P13 cl119 | D10 | 1201500 | sample | 0.7918149 | -1.8584530 |
| P13 cl119 | D11 | 1575000 | sample | 1.0379597 | 0.3388632 |
| P13 cl119 | D12 | 1592200 | DMSO | 1.0492948 | 0.4400515 |
| P13 cl119 | E01 | 1342100 | DMSO | 0.8844734 | -1.0312973 |
| P13 cl119 | E02 | 1620700 | sample | 1.0680770 | 0.6077182 |
| P13 cl119 | E03 | 1664900 | sample | 1.0972057 | 0.8677487 |
| P13 cl119 | E04 | 1514300 | sample | 0.9979570 | -0.0182374 |
| P13 cl119 | E05 | 1657500 | sample | 1.0923290 | 0.8242142 |
| P13 cl119 | E06 | 1687900 | sample | 1.1123633 | 1.0030587 |
| P13 cl119 | E07 | 1671100 | sample | 1.1012917 | 0.9042235 |
| P13 cl119 | E08 | 1602800 | sample | 1.0562805 | 0.5024118 |
| P13 cl119 | E09 | 1670100 | sample | 1.1006327 | 0.8983405 |
| P13 cl119 | E10 | 1643700 | sample | 1.0832345 | 0.7430282 |
| P13 cl119 | E11 | 1659200 | sample | 1.0934493 | 0.8342153 |
| P13 cl119 | E12 | 1606800 | DMSO | 1.0589166 | 0.5259440 |
| P13 cl119 | F01 | 1404900 | DMSO | 0.9258600 | -0.6618422 |
| P13 cl119 | F02 | 1637800 | sample | 1.0793463 | 0.7083183 |
| P13 cl119 | F03 | 1511800 | sample | 0.9963095 | -0.0329450 |
| P13 cl119 | F04 | 776680 | sample | 0.5118492 | -4.3576868 |
| P13 cl119 | F05 | 1111300 | sample | 0.7323712 | -2.3891033 |
| P13 cl119 | F06 | 1640800 | sample | 1.0813233 | 0.7259674 |
| P13 cl119 | F07 | 1599900 | sample | 1.0543693 | 0.4853510 |
| P13 cl119 | F08 | 1638100 | sample | 1.0795440 | 0.7100832 |
| P13 cl119 | F09 | 1693400 | sample | 1.1159879 | 1.0354154 |
| P13 cl119 | F10 | 1624200 | sample | 1.0703836 | 0.6283089 |
| P13 cl119 | F11 | 1629700 | sample | 1.0740082 | 0.6606656 |

*(continued)*

| Plate | Well | Value | Treatment | Norm | Score |
|---|---|---|---|---|---|
| P13 cl119 | F12 | 400790 | SAL | 0.2641294 | -6.5690635 |
| P13 cl119 | G01 | 754160 | empty | 0.4970080 | -4.4901729 |
| P13 cl119 | G02 | 1635500 | sample | 1.0778305 | 0.6947873 |
| P13 cl119 | G03 | 1597700 | sample | 1.0529195 | 0.4724083 |
| P13 cl119 | G04 | 1480200 | sample | 0.9754844 | -0.2188492 |
| P13 cl119 | G05 | 1548900 | sample | 1.0207592 | 0.1853158 |
| P13 cl119 | G06 | 1623700 | sample | 1.0700540 | 0.6253674 |
| P13 cl119 | G07 | 1542300 | sample | 1.0164096 | 0.1464877 |
| P13 cl119 | G08 | 1555300 | sample | 1.0249769 | 0.2229673 |
| P13 cl119 | G09 | 1575900 | sample | 1.0385528 | 0.3441580 |
| P13 cl119 | G10 | 1586900 | sample | 1.0458020 | 0.4088714 |
| P13 cl119 | G11 | 1599700 | sample | 1.0542375 | 0.4841744 |
| P13 cl119 | G12 | 393560 | SAL | 0.2593647 | -6.6115979 |
| P13 cl119 | H01 | 1063500 | empty | 0.7008699 | -2.6703127 |
| P13 cl119 | H02 | 1198200 | sample | 0.7896402 | -1.8778670 |
| P13 cl119 | H03 | 1441500 | sample | 0.9499802 | -0.4465229 |
| P13 cl119 | H04 | 1501000 | sample | 0.9891920 | -0.0964819 |
| P13 cl119 | H05 | 1459600 | sample | 0.9619085 | -0.3400398 |
| P13 cl119 | H06 | 1271500 | sample | 0.8379465 | -1.4466400 |
| P13 cl119 | H07 | 1419600 | sample | 0.9355476 | -0.5753615 |
| P13 cl119 | H08 | 1334300 | sample | 0.8793331 | -1.0771850 |
| P13 cl119 | H09 | 1469100 | sample | 0.9681692 | -0.2841509 |
| P13 cl119 | H10 | 1485000 | sample | 0.9786477 | -0.1906106 |
| P13 cl119 | H11 | 145480 | sample | 0.0958745 | -8.0710629 |
| P13 cl119 | H12 | 287690 | SAL | 0.1895940 | -7.2344355 |
| P14 cl119 | A01 | 218910 | SAL | 0.1413736 | -6.8272740 |
| P14 cl119 | A02 | 1392000 | sample | 0.8989635 | -0.8033809 |
| P14 cl119 | A03 | 1345600 | sample | 0.8689980 | -1.0416479 |
| P14 cl119 | A04 | 1037000 | sample | 0.6697020 | -2.6263289 |
| P14 cl119 | A05 | 1060800 | sample | 0.6850722 | -2.5041143 |
| P14 cl119 | A06 | 1408100 | sample | 0.9093610 | -0.7207063 |
| P14 cl119 | A07 | 1552400 | sample | 1.0025509 | 0.0202835 |
| P14 cl119 | A08 | 1391200 | sample | 0.8984468 | -0.8074889 |
| P14 cl119 | A09 | 873460 | sample | 0.5640867 | -3.4661174 |
| P14 cl119 | A10 | 1452900 | sample | 0.9382931 | -0.4906554 |
| P14 cl119 | A11 | 1385800 | sample | 0.8949595 | -0.8352183 |
| P14 cl119 | A12 | 1288000 | empty | 0.8317995 | -1.3374276 |
| P14 cl119 | B01 | 254830 | SAL | 0.1645710 | -6.6428225 |
| P14 cl119 | B02 | 1450200 | sample | 0.9365495 | -0.5045201 |
| P14 cl119 | B03 | 1472600 | sample | 0.9510155 | -0.3894947 |
| P14 cl119 | B04 | 1394500 | sample | 0.9005780 | -0.7905432 |
| P14 cl119 | B05 | 1454600 | sample | 0.9393910 | -0.4819258 |
| P14 cl119 | B06 | 1448800 | sample | 0.9356453 | -0.5117092 |
| P14 cl119 | B07 | 515710 | sample | 0.3330492 | -5.3031868 |
| P14 cl119 | B08 | 1513400 | sample | 0.9773645 | -0.1799840 |
| P14 cl119 | B09 | 1531500 | sample | 0.9890536 | -0.0870393 |
| P14 cl119 | B10 | 1698000 | sample | 1.0965805 | 0.7679489 |
| P14 cl119 | B11 | 1655900 | sample | 1.0693920 | 0.5517627 |
| P14 cl119 | B12 | 1705600 | empty | 1.1014886 | 0.8069754 |
| P14 cl119 | C01 | 278570 | SAL | 0.1799025 | -6.5209161 |
| P14 cl119 | C02 | 1278100 | sample | 0.8254061 | -1.3882648 |
| P14 cl119 | C03 | 1473800 | sample | 0.9517905 | -0.3833326 |
| P14 cl119 | C04 | 1336500 | sample | 0.8631212 | -1.0883770 |
| P14 cl119 | C05 | 1454300 | sample | 0.9391973 | -0.4834663 |
| P14 cl119 | C06 | 1610800 | sample | 1.0402661 | 0.3201713 |
| P14 cl119 | C07 | 1665600 | sample | 1.0756563 | 0.6015728 |
| P14 cl119 | C08 | 1603300 | sample | 1.0354225 | 0.2816583 |
| P14 cl119 | C09 | 642460 | sample | 0.4149052 | -4.6523173 |
| P14 cl119 | C10 | 1763700 | sample | 1.1390100 | 1.1053227 |
| P14 cl119 | C11 | 204520 | sample | 0.1320805 | -6.9011676 |
| P14 cl119 | C12 | 1664300 | DMSO | 1.0748168 | 0.5948973 |
| P14 cl119 | D01 | 996980 | DMSO | 0.6438568 | -2.8318342 |

*(continued)*

| Plate | Well | Value | Treatment | Norm | Score |
|---|---|---|---|---|---|
| P14 cl119 | D02 | 1489400 | sample | 0.9618651 | -0.3032256 |
| P14 cl119 | D03 | 1546100 | sample | 0.9984824 | -0.0120674 |
| P14 cl119 | D04 | 1283100 | sample | 0.8286351 | -1.3625894 |
| P14 cl119 | D05 | 1533200 | sample | 0.9901514 | -0.0783097 |
| P14 cl119 | D06 | 1730500 | sample | 1.1175692 | 0.9348385 |
| P14 cl119 | D07 | 1680200 | sample | 1.0850851 | 0.6765448 |
| P14 cl119 | D08 | 1645100 | sample | 1.0624173 | 0.4963040 |
| P14 cl119 | D09 | 1679600 | sample | 1.0846976 | 0.6734637 |
| P14 cl119 | D10 | 1627800 | sample | 1.0512448 | 0.4074674 |
| P14 cl119 | D11 | 771940 | sample | 0.4985243 | -3.9874292 |
| P14 cl119 | D12 | 1548100 | DMSO | 0.9997740 | -0.0017973 |
| P14 cl119 | E01 | 950050 | DMSO | 0.6135490 | -3.0728228 |
| P14 cl119 | E02 | 1355500 | sample | 0.8753915 | -0.9908108 |
| P14 cl119 | E03 | 1391000 | sample | 0.8983177 | -0.8085160 |
| P14 cl119 | E04 | 1345300 | sample | 0.8688043 | -1.0431884 |
| P14 cl119 | E05 | 1559600 | sample | 1.0072007 | 0.0572560 |
| P14 cl119 | E06 | 1776600 | sample | 1.1473409 | 1.1715650 |
| P14 cl119 | E07 | 1752400 | sample | 1.1317124 | 1.0472965 |
| P14 cl119 | E08 | 1702000 | sample | 1.0991637 | 0.7884892 |
| P14 cl119 | E09 | 1709100 | sample | 1.1037489 | 0.8249482 |
| P14 cl119 | E10 | 1719500 | sample | 1.1104653 | 0.8783528 |
| P14 cl119 | E11 | 1705100 | sample | 1.1011657 | 0.8044079 |
| P14 cl119 | E12 | 1689700 | DMSO | 1.0912203 | 0.7253279 |
| P14 cl119 | F01 | 990630 | DMSO | 0.6397559 | -2.8644418 |
| P14 cl119 | F02 | 1620400 | sample | 1.0464658 | 0.3694679 |
| P14 cl119 | F03 | 1529800 | sample | 0.9879557 | -0.0957690 |
| P14 cl119 | F04 | 1305000 | sample | 0.8427783 | -1.2501315 |
| P14 cl119 | F05 | 1650100 | sample | 1.0656463 | 0.5219793 |
| P14 cl119 | F06 | 1757800 | sample | 1.1351997 | 1.0750258 |
| P14 cl119 | F07 | 1760400 | sample | 1.1368788 | 1.0883770 |
| P14 cl119 | F08 | 1659800 | sample | 1.0719106 | 0.5717895 |
| P14 cl119 | F09 | 1685600 | sample | 1.0885724 | 0.7042741 |
| P14 cl119 | F10 | 1651600 | sample | 1.0666150 | 0.5296819 |
| P14 cl119 | F11 | 1673200 | sample | 1.0805644 | 0.6405993 |
| P14 cl119 | F12 | 400820 | SAL | 0.2588524 | -5.8931544 |
| P14 cl119 | G01 | 1005800 | empty | 0.6495528 | -2.7865429 |
| P14 cl119 | G02 | 1562600 | sample | 1.0091382 | 0.0726612 |
| P14 cl119 | G03 | 1583000 | sample | 1.0223126 | 0.1774165 |
| P14 cl119 | G04 | 1268000 | sample | 0.8188834 | -1.4401289 |
| P14 cl119 | G05 | 1728500 | sample | 1.1162776 | 0.9245684 |
| P14 cl119 | G06 | 1607400 | sample | 1.0380703 | 0.3027121 |
| P14 cl119 | G07 | 1644500 | sample | 1.0620298 | 0.4932230 |
| P14 cl119 | G08 | 1680000 | sample | 1.0849559 | 0.6755178 |
| P14 cl119 | G09 | 1610400 | sample | 1.0400077 | 0.3181173 |
| P14 cl119 | G10 | 1672900 | sample | 1.0803707 | 0.6390588 |
| P14 cl119 | G11 | 650620 | sample | 0.4201750 | -4.6104152 |
| P14 cl119 | G12 | 390980 | SAL | 0.2524977 | -5.9436834 |
| P14 cl119 | H01 | 903260 | empty | 0.5833317 | -3.3130924 |
| P14 cl119 | H02 | 1452600 | sample | 0.9380994 | -0.4921960 |
| P14 cl119 | H03 | 1409100 | sample | 0.9100068 | -0.7155713 |
| P14 cl119 | H04 | 1252200 | sample | 0.8086796 | -1.5212629 |
| P14 cl119 | H05 | 1547800 | sample | 0.9995802 | -0.0033378 |
| P14 cl119 | H06 | 1549100 | sample | 1.0004198 | 0.0033378 |
| P14 cl119 | H07 | 1269400 | sample | 0.8197875 | -1.4329398 |
| P14 cl119 | H08 | 1614100 | sample | 1.0423972 | 0.3371170 |
| P14 cl119 | H09 | 1618500 | sample | 1.0452388 | 0.3597113 |
| P14 cl119 | H10 | 1589800 | sample | 1.0267041 | 0.2123349 |
| P14 cl119 | H11 | 1593800 | sample | 1.0292874 | 0.2328752 |
| P14 cl119 | H12 | 359180 | SAL | 0.2319610 | -6.1069785 |
| P15 cl119 | A01 | 311330 | SAL | 0.2076710 | -5.5656382 |
| P15 cl119 | A02 | 1314500 | sample | 0.8768302 | -0.8651943 |

*(continued)*

| Plate | Well | Value | Treatment | Norm | Score |
|---|---|---|---|---|---|
| P15 cl119 | A03 | 1429400 | sample | 0.9534736 | -0.3268199 |
| P15 cl119 | A04 | 1389100 | sample | 0.9265917 | -0.5156492 |
| P15 cl119 | A05 | 1239000 | sample | 0.8264683 | -1.2189564 |
| P15 cl119 | A06 | 1306300 | sample | 0.8713604 | -0.9036161 |
| P15 cl119 | A07 | 1309300 | sample | 0.8733616 | -0.8895594 |
| P15 cl119 | A08 | 1322500 | sample | 0.8821666 | -0.8277096 |
| P15 cl119 | A09 | 1282200 | sample | 0.8552847 | -1.0165389 |
| P15 cl119 | A10 | 1213200 | sample | 0.8092586 | -1.3398446 |
| P15 cl119 | A11 | 1387900 | sample | 0.9257913 | -0.5212719 |
| P15 cl119 | A12 | 1603000 | empty | 1.0692726 | 0.4865986 |
| P15 cl119 | B01 | 341630 | SAL | 0.2278825 | -5.4236648 |
| P15 cl119 | B02 | 1535700 | sample | 1.0243805 | 0.1712583 |
| P15 cl119 | B03 | 1362100 | sample | 0.9085815 | -0.6421602 |
| P15 cl119 | B04 | 1499800 | sample | 1.0004336 | 0.0030456 |
| P15 cl119 | B05 | 1569900 | sample | 1.0471934 | 0.3315055 |
| P15 cl119 | B06 | 1568600 | sample | 1.0463263 | 0.3254143 |
| P15 cl119 | B07 | 1426400 | sample | 0.9514725 | -0.3408767 |
| P15 cl119 | B08 | 1232000 | sample | 0.8217990 | -1.2517555 |
| P15 cl119 | B09 | 1525400 | sample | 1.0175099 | 0.1229968 |
| P15 cl119 | B10 | 1590200 | sample | 1.0607344 | 0.4266230 |
| P15 cl119 | B11 | 1495400 | sample | 0.9974986 | -0.0175710 |
| P15 cl119 | B12 | 1614600 | empty | 1.0770103 | 0.5409514 |
| P15 cl119 | C01 | 387760 | SAL | 0.2586532 | -5.2075185 |
| P15 cl119 | C02 | 1469000 | sample | 0.9798886 | -0.1412706 |
| P15 cl119 | C03 | 1624200 | sample | 1.0834139 | 0.5859331 |
| P15 cl119 | C04 | 1510800 | sample | 1.0077711 | 0.0545871 |
| P15 cl119 | C05 | 1644800 | sample | 1.0971551 | 0.6824563 |
| P15 cl119 | C06 | 1667100 | sample | 1.1120302 | 0.7869449 |
| P15 cl119 | C07 | 1600800 | sample | 1.0678051 | 0.4762903 |
| P15 cl119 | C08 | 1507100 | sample | 1.0053030 | 0.0372504 |
| P15 cl119 | C09 | 284350 | sample | 0.1896741 | -5.6920554 |
| P15 cl119 | C10 | 1558000 | sample | 1.0392556 | 0.2757470 |
| P15 cl119 | C11 | 1718600 | sample | 1.1463830 | 1.0282528 |
| P15 cl119 | C12 | 1730100 | DMSO | 1.1540540 | 1.0821371 |
| P15 cl119 | D01 | 1431800 | DMSO | 0.9550745 | -0.3155745 |
| P15 cl119 | D02 | 507100 | sample | 0.3382583 | -4.6483401 |
| P15 cl119 | D03 | 1537400 | sample | 1.0255145 | 0.1792238 |
| P15 cl119 | D04 | 1498500 | sample | 0.9995664 | -0.0030456 |
| P15 cl119 | D05 | 1585900 | sample | 1.0578661 | 0.4064750 |
| P15 cl119 | D06 | 1564000 | sample | 1.0432578 | 0.3038605 |
| P15 cl119 | D07 | 1613700 | sample | 1.0764100 | 0.5367344 |
| P15 cl119 | D08 | 1626000 | sample | 1.0846146 | 0.5943672 |
| P15 cl119 | D09 | 1651900 | sample | 1.1018911 | 0.7157240 |
| P15 cl119 | D10 | 1658200 | sample | 1.1060935 | 0.7452432 |
| P15 cl119 | D11 | 1681400 | sample | 1.1215689 | 0.8539489 |
| P15 cl119 | D12 | 1658100 | DMSO | 1.1060267 | 0.7447746 |
| P15 cl119 | E01 | 1444200 | DMSO | 0.9633459 | -0.2574732 |
| P15 cl119 | E02 | 1615600 | sample | 1.0776774 | 0.5456370 |
| P15 cl119 | E03 | 1269500 | sample | 0.8468132 | -1.0760459 |
| P15 cl119 | E04 | 1685000 | sample | 1.1239702 | 0.8708170 |
| P15 cl119 | E05 | 1641400 | sample | 1.0948871 | 0.6665253 |
| P15 cl119 | E06 | 1545200 | sample | 1.0307174 | 0.2157714 |
| P15 cl119 | E07 | 1680000 | sample | 1.1206350 | 0.8473891 |
| P15 cl119 | E08 | 1476100 | sample | 0.9846246 | -0.1080029 |
| P15 cl119 | E09 | 1697500 | sample | 1.1323083 | 0.9293869 |
| P15 cl119 | E10 | 1674300 | sample | 1.1168329 | 0.8206812 |
| P15 cl119 | E11 | 1709300 | sample | 1.1401794 | 0.9846769 |
| P15 cl119 | E12 | 1665400 | DMSO | 1.1108962 | 0.7789794 |
| P15 cl119 | F01 | 1318300 | DMSO | 0.8793650 | -0.8473891 |
| P15 cl119 | F02 | 473560 | sample | 0.3158857 | -4.8054948 |
| P15 cl119 | F03 | 1346000 | sample | 0.8978421 | -0.7175982 |
| P15 cl119 | F04 | 1468600 | sample | 0.9796218 | -0.1431448 |

*(continued)*

| Plate | Well | Value | Treatment | Norm | Score |
|---|---|---|---|---|---|
| P15 cl119 | F05 | 1614300 | sample | 1.0768102 | 0.5395458 |
| P15 cl119 | F06 | 1661500 | sample | 1.1082947 | 0.7607056 |
| P15 cl119 | F07 | 915920 | sample | 0.6109595 | -2.7327770 |
| P15 cl119 | F08 | 1594200 | sample | 1.0634026 | 0.4453654 |
| P15 cl119 | F09 | 1594800 | sample | 1.0638028 | 0.4481767 |
| P15 cl119 | F10 | 1661200 | sample | 1.1080946 | 0.7592999 |
| P15 cl119 | F11 | 1497900 | sample | 0.9991662 | -0.0058570 |
| P15 cl119 | F12 | 422040 | SAL | 0.2815195 | -5.0468964 |
| P15 cl119 | G01 | 1355900 | empty | 0.9044459 | -0.6712108 |
| P15 cl119 | G02 | 1491000 | sample | 0.9945636 | -0.0381876 |
| P15 cl119 | G03 | 1523200 | sample | 1.0160424 | 0.1126885 |
| P15 cl119 | G04 | 287500 | sample | 0.1917753 | -5.6772958 |
| P15 cl119 | G05 | 1570600 | sample | 1.0476603 | 0.3347854 |
| P15 cl119 | G06 | 1546500 | sample | 1.0315846 | 0.2218627 |
| P15 cl119 | G07 | 1482600 | sample | 0.9889604 | -0.0775465 |
| P15 cl119 | G08 | 934710 | sample | 0.6234933 | -2.6447347 |
| P15 cl119 | G09 | 1328800 | sample | 0.8863689 | -0.7981903 |
| P15 cl119 | G10 | 1605000 | sample | 1.0706067 | 0.4959698 |
| P15 cl119 | G11 | 1614900 | sample | 1.0772104 | 0.5423571 |
| P15 cl119 | G12 | 393300 | SAL | 0.2623487 | -5.1815603 |
| P15 cl119 | H01 | 1208400 | empty | 0.8060568 | -1.3623355 |
| P15 cl119 | H02 | 839850 | sample | 0.5602175 | -3.0892098 |
| P15 cl119 | H03 | 1333900 | sample | 0.8897709 | -0.7742938 |
| P15 cl119 | H04 | 1271000 | sample | 0.8478138 | -1.0690175 |
| P15 cl119 | H05 | 1324800 | sample | 0.8837008 | -0.8169327 |
| P15 cl119 | H06 | 940990 | sample | 0.6276824 | -2.6153092 |
| P15 cl119 | H07 | 59410 | sample | 0.0396291 | -6.7460321 |
| P15 cl119 | H08 | 53309 | sample | 0.0355595 | -6.7746189 |
| P15 cl119 | H09 | 1087500 | sample | 0.7254111 | -1.9288233 |
| P15 cl119 | H10 | 1310900 | sample | 0.8744288 | -0.8820624 |
| P15 cl119 | H11 | 1093000 | sample | 0.7290798 | -1.9030526 |
| P15 cl119 | H12 | 339530 | SAL | 0.2264817 | -5.4335045 |
| P16 cl119 | A01 | 327550 | SAL | 0.2155572 | -6.3331468 |
| P16 cl119 | A02 | 1095600 | sample | 0.7210029 | -2.2524644 |
| P16 cl119 | A03 | 1387000 | sample | 0.9127702 | -0.7042438 |
| P16 cl119 | A04 | 1154100 | sample | 0.7595012 | -1.9416514 |
| P16 cl119 | A05 | 1358100 | sample | 0.8937514 | -0.8577907 |
| P16 cl119 | A06 | 1385300 | sample | 0.9116515 | -0.7132760 |
| P16 cl119 | A07 | 1238100 | sample | 0.8147807 | -1.4953558 |
| P16 cl119 | A08 | 1128400 | sample | 0.7425883 | -2.0781966 |
| P16 cl119 | A09 | 1444900 | sample | 0.9508736 | -0.3966186 |
| P16 cl119 | A10 | 1368000 | sample | 0.9002665 | -0.8051916 |
| P16 cl119 | A11 | 1437900 | sample | 0.9462670 | -0.4338099 |
| P16 cl119 | A12 | 1162000 | empty | 0.7647001 | -1.8996784 |
| P16 cl119 | B01 | 390600 | SAL | 0.2570498 | -5.9981595 |
| P16 cl119 | B02 | 1532700 | sample | 1.0086539 | 0.0698665 |
| P16 cl119 | B03 | 1580500 | sample | 1.0401106 | 0.3238299 |
| P16 cl119 | B04 | 1572400 | sample | 1.0347800 | 0.2807943 |
| P16 cl119 | B05 | 97123 | sample | 0.0639156 | -7.5574153 |
| P16 cl119 | B06 | 1705000 | sample | 1.1220427 | 0.9853038 |
| P16 cl119 | B07 | 1460500 | sample | 0.9611398 | -0.3137352 |
| P16 cl119 | B08 | 1589100 | sample | 1.0457701 | 0.3695221 |
| P16 cl119 | B09 | 1417300 | sample | 0.9327103 | -0.5432586 |
| P16 cl119 | B10 | 1608600 | sample | 1.0586029 | 0.4731264 |
| P16 cl119 | B11 | 837200 | sample | 0.5509526 | -3.6253546 |
| P16 cl119 | B12 | 1649500 | empty | 1.0855187 | 0.6904299 |
| P16 cl119 | C01 | 383250 | SAL | 0.2522128 | -6.0372103 |
| P16 cl119 | C02 | 1368700 | sample | 0.9007272 | -0.8014725 |
| P16 cl119 | C03 | 1419100 | sample | 0.9338949 | -0.5336951 |
| P16 cl119 | C04 | 1433500 | sample | 0.9433714 | -0.4571873 |
| P16 cl119 | C05 | 1393700 | sample | 0.9171794 | -0.6686464 |
| P16 cl119 | C06 | 1627700 | sample | 1.0711724 | 0.5746056 |

*(continued)*

| Plate | Well | Value | Treatment | Norm | Score |
|---|---|---|---|---|---|
| P16 cl119 | C07 | 1601100 | sample | 1.0536672 | 0.4332786 |
| P16 cl119 | C08 | 1449900 | sample | 0.9541641 | -0.3700534 |
| P16 cl119 | C09 | 1644200 | sample | 1.0820309 | 0.6622708 |
| P16 cl119 | C10 | 1638300 | sample | 1.0781481 | 0.6309238 |
| P16 cl119 | C11 | 1672600 | sample | 1.1007206 | 0.8131612 |
| P16 cl119 | C12 | 1642400 | DMSO | 1.0808463 | 0.6527073 |
| P16 cl119 | D01 | 1424000 | DMSO | 0.9371195 | -0.5076612 |
| P16 cl119 | D02 | 342470 | sample | 0.2253759 | -6.2538762 |
| P16 cl119 | D03 | 1609700 | sample | 1.0593268 | 0.4789708 |
| P16 cl119 | D04 | 1332900 | sample | 0.8771676 | -0.9916794 |
| P16 cl119 | D05 | 1165400 | sample | 0.7669376 | -1.8816140 |
| P16 cl119 | D06 | 1568200 | sample | 1.0320161 | 0.2584795 |
| P16 cl119 | D07 | 1560000 | sample | 1.0266197 | 0.2149126 |
| P16 cl119 | D08 | 1555000 | sample | 1.0233293 | 0.1883474 |
| P16 cl119 | D09 | 1675000 | sample | 1.1023000 | 0.8259125 |
| P16 cl119 | D10 | 1524300 | sample | 1.0031259 | 0.0252370 |
| P16 cl119 | D11 | 1734800 | sample | 1.1416538 | 1.1436324 |
| P16 cl119 | D12 | 294890 | DMSO | 0.1940640 | -6.5066708 |
| P16 cl119 | E01 | 1418200 | DMSO | 0.9333026 | -0.5384769 |
| P16 cl119 | E02 | 1679000 | sample | 1.1049324 | 0.8471646 |
| P16 cl119 | E03 | 1729800 | sample | 1.1383633 | 1.1170672 |
| P16 cl119 | E04 | 1550300 | sample | 1.0202363 | 0.1633761 |
| P16 cl119 | E05 | 1251700 | sample | 0.8237307 | -1.4230985 |
| P16 cl119 | E06 | 1691200 | sample | 1.1129611 | 0.9119838 |
| P16 cl119 | E07 | 1678700 | sample | 1.1047350 | 0.8455707 |
| P16 cl119 | E08 | 1573100 | sample | 1.0352407 | 0.2845134 |
| P16 cl119 | E09 | 1660200 | sample | 1.0925603 | 0.7472794 |
| P16 cl119 | E10 | 1749700 | sample | 1.1514593 | 1.2227968 |
| P16 cl119 | E11 | 1617000 | sample | 1.0641308 | 0.5177560 |
| P16 cl119 | E12 | 1429800 | DMSO | 0.9409365 | -0.4768456 |
| P16 cl119 | F01 | 1437200 | DMSO | 0.9458063 | -0.4375291 |
| P16 cl119 | F02 | 1598500 | sample | 1.0519562 | 0.4194647 |
| P16 cl119 | F03 | 1588400 | sample | 1.0453095 | 0.3658030 |
| P16 cl119 | F04 | 1551000 | sample | 1.0206969 | 0.1670952 |
| P16 cl119 | F05 | 1642400 | sample | 1.0808463 | 0.6527073 |
| P16 cl119 | F06 | 1652900 | sample | 1.0877562 | 0.7084942 |
| P16 cl119 | F07 | 1573900 | sample | 1.0357672 | 0.2887639 |
| P16 cl119 | F08 | 1520300 | sample | 1.0004936 | 0.0039848 |
| P16 cl119 | F09 | 1621500 | sample | 1.0670922 | 0.5416647 |
| P16 cl119 | F10 | 1647600 | sample | 1.0842684 | 0.6803351 |
| P16 cl119 | F11 | 1689700 | sample | 1.1119739 | 0.9040142 |
| P16 cl119 | F12 | 416880 | SAL | 0.2743444 | -5.8585327 |
| P16 cl119 | G01 | 1336600 | empty | 0.8796025 | -0.9720211 |
| P16 cl119 | G02 | 246300 | sample | 0.1620875 | -6.7648315 |
| P16 cl119 | G03 | 1483700 | sample | 0.9764075 | -0.1904726 |
| P16 cl119 | G04 | 1385100 | sample | 0.9115199 | -0.7143386 |
| P16 cl119 | G05 | 1302000 | sample | 0.8568326 | -1.1558524 |
| P16 cl119 | G06 | 1589900 | sample | 1.0462966 | 0.3737725 |
| P16 cl119 | G07 | 1527000 | sample | 1.0049028 | 0.0395822 |
| P16 cl119 | G08 | 1512000 | sample | 0.9950314 | -0.0401135 |
| P16 cl119 | G09 | 1432900 | sample | 0.9429765 | -0.4603751 |
| P16 cl119 | G10 | 1607800 | sample | 1.0580764 | 0.4688760 |
| P16 cl119 | G11 | 1518800 | sample | 0.9995064 | -0.0039848 |
| P16 cl119 | G12 | 366420 | SAL | 0.2411372 | -6.1266288 |
| P16 cl119 | H01 | 1312900 | empty | 0.8640058 | -1.0979403 |
| P16 cl119 | H02 | 1351200 | sample | 0.8892106 | -0.8944507 |
| P16 cl119 | H03 | 347980 | sample | 0.2290020 | -6.2246013 |
| P16 cl119 | H04 | 304930 | sample | 0.2006713 | -6.4533278 |
| P16 cl119 | H05 | 1464700 | sample | 0.9639038 | -0.2914204 |
| P16 cl119 | H06 | 1339100 | sample | 0.8812477 | -0.9587385 |
| P16 cl119 | H07 | 1285800 | sample | 0.8461716 | -1.2419237 |

*(continued)*

| Plate | Well | Value | Treatment | Norm | Score |
|---|---|---|---|---|---|
| P16 cl119 | H08 | 1239400 | sample | 0.8156362 | -1.4884489 |
| P16 cl119 | H09 | 1367500 | sample | 0.8999375 | -0.8078481 |
| P16 cl119 | H10 | 1302000 | sample | 0.8568326 | -1.1558524 |
| P16 cl119 | H11 | 1353600 | sample | 0.8907900 | -0.8816994 |
| P16 cl119 | H12 | 338880 | SAL | 0.2230134 | -6.2729500 |

#### Hitmap for the median based normalization

For the *median based normalization* a threshold at was chosen arbitrarily according to the positives controls.

#### Clone WT

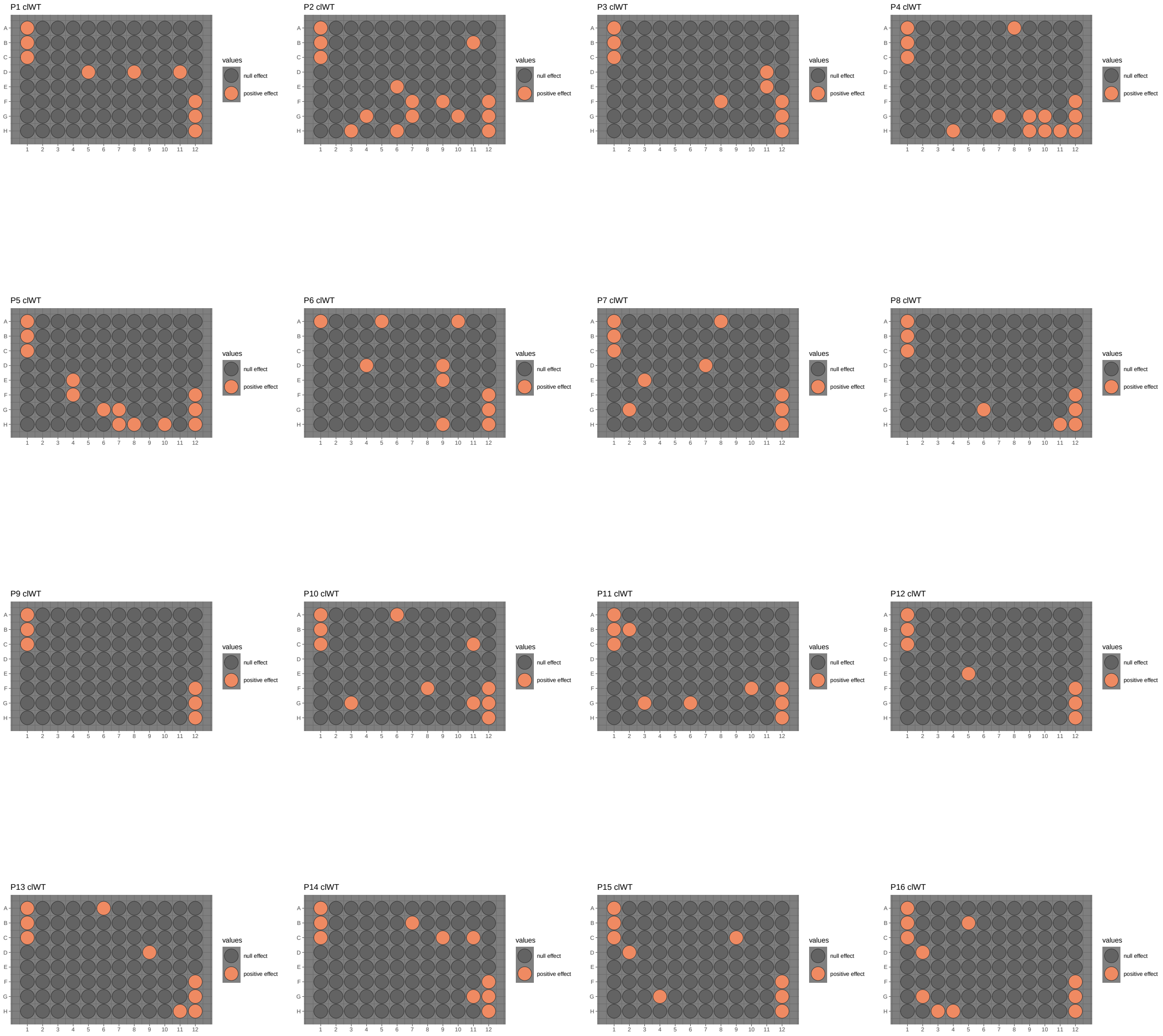

### Clone 101

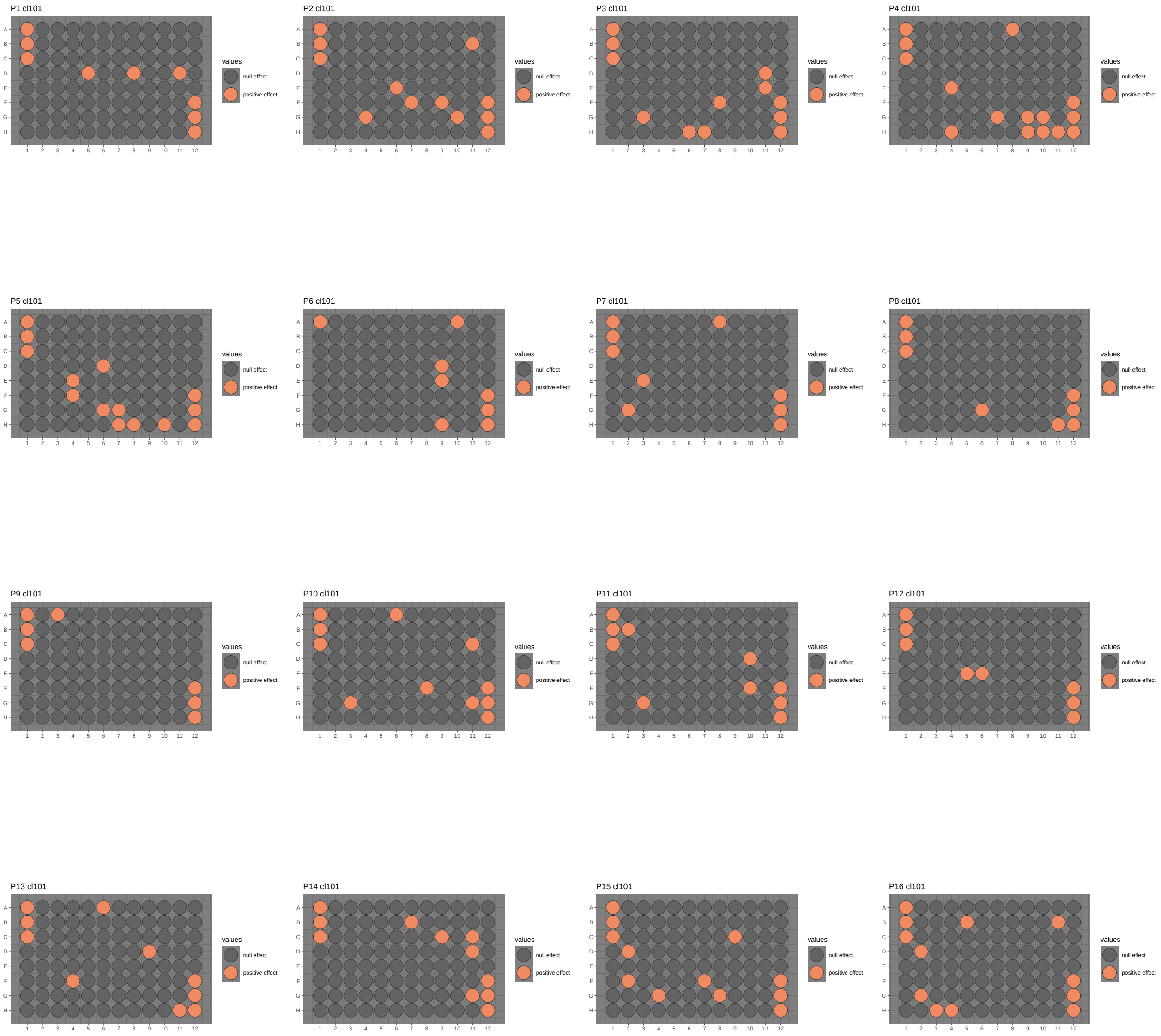
P1 cl101
P2 cl101
P3 cl101
P4 cl101
P5 cl101
P6 cl101
P7 cl101
P8 cl101
P9 cl101
P10 cl101
P11 cl101
P12 cl101
P13 cl101
P14 cl101
P15 cl101
P16 cl101
values
null effect
positive effect

### Clone 119

P1 cl119

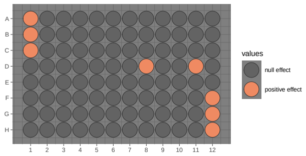


P2 cl119

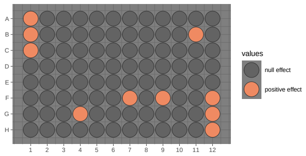


P3 cl119

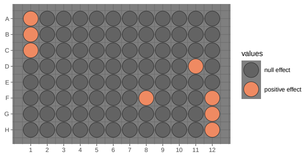


P4 cl119

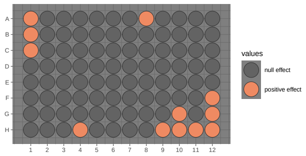


P5 cl119

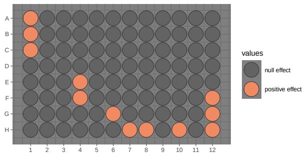


P6 cl119

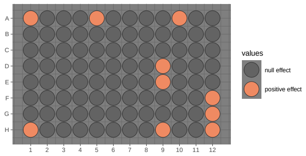


P7 cl119

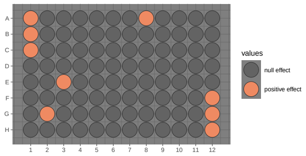


P8 cl119

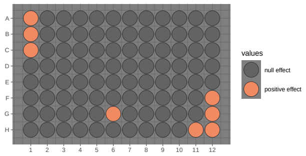


P9 cl119

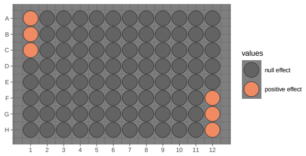


P10 cl119

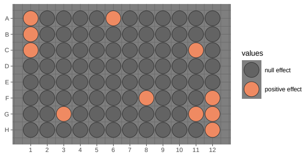


P11 cl119

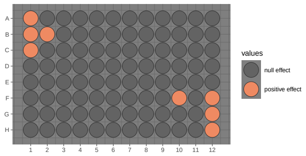


P12 cl119

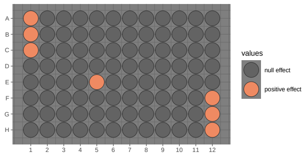


P13 cl119

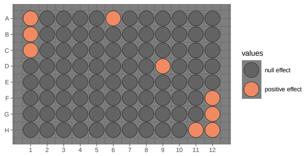


P14 cl119

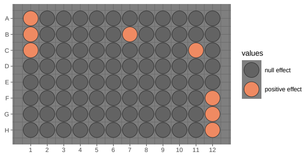


P15 cl119

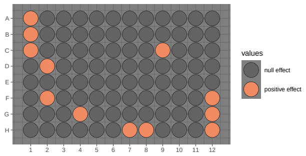


P16 cl119

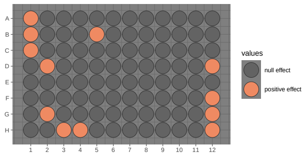

##### Hit list for the median based normalization

##### Clone WT

| PlateNumber__PositionNumber__96 | score | Chemical name | PlateNumber__PositionNumber__384 |
|---|---|---|---|
| 01D05 | -11.564834 | Triamterene | 1G9 |
| 01D08 | -16.317084 | Pyrimethamine | 1G15 |
| 01D11 | -14.783788 | Niclosamide | 1G21 |
| 02B11 | -7.934406 | Nocodazole | 1C22 |
| 02E06 | -5.513436 | Perphenazine | 1I12 |
| 02F07 | -10.059853 | Astemizole | 1K14 |
| 02F09 | -8.747999 | Terfenadine | 1K18 |
| 02G04 | -8.089970 | Chlorhexidine | 1M8 |
| 02G07 | -4.847568 | Tamoxifen citrate | 1M14 |
| 02G10 | -5.137318 | Thioproperazine dimesylate | 1M20 |
| 02H03 | -4.191178 | Chloroxine | 1O6 |
| 02H06 | -5.502320 | Paclitaxel | 1O12 |
| 03D11 | -12.226456 | Camptothecine (S,+) | 1H21 |
| 03E11 | -10.277019 | Fenbendazole | 1J21 |
| 03F08 | -11.926046 | Mebendazole | 1L15 |
| 04A08 | -8.538668 | Albendazole | 1B16 |
| 04G07 | -5.400979 | Clemastine fumarate | 1N14 |
| 04G09 | -7.131273 | Pimozide | 1N18 |
| 04G10 | -7.975864 | Amodiaquin dihydrochloride dihydrate | 1N20 |
| 04H04 | -11.217445 | Trifluoperazine dihydrochloride | 1P8 |
| 04H09 | -12.767181 | Quinacrine dihydrochloride hydrate | 1P18 |
| 04H10 | -6.328907 | Clofilium tosylate | 1P20 |
| 04H11 | -11.477563 | Fluphenazine dihydrochloride | 1P22 |
| 05E04 | -9.662776 | Colchicine | 2I7 |
| 05F04 | -12.844355 | Amethopterin (R,S) | 2K7 |
| 05G06 | -12.765172 | Mitoxantrone dihydrochloride | 2M11 |
| 05G07 | -6.391843 | GBR 12909 dihydrochloride | 2M13 |
| 05H07 | -6.738562 | Etoposide | 2O13 |
| 05H08 | -12.972022 | Clomiphene citrate (Z,E) | 2O15 |
| 05H10 | -11.270341 | Prochlorperazine dimaleate | 2O19 |
| 06A05 | -5.041428 | Idebenone | 2A10 |
| 06A10 | -7.638389 | Amiodarone hydrochloride | 2A20 |
| 06D04 | -5.665189 | Cyanocobalamin | 2G8 |
| 06D09 | -10.019558 | Doxorubicin hydrochloride | 2G18 |
| 06E09 | -9.096827 | Pitavastatin calcium | 2I18 |
| 06H09 | -4.539997 | Felodipine | 2O18 |
| 07A08 | -17.314677 | Daunorubicin hydrochloride | 2B15 |
| 07D07 | -9.443219 | Lovastatin | 2H13 |
| 07E03 | -15.462207 | Thiostrepton | 2J5 |
| 07G02 | -16.809447 | Ciclopirox ethanolamine | 2N3 |
| 08G06 | -13.772521 | Cytarabine | 2N12 |
| 08H11 | -16.971566 | Sertindole | 2P22 |
| 10A06 | -16.379848 | Deferoxamine mesylate | 3A12 |
| 10C11 | -20.050253 | Dronedarone hydrochloride | 3E22 |
| 10F08 | -21.803583 | Alexidine dihydrochloride | 3K16 |
| 10G03 | -20.294658 | Podophyllotoxin | 3M6 |
| 10G11 | -23.353789 | Cycloheximide | 3M22 |
| 11B02 | -15.673366 | Auranofin | 3D3 |
| 11F10 | -12.457392 | Fluvastatin sodium salt | 3L19 |
| 11G03 | -10.360829 | Raloxifene hydrochloride | 3N5 |
| 11G06 | -10.763388 | Simvastatin | 3N11 |
| 12E05 | -12.589553 | Posaconazole | 3J10 |
| 13A06 | -21.705433 | Floxuridine | 4A11 |
| 13D09 | -19.874769 | Zuclopenthixol dihydrochloride | 4G17 |
| 13H11 | -25.727509 | Pyrvinium pamoate | 4O21 |
| 14B07 | -16.802647 | Trifluridine | 4C14 |
| 14C09 | -16.487718 | Thiethylperazine dimalate | 4E18 |
| 14C11 | -19.055113 | Vorinostat | 4E22 |
| 14G11 | -15.732455 | Parbendazole | 4M22 |

*(continued)*

| PlateNumber__PositionNumber__96 | score | Chemical name | PlateNumber__PositionNumber__384 |
|---|---|---|---|
| 15C09 | -18.053010 | Cladribine | 4F17 |
| 15D02 | -14.889960 | 5-fluorouracil | 4H3 |
| 15G04 | -18.329320 | Gemcitabine | 4N7 |
| 16B05 | -18.068195 | Tegaserod maleate | 4D10 |
| 16D02 | -14.959508 | Aminacrine | 4H4 |
| 16G02 | -16.454619 | Epirubicin hydrochloride | 4N4 |
| 16H03 | -14.149764 | Pemetrexed disodium | 4P6 |
| 16H04 | -15.460440 | Raltitrexed | 4P8 |

#### Clone 101

| PlateNumber__PositionNumber__96 | score | Chemical name | PlateNumber__PositionNumber__384 |
|---|---|---|---|
| 01D05 | -9.827514 | Triamterene | 1G9 |
| 01D08 | -13.214560 | Pyrimethamine | 1G15 |
| 01D11 | -13.662821 | Niclosamide | 1G21 |
| 02B11 | -8.716801 | Nocodazole | 1C22 |
| 02E06 | -7.948564 | Perphenazine | 1I12 |
| 02F07 | -11.109444 | Astemizole | 1K14 |
| 02F09 | -11.246280 | Terfenadine | 1K18 |
| 02G04 | -11.113043 | Chlorhexidine | 1M8 |
| 02G10 | -6.408954 | Thioproperazine dimesylate | 1M20 |
| 03D11 | -6.223312 | Camptothecine (S,+) | 1H21 |
| 03E11 | -4.594220 | Fenbendazole | 1J21 |
| 03F08 | -5.335434 | Mebendazole | 1L15 |
| 03G03 | -4.334846 | Antimycin A | 1N5 |
| 03H06 | -4.191029 | Carmofur | 1P11 |
| 03H07 | -4.174483 | Dilazep dihydrochloride | 1P13 |
| 04A08 | -5.654045 | Albendazole | 1B16 |
| 04E04 | -4.928705 | Homochlorcyclizine dihydrochloride | 1J8 |
| 04G07 | -4.883150 | Clemastine fumarate | 1N14 |
| 04G09 | -4.907591 | Pimozide | 1N18 |
| 04G10 | -5.131425 | Amodiaquin dihydrochloride dihydrate | 1N20 |
| 04H04 | -8.273120 | Trifluoperazine dihydrochloride | 1P8 |
| 04H09 | -9.325285 | Quinacrine dihydrochloride hydrate | 1P18 |
| 04H10 | -4.737186 | Clofilium tosylate | 1P20 |
| 04H11 | -8.535855 | Fluphenazine dihydrochloride | 1P22 |
| 05D06 | -5.743307 | Hycanthone | 2G11 |
| 05E04 | -7.345533 | Colchicine | 2I7 |
| 05F04 | -9.696188 | Amethopterin (R,S) | 2K7 |
| 05G06 | -9.636030 | Mitoxantrone dihydrochloride | 2M11 |
| 05G07 | -5.452073 | GBR 12909 dihydrochloride | 2M13 |
| 05H07 | -5.344811 | Etoposide | 2O13 |
| 05H08 | -9.880816 | Clomiphene citrate (Z,E) | 2O15 |
| 05H10 | -8.713254 | Prochlorperazine dimaleate | 2O19 |
| 06A10 | -9.589215 | Amiodarone hydrochloride | 2A20 |
| 06D09 | -12.771789 | Doxorubicin hydrochloride | 2G18 |
| 06E09 | -6.798254 | Pitavastatin calcium | 2I18 |
| 06H09 | -8.697456 | Felodipine | 2O18 |
| 07A08 | -10.256859 | Daunorubicin hydrochloride | 2B15 |
| 07E03 | -9.848150 | Thiostrepton | 2J5 |
| 07G02 | -9.955283 | Ciclopirox ethanolamine | 2N3 |
| 08G06 | -11.940727 | Cytarabine | 2N12 |
| 08H11 | -15.401582 | Sertindole | 2P22 |
| 09A03 | -10.845546 | Gefitinib | 3A5 |
| 10A06 | -7.501786 | Deferoxamine mesylate | 3A12 |
| 10C11 | -8.221544 | Dronedarone hydrochloride | 3E22 |
| 10F08 | -9.708469 | Alexidine dihydrochloride | 3K16 |
| 10G03 | -8.359076 | Podophyllotoxin | 3M6 |
| 10G11 | -10.086497 | Cycloheximide | 3M22 |
| 11B02 | -17.233320 | Auranofin | 3D3 |
| 11D10 | -16.694577 | Ganciclovir | 3H19 |
| 11F10 | -13.959517 | Fluvastatin sodium salt | 3L19 |
| 11G03 | -11.353424 | Raloxifene hydrochloride | 3N5 |
| 12E05 | -9.430853 | Posaconazole | 3J10 |
| 12E06 | -9.864021 | Thonzonium bromide | 3J12 |
| 13A06 | -11.088702 | Floxuridine | 4A11 |
| 13D09 | -10.232316 | Zuclopenthixol dihydrochloride | 4G17 |
| 13F04 | -7.313400 | Deptropine citrate | 4K7 |
| 13H11 | -13.081335 | Pyrvinium pamoate | 4O21 |
| 14B07 | -6.908868 | Trifluridine | 4C14 |
| 14C09 | -5.213765 | Thiethylperazine dimalate | 4E18 |
| 14C11 | -10.020738 | Vorinostat | 4E22 |
| 14D11 | -5.473225 | Methiazole | 4G22 |

*(continued)*

| PlateNumber__PositionNumber__96 | score | Chemical name | PlateNumber__PositionNumber__384 |
|---|---|---|---|
| 14G11 | -6.808874 | Parbendazole | 4M22 |
| 15C09 | -10.310047 | Cladribine | 4F17 |
| 15D02 | -8.097124 | 5-fluorouracil | 4H3 |
| 15F02 | -9.147820 | Topotecan | 4L3 |
| 15F07 | -6.072843 | Benztropine mesylate | 4L13 |
| 15G04 | -10.791852 | Gemcitabine | 4N7 |
| 15G08 | -6.371668 | Docetaxel | 4N15 |
| 16B05 | -8.806575 | Tegaserod maleate | 4D10 |
| 16B11 | -4.532988 | Estramustine | 4D22 |
| 16D02 | -7.309353 | Aminacrine | 4H4 |
| 16G02 | -7.663125 | Epirubicin hydrochloride | 4N4 |
| 16H03 | -7.086188 | Pemetrexed disodium | 4P6 |
| 16H04 | -7.584274 | Raltitrexed | 4P8 |

#### Clone 119

| PlateNumber_PositionNumber_96 | score | Chemical name | PlateNumber_PositionNumber_384 |
|---|---|---|---|
| 01D08 | -8.001103 | Pyrimethamine | 1G15 |
| 01D11 | -8.390415 | Niclosamide | 1G21 |
| 02B11 | -3.725538 | Nocodazole | 1C22 |
| 02F07 | -5.055972 | Astemizole | 1K14 |
| 02F09 | -4.799566 | Terfenadine | 1K18 |
| 02G04 | -4.224700 | Chlorhexidine | 1M8 |
| 03D11 | -5.709235 | Camptothecine (S,+) | 1H21 |
| 03F08 | -4.218859 | Mebendazole | 1L15 |
| 04A08 | -4.087522 | Albendazole | 1B16 |
| 04G10 | -3.043978 | Amodiaquin dihydrochloride dihydrate | 1N20 |
| 04H04 | -4.576428 | Trifluoperazine dihydrochloride | 1P8 |
| 04H09 | -5.869316 | Quinacrine dihydrochloride hydrate | 1P18 |
| 04H10 | -3.074822 | Clofilium tosylate | 1P20 |
| 04H11 | -4.767749 | Fluphenazine dihydrochloride | 1P22 |
| 05E04 | -3.454363 | Colchicine | 2I7 |
| 05F04 | -5.281988 | Amethopterin (R,S) | 2K7 |
| 05G06 | -5.232155 | Mitoxantrone dihydrochloride | 2M11 |
| 05H07 | -3.054473 | Etoposide | 2O13 |
| 05H08 | -5.291126 | Clomiphene citrate (Z,E) | 2O15 |
| 05H10 | -4.063901 | Prochlorperazine dimaleate | 2O19 |
| 06A05 | -3.291986 | Idebenone | 2A10 |
| 06A10 | -3.937398 | Amiodarone hydrochloride | 2A20 |
| 06D09 | -5.673127 | Doxorubicin hydrochloride | 2G18 |
| 06E09 | -3.137130 | Pitavastatin calcium | 2I18 |
| 06H09 | -4.180960 | Felodipine | 2O18 |
| 07A08 | -6.370301 | Daunorubicin hydrochloride | 2B15 |
| 07E03 | -5.762711 | Thiostrepton | 2J5 |
| 07G02 | -6.289685 | Ciclopirox ethanolamine | 2N3 |
| 08G06 | -4.697005 | Cytarabine | 2N12 |
| 08H11 | -5.511178 | Sertindole | 2P22 |
| 10A06 | -5.285447 | Deferoxamine mesylate | 3A12 |
| 10C11 | -4.565308 | Dronedarone hydrochloride | 3E22 |
| 10F08 | -5.320242 | Alexidine dihydrochloride | 3K16 |
| 10G03 | -4.959530 | Podophyllotoxin | 3M6 |
| 10G11 | -5.712561 | Cycloheximide | 3M22 |
| 11B02 | -10.199906 | Auranofin | 3D3 |
| 11F10 | -7.469432 | Fluvastatin sodium salt | 3L19 |
| 12E05 | -5.231768 | Posaconazole | 3J10 |
| 13A06 | -6.786501 | Floxuridine | 4A11 |
| 13D09 | -5.479877 | Zuclopenthixol dihydrochloride | 4G17 |
| 13H11 | -8.071063 | Pyrvinium pamoate | 4O21 |
| 14B07 | -5.303187 | Trifluridine | 4C14 |
| 14C11 | -6.901168 | Vorinostat | 4E22 |
| 15C09 | -5.692055 | Cladribine | 4F17 |
| 15D02 | -4.648340 | 5-fluorouracil | 4H3 |
| 15F02 | -4.805495 | Topotecan | 4L3 |
| 15G04 | -5.677296 | Gemcitabine | 4N7 |
| 15H07 | -6.746032 | Rosiglitazone Hydrochloride | 4P13 |
| 15H08 | -6.774619 | Rivastigmine | 4P15 |
| 16B05 | -7.557415 | Tegaserod maleate | 4D10 |
| 16D02 | -6.253876 | Aminacrine | 4H4 |
| 16G02 | -6.764831 | Epirubicin hydrochloride | 4N4 |
| 16H03 | -6.224601 | Pemetrexed disodium | 4P6 |
| 16H04 | -6.453328 | Raltitrexed | 4P8 |

#### Common Hit list for the median based normalization

| PlateNumber__PositionNumber__96 | Chemical name | PlateNumber__PositionNumber__384 |
|---|---|---|
| 01D08 | Pyrimethamine | 1G15 |
| 01D11 | Niclosamide | 1G21 |
| 02B11 | Nocodazole | 1C22 |
| 02F07 | Astemizole | 1K14 |
| 02F09 | Terfenadine | 1K18 |
| 02G04 | Chlorhexidine | 1M8 |
| 03D11 | Camptothecine (S,+) | 1H21 |
| 03F08 | Mebendazole | 1L15 |
| 04A08 | Albendazole | 1B16 |
| 04G10 | Amodiaquin dihydrochloride dihydrate | 1N20 |
| 04H04 | Trifluoperazine dihydrochloride | 1P8 |
| 04H09 | Quinacrine dihydrochloride hydrate | 1P18 |
| 04H10 | Clofilium tosylate | 1P20 |
| 04H11 | Fluphenazine dihydrochloride | 1P22 |
| 05E04 | Colchicine | 2I7 |
| 05F04 | Amethopterin (R,S) | 2K7 |
| 05G06 | Mitoxantrone dihydrochloride | 2M11 |
| 05H07 | Etoposide | 2O13 |
| 05H08 | Clomiphene citrate (Z,E) | 2O15 |
| 05H10 | Prochlorperazine dimaleate | 2O19 |
| 06A10 | Amiodarone hydrochloride | 2A20 |
| 06D09 | Doxorubicin hydrochloride | 2G18 |
| 06E09 | Pitavastatin calcium | 2I18 |
| 06H09 | Felodipine | 2O18 |
| 07A08 | Daunorubicin hydrochloride | 2B15 |
| 07E03 | Thiostrepton | 2J5 |
| 07G02 | Ciclopirox ethanolamine | 2N3 |
| 08G06 | Cytarabine | 2N12 |
| 08H11 | Sertindole | 2P22 |
| 10A06 | Deferoxamine mesylate | 3A12 |
| 10C11 | Dronedarone hydrochloride | 3E22 |
| 10F08 | Alexidine dihydrochloride | 3K16 |
| 10G03 | Podophyllotoxin | 3M6 |
| 10G11 | Cycloheximide | 3M22 |
| 11B02 | Auranofin | 3D3 |
| 11F10 | Fluvastatin sodium salt | 3L19 |
| 12E05 | Posaconazole | 3J10 |
| 13A06 | Floxuridine | 4A11 |
| 13D09 | Zuclopenthixol dihydrochloride | 4G17 |
| 13H11 | Pyrvinium pamoate | 4O21 |
| 14B07 | Trifluridine | 4C14 |
| 14C11 | Vorinostat | 4E22 |
| 15C09 | Cladribine | 4F17 |
| 15D02 | 5-fluorouracil | 4H3 |
| 15G04 | Gemcitabine | 4N7 |
| 16B05 | Tegaserod maleate | 4D10 |
| 16D02 | Aminacrine | 4H4 |
| 16G02 | Epirubicin hydrochloride | 4N4 |
| 16H03 | Pemetrexed disodium | 4P6 |
| 16H04 | Raltitrexed | 4P8 |

#### Hitmap for the normalized percent inhibition

For the *normalized percent inhibition* a threshold of 40% has been chosen.

#### Clone WT

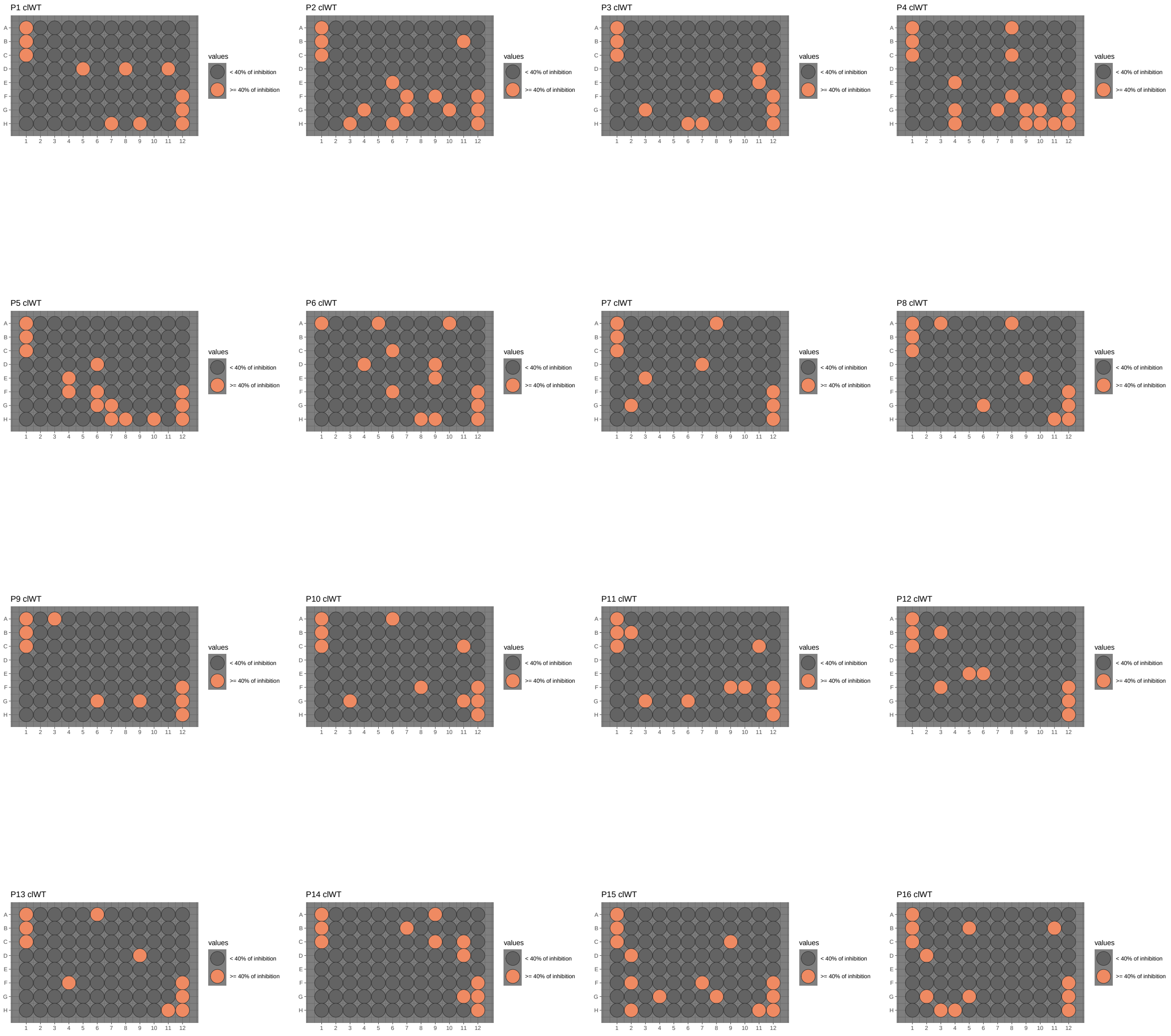

### Clone 101

P1 cl101

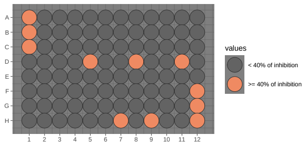


P2 cl101

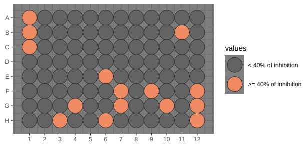


P3 cl101

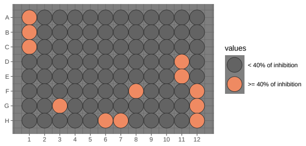


P4 cl101

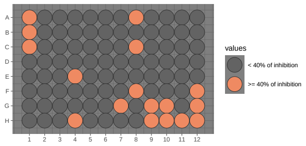


P5 cl101

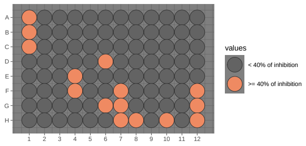


P6 cl101

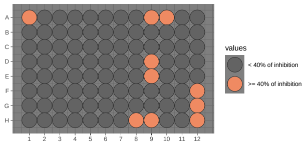


P7 cl101

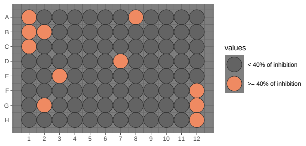


P8 cl101

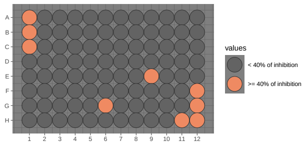


P9 cl101

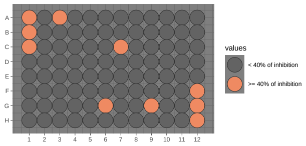


P10 cl101

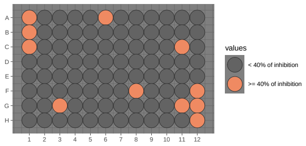


P11 cl101

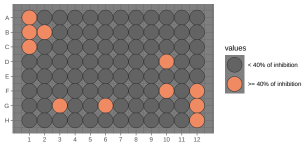


P12 cl101

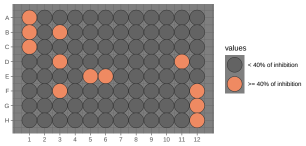


P13 cl101

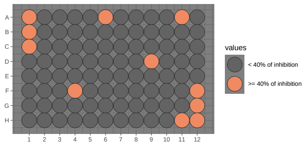


P14 cl101

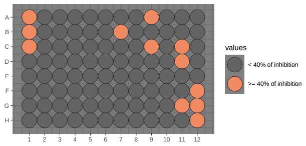


P15 cl101

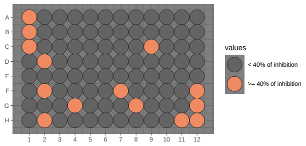


P16 cl101

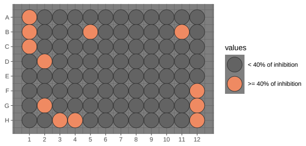

### Clone 119

P1 cl119

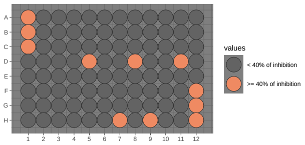


P2 cl119

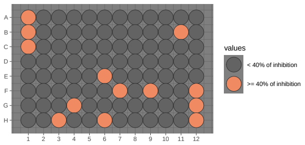


P3 cl119

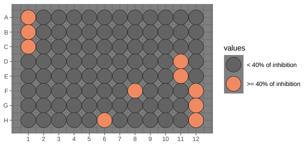


P4 cl119

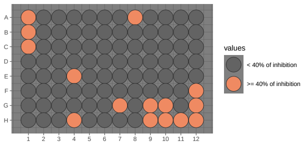


P5 cl119

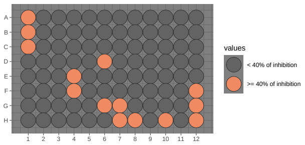


P6 cl119

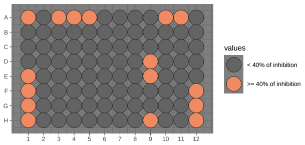


P7 cl119

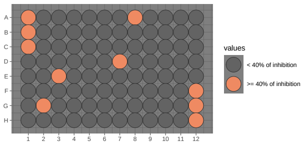


P8 cl119

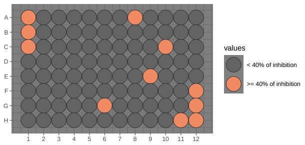


P9 cl119

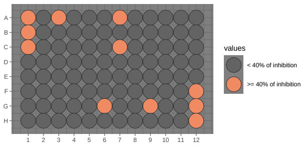


P10 cl119

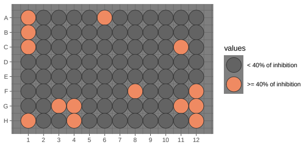


P11 cl119

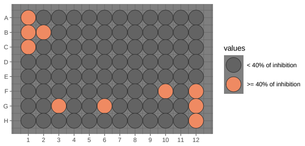


P12 cl119

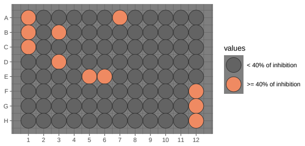


P13 cl119

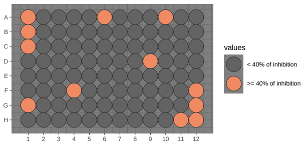


P14 cl119

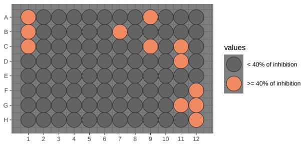


P15 cl119

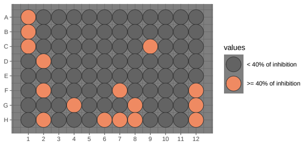


P16 cl119

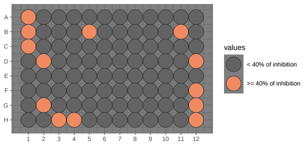

#### Hit list for the normalized percent inhibition

#### Clone WT

| PlateNumber_PositionNumber_96 | norm | Chemical name | PlateNumber_PositionNumber_384 |
|---|---|---|---|
| 01D05 | 83.73947 | Triamterene | 1G9 |
| 01D08 | 118.03921 | Pyrimethamine | 1G15 |
| 01D11 | 106.97252 | Niclosamide | 1G21 |
| 01H07 | 47.72804 | Dibucaine | 1O13 |
| 01H09 | 45.64403 | Thioridazine hydrochloride | 1O17 |
| 02B11 | 94.34121 | Nocodazole | 1C22 |
| 02E06 | 64.13364 | Perphenazine | 1I12 |
| 02F07 | 120.86142 | Astemizole | 1K14 |
| 02F09 | 104.49280 | Terfenadine | 1K18 |
| 02G04 | 96.28226 | Chlorhexidine | 1M8 |
| 02G07 | 55.82529 | Tamoxifen citrate | 1M14 |
| 02G10 | 59.44063 | Thioproperazine dimesylate | 1M20 |
| 02H03 | 47.63519 | Chloroxine | 1O6 |
| 02H06 | 63.99493 | Paclitaxel | 1O12 |
| 03D11 | 86.00897 | Camptothecine (S,+) | 1H21 |
| 03E11 | 72.00904 | Fenbendazole | 1J21 |
| 03F08 | 83.85157 | Mebendazole | 1L15 |
| 03G03 | 56.80296 | Antimycin A | 1N5 |
| 03H06 | 59.28270 | Carmofur | 1P11 |
| 03H07 | 51.32053 | Dilazep dihydrochloride | 1P13 |
| 04A08 | 88.45771 | Albendazole | 1B16 |
| 04C08 | 54.25708 | Clotrimazole | 1F16 |
| 04E04 | 51.26197 | Homochlorcyclizine dihydrochloride | 1J8 |
| 04F08 | 52.07387 | Benzydamine hydrochloride | 1L16 |
| 04G04 | 41.15377 | Telenzepine dihydrochloride | 1N8 |
| 04G07 | 58.71443 | Clemastine fumarate | 1N14 |
| 04G09 | 75.11651 | Pimozide | 1N18 |
| 04G10 | 83.12269 | Amodiaquin dihydrochloride dihydrate | 1N20 |
| 04H04 | 113.85080 | Trifluoperazine dihydrochloride | 1P8 |
| 04H09 | 128.54131 | Quinacrine dihydrochloride hydrate | 1P18 |
| 04H10 | 67.51060 | Clofilium tosylate | 1P20 |
| 04H11 | 116.31656 | Fluphenazine dihydrochloride | 1P22 |
| 05D06 | 55.49513 | Hycanthone | 2G11 |
| 05E04 | 93.39350 | Colchicine | 2I7 |
| 05F04 | 126.13440 | Amethopterin (R,S) | 2K7 |
| 05F06 | 46.87698 | Methiothepin maleate | 2K11 |
| 05G06 | 125.31955 | Mitoxantrone dihydrochloride | 2M11 |
| 05G07 | 59.73309 | GBR 12909 dihydrochloride | 2M13 |
| 05H07 | 63.30110 | Etoposide | 2O13 |
| 05H08 | 127.44819 | Clomiphene citrate (Z,E) | 2O15 |
| 05H10 | 109.93658 | Prochlorperazine dimaleate | 2O19 |
| 06A05 | 79.74603 | Idebenone | 2A10 |
| 06A10 | 119.42875 | Amiodarone hydrochloride | 2A20 |
| 06C06 | 41.30057 | Chicago sky blue 6B | 2E12 |
| 06D04 | 89.27738 | Cyanocobalamin | 2G8 |
| 06D09 | 155.81408 | Doxorubicin hydrochloride | 2G18 |
| 06E09 | 141.71434 | Pitavastatin calcium | 2I18 |
| 06F06 | 44.71085 | Mebhydroline 1,5-naphtalenedisulfonate | 2K12 |
| 06H08 | 40.84664 | Progesterone | 2O16 |
| 06H09 | 72.08394 | Felodipine | 2O18 |
| 07A08 | 126.59544 | Daunorubicin hydrochloride | 2B15 |
| 07D07 | 67.75679 | Lovastatin | 2H13 |
| 07E03 | 112.74835 | Thiostrepton | 2J5 |
| 07G02 | 122.81888 | Ciclopirox ethanolamine | 2N3 |
| 08A03 | 46.96320 | Mevastatin | 2B6 |
| 08A08 | 46.49373 | Itraconazole | 2B16 |
| 08E09 | 58.25900 | Oxibendazol | 2J18 |
| 08G06 | 103.25840 | Cytarabine | 2N12 |
| 08H11 | 126.52477 | Sertindole | 2P22 |

*(continued)*

| PlateNumber__PositionNumber__96 | norm | Chemical name | PlateNumber__PositionNumber__384 |
|---|---|---|---|
| 09A03 | 73.16130 | Gefitinib | 3A5 |
| 09G06 | 50.42801 | Methyl benzethonium chloride | 3M11 |
| 09G09 | 56.15164 | Benzethonium chloride | 3M17 |
| 10A06 | 82.14846 | Deferoxamine mesylate | 3A12 |
| 10C11 | 98.99028 | Dronedarone hydrochloride | 3E22 |
| 10F08 | 107.03552 | Alexidine dihydrochloride | 3K16 |
| 10G03 | 100.11175 | Podophyllotoxin | 3M6 |
| 10G11 | 114.14872 | Cycloheximide | 3M22 |
| 11B02 | 128.48855 | Auranofin | 3D3 |
| 11C11 | 41.53060 | Ethaverine hydrochloride | 3F21 |
| 11F09 | 44.14922 | Doxazosin mesylate | 3L17 |
| 11F10 | 103.38490 | Fluvastatin sodium salt | 3L19 |
| 11G03 | 87.01929 | Raloxifene hydrochloride | 3N5 |
| 11G06 | 90.16164 | Simvastatin | 3N11 |
| 12B03 | 67.50988 | Flubendazol | 3D6 |
| 12E05 | 77.00806 | Posaconazole | 3J10 |
| 12E06 | 67.27153 | Thonzonium bromide | 3J12 |
| 12F03 | 45.46412 | S(-)Eticlopride hydrochloride | 3L6 |
| 13A06 | 109.11184 | Floxuridine | 4A11 |
| 13D09 | 100.44292 | Zuclopenthixol dihydrochloride | 4G17 |
| 13F04 | 74.62324 | Deptropine citrate | 4K7 |
| 13H11 | 128.15796 | Pyrvinium pamoate | 4O21 |
| 14A09 | 48.70781 | Piperacetazine | 4A18 |
| 14B07 | 88.21279 | Trifluridine | 4C14 |
| 14C09 | 86.48377 | Thiethylperazine dimalate | 4E18 |
| 14C11 | 100.57923 | Vorinostat | 4E22 |
| 14D11 | 61.15198 | Methiazole | 4G22 |
| 14G11 | 82.33724 | Parbendazole | 4M22 |
| 15C09 | 112.85210 | Cladribine | 4F17 |
| 15D02 | 94.25132 | 5-fluorouracil | 4H3 |
| 15F02 | 66.68493 | Topotecan | 4L3 |
| 15F07 | 70.27389 | Benztropine mesylate | 4L13 |
| 15G04 | 114.47699 | Gemcitabine | 4N7 |
| 15G08 | 67.14542 | Docetaxel | 4N15 |
| 15H02 | 58.18860 | Imatinib | 4P3 |
| 15H11 | 47.74576 | Hexachlorophene | 4P21 |
| 16B05 | 127.08771 | Tegaserod maleate | 4D10 |
| 16B11 | 61.49254 | Estramustine | 4D22 |
| 16D02 | 106.20609 | Aminacrine | 4H4 |
| 16G02 | 116.24903 | Epirubicin hydrochloride | 4N4 |
| 16G05 | 65.52290 | Lomerizine hydrochloride | 4N10 |
| 16H03 | 100.76689 | Pemetrexed disodium | 4P6 |
| 16H04 | 109.57094 | Raltitrexed | 4P8 |

#### Clone 101

| PlateNumber_PositionNumber_96 | norm | Chemical name | PlateNumber_PositionNumber_384 |
|---|---|---|---|
| 01D05 | 79.12048 | Triamterene | 1G9 |
| 01D08 | 108.63059 | Pyrimethamine | 1G15 |
| 01D11 | 112.53613 | Niclosamide | 1G21 |
| 01H07 | 49.39999 | Dibucaine | 1O13 |
| 01H09 | 55.54641 | Thioridazine hydrochloride | 1O17 |
| 02B11 | 87.67199 | Nocodazole | 1C22 |
| 02E06 | 79.72584 | Perphenazine | 1I12 |
| 02F07 | 112.41995 | Astemizole | 1K14 |
| 02F09 | 113.83529 | Terfenadine | 1K18 |
| 02G04 | 112.45717 | Chlorhexidine | 1M8 |
| 02G07 | 58.90514 | Tamoxifen citrate | 1M14 |
| 02G10 | 63.80109 | Thioproperazine dimesylate | 1M20 |
| 02H03 | 41.98975 | Chloroxine | 1O6 |
| 02H06 | 58.91277 | Paclitaxel | 1O12 |
| 03D11 | 94.10429 | Camptothecine (S,+) | 1H21 |
| 03E11 | 67.71781 | Fenbendazole | 1J21 |
| 03F08 | 79.72329 | Mebendazole | 1L15 |
| 03G03 | 63.51672 | Antimycin A | 1N5 |
| 03H06 | 61.18732 | Carmofur | 1P11 |
| 03H07 | 60.91932 | Dilazep dihydrochloride | 1P13 |
| 04A08 | 78.25296 | Albendazole | 1B16 |
| 04C08 | 42.39925 | Clotrimazole | 1F16 |
| 04E04 | 68.23829 | Homochlorcyclizine dihydrochloride | 1J8 |
| 04F08 | 48.44798 | Benzydamine hydrochloride | 1L16 |
| 04G07 | 67.60933 | Clemastine fumarate | 1N14 |
| 04G09 | 67.94678 | Pimozide | 1N18 |
| 04G10 | 71.03723 | Amodiaquin dihydrochloride dihydrate | 1N20 |
| 04H04 | 114.41413 | Trifluoperazine dihydrochloride | 1P8 |
| 04H09 | 128.94121 | Quinacrine dihydrochloride hydrate | 1P18 |
| 04H10 | 65.59402 | Clofilium tosylate | 1P20 |
| 04H11 | 118.04168 | Fluphenazine dihydrochloride | 1P22 |
| 05D06 | 70.24988 | Hycanthone | 2G11 |
| 05E04 | 91.17183 | Colchicine | 2I7 |
| 05F04 | 121.86682 | Amethopterin (R,S) | 2K7 |
| 05F07 | 50.30485 | Clofazimine | 2K13 |
| 05G06 | 121.08128 | Mitoxantrone dihydrochloride | 2M11 |
| 05G07 | 66.44693 | GBR 12909 dihydrochloride | 2M13 |
| 05H07 | 65.04629 | Etoposide | 2O13 |
| 05H08 | 124.27770 | Clomiphene citrate (Z,E) | 2O15 |
| 05H10 | 109.03162 | Prochlorperazine dimaleate | 2O19 |
| 06A09 | 42.64325 | Butoconazole nitrate | 2A18 |
| 06A10 | 127.42572 | Amiodarone hydrochloride | 2A20 |
| 06D09 | 171.51736 | Doxorubicin hydrochloride | 2G18 |
| 06E09 | 88.75952 | Pitavastatin calcium | 2I18 |
| 06H08 | 40.47704 | Progesterone | 2O16 |
| 06H09 | 115.07123 | Felodipine | 2O18 |
| 07A08 | 116.91037 | Daunorubicin hydrochloride | 2B15 |
| 07B02 | 47.19836 | Metixene hydrochloride | 2D3 |
| 07D07 | 54.77119 | Lovastatin | 2H13 |
| 07E03 | 112.23227 | Thiostrepton | 2J5 |
| 07G02 | 113.45851 | Ciclopirox ethanolamine | 2N3 |
| 08E09 | 60.01743 | Oxibendazol | 2J18 |
| 08G06 | 95.33591 | Cytarabine | 2N12 |
| 08H11 | 123.51935 | Sertindole | 2P22 |
| 09A03 | 79.30716 | Gefitinib | 3A5 |
| 09C07 | 42.63809 | Nisoldipine | 3E13 |
| 09G06 | 53.30206 | Methyl benzethonium chloride | 3M11 |
| 09G09 | 54.40884 | Benzethonium chloride | 3M17 |
| 10A06 | 82.54196 | Deferoxamine mesylate | 3A12 |
| 10C11 | 90.26983 | Dronedarone hydrochloride | 3E22 |
| 10F08 | 106.23458 | Alexidine dihydrochloride | 3K16 |

*(continued)*

| PlateNumber__PositionNumber__96 | norm | Chemical name | PlateNumber__PositionNumber__384 |
|---|---|---|---|
| 10G03 | 91.74648 | Podophyllotoxin | 3M6 |
| 10G11 | 110.29337 | Cycloheximide | 3M22 |
| 11B02 | 130.34943 | Auranofin | 3D3 |
| 11D10 | 126.16411 | Ganciclovir | 3H19 |
| 11F10 | 104.91632 | Fluvastatin sodium salt | 3L19 |
| 11G03 | 84.67044 | Raloxifene hydrochloride | 3N5 |
| 11G06 | 71.62761 | Simvastatin | 3N11 |
| 12B03 | 70.71231 | Flubendazol | 3D6 |
| 12D03 | 60.05857 | Monobenzone | 3H6 |
| 12D11 | 47.81443 | Gemifloxacin mesylate | 3H22 |
| 12E05 | 78.59745 | Posaconazole | 3J10 |
| 12E06 | 81.98392 | Thonzonium bromide | 3J12 |
| 12F03 | 43.45543 | S(-)Eticlopride hydrochloride | 3L6 |
| 13A06 | 106.33508 | Floxuridine | 4A11 |
| 13A11 | 44.18048 | Indatraline hydrochloride | 4A21 |
| 13D09 | 98.60378 | Zuclopenthixol dihydrochloride | 4G17 |
| 13F04 | 72.25237 | Deptropine citrate | 4K7 |
| 13H11 | 124.32419 | Pyrvinium pamoate | 4O21 |
| 14A09 | 43.59990 | Piperacetazine | 4A18 |
| 14B07 | 82.59355 | Trifluridine | 4C14 |
| 14C09 | 61.91432 | Thiethylperazine dimalate | 4E18 |
| 14C11 | 120.55648 | Vorinostat | 4E22 |
| 14D11 | 65.07957 | Methiazole | 4G22 |
| 14G11 | 81.37368 | Parbendazole | 4M22 |
| 15C09 | 107.73436 | Cladribine | 4F17 |
| 15D02 | 84.71267 | 5-fluorouracil | 4H3 |
| 15F02 | 95.64337 | Topotecan | 4L3 |
| 15F07 | 63.65348 | Benztropine mesylate | 4L13 |
| 15G04 | 112.74671 | Gemcitabine | 4N7 |
| 15G08 | 66.76224 | Docetaxel | 4N15 |
| 15H02 | 50.83415 | Imatinib | 4P3 |
| 15H11 | 41.60509 | Hexachlorophene | 4P21 |
| 16B05 | 125.80984 | Tegaserod maleate | 4D10 |
| 16B11 | 65.48225 | Estramustine | 4D22 |
| 16D02 | 104.67448 | Aminacrine | 4H4 |
| 16G02 | 109.66845 | Epirubicin hydrochloride | 4N4 |
| 16H03 | 101.52420 | Pemetrexed disodium | 4P6 |
| 16H04 | 108.55537 | Raltitrexed | 4P8 |

#### Clone 119

| PlateNumber__PositionNumber__96 | norm | Chemical name | PlateNumber__PositionNumber__384 |
|---|---|---|---|
| 01D05 | 66.44099 | Triamterene | 1G9 |
| 01D08 | 102.52533 | Pyrimethamine | 1G15 |
| 01D11 | 108.30052 | Niclosamide | 1G21 |
| 01H07 | 42.90759 | Dibucaine | 1O13 |
| 01H09 | 56.52725 | Thioridazine hydrochloride | 1O17 |
| 02B11 | 77.66857 | Nocodazole | 1C22 |
| 02E06 | 42.50339 | Perphenazine | 1I12 |
| 02F07 | 115.20694 | Astemizole | 1K14 |
| 02F09 | 107.97241 | Terfenadine | 1K18 |
| 02G04 | 91.75249 | Chlorhexidine | 1M8 |
| 02H03 | 50.83588 | Chloroxine | 1O6 |
| 02H06 | 41.81870 | Paclitaxel | 1O12 |
| 03D11 | 98.21048 | Camptothecine (S,+) | 1H21 |
| 03E11 | 44.56844 | Fenbendazole | 1J21 |
| 03F08 | 61.00553 | Mebendazole | 1L15 |
| 03H06 | 48.89376 | Carmofur | 1P11 |
| 04A08 | 85.94884 | Albendazole | 1B16 |
| 04E04 | 43.04360 | Homochlorcyclizine dihydrochloride | 1J8 |
| 04G07 | 42.65304 | Clemastine fumarate | 1N14 |
| 04G09 | 41.53369 | Pimozide | 1N18 |
| 04G10 | 57.62188 | Amodiaquin dihydrochloride dihydrate | 1N20 |
| 04H04 | 99.22016 | Trifluoperazine dihydrochloride | 1P8 |
| 04H09 | 134.31551 | Quinacrine dihydrochloride hydrate | 1P18 |
| 04H10 | 58.45916 | Clofilium tosylate | 1P20 |
| 04H11 | 104.41356 | Fluphenazine dihydrochloride | 1P22 |
| 05D06 | 59.93876 | Hycanthone | 2G11 |
| 05E04 | 79.95940 | Colchicine | 2I7 |
| 05F04 | 129.94272 | Amethopterin (R,S) | 2K7 |
| 05G06 | 128.57984 | Mitoxantrone dihydrochloride | 2M11 |
| 05G07 | 59.58194 | GBR 12909 dihydrochloride | 2M13 |
| 05H07 | 69.02289 | Etoposide | 2O13 |
| 05H08 | 130.19262 | Clomiphene citrate (Z,E) | 2O15 |
| 05H10 | 96.62952 | Prochlorperazine dimaleate | 2O19 |
| 06A03 | 40.59665 | Haloprogin | 2A6 |
| 06A04 | 42.78559 | Thyroxine (L) | 2A8 |
| 06A05 | 68.89656 | Idebenone | 2A10 |
| 06A10 | 92.01926 | Amiodarone hydrochloride | 2A20 |
| 06A11 | 47.09848 | Amphotericin B | 2A22 |
| 06D09 | 154.20388 | Doxorubicin hydrochloride | 2G18 |
| 06E09 | 63.34866 | Pitavastatin calcium | 2I18 |
| 06H09 | 100.74517 | Felodipine | 2O18 |
| 07A08 | 123.20350 | Daunorubicin hydrochloride | 2B15 |
| 07D07 | 53.13669 | Lovastatin | 2H13 |
| 07E03 | 110.02653 | Thiostrepton | 2J5 |
| 07G02 | 121.45517 | Ciclopirox ethanolamine | 2N3 |
| 08A08 | 44.09169 | Itraconazole | 2B16 |
| 08C10 | 44.31754 | Azelastine hydrochloride | 2F20 |
| 08E09 | 44.19885 | Oxibendazol | 2J18 |
| 08G06 | 99.49913 | Cytarabine | 2N12 |
| 08H11 | 119.89032 | Sertindole | 2P22 |
| 09A03 | 70.77601 | Gefitinib | 3A5 |
| 09A07 | 42.29514 | Oxiconazole Nitrate | 3A13 |
| 09C07 | 61.34671 | Nisoldipine | 3E13 |
| 09G06 | 42.23501 | Methyl benzethonium chloride | 3M11 |
| 09G09 | 44.34410 | Benzethonium chloride | 3M17 |
| 10A06 | 96.56952 | Deferoxamine mesylate | 3A12 |
| 10C11 | 81.24407 | Dronedarone hydrochloride | 3E22 |
| 10F08 | 97.31002 | Alexidine dihydrochloride | 3K16 |
| 10G03 | 89.63361 | Podophyllotoxin | 3M6 |
| 10G04 | 42.29072 | Clofibric acid | 3M8 |
| 10G11 | 105.65904 | Cycloheximide | 3M22 |

*(continued)*

| PlateNumber_PositionNumber_96 | norm | Chemical name | PlateNumber_PositionNumber_384 |
|---|---|---|---|
| 10H04 | 40.08951 | Cloperastine hydrochloride | 3O8 |
| 11B02 | 129.35233 | Auranofin | 3D3 |
| 11F10 | 93.90875 | Fluvastatin sodium salt | 3L19 |
| 11G03 | 80.43842 | Raloxifene hydrochloride | 3N5 |
| 11G06 | 66.24727 | Simvastatin | 3N11 |
| 12A07 | 45.51637 | Clioquinol | 3B14 |
| 12B03 | 58.81574 | Flubendazol | 3D6 |
| 12D03 | 68.46277 | Monobenzone | 3H6 |
| 12E05 | 77.65903 | Posaconazole | 3J10 |
| 12E06 | 73.54838 | Thonzonium bromide | 3J12 |
| 13A06 | 96.13794 | Floxuridine | 4A11 |
| 13A10 | 44.27796 | Darifenacin hydrobromide | 4A19 |
| 13D09 | 75.18107 | Zuclopenthixol dihydrochloride | 4G17 |
| 13F04 | 57.18233 | Deptropine citrate | 4K7 |
| 13H11 | 116.74096 | Pyrvinium pamoate | 4O21 |
| 14A09 | 41.87782 | Piperacetazine | 4A18 |
| 14B07 | 79.25921 | Trifluridine | 4C14 |
| 14C09 | 66.01507 | Thiethylperazine dimalate | 4E18 |
| 14C11 | 111.77553 | Vorinostat | 4E22 |
| 14D11 | 52.48567 | Methiazole | 4G22 |
| 14G11 | 65.16242 | Parbendazole | 4M22 |
| 15C09 | 107.17486 | Cladribine | 4F17 |
| 15D02 | 87.58467 | 5-fluorouracil | 4H3 |
| 15F02 | 90.53441 | Topotecan | 4L3 |
| 15F07 | 51.63017 | Benztropine mesylate | 4L13 |
| 15G04 | 106.89783 | Gemcitabine | 4N7 |
| 15G08 | 49.97765 | Docetaxel | 4N15 |
| 15H02 | 58.32030 | Imatinib | 4P3 |
| 15H06 | 49.42534 | Pravastatin | 4P11 |
| 15H07 | 126.95766 | Rosiglitazone Hydrochloride | 4P13 |
| 15H08 | 127.49423 | Rivastigmine | 4P15 |
| 16B05 | 129.08823 | Tegaserod maleate | 4D10 |
| 16B11 | 50.36939 | Estramustine | 4D22 |
| 16D02 | 102.99171 | Aminacrine | 4H4 |
| 16G02 | 113.22091 | Epirubicin hydrochloride | 4N4 |
| 16H03 | 102.40564 | Pemetrexed disodium | 4P6 |
| 16H04 | 106.98468 | Raltitrexed | 4P8 |

#### Common Hit list for the normalized percent inhibition

| PlateNumber__PositionNumber__96 | Chemical name | PlateNumber__PositionNumber__384 |
|---|---|---|
| 01D05 | Triamterene | 1G9 |
| 01D08 | Pyrimethamine | 1G15 |
| 01D11 | Niclosamide | 1G21 |
| 01H07 | Dibucaine | 1O13 |
| 01H09 | Thioridazine hydrochloride | 1O17 |
| 02B11 | Nocodazole | 1C22 |
| 02E06 | Perphenazine | 1I12 |
| 02F07 | Astemizole | 1K14 |
| 02F09 | Terfenadine | 1K18 |
| 02G04 | Chlorhexidine | 1M8 |
| 02H03 | Chloroxine | 1O6 |
| 02H06 | Paclitaxel | 1O12 |
| 03D11 | Camptothecine (S,+) | 1H21 |
| 03E11 | Fenbendazole | 1J21 |
| 03F08 | Mebendazole | 1L15 |
| 03H06 | Carmofur | 1P11 |
| 04A08 | Albendazole | 1B16 |
| 04E04 | Homochlorcyclizine dihydrochloride | 1J8 |
| 04G07 | Clemastine fumarate | 1N14 |
| 04G09 | Pimozide | 1N18 |
| 04G10 | Amodiaquin dihydrochloride dihydrate | 1N20 |
| 04H04 | Trifluoperazine dihydrochloride | 1P8 |
| 04H09 | Quinacrine dihydrochloride hydrate | 1P18 |
| 04H10 | Clofilium tosylate | 1P20 |
| 04H11 | Fluphenazine dihydrochloride | 1P22 |
| 05D06 | Hycanthone | 2G11 |
| 05E04 | Colchicine | 2I7 |
| 05F04 | Amethopterin (R,S) | 2K7 |
| 05G06 | Mitoxantrone dihydrochloride | 2M11 |
| 05G07 | GBR 12909 dihydrochloride | 2M13 |
| 05H07 | Etoposide | 2O13 |
| 05H08 | Clomiphene citrate (Z,E) | 2O15 |
| 05H10 | Prochlorperazine dimaleate | 2O19 |
| 06A10 | Amiodarone hydrochloride | 2A20 |
| 06D09 | Doxorubicin hydrochloride | 2G18 |
| 06E09 | Pitavastatin calcium | 2I18 |
| 06H09 | Felodipine | 2O18 |
| 07A08 | Daunorubicin hydrochloride | 2B15 |
| 07D07 | Lovastatin | 2H13 |
| 07E03 | Thiostrepton | 2J5 |
| 07G02 | Ciclopirox ethanolamine | 2N3 |
| 08E09 | Oxibendazol | 2J18 |
| 08G06 | Cytarabine | 2N12 |
| 08H11 | Sertindole | 2P22 |
| 09A03 | Gefitinib | 3A5 |
| 09G06 | Methyl benzethonium chloride | 3M11 |
| 09G09 | Benzethonium chloride | 3M17 |
| 10A06 | Deferoxamine mesylate | 3A12 |
| 10C11 | Dronedarone hydrochloride | 3E22 |
| 10F08 | Alexidine dihydrochloride | 3K16 |
| 10G03 | Podophyllotoxin | 3M6 |
| 10G11 | Cycloheximide | 3M22 |
| 11B02 | Auranofin | 3D3 |
| 11F10 | Fluvastatin sodium salt | 3L19 |
| 11G03 | Raloxifene hydrochloride | 3N5 |
| 11G06 | Simvastatin | 3N11 |
| 12B03 | Flubendazol | 3D6 |
| 12E05 | Posaconazole | 3J10 |
| 12E06 | Thonzonium bromide | 3J12 |
| 13A06 | Floxuridine | 4A11 |

*(continued)*

| PlateNumber__PositionNumber__96 | Chemical name | PlateNumber__PositionNumber__384 |
|---|---|---|
| 13D09 | Zuclopenthixol dihydrochloride | 4G17 |
| 13F04 | Deptropine citrate | 4K7 |
| 13H11 | Pyrvinium pamoate | 4O21 |
| 14A09 | Piperacetazine | 4A18 |
| 14B07 | Trifluridine | 4C14 |
| 14C09 | Thiethylperazine dimalate | 4E18 |
| 14C11 | Vorinostat | 4E22 |
| 14D11 | Methiazole | 4G22 |
| 14G11 | Parbendazole | 4M22 |
| 15C09 | Cladribine | 4F17 |
| 15D02 | 5-fluorouracil | 4H3 |
| 15F02 | Topotecan | 4L3 |
| 15F07 | Benztropine mesylate | 4L13 |
| 15G04 | Gemcitabine | 4N7 |
| 15G08 | Docetaxel | 4N15 |
| 15H02 | Imatinib | 4P3 |
| 16B05 | Tegaserod maleate | 4D10 |
| 16B11 | Estramustine | 4D22 |
| 16D02 | Aminacrine | 4H4 |
| 16G02 | Epirubicin hydrochloride | 4N4 |
| 16H03 | Pemetrexed disodium | 4P6 |
| 16H04 | Raltitrexed | 4P8 |
